# Malignancy and NF-kB signalling strengthen coordination between the expression of mitochondrial and nuclear-encoded oxidative phosphorylation genes

**DOI:** 10.1101/2021.06.30.450588

**Authors:** Marcos Francisco Perez, Peter Sarkies

## Abstract

Mitochondria are ancient endosymbiotic organelles crucial to eukaryotic growth and metabolism. Mammalian mitochondria carry a small genome containing thirteen protein-coding genes with the remaining mitochondrial proteins encoded by the nuclear genome. Little is known about how coordination between the two sets of genes is achieved. Correlation analysis of RNA-seq expression data from large publicly-available datasets is a common method to leverage genetic diversity to infer gene co-expression modules. Here we use this method to investigate nuclear-mitochondrial gene expression coordination. We identify a pitfall in correlation analysis that results from the large variation in the proportion of transcripts from the mitochondrial genome in RNA-seq data. Commonly used normalization techniques based on total read count (such as FPKM or TPM) produce artefactual negative correlations between mitochondrial- and nuclear-encoded transcripts. This also results in artefactual correlations between pairs of nuclear-encoded genes, thus having important consequences for inferring co-expression modules beyond mitochondria. We show that these effects can be overcome by normalizing using the median-ratio normalization (MRN) or trimmed mean of M values (TMM) methods. Using these normalizations, we find only weak and inconsistent correlations between mitochondrial and nuclear-encoded mitochondrial genes in the majority of healthy human tissues from the GTEx database. However, a subset of healthy tissues with high expression of NF-κB show significant coordination supporting a role for NF-κB in retrograde signalling. Contrastingly, most cancer types show robust coordination of nuclear and mitochondrial OXPHOS gene expression, identifying this as a feature of gene regulation in cancer.

## Introduction

Human mtDNA contains 24 genes encoding for ribosomal and transfer RNAs and 13 protein-coding genes, all of which are involved in oxidative phosphorylation (hereafter mtOXPHOS genes). Mammalian mitochondrial genome-encoded RNAs (mtRNAs) are polyadenylated (Taanman 1999) and are thus robustly represented in polyA+ selected RNA-seq libraries, comprising a large fraction of total reads in many human tissues (Mercer, Neph et al. 2011). However, the majority of proteins with mitochondrial localisation are encoded by the nuclear genome, including over 100 genes involved in oxidative phosphorylation (nuOXPHOS genes) in humans.

As the mtOXPHOS and nuOXPHOS genes are obligate partners in catalysis within protein complexes with a defined stoichiometry, one might expect their expression to be coordinated in order to maximise cell growth and function. This would require mitochondrial-to-nuclear communication, known as ‘retrograde signalling’. While retrograde signalling has been characterised in the case of overt mitochondrial function dysfunction or depletion in yeast (Parikh, Morgan et al. 1987), *C. elegans* (Haynes, Petrova et al. 2007), *Drosophila* (Owusu-Ansah, Song et al. 2013) and mammals (Zhao, Wang et al. 2002, Münch and Harper 2016), little is known about whether retrograde signalling functions to co-ordinate mtOXPHOS and nuOXPHOS gene expression under normal physiological conditions.

One possible method to investigate the coordination between mtRNA and nuclear mitochondrial gene expression is to examine the correlation between mtOXPHOS and nuOXPHOS expression across different individuals to investigate whether natural differences in mtOXPHOS levels are linked to differences in nuOXPHOS expression. Large datasets such as GTEx and TCGA are ideal for this purpose, enabling examination of normal and diseased tissues respectively. Previous approaches utilising this method to examine GTEx data have suggested that there is a weak positive correlation on average between mtOXPHOS and mtOXPHOS expression, indicating coregulation. Surprisingly, however, many tissues were found to have strongly negative correlations (Barshad, Blumberg et al. 2018). This question has yet to be directly investigated using TCGA data, but expression of nuclear-encoded mitochondrial genes was shown to correlate, weakly, with predicted mtDNA copy number (Reznik, Miller et al. 2016).

An important consideration in use of gene expression datasets across a number of different samples drawn from different individuals is how to normalize the data to enable comparison across genes and samples. Currently, the dominant technology for high-throughput quantification of gene expression is sequencing RNA (RNA-seq) (Wang, Gerstein et al. 2009). The expression level of a gene is indicated by the number of independent sequencing reads mapping to the gene, known as the read count (Mortazavi, Williams et al. 2008). However, raw read counts are not comparable for genes within a sample, as longer transcripts accumulate more mapped reads. Raw counts are also not comparable for a given gene across samples, due to potential differences in sequencing depth/library size and, crucially, library composition.

A common normalization method is RPKM (Reads Per Kilobase of transcript, per Million; also known as FPKM for Fragments Per Kilobase of transcript, per Million) for single- or paired-end sequencing, respectively (Mortazavi, Williams et al. 2008). This method normalizes read counts by library size and gene length. However, the average FPKM still varies from sample to sample, leading to the introduction of the similar TPM (transcripts per million) normalization (Li, Ruotti et al. 2010). The sum of TPM values across samples is invariant, and thus TPM is argued to be a better normalization for comparison of expression levels across samples (Li, Ruotti et al. 2010, Zhao, Ye et al. 2020). The Genotype Expression Project GTEx) provides data as TPM, and the Cancer Genome Atlas (TCGA) provides data as FPKM. Many researchers use data normalized in this way for correlation analysis (Barshad, Blumberg et al. 2018, Galai, Ben-David et al. 2020, Sakharkar, Kaur Dhillon et al. 2020, Yang, Zhang et al. 2020, Boardman, Migally et al. 2021, Marquardt, Solimando et al. 2021), sometimes aided by web-based database interrogation tools (Tang, Li et al. 2017). However, other normalization methods have been proposed which aim to account for biases in library composition, in addition to sequencing depth. The median ratio normalization (MRN; also known as relative log expression – RLE) provides scaling factors for normalizing library read counts in a way that controls for library size and is insensitive to the presence of a minority of highly expressed transcripts, thus controlling for library composition biases across samples (Anders and Huber 2010). The trimmed mean of M-values (TMM) is another algorithm which performs similarly (Robinson and Oshlack 2010). An alternative normalization, the upper quartile (UQ) method, adjusts gene read counts by the library 75^th^ percentile of non-zero read counts rather than total read count, so as to exclude the influence of a minority of highly expressed genes (Bullard, Purdom et al. 2010). The MRN and TMM normalization methods are employed in differential expression analysis by the commonly used *R* packages *DESeq2* and *edgeR*, respectively, in which they are used in statistical models.

In this study we set out to investigate the relationship between mtOXPHOS and nuOXPHOS expression using GTEx and TCGA data. We discover that the apparent relationship depends strongly on the choice of normalization method used for RNA-seq data. We show that this is due to the fact that mtRNA levels, including mtOXPHOS genes, not only often make up more than 50% of the total sequenced transcripts in tissue or cancer types but also show a very large range of variation across samples within a cohort. We show that this leads to artefactually negative correlations between mtOXPHOS and nuOXPHOS expression across samples within a cohort when using TPM or FPKM normalizations. Importantly, we also show that the effect of mtRNA expression variation between tissue samples leads to artefactual positive correlations between TPM values for random nuclear-encoded genes unrelated to the mitochondria.

Normalization using methods that correct for library composition bias, such as MRN or TMM, removes these spurious correlations. This analysis shows that in most tissue types, there is little correlation between mtOXPHOS and nuOXPHOS gene expression across individuals. Nevertheless, there are some tissues showing stronger coordination, and we show that this is linked to higher expression of the NF-κB signalling pathway. Contrastingly, we show a consistent positive correlation between mtOXPHOS and nuOXPHOS expression in many cancers. Interestingly, further, we show that, whilst mostly positive, coordination decreases in cancers with the highest proliferation rates. Our work highlights the dangers in performing correlation analysis across GTEx and TCGA data to identify coregulated genes and evaluates methods to solve these problems, thereby providing new more robust insights into the fundamental problem of how to balance mitochondrial gene regulation across two distinct genomes.

## Results

### mtOXPHOS-nuOXPHOS gene expression appears anticorrelated using TPM normalization

To examine the expression correlation between the mtOXPHOS and nuOXPHOS genes, we downloaded data for 54 human tissues from the GTEx website, normalized to TPM (version 8). We excluded tissues with fewer than 100 samples, leaving 48 tissues remaining. We then applied a linear model to each gene in each tissue to regress out the influence of known confounders (age bracket, sex, cause of death, sequencing batch and ischemic time (Ferreira, Muñoz-Aguirre et al. 2018)), continuing downstream analysis with the resulting residuals. We downsampled to 100 samples per tissue to ensure equal representation across tissues. To ensure that the correlations reflect differences among donors and not expression levels between tissues, we ranked the samples for each gene within each tissue, to reflect the relative level of expression within the tissue-specific randomly-sampled cohort of each gene within a sample. Finally, we determined the correlation of the ranks of all pairs of genes using Spearman’s rank-correlation coefficient across the 4800 chosen samples. We repeated this process 100 times and plotted the median value of each correlation coefficient.

Using this approach, expression of mtOXPHOS genes showed a high degree of positive correlation with each other (ρ = 0.617), as did nuOXPHOS genes (ρ = 0.494). However, mtOXPHOS and nuOXPHOS expression appeared to be strongly anticorrelated (Fig 1a; ρ = - 0.336). As nuclear and mitochondrial gene expression have been reported to correlate positively across tissues (Mercer, Neph et al. 2011), this result was unexpected and so we set out to investigate the source of the apparent anticorrelation in more depth.

**Figure 1.**
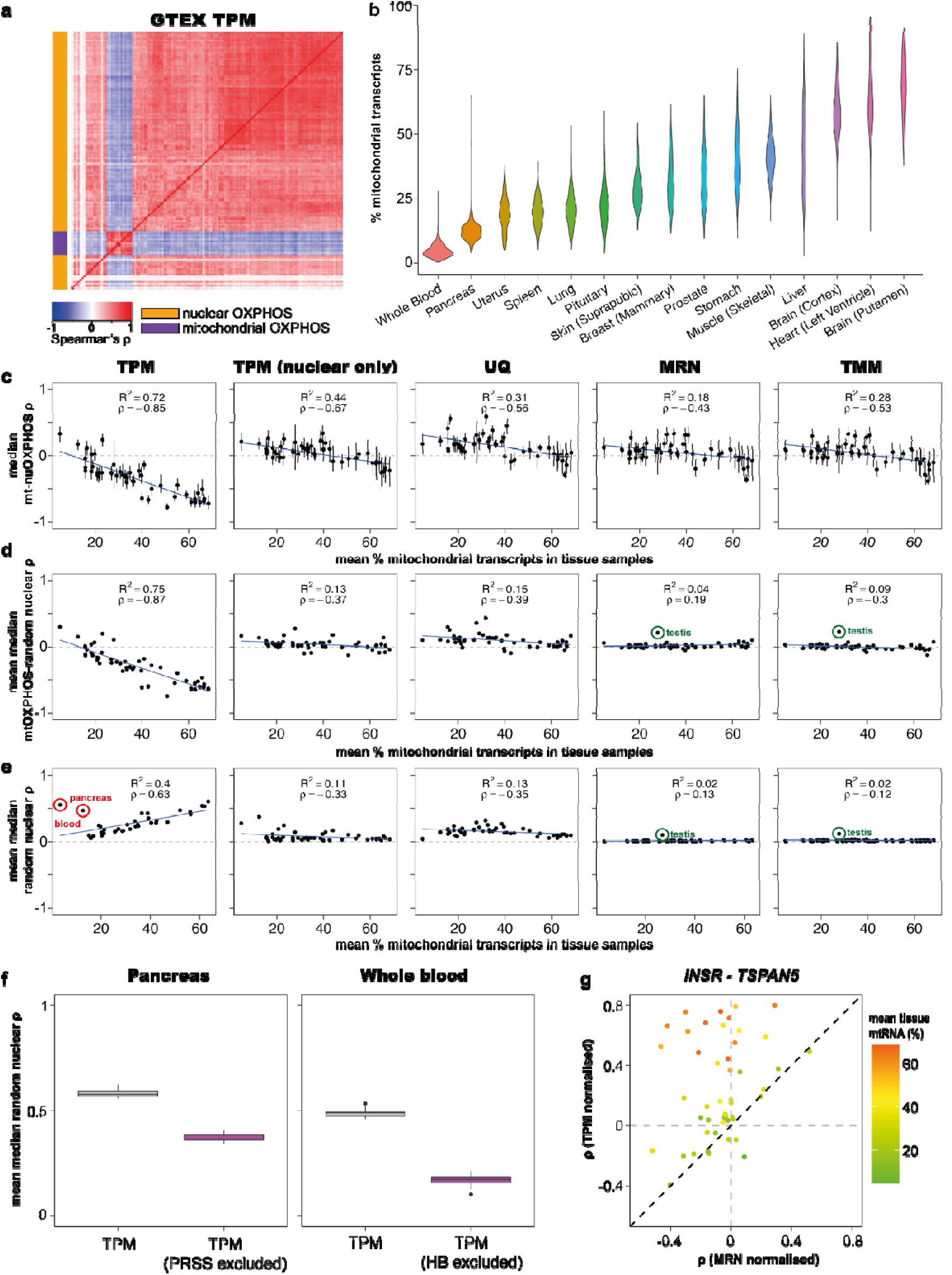
High mitochondrial reads in GTEx libraries leads to artefactual correlations between genes using TPM normalization, obscuring biological relationships. **a)** Heatmap showing median Spearman’s correlation for mtOXPHOS and nuOXPHOS gene expression using TPM normalization in 48 GTEx tissues combined (100 samples from each tissue, sampled 100 times). **b)** Violin plot showing total expression from mtDNA-encoded genes in samples from a selection of 15 healthy human tissues from the GTEx database. **c)** Scatterplot shows median Spearman’s correlation (ρ) of mtOXPHOS-nuOXPHOS gene pairs within 48 GTEx tissues vs tissue mean of mtRNA levels as a percentage of total transcripts. Shown for data normalized by TPM, TPM excluding mitochondrial reads for normalizing nuclear genes, UQ, MRN and TMM methods. Error bars indicate IQR. Blue line shows linear regression with R^2^ and ρ noted within panel. **d)** Scatterplot shows mean over 100 samples of the tissue median ρ of mtOXPHOS with 126 random expressed nuclear genes vs tissue mean of mtRNA levels as a percentage of total transcripts. Shown for data normalized by TPM, TPM excluding mitochondrial reads for normalizing nuclear genes, UQ, MRN and TMM methods. 95% confidence interval error bars are smaller than the plotted symbols. Blue line shows linear regression with R^2^ and ρ noted within panel. Green circles for MRN and TMM highlight unusually high correlations in the testis. **e)** Scatterplot shows mean over 100 samples of the tissue median ρ within 100 random expressed nuclear genes vs tissue mean of mtRNA levels as a percentage of total transcripts. Shown for data normalized by TPM, TPM excluding mitochondrial reads for normalizing nuclear genes, UQ, MRN and TMM methods. 95% confidence interval error bars are smaller than the plotted symbols. Blue line shows linear regression with R^2^ and ρ noted within panel. Red circles for TPM highlight whole blood and pancreas; green circles for MRN and TMM highlight unusually high correlations in the testis. **f)** Boxplot shows mean values for 10 samples of the median ρ of 100 random expressed nuclear genes with TPM normalization or TPM excluding the read counts for *PRSS1* & *PRSS2* (pancreas) or *HBA1*, *HBA2*, *HBB* and *HBD* (whole blood). **g)** Scatterplot shows Spearman’s ρ within tissues for two genes, *INSR* and *TSPAN5,* for TPM normalized data and MRN normalized data. Colour indicates the tissue mean mtRNA level (% total transcripts). Tissues that fall in the top left quadrant are those in which the negative correlation observed using MRN normalization has switched sign and appears to be positive using TPM normalization.

### Variance in mtRNA expression within tissues drives negative mito-nuclear correlations

The total proportion of transcripts derived from mitochondrial DNA ranges from more than 50% of the total transcripts on average in the heart and most brain tissues to just 4.8% of transcripts in whole blood samples (Fig 1b; Fig S1a for all tissues). Variance was also high within tissues, with mtRNAs in the 861 heart (left ventricle) samples ranging from 12.5% to 95.4% of total transcripts (Fig 1b). As the total of TPM values for each sample must sum to 1 million, a sample with 95.4% mtRNAs leaves fewer than 50,000 transcripts to be shared among the nuclear genome, while a sample with 12.5% mtRNAs has ∼875,000. We reasoned that the variation in total mtRNA proportion between donors might therefore dominate the variance in other genes, affecting correlations between genes. Samples with high mtRNA levels will exhibit lower levels of all nuclear genome-derived transcripts and vice-versa, which would result in a spurious negative correlation between mtRNAs and nuclear-encoded RNAs.

For each tissue we plotted the total percentage of mtRNAs against the median Spearman’s correlation of mtOXPHOS-nuOXPHOS gene pairs, using data for all available samples within each tissue (Fig 1c). We observed that the average mtRNA proportion was anticorrelated to the average correlation between mtOXPHOS and nuOXPHOS genes, such that high mtRNA proportions led to negative correlations between mtOXPHOS and nuOXPHOS (ρ = -0.85, p < 2.2 x 10^-16^).

If this was a result of a technical artefact, we should observe the same pattern when we consider the correlations of mtOXPHOS with randomly selected nuclear genes, instead of the nuOXPHOS genes. Indeed, the median Spearman’s correlation of mtOXPHOS-random nuclear gene pairs also shows a strong negative correlation with total mtRNA percentage (Fig 1d; (ρ = -0.87, p < 2.2 x 10^-16^). Thus, as tissue mtRNA levels climb, their variance between samples increasingly comes to dominate the variance of other genes, leading to artefactual negative correlations.

### mtRNA expression variance drives positive correlations between random nuclear transcripts

If sample mtRNA level is a dominant factor in the variance of nuclear gene TPM values, then we reasoned that this might drive artefactual correlations between random nuclear genes which are functionally unrelated to the mitochondria. As the mean tissue mtRNA level rises, variance in mtRNA levels between samples would become an increasingly dominant factor in the variance of nuclear gene TPMs, which are all artificially depressed or inflated in each sample according to mtRNA level.

Supporting this hypothesis, we find that the average correlation for TPM values of pairs of random expressed nuclear genes is positive and increases with mtRNA transcript levels (Fig 1e). The average correlation of random nuclear genes for a tissue was predicted by not only the tissue mean mtRNA level but also the coefficient of variation (CV) of mtRNA levels (linear model, p(mtRNA levels) = 1.37 x 10^-8,^ p(CV) = 3.46 x 10^-5^, p(mtRNA levels * CV) = 0.0136). Stronger artefactual correlations are therefore produced when analysing tissues with higher and more variable mtRNA levels.

We noticed two outlying tissues with low mtRNA levels and high correlations between random nuclear genes (Fig 1e; red circles) - whole blood and pancreas. We looked at the highest expressing genes in these tissues to see if these correlations might be driven by different library composition biases. In whole blood samples just 4 genes encoding haemoglobin subunits – *HBB*, *HBA2*, *HBA1* and *HBD* – make up 40.8% of transcripts on average. Meanwhile in pancreas samples, there is high expression of many digestive enzymes, with two paralogous genes encoding serine proteases – *PRSS1* and *PRSS2* – comprising an average of 18.4% of transcripts. Removing the read counts for these genes from whole blood and pancreas samples before computing sample sums for TPM normalization leads to large and significant decreases in correlation coefficients between random genes (Fig 1f). This shows that the effect we observe is not specific for mitochondrial transcripts but rather applies generally for any transcripts that tend to comprise a high proportion of the total RNA-seq read count.

The inflation of correlations using TPM normalization is a problem for all tissue datasets, but much more so for those with a high mean mtRNA level (or other strong library composition bias), leading in some cases to gene pairs with true negative correlations switching sign and giving the appearance of being significantly and robustly co-expressed (Fig 1g).

Notably, all these analyses are entirely consistent when using Pearson’s correlations instead of Spearman’s rank correlation coefficient (Fig S1). Similar artefactual correlations between mito-nuclear gene pairs or pairs of random nuclear genes are also found using data subjected to FPKM normalization instead of TPM (Fig S2).

### Normalizing libraries with MRN or TMM scaling factors abrogates correlation artefacts

We sought to determine if alternative normalization methods can mitigate these artefacts during correlation analysis. We tried four alternative normalizations. First, we tried removing the mtRNA reads when computing sample read sums for the purposes of normalizing nuclear gene expression as with TPM (mtRNAs are still normalized by the total sample sums). We also tried UQ normalization, which is already applied to some pre-normalized expression datasets available from TCGA. Lastly, we employed MRN and TMM algorithms, which use different procedures to arrive at a scaling factor for library normalization that is designed to be insensitive to library composition bias.

When computing TPM values for nuclear genes with mitochondrial reads excluded, we find that the strong negative correlation of tissue mitochondrial level with average mtOXPHOS- nuOXPHOS is successfully mitigated (Fig 1c). Indeed, there is no relationship between tissue mtRNA level with average mtOXPHOS genes-random nuclear correlations (Fig 1d) or average correlations between random nuclear genes (Fig 1e). However, in tissues with alternative marked library composition biases such as whole blood or pancreas, average correlations of random nuclear genes are still strongly positive, while they remain at least slightly positive in all tissues. Excluding mitochondrial reads alone therefore fails to account for other library composition biases.

UQ normalization performs similarly to TPM with mitochondrial reads excluded with respect to artefacts driven by high mitochondrial transcript levels (Fig 1c-e) and abrogates the strong positive correlation between random nuclear genes driven by alternative library composition biases in whole blood and pancreas samples (Fig 1e). However, UQ normalization results in a stronger inflation of correlations between random nuclear genes in most tissues (Fig 1e), and therefore cannot be recommended for performing correlation analyses.

MRN and TMM normalizations abrogate the artefactual correlations of mitochondrial genes with random nuclear genes (Fig 1d) and between random nuclear genes (Fig 1e). We conclude that scaling read counts by library scaling factors produced by MRN or TMM algorithms is a simple and appropriate way of normalizing RNA-seq data before correlation analysis. We note that the processed data used for QTL analysis available for download on the GTEx portal are already TMM-normalized (in addition to further processing) and are thus appropriate for use in correlation analysis.

We note that one tissue, the testis (green circles, Fig 1d, e; Fig S1c, d), displays positive correlations between mtOXPHOS genes and random nuclear genes and also between pairs of random nuclear genes when using MRN and TMM normalizations. The distribution of correlations across random nuclear gene pairs differs in width across tissues (Fig S3a); in some tissues more extreme positive and negative correlations are more common, which is apparent when inspecting correlation heatmaps between random nuclear genes (Fig S3b). Nonetheless, the distributions are centred around 0 and are unimodal for all tissues but the testis (Fig S3a), which displayed a significantly bimodal distribution of correlations between gene pairs for 7/100 samples of 100 random nuclear genes (Hartigan’s dip test; FDR < 0.05; no significant samples in any of the 47 other tissues). This unusual property may be related to the unique transcriptional complexity of the testis, with over 80% of protein-coding genes expressed (Soumillon, Necsulea et al. 2013). This was recently suggested to play a role in supressing mutagenesis in the germline (Xia, Yan et al. 2020).

### Correlations of mtOXPHOS-nuOXPHOS genes are weak and inconsistent across tissues

Having established appropriate normalizations for the data, we return to the question of the correlation of the mtOXPHOS and nuOXPHOS genes. We normalized GTEx read counts using MRN or TMM and applied a linear model for each gene in each tissue individually to control for known confounders. It is crucial to account for tissue-specific trends with respect to confounding variables, as to apply a linear model across samples from multiple tissues can generate further severe artefacts. As previously, we sampled 100 samples from each tissue, ranked the residuals for each gene within the tissue sample and aggregated these ranks across tissues before computing the Spearman’s correlation across 4800 samples. We repeated this process 100 times and plotted the median coefficient of the 100 iterations for each gene pair (Fig 2a).

**Figure 2.**
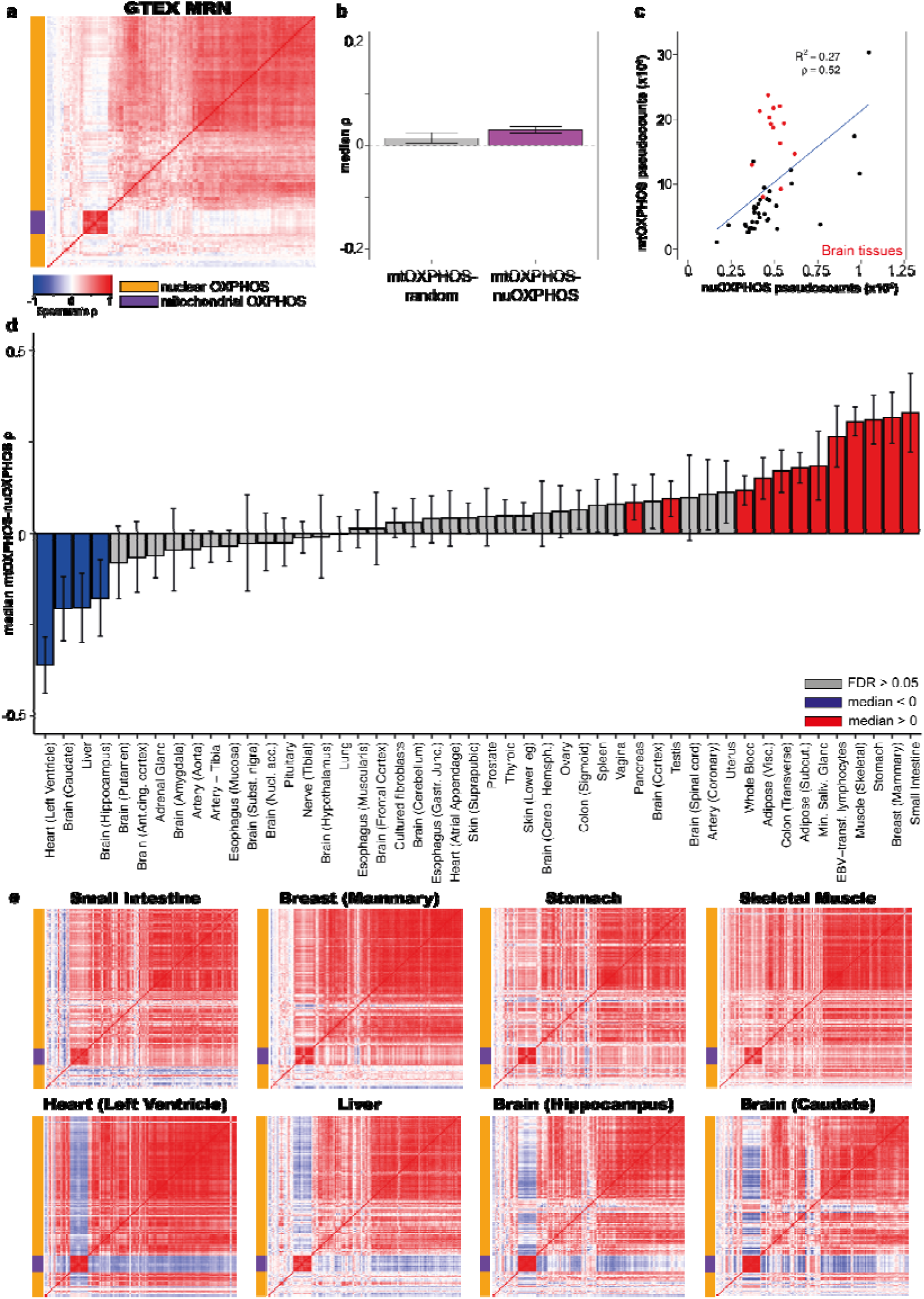
MRN normalization reveals weak and inconsistent correlations between mtOXPHOS and nuOXPHOS within tissues. **a)** Heatmap showing median Spearman’s ρ for mtOXPHOS and nuOXPHOS gene expression using MRN normalization in 48 GTEx tissues combined (100 samples from each tissue, sampled 100 times). **b)** Boxplot showing median Spearman’s ρ for 100 iterations of mtOXPHOS genes with random nuclear genes or mtOXPHOS with nuOXPHOS genes for 48 GTEx tissues combined. **c)** Scatterplot showing mean MRN-normalized pseudocounts for mtOXPHOS and nuOXPHOS genes in 48 GTEx tissues. The 13 brain tissues display a high mtOXPHOS count and are shown in red. **d)** Observed median Spearman’s ρ between mtOXPHOS and nuOXPHOS genes for 48 GTEx tissues. Error bars show 95% bootstrap confidence interval. Blue bars indicate observed correlation significantly lower than 0 (bootstrap empirical p-value, FDR-adjusted < 0.05), red bars indicate observed correlation significantly higher than 0 and grey bars indicate FDR > 0.05. Z-statistics and p-values for observed value relative to mtOXPHOS with random nuclear genes are shown in Table S1. **e)** Heatmap showing Spearman’s ρ for mtOXPHOS and nuOXPHOS genes for 8 GTEx tissues showing clear positive or negative correlations.

When employing MRN or TMM normalizations, we find that the overall correlation between mtOXPHOS and nuOXPHOS genes between donors in the GTEx data is very weak. The mean median Spearman’s correlation for mtOXPHOS-nuOXPHOS gene pairs is only slightly higher than the mean median Spearman’s correlation between mtOXPHOS genes and random nuclear genes (0.0302 vs 0.0144 for MRN normalization; 0.0157 vs -0.00326 for TMM normalization; Fig 2b). Although significant (t-test p = 2.82 x 10^-27^ for MRN; 2.16 x 10^-38^ for TMM), only a tiny fraction of the variation in nuOXPHOS gene expression can be explained by mtOXPHOS gene expression. This suggests that despite broad coordination of mtOXPHOS-nuOXPHOS expression across tissues (Fig 2c)(Mercer, Neph et al. 2011) there is limited coordination that can be detected between the nuclear and mitochondrial gene expression programs within tissues. Notably, this contrasts to very clear coregulation within the mtOXPHOS (ρ = 0.775) and nuOXPHOS (ρ = 0.329) gene sets.

With MRN normalized data, we observed that the relationship between mtOXPHOS and nuOXPHOS gene expression in individual tissues is inconsistent (Fig 2d, Table S1, Supplementary Dataset 1). 32/48 tissues exhibit a median mtOXPHOS-nuOXPHOS correlation that is not significantly different from 0 (FDR > 0.05). 12/48 tissues exhibit a median mtOXPHOS-nuOXPHOS correlation significantly higher from 0, the most striking being the small intestine (terminal ileum), breast (mammary tissue), stomach and skeletal muscle all with median correlations between 0.30 and 0.32 (Fig 2e). 4/48 tissues display a significantly lower correlation with respect to random expectation; the left ventricle of the heart, liver, hippocampus (brain) and caudate (basal ganglia of the brain), with medians - 0.361, -0.204, -0.206 and -0.177 respectively (Fig 2e). Note that although the testis has a positive mtOXPHOS-nuOXPHOS correlation, it is significantly lower than that observed between mtOXPHOS and random nuclear genes (Fig S4a; Table S1), which is unusually high as noted above. Very similar results are obtained starting with a TMM normalization (Fig S5).

### Positive correlation between mtOXPHOS and nuOXPHOS gene expression in cancer

We next explored whether any relationship between mtOXPHOS and nuOXPHOS gene expression exists in cancer samples. We downloaded data from TCGA (harmonized) for 31 cancer types with more than 50 primary tumour samples.

We observed substantial variation in mtRNA across and within cancer types (Fig 3b). As we observed for GTEx data, normalization of TCGA gene expression values using TPM leads to spurious correlations, both between mtOXPHOS-nuOXPHOS genes and random nuclear gene pairs (Fig 3a, c-e). We then proceeded to apply MRN/TMM normalization and repeated the analysis. These normalization methods correct the spurious correlations between mtOXPHOS genes and random nuclear genes and between pairs of random nuclear genes (Fig 3a, d, e).

**Figure 3.**
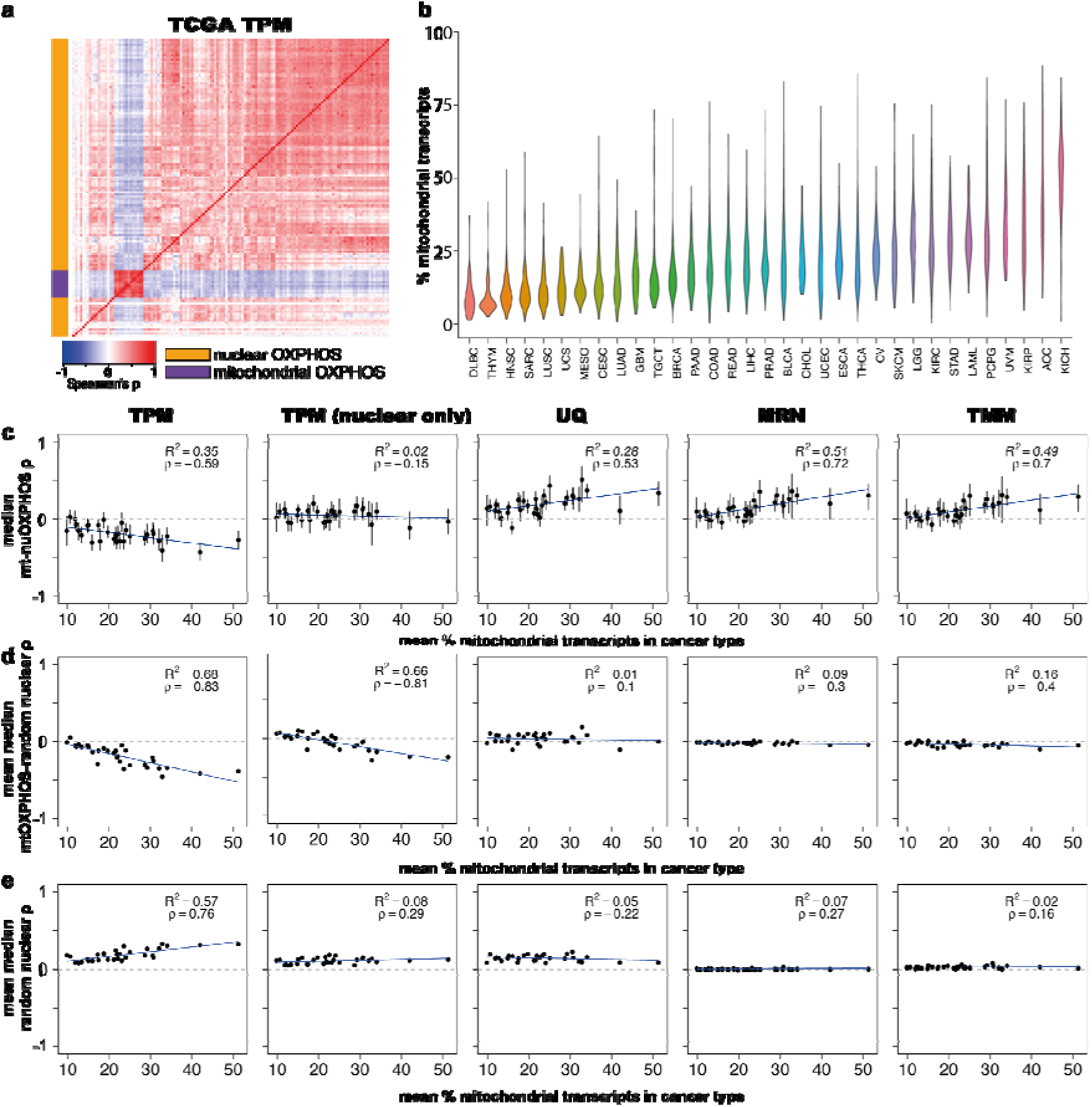
Artefactual correlations are also driven by mtRNA expression in the Cancer Genome Atlas (TCGA) database. **a)** Heatmap showing median Spearman’s correlation for mtOXPHOS and nuOXPHOS gene expression using TPM normalization in 31 TCGA cancer types combined (50 samples from each tissue, sampled 100 times). **b)** Violin plot showing total expression from mtDNA-encoded genes in samples from a selection of 13 cancer types from TCGA database. **c)** Scatterplot shows median Spearman’s correlation (ρ) of mtOXPHOS-nuOXPHOS gene pairs within 31 TCGA cancer types vs cancer type mean of mtRNA expression as a percentage of total transcripts. Shown for data normalized by TPM, TPM excluding mitochondrial reads for normalizing nuclear genes, UQ, MRN and TMM methods. Error bars indicate IQR. Blue line shows linear regression with R^2^ noted within panel. **d)** Scatterplot shows mean over 100 samples of the cancer type median ρ of mtOXPHOS with 126 random expressed nuclear genes vs cancer type mean of mtRNA expression as a percentage of total transcripts. Shown for data normalized by TPM, TPM excluding mitochondrial reads for normalizing nuclear genes, UQ, MRN and TMM methods. 95% confidence interval error bars are smaller than the plotted symbols. Blue line shows linear regression with R^2^ and ρ noted within panel. **e)** Scatterplot shows mean over 100 samples of the cancer type median ρ within 100 random expressed nuclear genes vs cancer type mean of mtRNA expression as a percentage of total transcripts. Shown for data normalized by TPM, TPM excluding mitochondrial reads for normalizing nuclear genes, UQ, MRN and TMM methods. 95% confidence interval error bars are smaller than the plotted symbols. Blue line shows linear regression with R^2^ and ρ noted within panel. Abbreviations for cancer type are given in Table S5.

Nevertheless, with MRN normalization 23 of 31 individual cancer types tested displayed a significantly positive correlation (FDR < 0.05), with none displaying a significant negative correlation (Fig 4c, Table S2; Supplementary Dataset 2). Median correlations of mtOXPHOS gene expression with random nuclear genes was around 0 for all cancer types (Fig S4b). Analysing across cancer types, the mean median Spearman’s correlation for mtOXPHOS-nuOXPHOS genes in 0.110, while for mtOXPHOS with random nuclear genes it is -0.0147 (Fig 4a, b; t-test p-value = 5.53 x 10^-158^). Interestingly, the degree of mtOXPHOS-nuOXPHOS correlation within a cancer type increases with mean mtRNA expression level (Fig 3a), in a reversal of the apparent relationship when using TPM-normalized data.

**Figure 4.**
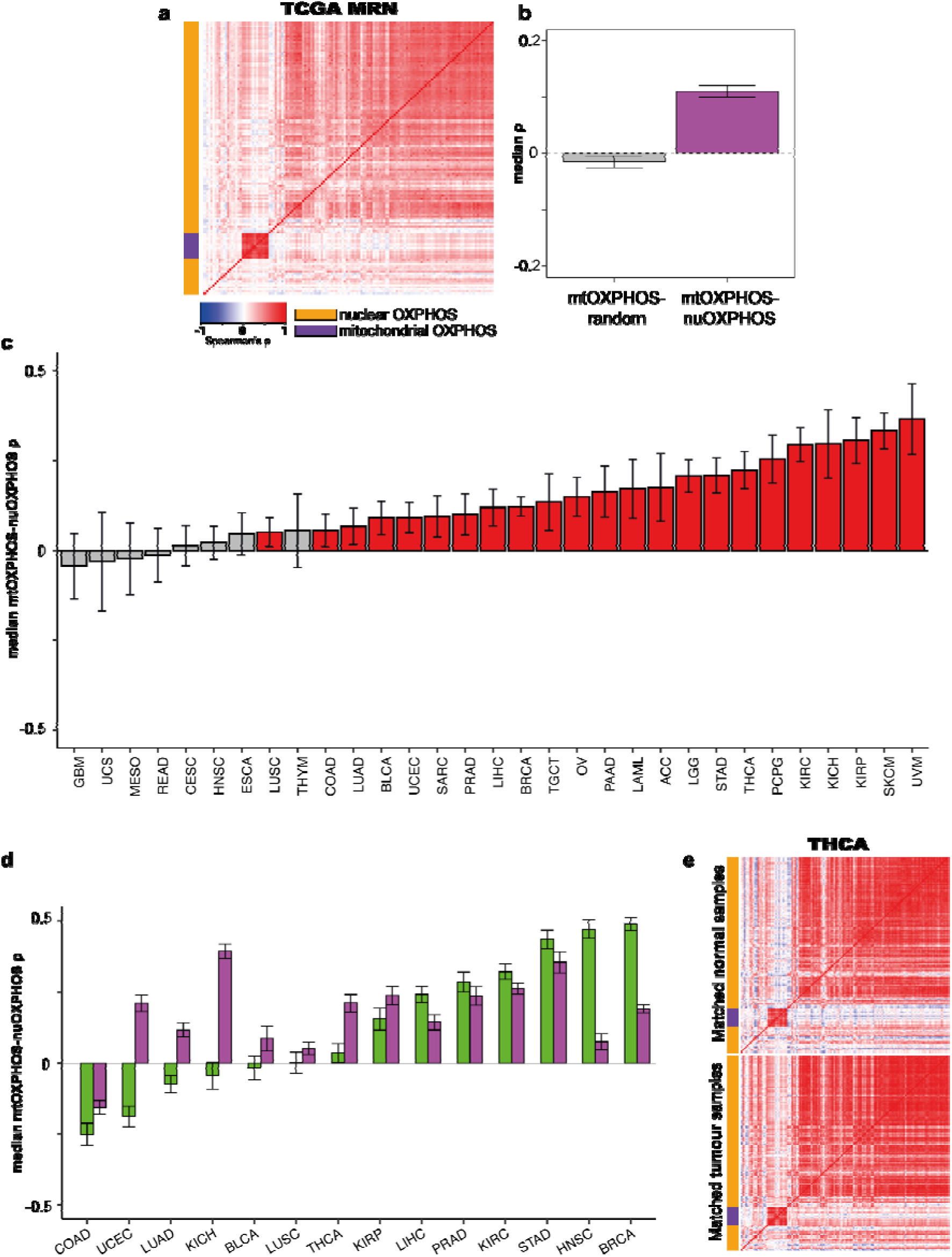
Most cancer types display a positive correlation between mtOXPHOS and nuOXPHOS expression. **a)** Heatmap showing median Spearman’s ρ for mtOXPHOS and nuOXPHOS gene expression using MRN normalization in 31 TCGA cancer types combined (50 samples from each tissue, sampled 100 times). **b)** Boxplot showing median Spearman’s ρ for 100 iterations of mtOXPHOS genes with random nuclear genes or mtOXPHOS with nuOXPHOS genes for 31 TCGA cancer types combined. **c)** Observed median Spearman’s ρ between mtOXPHOS and nuOXPHOS genes for 31 TCGA cancer types. Error bars show bootstrap 95% confidence interval. Blue bar indicates observed correlation significantly lower 0 (bootstrap empirical p-value, FDR-adjusted < 0.05), red bars indicate observed correlation significantly higher than 0 and grey bars indicate FDR > 0.05. Z-statistics and p-values for observed value are shown in Table S2. **d)** Median mtOXPHOS-nuOXPHOS ρ within matched samples of tumour (red) or normal tissue (grey) from TCGA projects. Error bars show bootstrapped standard error. **e)** mtOXPHOS-nuOXPHOS correlation heatmaps for thyroid cancer (TCGA) matched normal tissue samples (above) and tumour samples (below). Abbreviations for cancer types are given in Table S5.

We analysed 14 cancer types for which data for at least 10 normal adjacent tissue samples was available, and identified the matched tumour counterpart samples. Using MRN-normalized data, we then computed the correlations between mtOXPHOS and nuOXPHOS genes within the normal and tumour samples to compare them (Fig 4d). Although direct comparison between the two databases is not possible, we note that the mtOXPHOS-nuOXPHOS correlations we observe for normal samples in TCGA are broadly congruous with those observed in the GTEx tissues, in particular the strong positive correlations in TCGA adjacent normal tissue samples from the stomach (STAD) and breast (BRCA). We find that the mtOXPHOS-nuOXPHOS correlation tends to be positive in diseased samples in those tissues in which the normal cohort displays a negative or insignificant correlation, such as in thyroid cancer (Fig 4e; Supplementary Dataset 3, 4). Overall, 13/14 matched tumour samples display a significantly positive mtOXPHOS-nuOXPHOS correlation, compared to 7/14 of the matched normal tumour samples (p = 0.0329, Fisher’s exact test).

### NF-κB expression is correlated with OXPHOS co-ordination in healthy tissues

To understand what might underlie the variation in mtOXPHOS-nuOXPHOS correlation across tissues, we correlated the mean expression of all genes within each GTEx tissue to the observed tissue OXPHOS correlation. We selected the 1000 genes whose expression across tissues most strongly correlated positively to mtOXPHOS-nuOXPHOS correlation across 48 tissues (the top 1000, Spearman’s ρ > 0.487) and performed a gene ontology and KEGG pathway enrichment analysis; we did the same for the 1000 most anticorrelated genes (the bottom 1000, Spearman’s ρ < -0.443). While the anticorrelated genes yielded no significantly enriched terms, the positively correlated genes were strongly enriched for terms related to immune function and B cell activation (Supplementary Dataset 5, 6). On closer inspection, we found that this was mostly due to a large number (221) of immunoglobulin-encoding genes within the top 1000 genes. These immunoglobulin genes represented all three major immunoglobulin clusters: IGH (107 genes), IGK (50) and IGL (42), as well as IGH (15) and IGK (7) orphons located outside the main clusters.

We used estimates of immune cell fraction for GTEx samples produced by GEDIT, a gene expression deconvolution tool (Nadel, Lopez et al. 2021) to see if this was predictive of nuOXPHOS-mtOXPHOS correlation. Immune cell fraction could explain little of the OXPHOS correlation variance (R^2^ = 0.0199, p < 2.2 x 10^-16^) (Fig S6).

In addition to immune-related terms, there was an enrichment for terms that implied a possible role for cellular proliferation, such as ribosome biogenesis and DNA replication. To assess this contribution to mtOXPHOS-nuOXPHOS correlation, we used the expression data to infer a measure of active proliferation, the Proliferative Index (PI) (Ramaker, Lasseigne et al. 2017), for all GTEx samples. Despite a significant relationship, the PI explained a tiny fraction of the variance across tissues in OXPHOS correlation (R^2^ = 0.00759, p < 2.2 x 10^- 16^)(Fig 5b).

**Figure 5.**
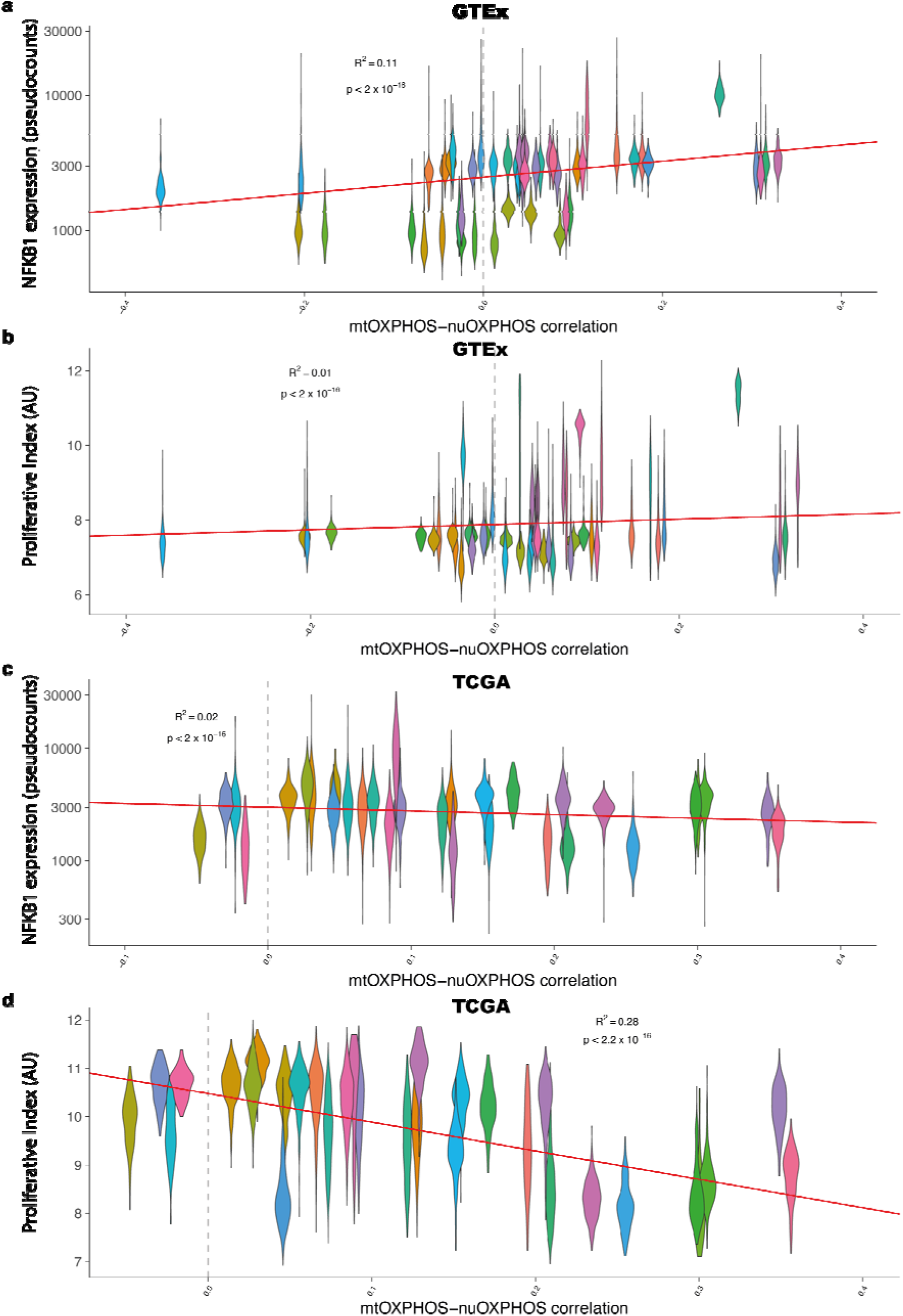
mtOXPHOS-nuOXPHOS co-ordination is correlated to NF-κB expression and cellular proliferation in the GTEx and TCGA respectively. **a)** Within-tissue mtOXPHOS-nuOXPHOS correlation against NFKB1 expression (MRN pseudocounts) for GTEx tissues. Each violin represents the samples within a tissue. **b)** Within-tissue mtOXPHOS-nuOXPHOS correlation against Proliferative Index for GTEx tissues. Each violin represents the samples within a tissue. **c)** Within-tissue mtOXPHOS-nuOXPHOS correlation against NFKB1 expression (MRN pseudocounts) for TCGA cancer types. Each violin represents the samples within a tissue. **d)** Within-tissue mtOXPHOS-nuOXPHOS correlation against Proliferative Index for TCGA cancer types. Each violin represents the samples within a tissue. Dashed grey vertical line shows no correlation. AU, arbitrary units.

Another possibility was that immunoglobulin expression was driven by NF-κB, a protein complex present in almost all cell types that can drive immunoglobulin expression in non-immune cells (Liu, Zheng et al. 2009) and has been implicated in mitochondrial signalling to the nucleus in mammals (Butow and Avadhani 2004) and in mitochondrial gene regulation (Cogswell, Kashatus et al. 2003). 3 of the 5 NF-κB members (NFKB1, REL and RELB) were found in the top 1000 genes. Indeed, together the expression of the 5 NF-κB members explain much of the variance in mtOXPHOS-nuOXPHOS correlation, with NFKB1 expression alone the most important contribution (Fig 5a; Table S3).

### Proliferation is negatively correlated to OXPHOS co-ordination in cancer

Having established the importance of NF-κB expression in OXPHOS coordination in healthy tissues we next tested whether NF-κB could play a similar role in cancer. However, we found only a weak, negative relationship between NFKB1 expression and mtOXPHOS-nuOXPHOS correlation in the TCGA data (R^2^ = 0.0194 p < 2.2 x 10^-16^; Fig 5c).

As for the GTEx data, we correlated the mean expression of all genes within 31 TCGA cancer type to the observed mtOXPHOS-nuOXPHOS correlation, before performing a gene ontology enrichment analysis on the top and bottom 1000 genes (Spearman’s ρ > 0.467 and < -0.502 respectively; Supplementary Dataset 7, 8). Overall, there was no overlap at all between the gene ontology terms that were positively or negatively related to OXPHOS coordination in both GTEx and TCGA databases. For the TCGA, the top 1000 genes yielded only 7 enriched terms, related to the lysosome and Golgi apparatus. On the contrary, the 1000 most negatively correlated genes were associated with 220 significant terms, many associated with replication and mitosis.

In agreement with the enrichment of proliferation-related terms in the bottom 1000 genes, we found a strong negative relationship between proliferative index and coordination (R^2^ = 0.279 p < 2.2 x 10^-16^; Fig 5d; Table S4), suggesting that variation in the rate of proliferation in cancer might explain differences in OXPHOS coordination (see Discussion).

## Discussion

Here we have set out to investigate the coordination between the expression of mitochondrial and nuclear transcripts, in particular the mtOXPHOS genes which comprise all 13 mitochondrially-encoded protein coding genes and their nuclear-encoded catalytic partners, the nuOXPHOS genes. Using tens of thousands of RNA-seq samples across 48 healthy human tissues from the GTEx database and 31 distinct cancer types from TCGA, we examined the correlation of mtOXPHOS and nuOXPHOS expression to identify signs of co-regulation. We find that most healthy tissues in the GTEx database show little correlation between mtOXPHOS and nuOXPHOS expression. In contrast, in TCGA we find that cancers show a clear tendency towards positive correlations between mtOXPHOS and nuOXPHOS expression. Our results have implications for analysis of gene expression coordination as well as the potential for retrograde signalling to balance nuclear and mitochondrial expression of genes with mitochondrial function.

### Avoiding artefacts when analysing correlations between expressed genes

In the course of our investigation, we realised that the use of TPM or FPKM normalizations for RNA-seq data produces grave and ubiquitous artefacts in correlation analysis owing to uncontrolled library composition biases. These artefacts are more severe for tissues or cancers with strong biases, of which the most common is high levels of expression from mitochondrial DNA. Because it allows direct comparison across samples, TPM is often considered to be a superior normalization to FPKM (Li, Ruotti et al. 2010, Zhao, Ye et al. 2020). However, as the total sum of TPM values is invariant, the strength of the artefactual correlations introduced are actually more acute than those observed with FPKM normalization.

It has previously been pointed out that library composition biases make comparison of samples from different tissues problematic when using TPM or FPKM normalization (Robinson and Oshlack 2010, Zhao, Ye et al. 2020) Here we have demonstrated that library composition biases make comparisons and correlations of TPM/FPKM values problematic even when the analysis is limited to samples of the same tissue type.

These results have severe implications for any reported correlation analyses using expression values normalized by TPM or FPKM. As correlation coefficients between all pairs of nuclear genes are artificially inflated in every tissue, these artefacts can obscure the true relationship between genes, with negatively correlated gene pairs appearing to be strongly co-expressed. This also presents difficulties in understanding changes in the interactions of pairs of genes across tissues, as changes may be simply due to different library composition biases in those tissues. Lastly, this adds additional difficulties in interpretation of the already problematic (Lovén, Orlando et al. 2012) but frequently observed practice of plotting TPM values of a favourite gene for each tissue from the GTEx database side-by-side as a qualitative measure of tissue expression level, given that a gene’s mean TPM value for a tissue will in large part be determined by the mean mtRNA level of that tissue.

We demonstrate that scaling libraries using MRN or TMM algorithms is a simple and effective way to account for these biases prior to correlation analysis. We strongly dissuade researchers from using TPM or FPKM for gene expression correlation analysis and recommend that researchers employ alternative scaling normalizations that account for library composition bias before embarking on correlation analyses.

### Coordination between mitochondrial and nuclear gene expression

Our analysis using correct normalization substantially updates our view of how gene expression is coordinated between mitochondrial and nuclear genomes. The lack of strong correlations between mtOXPHOS and nuOXPHOS gene expression in the majority of healthy tissues in the GTEx database implies that these genes are generally not strongly co-ordinated by retrograde signalling pathways. The broad co-ordination afforded by tissue-specific transcriptional programs (Mercer, Neph et al. 2011) is likely sufficient for supporting tissue function under normal physiological conditions. However, we found that the tissue expression level of NF-κB genes is associated with the strength of co-ordination between mtOXPHOS and nuOXPHOS expression, supporting a role for this complex in mito-nuclear communication as proposed previously (Butow and Avadhani 2004). The biological significance of strong negative correlations, such as observed in the left ventricle of the heart and the liver, remains unclear.

It is clear from our analysis that the mechanisms by which apparent co-ordination between the mitochondrial and nuclear OXPHOS genes arises differ sharply between healthy tissues and tumour cells. The positive correlation of mtOXPHOS and nuOXPHOS genes across samples within many cancer types does not necessarily imply that retrograde signalling is active in cancers, as other differences between healthy tissues and tumours could play a role. Both mtOXPHOS and nuOXPHOS gene expression is often altered in cancers (Reznik, Wang et al. 2017). Positive correlations might result from selection within tumours for clones with concordant OXPHOS expression, with this selection becoming stronger with higher mtOXPHOS expression; this might explain the relationship between mtOXPHOS-nuOXPHOS correlation and mtRNA levels among TCGA cancer types.

Intriguingly, however, we found that although cancers tend to exhibit positive mtOXPHOS-nuOXPHOS correlation, the strength of this correlation in a particular cancer type relates strongly to its inferred proliferation status, with stronger proliferation related to lower mtOXPHOS-nuOXPHOS correlation. This may be due to the Warburg effect, in which metabolism in rapidly proliferating tumour cells is reported to shift towards aerobic glycolysis, thereby bypassing the mitochondrial role in respiration (Liberti and Locasale 2016). In the fastest proliferating cancer types, the importance of mitochondrial respiration may therefore be reduced, leading to reduced selection within tumours for concordant OXPHOS expression. Thus although selection between cells within proliferating tumours may act to produce a positive mtOXPHOS-nuOXPHOS correlation in most cancers, this selection may be weakened and increasingly dominated by the Warburg effect in the fastest proliferating cancers.

## Supporting information

Supplemental tables and scripts

---

**Figure S1.**
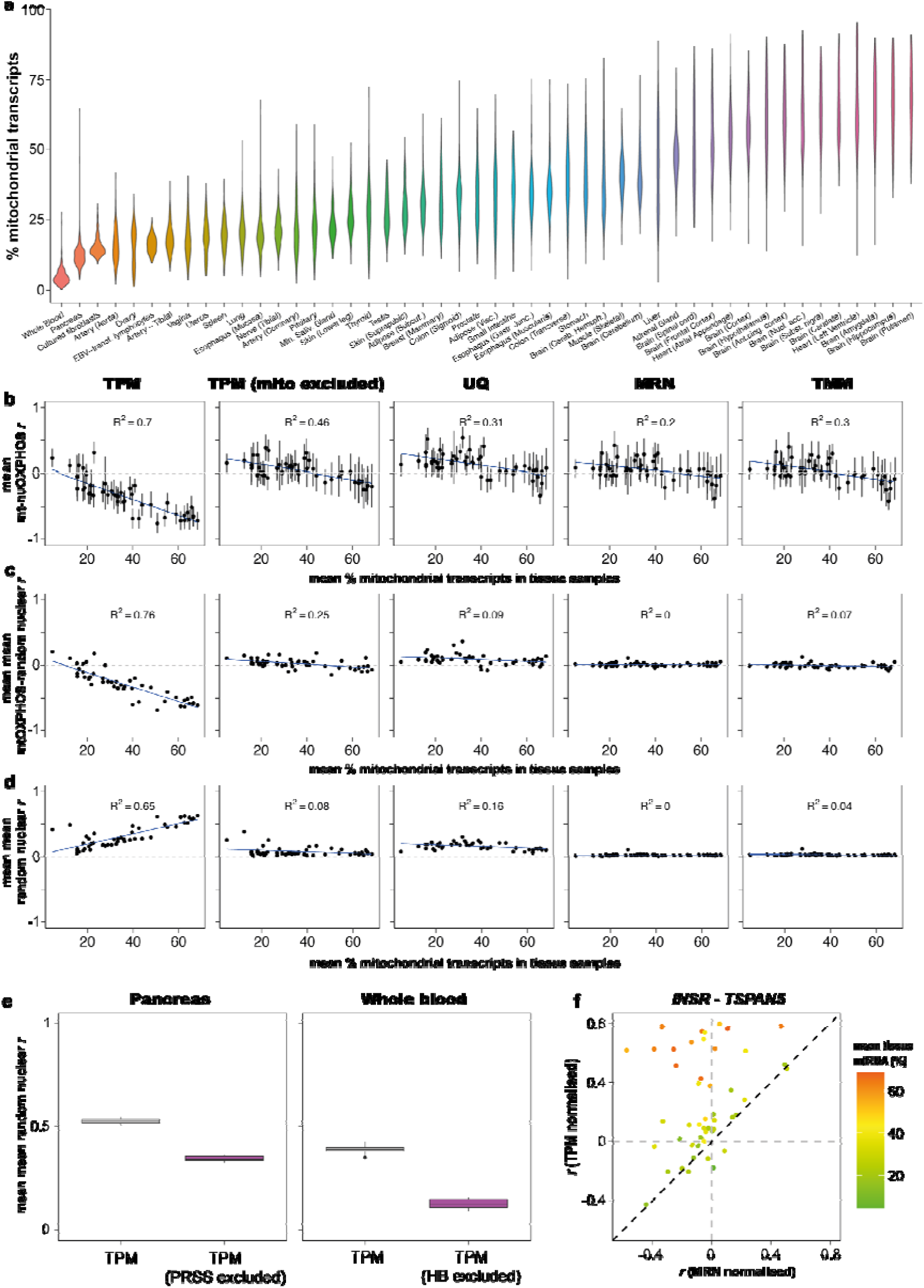
Mitochondrial library bias drive artefactual correlations using Pearson’s correlation. **a)** Violin plot showing total expression from mtDNA-encoded genes in samples from all 48 healthy human tissues with more than 100 samples from the GTEx database. **b)** Scatterplot shows mean Pearson’s correlation (*r*) of mtOXPHOS-nuOXPHOS gene pairs within 48 GTEx tissues vs tissue mean of mtRNA expression as a percentage of total transcripts. Shown for data normalized by TPM, TPM excluding mitochondrial reads for normalizing nuclear genes, UQ, MRN and TMM methods. Error bars indicate SD. Blue line shows linear regression with R^2^ noted within panel. **c)** Scatterplot shows mean over 100 samples of the tissue mean *r* of mtOXPHOS with 126 random expressed nuclear genes vs tissue mean of mtRNA expression as a percentage of total transcripts. Shown for data normalized by TPM, TPM excluding mitochondrial reads for normalizing nuclear genes, UQ, MRN and TMM methods. 95% confidence interval error bars are smaller than the plotted symbols. Blue line shows linear regression with R^2^ noted within panel. Green circles for MRN and TMM highlight unusually high correlations in the testis. **d)** Scatterplot shows mean over 100 samples of the tissue mean *r* within 100 random expressed nuclear genes within tissues vs tissue mean of mtRNA expression as a percentage of total transcripts. Shown for data normalized by TPM, TPM excluding mitochondrial reads for normalizing nuclear genes, UQ, MRN and TMM methods. 95% confidence interval error bars are smaller than the plotted symbols. Blue line shows linear regression with R^2^ noted within panel. Red circles for TPM highlight whole blood and pancreas; green circles for MRN and TMM highlight unusually high correlations in the testis. **e)** Boxplot shows mean values for 10 samples of the mean *r* within 100 random expressed nuclear genes with TPM normalization or TPM excluding the read counts for *PRSS1* & *PRSS2* (pancreas) or *HBA1*, *HBA2*, *HBB* and *HBD* (whole blood). **f)** Scatterplot shows Pearson’s *r* within tissues for two genes, *INSR* and *TSPAN5,* for TPM normalized data and MRN normalized data. Colour indicates the tissue mean mtRNA expression (% total transcripts).

**Figure S2.**
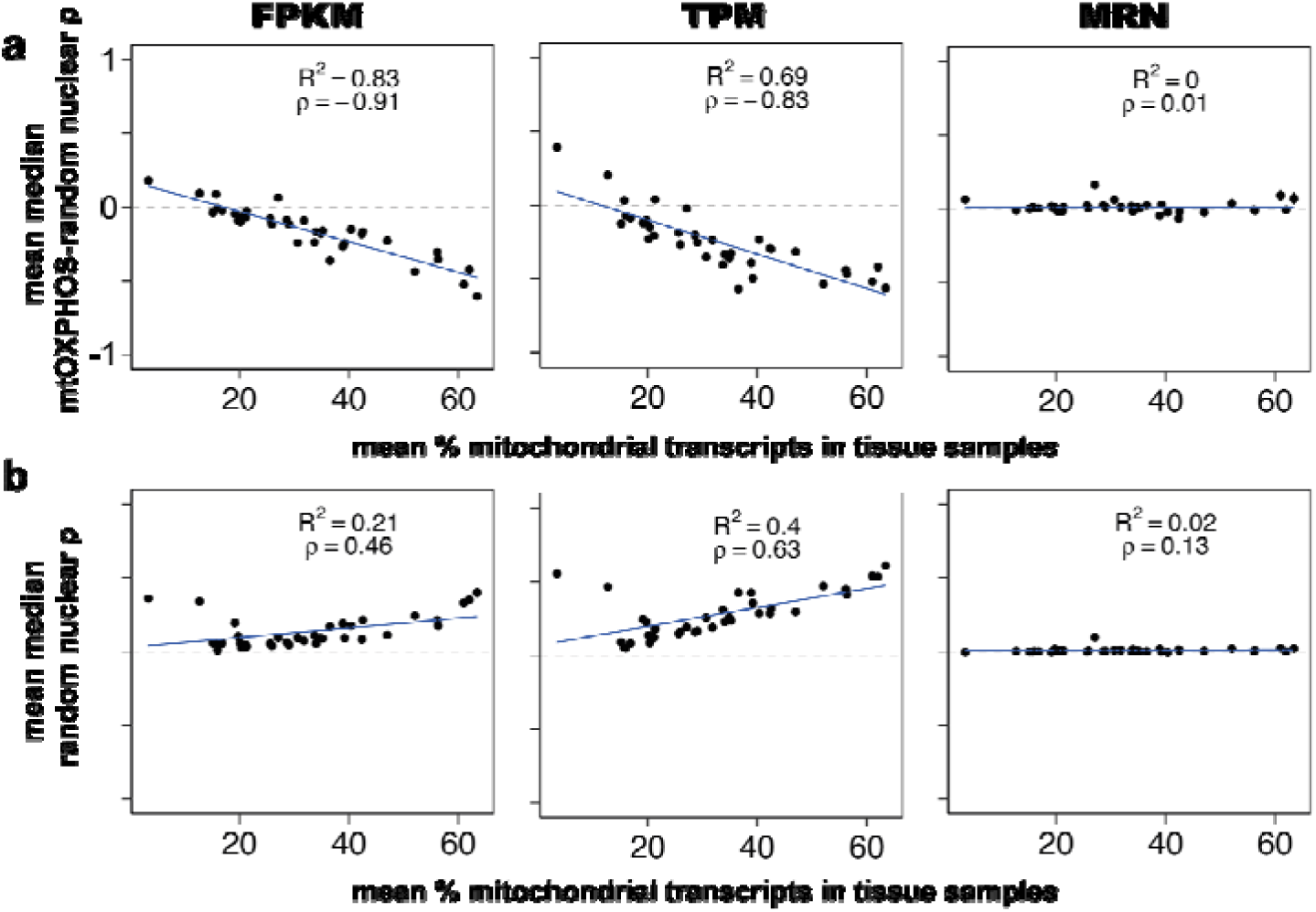
Mitochondrial library bias drive artefactual correlations using FPKM in addition to TPM. **a)** Scatterplot shows mean over 100 samples of the tissue median Spearman’s ρ of mtOXPHOS with 126 random expressed nuclear genes vs tissue mean of mtRNA expression as a percentage of total transcripts. Shown for data normalized by FPKM, TPM, or MRN. 95% confidence interval error bars are smaller than the plotted symbols. Blue line shows linear regression with R^2^ and Spearman’s ρ noted within panel. **b)** Scatterplot shows mean over 100 samples of the tissue median ρ within 100 random expressed nuclear genes vs tissue mean of mtRNA expression as a percentage of total transcripts. Shown for data normalized by TPM, TPM excluding mitochondrial reads for normalizing nuclear genes, UQ, MRN and TMM methods. 95% confidence interval error bars are smaller than the plotted symbols. Blue line shows linear regression with R^2^ noted within panel. Data in this figure are drawn from GTEx v6p, rather than GTEx v8 as for all other figures.

**Fig S3.**
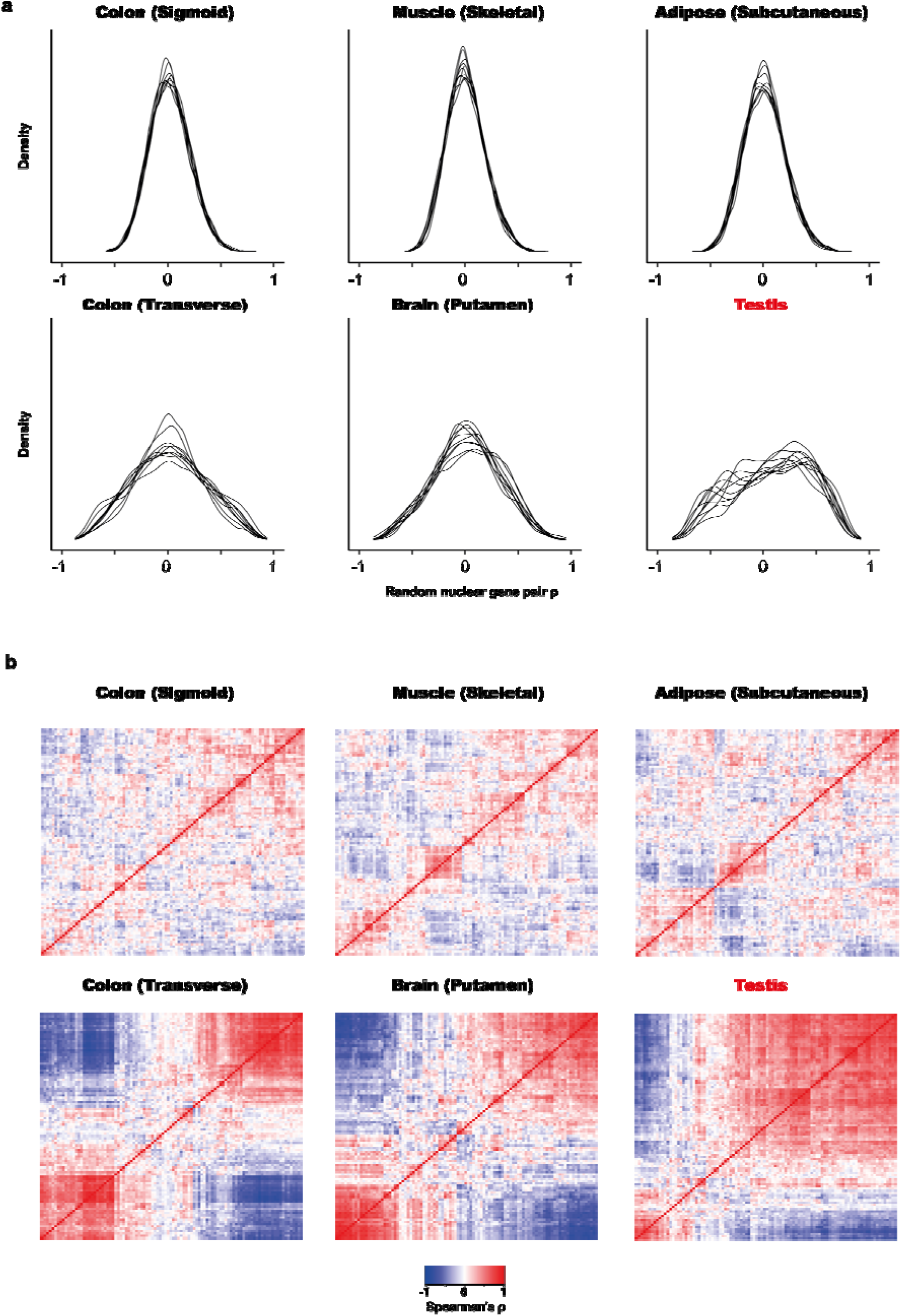
Correlation distributions vary consistently across tissues, with the testis unique in displaying signs of bimodal correlation distribution. **a)** Density of correlation coefficients within 10 random samples (overlaid lines) of 100 random nuclear genes for GTEx tissues with narrow distributions (above) and wide distributions (below). The testis shows clear signs of bimodality. Correlations performed on MRN-normalized data. **b)** Representative heatmaps for Spearman’s correlations of 100 random genes for the same tissues.

**Fig S4.**
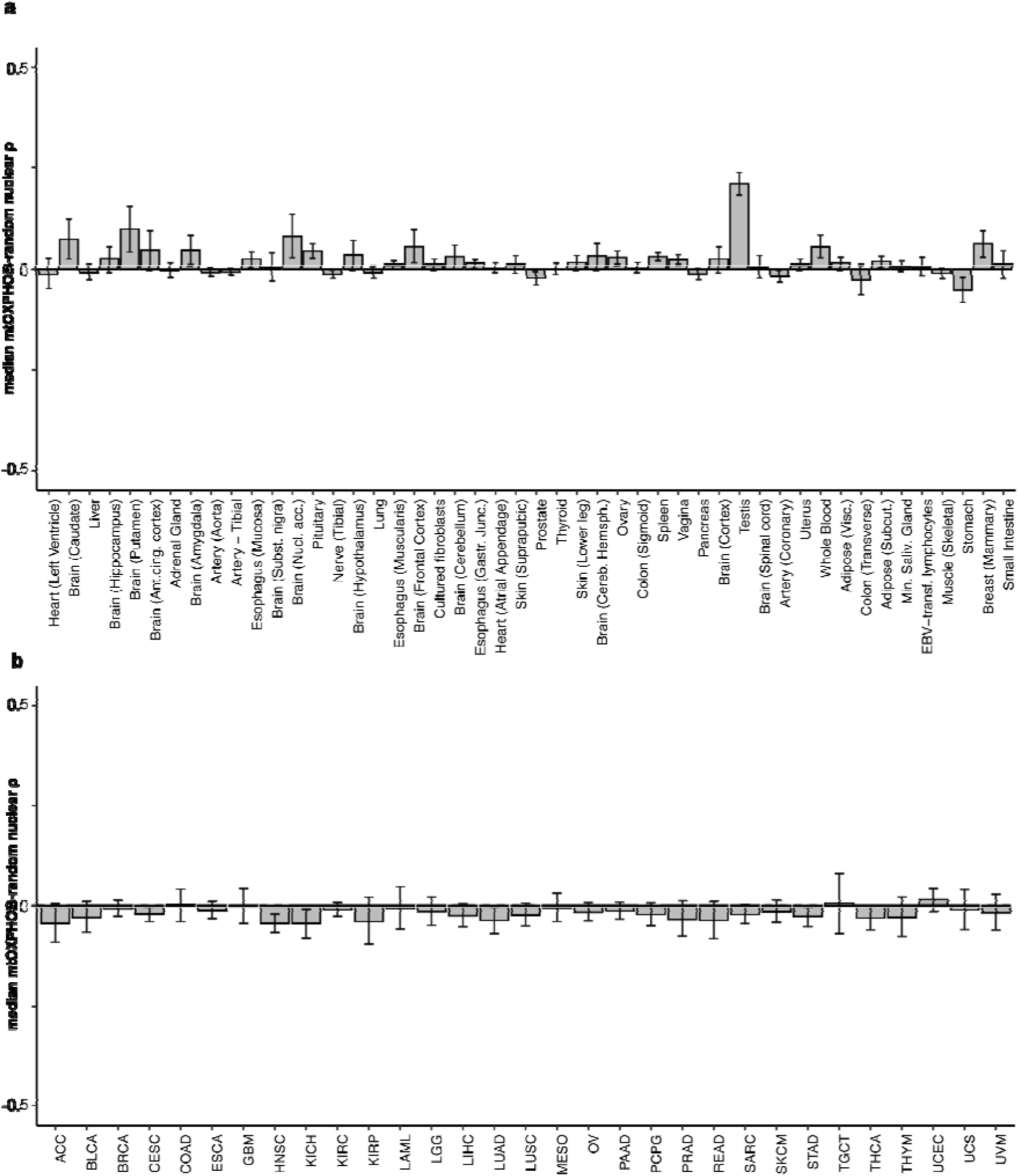
Correlations of mtOXPHOS with random nuclear genes do not resemble the observed mtOXPHOS-nuOXPHOS correlations across tissues or cancer types. **a)** Bars show mean median ρ for mtOXPHOS genes with 100 samples of 126 random nuclear genes in GTEx tissues; error bars indicate standard error. The testis has unusually high correlations between mtOXPHOS and random genes. The order of tissues matches the order of mtOXPHOS-nuOXPHOS correlation as shown in Fig 2d. The observed mtOXPHOS-nuOXPHOS correlation for each tissue is tested against this distribution in Table S1. **b)** Bars show mean median ρ for mtOXPHOS genes with 100 samples of 126 random nuclear genes in TCGA cancer types; error bars indicate standard error. The testis has unusually high correlations between mtOXPHOS and random genes. The order of tissues matches the order of mtOXPHOS-nuOXPHOS correlation as shown in Fig 4c. The observed mtOXPHOS-nuOXPHOS correlation for each cancer type is tested against this distribution in Table S2. Cancer abbreviations are found in Table S5.

**Fig S5.**
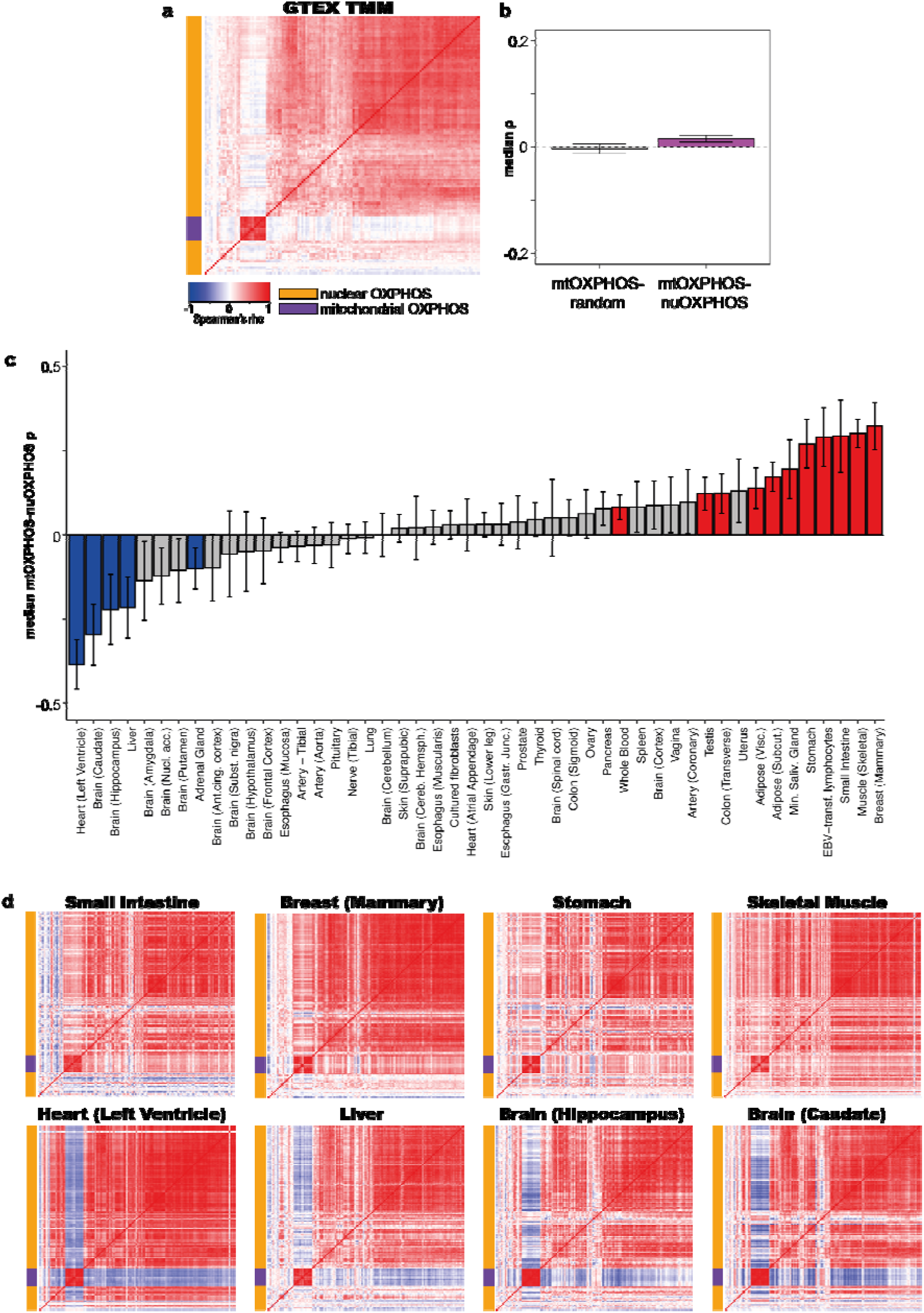
TMM normalization supports weak and inconsistent correlations between mtOXPHOS and nuOXPHOS within tissues. **a)** Heatmap showing median Spearman’s ρ for mtOXPHOS and nuOXPHOS gene expression using TMM normalization in 48 GTEx tissues combined (100 samples from each tissue, sampled 100 times). **b)** Boxplot showing median Spearman’s ρ for 100 iterations of mtOXPHOS genes with random nuclear genes or mtOXPHOS with nuOXPHOS genes for 48 GTEx tissues combined. **c)** Observed median Spearman’s ρ between mtOXPHOS and nuOXPHOS genes for 48 GTEx tissues. Error bars show 95% bootstrap confidence interval. Blue bars indicate observed correlation significantly lower than 0 (bootstrap empirical p-value, FDR-adjusted < 0.05), red bars indicate observed correlation significantly higher than 0 and grey bars indicate FDR > 0.05. **d)** Heatmap showing Spearman’s ρ for mtOXPHOS and nuOXPHOS genes for 8 GTEx tissues showing clear positive or negative correlations.

**Figure S6.**
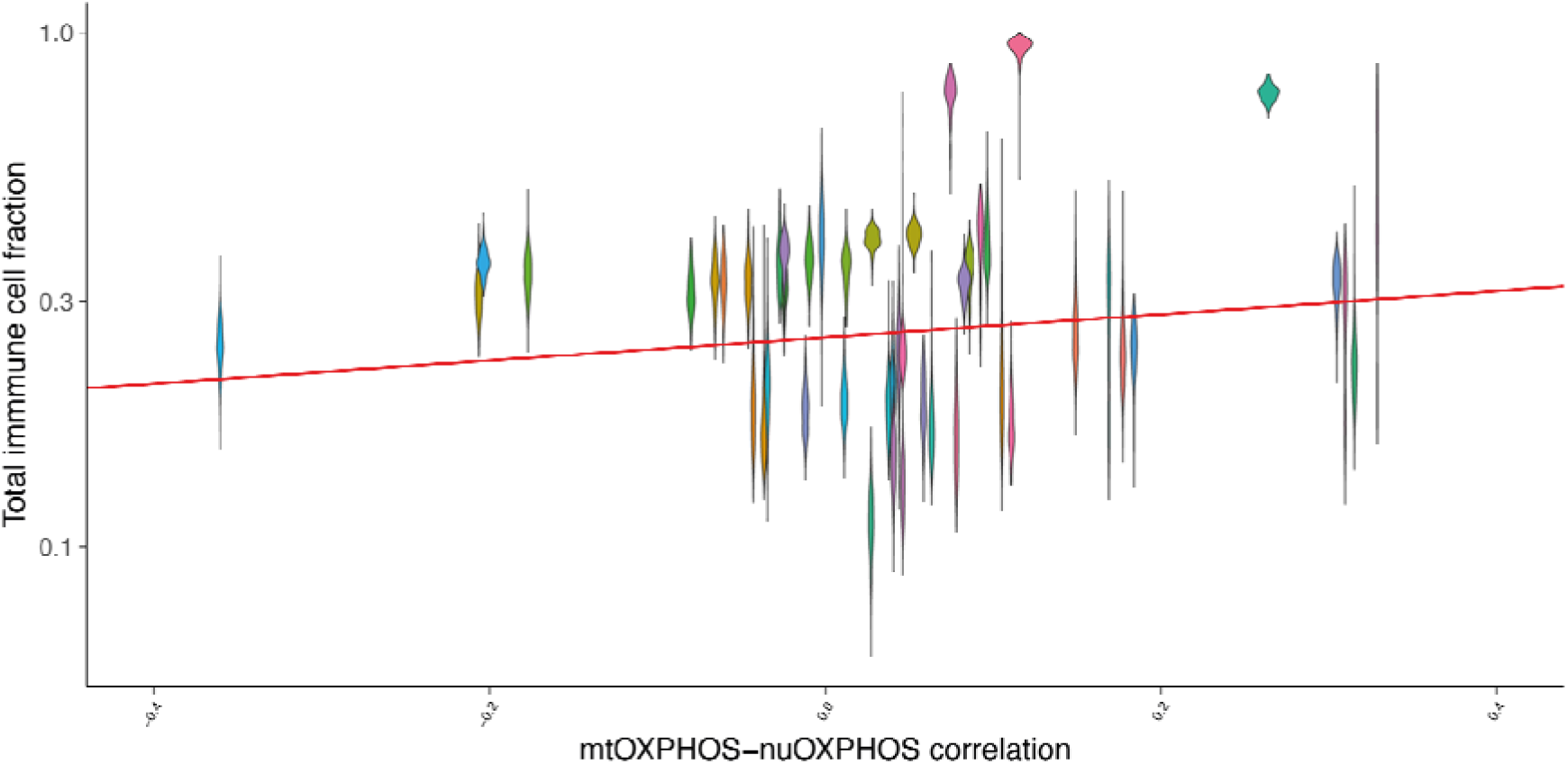
Total inferred immune cell fraction relates poorly to mtOXPHOS-nuOXPHOS correlation. Violins represent tissue types. Total inferred immune cell fraction for each sample was inferred from GEDIT, a gene expression deconvolution tool.

**Table 1.** Tissue mtOXPHOS-nuOXPHOS Spearman’s correlations for GTEx database relative to median mtOXPHOS-random nuclear gene correlations.

| **Tissue** | **Observed median mtOXPHOS-nuOXPHOS $\rho$** | **Mean median mtOXPHOS-random $\rho$** | **Shapiro-Wilk normality test FDR** | **Z-statistic (observed correlation)** | **FDR** |
| --- | --- | --- | --- | --- | --- |
| Small Intestine - Terminal Ileum | 0.329 | 0.010 | 0.517 | 9.88 | 3.52E-22 |
| Breast - Mammary Tissue | 0.315 | 0.059 | 0.878 | 7.84 | 1.91E-14 |
| Stomach | 0.310 | -0.051 | 0.992 | 11.7 | 1.89E-30 |
| Muscle - Skeletal | 0.305 | -0.010 | 0.946 | 25.3 | 5.16E-139 |
| Cells - EBV-transformed lymphocytes | 0.264 | 0.005 | 0.753 | 12.3 | 9.28E-34 |
| Minor Salivary Gland | 0.184 | 0.006 | 0.100 | 15.9 | 3.18E-55 |
| Adipose - Subcutaneous | 0.177 | 0.016 | 0.656 | 13.2 | 1.02E-38 |
| Colon - Transverse | 0.169 | -0.027 | 0.594 | 5.51 | 9.99E-08 |
| Adipose - Visceral (Omentum) | 0.149 | 0.012 | 0.869 | 9.23 | 1.43E-19 |
| Whole Blood | 0.116 | 0.053 | 0.517 | 2.20 | 3.91E-02 |
| Uterus | 0.111 | 0.010 | 0.878 | 7.79 | 2.74E-14 |
| Artery - Coronary | 0.105 | -0.017 | 0.878 | 8.10 | 2.63E-15 |
| Brain - Spinal cord (cervical c-1) | 0.096 | 0.004 | 0.953 | 3.55 | 7.25E-04 |
| Testis | 0.093 | 0.210 | 0.843 | -4.17 | 7.22E-05 |
| Brain - Cortex | 0.086 | 0.022 | 0.517 | 2.05 | 5.49E-02 |
| Pancreas | 0.083 | -0.011 | 0.926 | 7.05 | 6.67E-12 |
| Vagina | 0.078 | 0.021 | 0.656 | 4.83 | 3.58E-06 |
| Spleen | 0.075 | 0.028 | 0.628 | 4.73 | 5.58E-06 |
| Colon - Sigmoid | 0.063 | 0.002 | 0.953 | 6.00 | 6.29E-09 |
| Ovary | 0.058 | 0.025 | 0.878 | 1.96 | 6.69E-02 |
| Brain - Cerebellar Hemisphere | 0.053 | 0.029 | 0.517 | 0.746 | 4.97E-01 |
| Skin - Sun Exposed (Lower leg) | 0.046 | 0.013 | 0.753 | 1.91 | 7.17E-02 |
| Thyroid | 0.046 | -0.001 | 0.517 | 3.87 | 2.33E-04 |
| Prostate | 0.044 | -0.022 | 0.695 | 3.91 | 2.15E-04 |
| Skin - Not Sun Exposed (Suprapubic) | 0.041 | 0.010 | 0.946 | 1.53 | 1.54E-01 |
| Heart - Atrial Appendage | 0.040 | 0.002 | 0.882 | 3.85 | 2.42E-04 |
| Esophagus - Gastroesophageal Junction | 0.038 | 0.012 | 0.517 | 2.80 | 8.02E-03 |
| Brain - Cerebellum | 0.028 | 0.028 | 0.926 | 0.001 | 1.00E+00 |
| Cells - Cultured fibroblasts | 0.027 | 0.010 | 0.517 | 1.417 | 1.88E-01 |
| Brain - Frontal Cortex (BA9) | 0.012 | 0.053 | 0.992 | -0.989 | 3.69E-01 |
| Esophagus - Muscularis | 0.011 | 0.009 | 0.768 | 0.238 | 8.47E-01 |
| Lung | -0.002 | -0.008 | 0.594 | 0.400 | 7.35E-01 |
| Brain - Hypothalamus | -0.010 | 0.032 | 0.869 | -1.17 | 2.84E-01 |
| Nerve - Tibial | -0.012 | -0.012 | 0.517 | -0.021 | 1.00E+00 |
| Pituitary | -0.024 | 0.042 | 0.882 | -3.68 | 4.62E-04 |
| Brain - Nucleus accumbens (basal ganglia) | -0.025 | 0.079 | 0.517 | -1.91 | 7.17E-02 |
| Brain - Substantia nigra | -0.027 | 0.004 | 0.602 | -0.966 | 3.73E-01 |
| Esophagus - Mucosa | -0.034 | 0.022 | 0.953 | -3.10 | 3.16E-03 |
| Artery - Tibial | -0.036 | -0.006 | 0.869 | -3.48 | 9.02E-04 |
| Artery - Aorta | -0.043 | -0.007 | 0.594 | -3.42 | 1.06E-03 |
| Brain - Amygdala | -0.046 | 0.044 | 0.517 | -2.44 | 2.20E-02 |
| Adrenal Gland | -0.061 | -0.004 | 0.517 | -3.58 | 6.66E-04 |
| Brain - Anterior cingulate cortex (BA24) | -0.066 | 0.045 | 0.992 | -2.35 | 2.73E-02 |
| Brain - Putamen (basal ganglia) | -0.080 | 0.097 | 0.632 | -3.09 | 3.16E-03 |
| Brain - Hippocampus | -0.177 | 0.023 | 0.882 | -6.61 | 1.29E-10 |
| Liver | -0.204 | -0.008 | 0.882 | -10.9 | 8.41E-27 |
| Brain - Caudate (basal ganglia) | -0.206 | 0.071 | 0.602 | -5.62 | 5.89E-08 |
| Heart - Left Ventricle | -0.361 | -0.012 | 0.100 | -9.73 | 1.25E-21 |

**Table S2.**
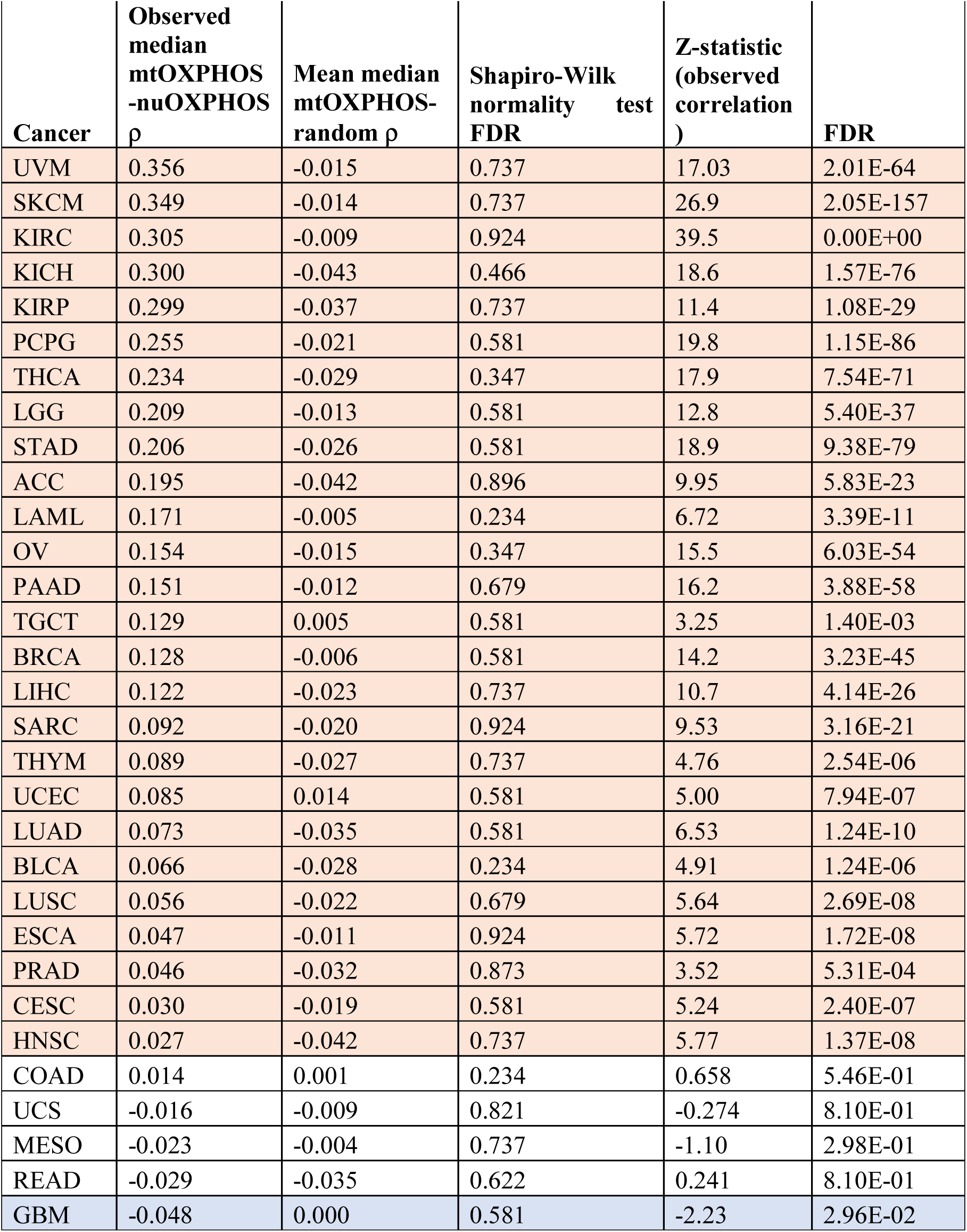
Cancer type mtOXPHOS-nuOXPHOS Spearman’s correlations for TCGA database relative to median mtOXPHOS-random nuclear gene correlations.

**Table S3.** Coefficients of a linear model predicting tissue mtOXPHOS-nuOXPHOS correlation by GTEx sample Proliferative Index (PI) and expression of NF-κB family members.

|  | Estimate | Std. Error | t value | Pr(>\|t\|) |
| --- | --- | --- | --- | --- |
| Intercept | -0.425181 | 0.024484 | -17.366 | < 2e-16 |
| Proliferative Index | -0.005170 | 0.001053 | -4.910 | 9.18e-07 |
| log<sub>10</sub>(NFKB1 expression) | 0.187359 | 0.007624 | 24.574 | < 2e-16 |
| log<sub>10</sub>(NFKB2 expression) | -0.077582 | 0.005614 | -13.819 | < 2e-16 |
| log<sub>10</sub>(REL expression) | 0.033591 | 0.003987 | 8.425 | < 2e-16 |
| log<sub>10</sub>(RELA expression) | -0.038051 | 0.008690 | -4.379 | 1.20e-05 |
| log<sub>10</sub>(RELB expression) | 0.068737 | 0.005065 | 13.570 | < 2e-16 |

Residual standard error: 0.1254 on 17237 degrees of freedom
Multiple R-squared: 0.1235, Adjusted R-squared: 0.1232
F-statistic: 404.7 on 6 and 17237 DF, p-value: < 2.2e-16

**Table S4.** Coefficients of a linear model predicting cancer type mtOXPHOS-nuOXPHOS correlation by TCGA sample Proliferative Index (PI) and expression of NF-κB family members.

|  | Estimate | Std. Error | t value | Pr(>\|t\|) |
| --- | --- | --- | --- | --- |
| Intercept | 0.7654839 | 0.0251287 | 30.463 | < 2e-16 |
| Proliferative Index | -0.0458482 | 0.0008316 | -55.135 | < 2e-16 |
| log<sub>10</sub>(NFKB1 expression) | -0.0310699 | 0.0047090 | -6.598 | 4.38e-11 |
| log<sub>10</sub>(NFKB2 expression) | 0.0412251 | 0.0043534 | 9.470 | < 2e-16 |
| log<sub>10</sub>(REL expression) | -0.0019117 | 0.0023865 | -0.801 | 0.423 |
| log<sub>10</sub>(RELA expression) | -0.0417820 | 0.0062731 | -6.661 | 2.88e-11 |
| log<sub>10</sub>(RELB expression) | -0.0227510 | 0.0046963 | -4.844 | 1.29e-06 |

Residual standard error: 0.08055 on 9714 degrees of freedom
Multiple R-squared: 0.294, Adjusted R-squared: 0.2935
F-statistic: 674.1 on 6 and 9714 DF, p-value: < 2.2e-16

**Table S5.** TCGA cancer type codes.

| **TCGA abbreviation** | **Cancer type** |
| --- | --- |
| LAML | Acute myeloid leukemia |
| ACC | Adrenocortical carcinoma |
| BLCA | Bladder urothelial carcinoma |
| LGG | Brain lower grade glioma |
| BRCA | Breast invasive carcinoma |
| CESC | Cervical squamous cell carcinoma and endocervical adenocarcinoma |
| COAD | Colon adenocarcinoma |
| ESCA | Esophageal carcinoma |
| GBM | Glioblastoma multiforme |
| HNSC | Head and neck squamous cell carcinoma |
| KICH | Kidney chromophobe |
| KIRC | Kidney renal clear cell carcinoma |
| KIRP | Kidney renal papillary cell carcinoma |
| LIHC | Liver hepatocellular carcinoma |
| LUAD | Lung adenocarcinoma |
| LUSC | Lung squamous cell carcinoma |
| MESO | Mesothelioma |
| OV | Ovarian serous cystadenocarcinoma |
| PAAD | Pancreatic adenocarcinoma |
| PCPG | Pheochromocytoma and paraganglioma |
| PRAD | Prostate adenocarcinoma |
| READ | Rectum adenocarcinoma |
| SARC | Sarcoma |
| SKCM | Skin cutaneous melanoma |
| STAD | Stomach adenocarcinoma |
| TGCT | Testicular germ cell tumors |
| THYM | Thymoma |
| THCA | Thyroid carcinoma |
| UCS | Uterine carcinosarcoma |
| UCEC | Uterine corpus endometrial carcinoma |
| UVM | Uveal melanoma |

## Supplementary datasets

**Supplementary dataset 1.** mtOXPHOS-nuOXPHOS expression heatmaps using MRN normalization for 48 GTEx tissues.

**Supplementary dataset 2.** mtOXPHOS-nuOXPHOS expression heatmaps using MRN normalization for 31 TCGA cancer types.

**Supplementary dataset 3.** mtOXPHOS-nuOXPHOS expression heatmaps for matched tumour samples for 14 cancer types.

**Supplementary dataset 4.** mtOXPHOS-nuOXPHOS expression heatmaps for matched normal samples for 14 cancer types.

**Supplementary dataset 5**. Gene Ontology enrichment for the top 1000 genes, whose expression across GTEx tissues is most positively correlated with tissue mtOXPHOS-nuOXPHOS correlation.

**Supplementary dataset 6**. Gene Ontology enrichment for the bottom 1000 genes, whose expression across GTEx tissues is most negatively correlated with tissue mtOXPHOS-nuOXPHOS correlation.

**Supplementary dataset 7**. Gene Ontology enrichment for the top 1000 genes whose expression across TCGA cancer types is most positively correlated with cancer type mtOXPHOS-nuOXPHOS correlation.

**Supplementary dataset 8**. Gene Ontology enrichment for the bottom 1000 genes, whose expression across TCGA cancer types is most negatively correlated with cancer type mtOXPHOS-nuOXPHOS correlation.

**Table S6.** Spreadsheet containing HGNC symbols and ENSEMBL gene IDs for mtOXPHOS and nuOXPHOS genes used in this study.

**Supplementary text 1.** R script used for all analyses, annotated according to corresponding figure panels.

## Methods

### Raw data

RNA-Seq data were downloaded from the GTEx data portal for GTEx v8, apart from data shown in Fig S2 from GTEx v6p. Data were downloaded as normalized TPM values (or FPKM values for GTEx v6p, later also converted to TPM) or as raw counts. ‘Harmonised’ (hg38) RNA-seq data were downloaded for TCGA projects using the ‘TCGAbiolinks’ package in R as normalized FPKM values or as raw counts.

For TCGA cancer type analyses, we only considered samples annotated as Primary Tumours, except where we explicitly note that we perform analyses on adjacent normal tissue samples.

We were mindful of the possibility of contamination of mitochondrial RNA reads by expression of nuclear integrations of mitochondrial DNA (NUMTs). Previous studies have established that NUMT contamination of mtDNA expression quantification in RNAseq data is negligible for the GTEx (Barshad, Blumberg et al. 2018) and TCGA (Reznik, Wang et al. 2017) databases.

We restricted our analyses to GTEx tissues with at least 100 samples or TCGA cancer types with at least 50 Primary Tumour samples.

### mtOXPHOS and nuOXPHOS genes

The list of mtOXPHOS and nuOXPHOS genes was taken from Barshad, Blumberg et al. (2018) and can be found in Table S6.

### Normalizations

For TCGA data and GTEx v6p, TPM values were obtained by converting FPKM values according to the following formula for gene *i*:

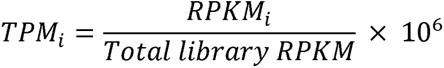

The MRN normalization was performed using the ‘DESeq2’ package in R. Normalizations were applied individually for each tissue or cancer type cohort. Normalized pseudocounts were obtained by converting raw counts data to a DEseq2DataSet using the ‘DESeqDataSetFromMatrix()’ function, applying the ‘estimateSizeFactors()’ function to the resulting dds object, and then retrieving the normalized pseudocounts with the function ‘counts()’ with ‘normalized = TRUE’.

The TMM and UQ normalizations were performed using the ‘edgeR’ package in R. Normalizations were applied individually for each tissue or cancer type cohort. Raw counts were converted into a DGEList object with the function ‘DGEList’. Normalization factors were returned using the function ‘calcNormFactors()’ with method set as either ‘TMM’ or ‘UQ’. Library scaling factors were then obtained by multiplying the resulting library sizes by the resulting normalization factors. All read counts for each library/sample were then multiplied by 10^6^ and divided by the scaling factor for that library.

TPM (nuclear only) values for nuclear-encoded genes were obtained by multiplying raw read counts for nuclear genes by 10^6^ and dividing by the sample sum of reads for all genes excluding those borne on mitochondrial DNA. mtDNA-encoded genes were normalized with the total sample sum instead, as for TPM. We did not normalize for transcript length; the values thus calculated should be considered as Counts Per Million. However, for correlation analysis, which does not involve direct comparison of genes within samples, this is equivalent to TPM.

TPM values excluding haemoglobin reads or *PRSS1/2* reads for GTEx Whole Blood or Pancreas samples respectively were obtained as for TPM nuclear only but excluding the read counts for those genes when computing the total sample read count, instead of the read counts for the mitochondrial genes.

### Correlations

Correlations were computed using the ‘cor.test()’ function in R, with ‘method = “spearman”’ or ‘method = “pearson”’.

### Linear model correction of data prior to correlation analysis

Before correlation analysis, we applied linear model corrections to normalized data to control for known confounders in the GTEx and TCGA datasets.

Sample and donor information for the GTEx were downloaded from the GTEx data portal in the form of the Subject Phenotypes and Subject Annotations files accompanying GTEx v8 (or v6p as appropriate). TCGA metadata was obtained using the ‘all_metadata(subset = “tcga”)’ function of the ‘recount’ package in R. Some TCGA donors provided multiple cancer samples; these were all excluded to leave only samples unique to a single donor.

Linear models were applied within each tissue or cancer type cohort, to account for tissue-specific trends. We found that to apply linear models pooling GTEx tissues was to introduce grave artifacts which can profoundly affect downstream analysis. A linear model was fitted for each gene using the ‘lm()’ function in R and the resulting residuals were used downstream in correlation analyses. GTEx data were corrected for age bracket of the patient, sex of the patient, cause of death (Hardy scale), sample ischemic time and sequencing batch. Age bracket was considered to be a numerical variable, and the midpoint of the appropriate age bracket was taken for each sample to be entered into the linear model. We opted to use the publicly available age-bracket data so that our analyses can be readily reproduced. TCGA data (including matched normal samples) was corrected for the gender, race, tumour stage and age of the patient and the sequencing centre where the samples were processed.

### Correlations combining tissues or cancer types

To perform a correlation analysis on combined tissues or cancer types, we took the residuals of expression data that had been normalized and corrected by fitting a linear model to control for confounding variables, as above. We filtered the tissue or cancer types for those with at least 100 (GTEx) or 50 (TCGA) samples, leaving 48 tissues and 31 cancers respectively. To ensure equal representation of each tissue or cancer type, we randomly sampled 100 (GTEx) or 50 (TCGA) from each tissue. Combining raw gene expression residuals for tissues or cancer types with different mtRNA levels may introduce artifacts, as high-expressing tissues will have much greater variance in the absolute value of the residuals, despite the linear model controlling for the tissue average gene expression level. To account for this, we ranked the sample residuals for each gene within the sample chosen for each tissue or cancer type. We then combined the ranks for the chosen samples across tissue types, using the ranks in place of the raw residuals; this gave us 4800 samples for GTEx or 1550 samples for TCGA. The Spearman’s correlation was then computed across these aggregated ranks. As the result varies slightly depending on the random sampling within each tissue, we repeated this process 100 times. Heatmaps show the median correlation for any gene pair over the 100 iterations. The order of mtOXPHOS/nuOXPHOS genes for all heatmaps for both GTEx and TCGA data throughout the manuscript is identical and was determined by clustering of the correlations for the MRN-normalized data combining GTEx tissues. Although the gene names are not shown in main figure panels, the genes are fully annotated with HGNC symbols in Supplementary Datasets 1-4.

In order to provide a null distribution for the correlation of mtOXPHOS genes with nuOXPHOS genes, we repeated the process as above, but replaced the 126 nuOXPHOS genes with a random sample of 126 expressed nuclear genes (median TPM > 5 across all samples for all tissues/cancer types) for each iteration. This process was repeated 100 times. In order to test the observed mtOXPHOS-nuOXPHOS correlation against this distribution, the 100 median correlations for mtOXPHOS-nuOXPHOS gene pairs for each iteration was compared to the 100 median correlations of mtOXPHOS genes with random nuclear genes with a two-tailed t test.

### Correlations within tissues or cancer types

For correlations reported within cancer types or tissue types, all available samples were utilised for the calculations. To provide a bootstrapped distribution, we calculated the median mtOXPHOS-nuOXPHOS for each tissue or cancer type 1000 times in samples constituted by resampling with replacement. The 95% confidence intervals shown in the main figure panels were calculated by multiplying the standard deviation of these 1000 bootstrap estimates of median mtOXPHOS-nuOXPHOS by 1.96. We used empirical p-values from the bootstrap distributions to test if observed median mtOXPHOS-nuOXPHOS correlations were different from 0. Raw p-values were the fraction of bootstrap values > 0 for tissue/cancer types s with observed negative correlations or < 0 for positive correlations. These p values were then corrected to FDR across tissue/cancer types.

To test the significance of mtOXPHOS-nuOXPHOS correlations against mtOXPHOS with random nuclear genes, correlations were computed for each tissue/cancer type for the mtOXPHOS genes and 126 random nuclear genes (median TPM > 5 across all samples from all tissues/cancer types). This was repeated 100 times and the medians of the correlations of mtOXPHOS genes with random nuclear genes from each iteration were taken to be the null distribution. We confirmed that these medians conformed to a normal distribution, as would be expected under the Central Limit Theorem, by performing a Shapiro-Wilk normality test on the 100 medians for each tissue. The adjusted p-values (FDR) for the Shapiro-Wilk tests are reported in Table S1 and Table S2. The Z-statistic for the observed median mtOXPHOS-nuOXPHOS correlation was then calculated by subtracting the mean and dividing by the standard deviation of the mtOXPHOS-random nuclear medians. p-values were then calculated for each tissue according to the normal cumulative distribution function and corrected to FDR (Table S1 and Table S2).

### Matched cancer and normal samples

Matched primary tumour and adjacent normal tissue samples were identified using TCGA metadata and barcodes. Tissues were identified with at least 10 normal samples. Using the donor portion of the TCGA barcode, matching primary tumour samples were identified. If multiple primary tumour samples matched the adjacent normal tissue sample, one was retained at random and the remainder were discarded. Additionally, normal tissue samples without lacking identifiable primary tumour samples in the expression data were discarded, such that all normal tissue samples had one matching primary tumour sample and vice versa. mtOXPHOS-nuOXPHOS correlations were computed using all samples. To calculate the bootstrapped standard error, we repeated the correlations using unique subsets of 90% of the datapoints 100 times. The bootstrapped standard error for the median mtOXPHOS-nuOXPHOS correlation is the standard deviation of the bootstrapped median mtOXPHOS-nuOXPHOS correlations.

### Dip test for bimodality

To test for bimodality in the distribution of correlations among random nuclear genes, correlations for 100 sets of random genes were computed for all GTEx tissues using MRN-normalized data. Hartigan’s dip test was then performed for each set of 100 random genes in each tissue using the ‘dip.test()’ function of the ‘diptest’ package in R. Dip test p-values were adjusted to FDR within each tissue.

### Gene ontology enrichment analysis

To test for genes whose expression was associated with mtOXPHOS-nuOXPHOS co-ordination, we calculated the mean expression of each gene in each tissue or cancer type (in pseudocounts generated by MRN normalisation of raw counts across all tissues/cancers). We then computed the Spearman’s correlation for each gene of the mean tissue/cancer pseudocounts with tissue/cancer mtOXPHOS-nuOXPHOS correlation calculated earlier using MRN normalisation. We ordered the gene lists in order of correlation and took the top or bottom 1000 genes. As these were ENSEMBL gene IDs, we retrieved the HGNC symbols for these genes using the ‘biomaRt’ package in R; this typically resulted in the loss of a fraction of the genes which lack HGNC symbols (such as some lncRNA genes). We then submitted the HGNC symbols to the ‘enrichr()’ function of the ‘enrichR’ package in R, using the reference databases ‘KEGG_2021_Human’, ‘GO_Molecular_Function_2018’, ‘GO_Cellular_Component_2018’ and ‘GO_Biological_Process_2018’.

### Proliferative Index

The Proliferative Index (PI) was calculated with the ‘ProliferativeIndex’ package in R. Briefly, the entire dataset across all tissues or cancer types was normalized by MRN and variance stabilising transformation using the ‘varianceStabilizingTransformation()’ function of DESeq2. Following the normalization, the PI was calculated by applying the ‘readDataForPI()’ function with a randomly selected gene specified in the ‘modelIDs’ argument, then running ‘calculatePI()’ on the resulting object.

### Estimations of total immune fraction

The estimates for the total immune fraction of GTEx samples were taken directly from the GitHub repository associated with the GEDIT tool (Nadel, Lopez et al. 2021).

## References

Anders, S. and W. Huber (2010). “Differential expression analysis for sequence count data.” Nature Precedings: 1–1.

Barshad, G., A. Blumberg, T. Cohen and D. Mishmar (2018). “Human primitive brain displays negative mitochondrial-nuclear expression correlation of respiratory genes.” Genome research 28(7): 952–967.

Boardman, N. T., B. Migally, C. Pileggi, G. S. Parmar, J. Y. Xuan, K. Menzies and M.-E. Harper (2021). “Glutaredoxin-2 and Sirtuin-3 deficiencies impair cardiac mitochondrial energetics but their effects are not additive.” Biochimica et Biophysica Acta (BBA)- Molecular Basis of Disease 1867(1): 165982.

Bullard, J. H., E. Purdom, K. D. Hansen and S. Dudoit (2010). “Evaluation of statistical methods for normalization and differential expression in mRNA-Seq experiments.” BMC bioinformatics 11(1): 1–13.

Butow, R. A. and N. G. Avadhani (2004). “Mitochondrial signaling: the retrograde response.” Molecular cell 14(1): 1–15.

Cogswell, P. C., D. F. Kashatus, J. A. Keifer, D. C. Guttridge, J. Y. Reuther, C. Bristow, S. Roy, D. W. Nicholson and A. S. Baldwin Jr (2003). “NF-κB and IκBα are found in the mitochondria: evidence for regulation of mitochondrial expression by NF-κB.” Journal of Biological Chemistry 278(5): 2963–2968.

Ferreira, P. G., M. Muñoz-Aguirre, F. Reverter, C. P. S. Godinho, A. Sousa, A. Amadoz, R. Sodaei, M. R. Hidalgo, D. Pervouchine and J. Carbonell-Caballero (2018). “The effects of death and post-mortem cold ischemia on human tissue transcriptomes.” Nature communications 9(1): 1–15.

Galai, G., H. Ben-David, L. Levin, M. F. Orth, T. G. Grünewald, S. Pilosof, S. Berstein and B. Rotblat (2020). “Pan-Cancer Analysis of Mitochondria Chaperone-Client Co-Expression Reveals Chaperone Functional Partitioning.” Cancers 12(4): 825.

Haynes, C. M., K. Petrova, C. Benedetti, Y. Yang and D. Ron (2007). “ClpP mediates activation of a mitochondrial unfolded protein response in C. elegans.” Developmental cell 13(4): 467–480.

Li, B., V. Ruotti, R. M. Stewart, J. A. Thomson and C. N. Dewey (2010). “RNA-Seq gene expression estimation with read mapping uncertainty.” Bioinformatics 26(4): 493–500.

Liberti, M. V. and J. W. Locasale (2016). “The Warburg effect: how does it benefit cancer cells?” Trends in biochemical sciences 41(3): 211–218.

Liu, H., H. Zheng, Z. Duan, D. Hu, M. Li, S. Liu, Z. Li, X. Deng, Z. Wang and M. Tang (2009). “LMP1-augmented kappa intron enhancer activity contributes to upregulation expression of Ig kappa light chain via NF-kappaB and AP-1 pathways in nasopharyngeal carcinoma cells.” Molecular Cancer 8(1): 1–18.

Lovén, J., D. A. Orlando, A. A. Sigova, C. Y. Lin, P. B. Rahl, C. B. Burge, D. L. Levens, T. I. Lee and R. A. Young (2012). “Revisiting global gene expression analysis.” Cell 151(3): 476–482.

Marquardt, A., A. G. Solimando, A. Kerscher, M. Bittrich, C. Kalogirou, H. Kübler, A. Rosenwald, R. Bargou, P. Kollmannsberger and B. Schilling (2021). “Subgroup-Independent Mapping of Renal Cell Carcinoma—Machine Learning Reveals Prognostic Mitochondrial Gene Signature Beyond Histopathologic Boundaries.” Frontiers in Oncology 11.

Mercer, T. R., S. Neph, M. E. Dinger, J. Crawford, M. A. Smith, A.-M. J. Shearwood, E. Haugen, C. P. Bracken, O. Rackham and J. A. Stamatoyannopoulos (2011). “The human mitochondrial transcriptome.” Cell 146(4): 645–658.

Mortazavi, A., B. A. Williams, K. McCue, L. Schaeffer and B. Wold (2008). “Mapping and quantifying mammalian transcriptomes by RNA-Seq.” Nature methods 5(7): 621–628.

Münch, C. and J. W. Harper (2016). “Mitochondrial unfolded protein response controls matrix pre-RNA processing and translation.” Nature 534(7609): 710–713.

Nadel, B. B., D. Lopez, D. J. Montoya, F. Ma, H. Waddel, M. M. Khan, S. Mangul and M. Pellegrini (2021). “The Gene Expression Deconvolution Interactive Tool (GEDIT): Accurate cell type quantification from gene expression data.” GigaScience 10(2): giab002.

Owusu-Ansah, E., W. Song and N. Perrimon (2013). “Muscle mitohormesis promotes longevity via systemic repression of insulin signaling.” Cell 155(3): 699–712.

Parikh, V. S., M. M. Morgan, R. Scott, L. S. Clements and R. A. Butow (1987). “The mitochondrial genotype can influence nuclear gene expression in yeast.” Science 235(4788): 576–580.

Ramaker, R. C., B. N. Lasseigne, A. A. Hardigan, L. Palacio, D. S. Gunther, R. M. Myers and S. J. Cooper (2017). “RNA sequencing-based cell proliferation analysis across 19 cancers identifies a subset of proliferation-informative cancers with a common survival signature.” Oncotarget 8(24): 38668.

Reznik, E., M. L. Miller, Y. Şenbabaoğlu, N. Riaz, J. Sarungbam, S. K. Tickoo, H. A. Al-Ahmadie, W. Lee, V. E. Seshan and A. Hakimi (2016). “Mitochondrial DNA copy number variation across human cancers.” elife 5: e10769.

Reznik, E., Q. Wang, K. La, N. Schultz and C. Sander (2017). “Mitochondrial respiratory gene expression is suppressed in many cancers.” elife 6: e21592.

Robinson, M. D. and A. Oshlack (2010). “A scaling normalization method for differential expression analysis of RNA-seq data.” Genome biology 11(3): 1–9.

Sakharkar, M. K., S. Kaur Dhillon, S. B. Chidambaram, M. M. Essa and J. Yang (2020). “Gene pair correlation coefficients in sphingolipid metabolic pathway as a potential prognostic biomarker for breast cancer.” Cancers 12(7): 1747.

Soumillon, M., A. Necsulea, M. Weier, D. Brawand, X. Zhang, H. Gu, P. Barthes, M. Kokkinaki, S. Nef and A. Gnirke (2013). “Cellular source and mechanisms of high transcriptome complexity in the mammalian testis.” Cell reports 3(6): 2179–2190.

Taanman, J.-W. (1999). “The mitochondrial genome: structure, transcription, translation and replication.” Biochimica et Biophysica Acta (BBA)-Bioenergetics 1410(2): 103–123.

Tang, Z., C. Li, B. Kang, G. Gao, C. Li and Z. Zhang (2017). “GEPIA: a web server for cancer and normal gene expression profiling and interactive analyses.” Nucleic acids research 45(W1): W98–W102.

Wang, Z., M. Gerstein and M. Snyder (2009). “RNA-Seq: a revolutionary tool for transcriptomics.” Nature reviews genetics 10(1): 57–63.

Xia, B., Y. Yan, M. Baron, F. Wagner, D. Barkley, M. Chiodin, S. Y. Kim, D. L. Keefe, J. P. Alukal and J. D. Boeke (2020). “Widespread transcriptional scanning in the testis modulates gene evolution rates.” Cell 180(2): 248–262. e221.

Yang, Y., Y. Zhang, L. Miao, W. Liao and W. Liao (2020). “LncRNA PPP1R14B-AS1 promotes tumor cell proliferation and migration via the enhancement of mitochondrial respiration.” Frontiers in Genetics 11.

Zhao, Q., J. Wang, I. V. Levichkin, S. Stasinopoulos, M. T. Ryan and N. J. Hoogenraad (2002). “A mitochondrial specific stress response in mammalian cells.” The EMBO journal 21(17): 4411–4419.

Zhao, S., Z. Ye and R. Stanton (2020). “Misuse of RPKM or TPM normalization when comparing across samples and sequencing protocols.” RNA 26(8): 903–909.

