## Supplemental tables and scripts for "Malignancy and NF-kB signalling strengthen coordination between the expression of mitochondrial and nuclear-encoded oxidative phosphorylation genes": Supplementary-Dataset-1_MRN-mtOXPHOS-nuOXPHOS-tissue-heatmaps.pdf

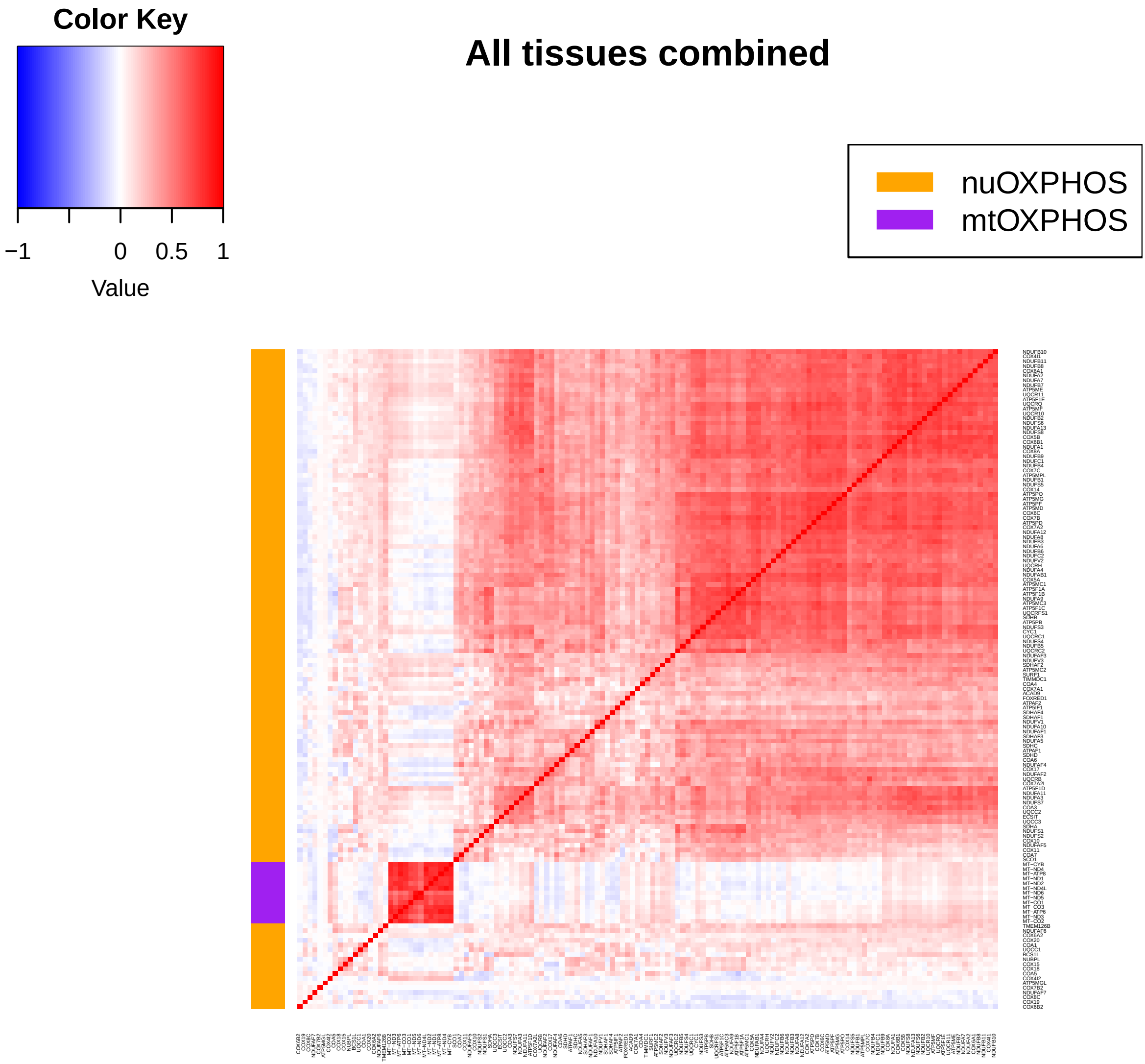
Color Key
−1
0
0.5
1
Value
All tissues combined
nuOXPHOS
mtOXPHOS

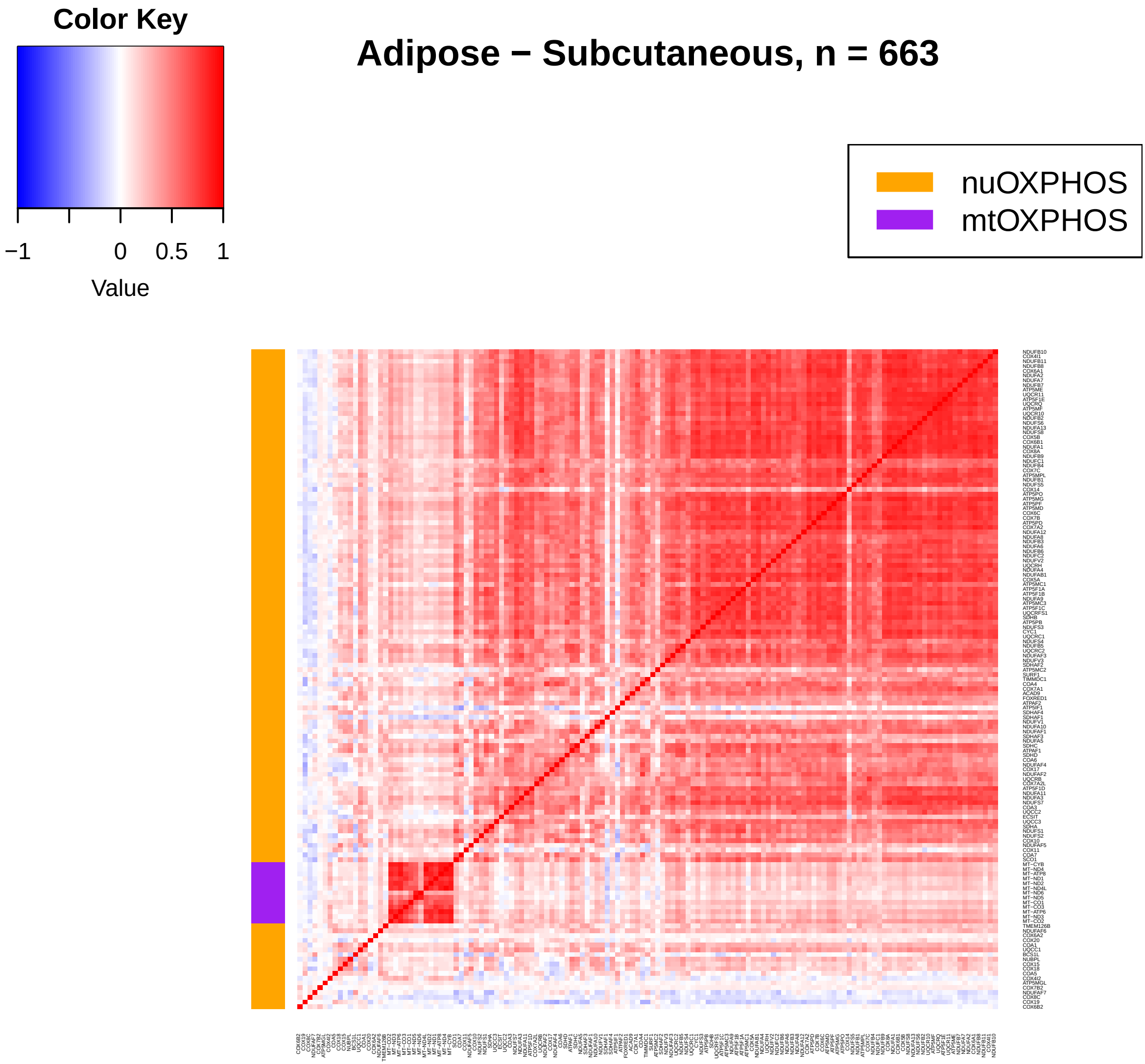
Color Key
−1
0
0.5
1
Value
Adipose − Subcutaneous, n = 663
nuOXPHOS
mtOXPHOS

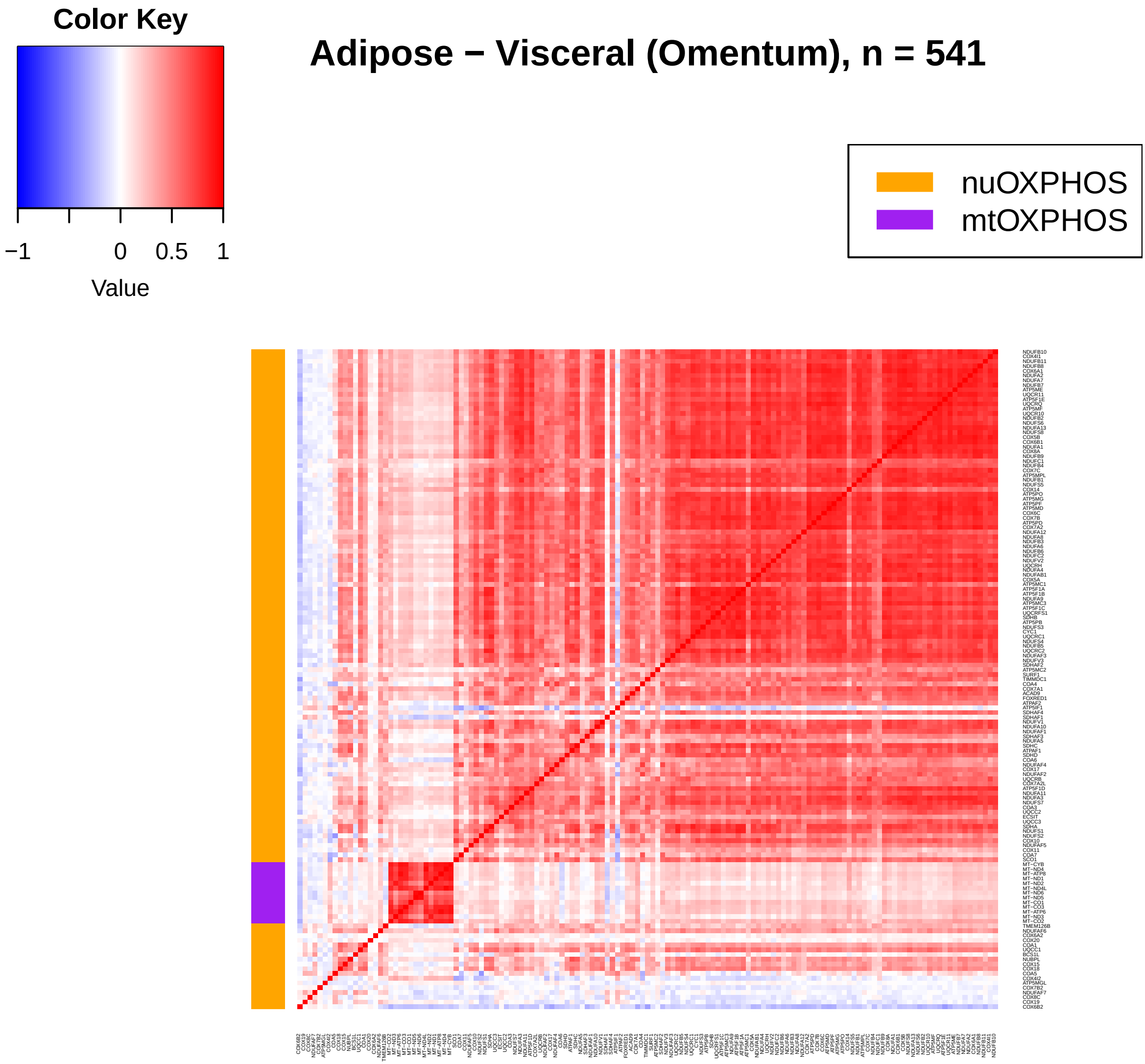
Color Key
−1
0
0.5
1
Value
Adipose − Visceral (Omentum), n = 541
nuOXPHOS
mtOXPHOS

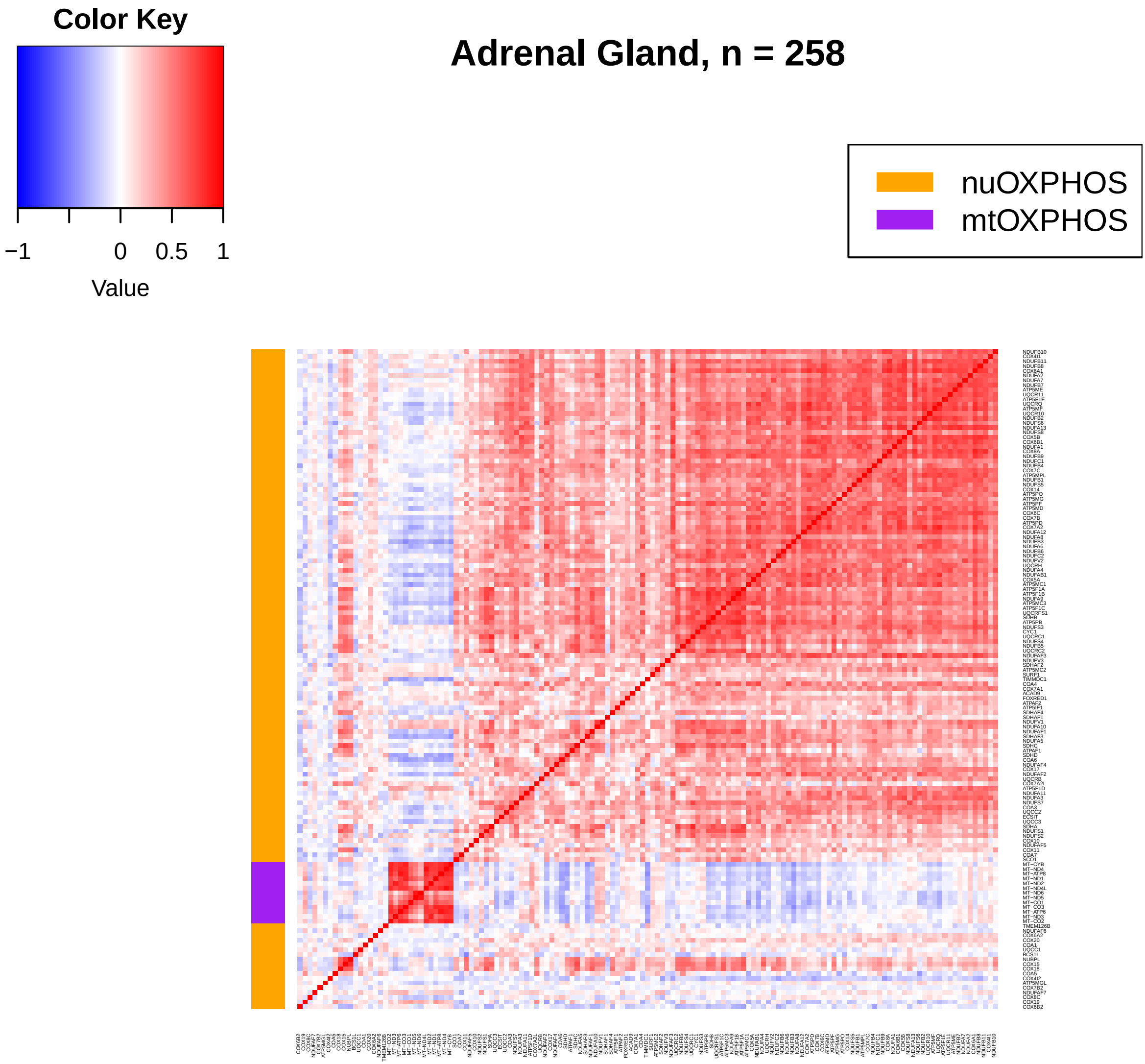

Color Key
−1
0
0.5
1
Value
Adrenal Gland, n = 258
nuOXPHOS
mtOXPHOS

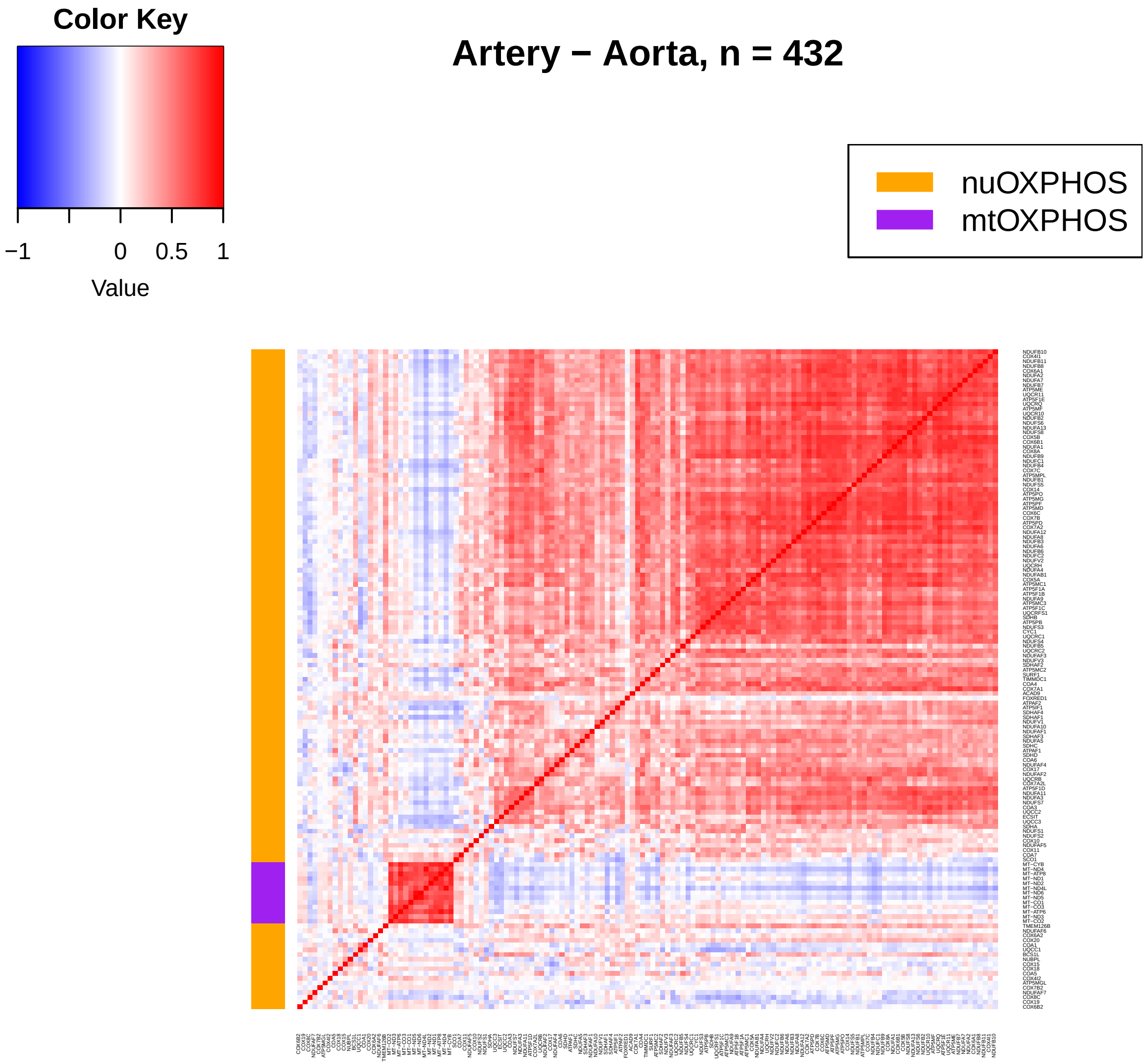
Color Key
−1
0
0.5
1
Value
Artery − Aorta, n = 432
nuOXPHOS
mtOXPHOS

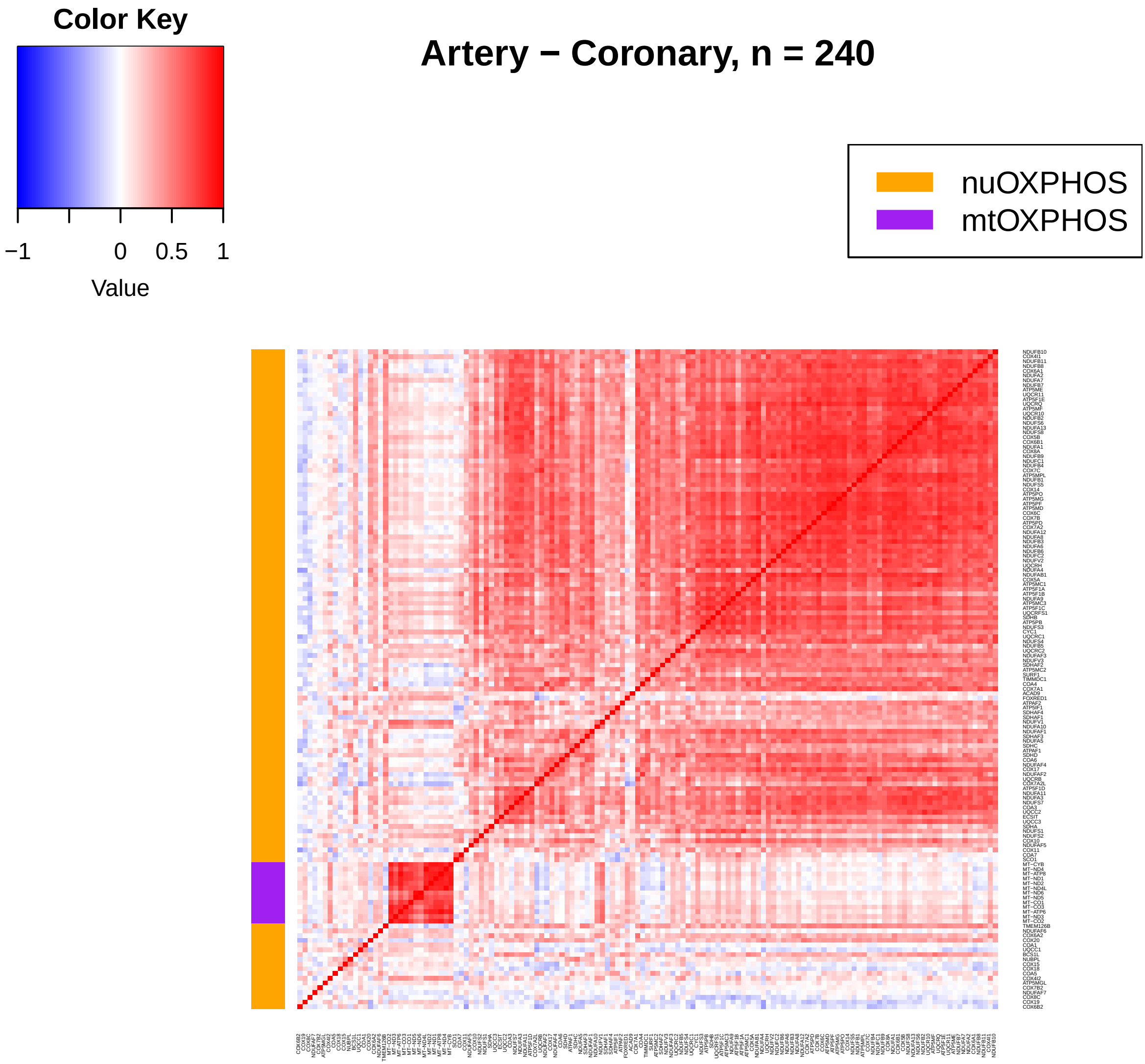

Color Key
−1
0
0.5
1
Value
Artery − Coronary, n = 240
nuOXPHOS
mtOXPHOS

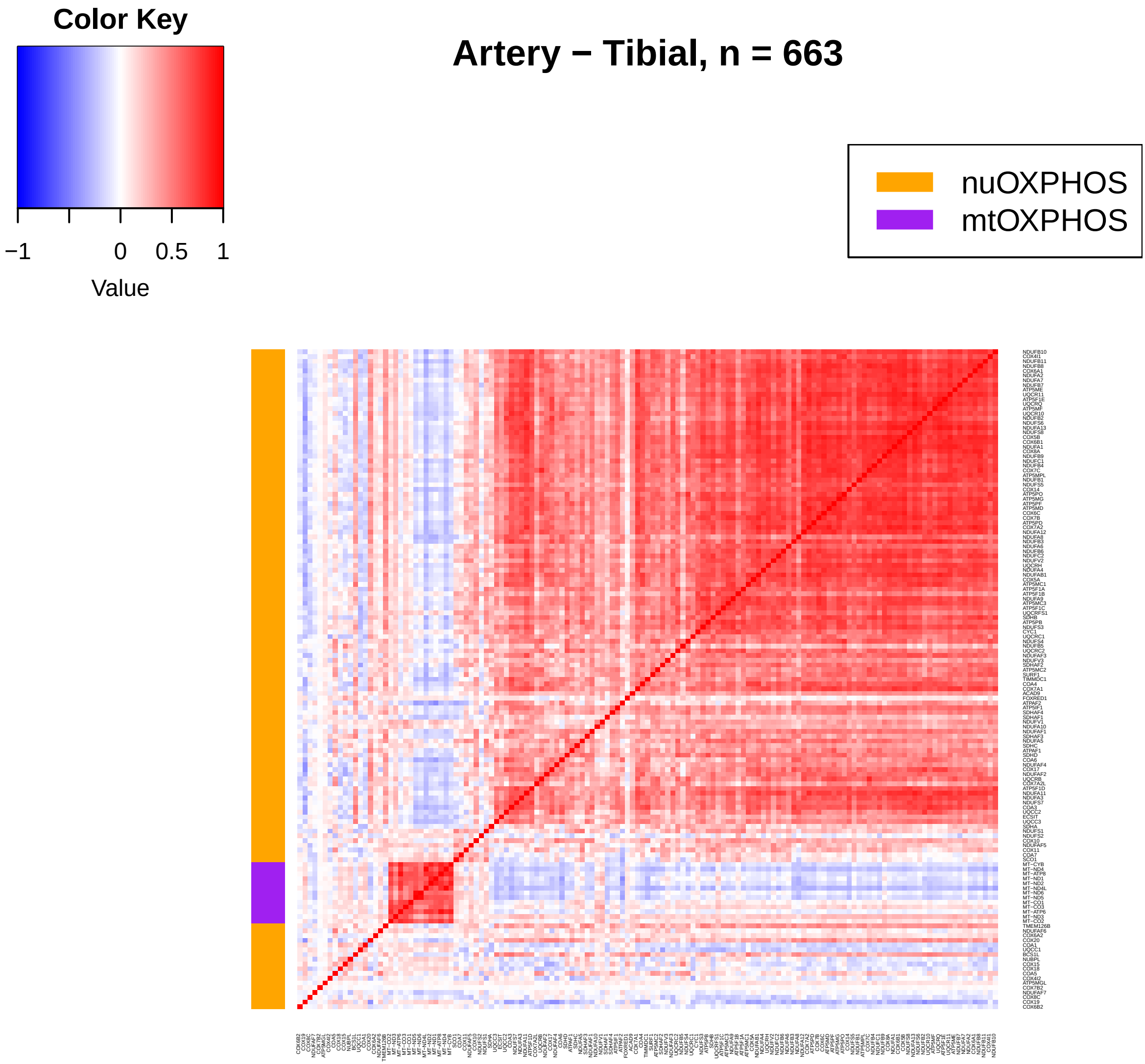
Color Key
−1
0
0.5
1
Value
Artery − Tibial, n = 663
nuOXPHOS
mtOXPHOS

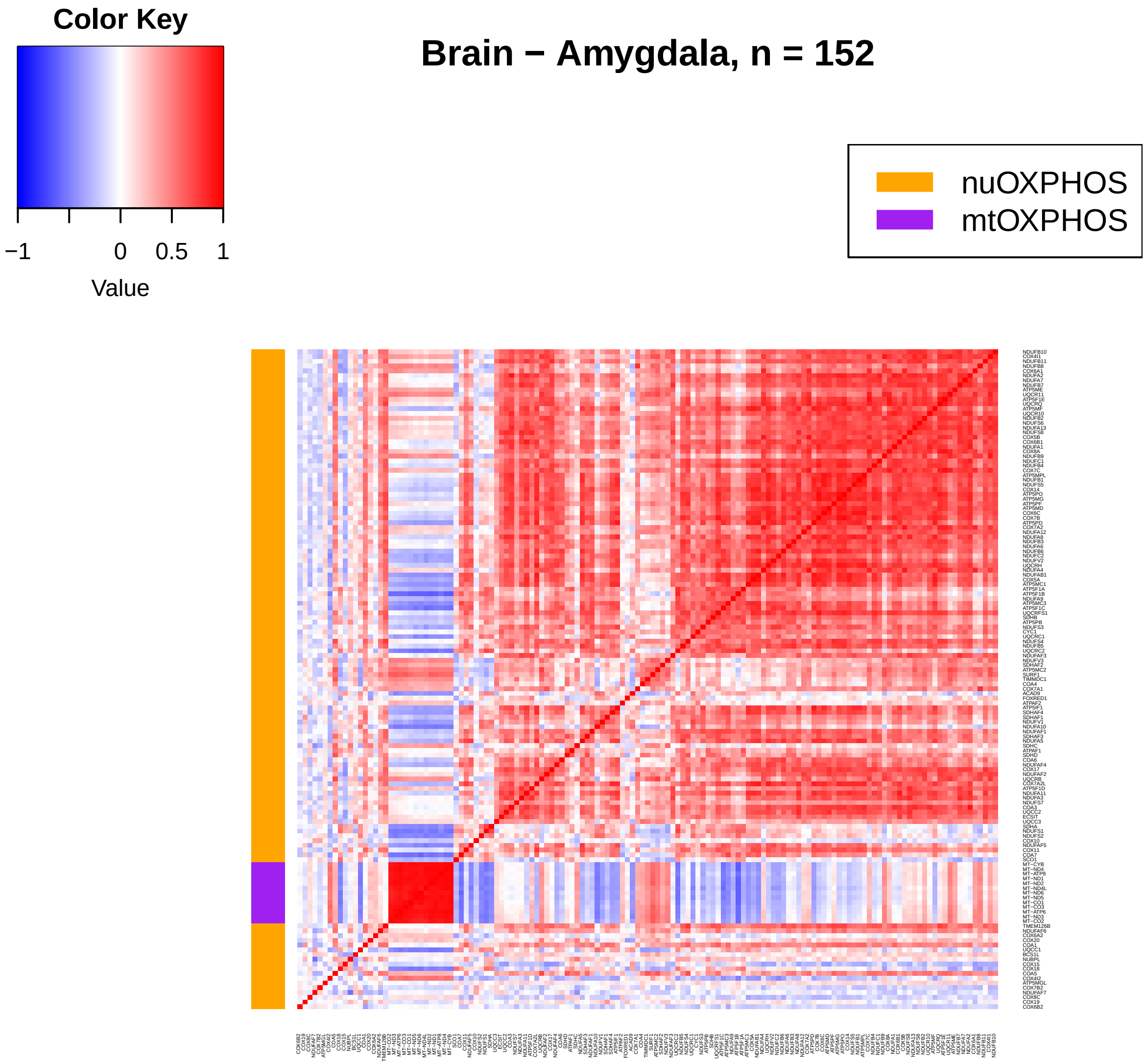
Color Key
−1
0
0.5
1
Value
Brain − Amygdala, n = 152
nuOXPHOS
mtOXPHOS

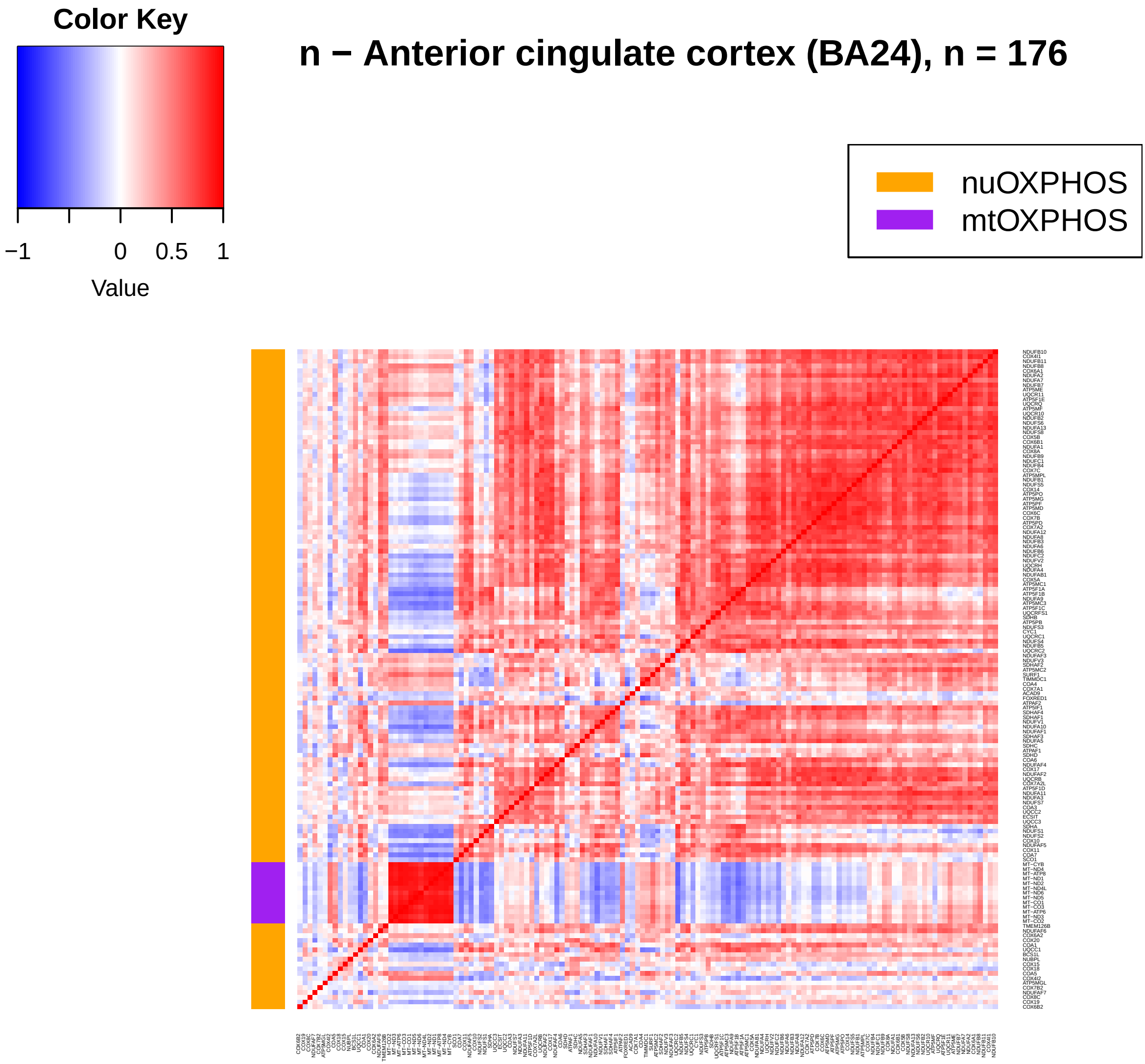
Color Key
−1
0
0.5
1
Value
n − Anterior cingulate cortex (BA24), n = 176
nuOXPHOS
mtOXPHOS

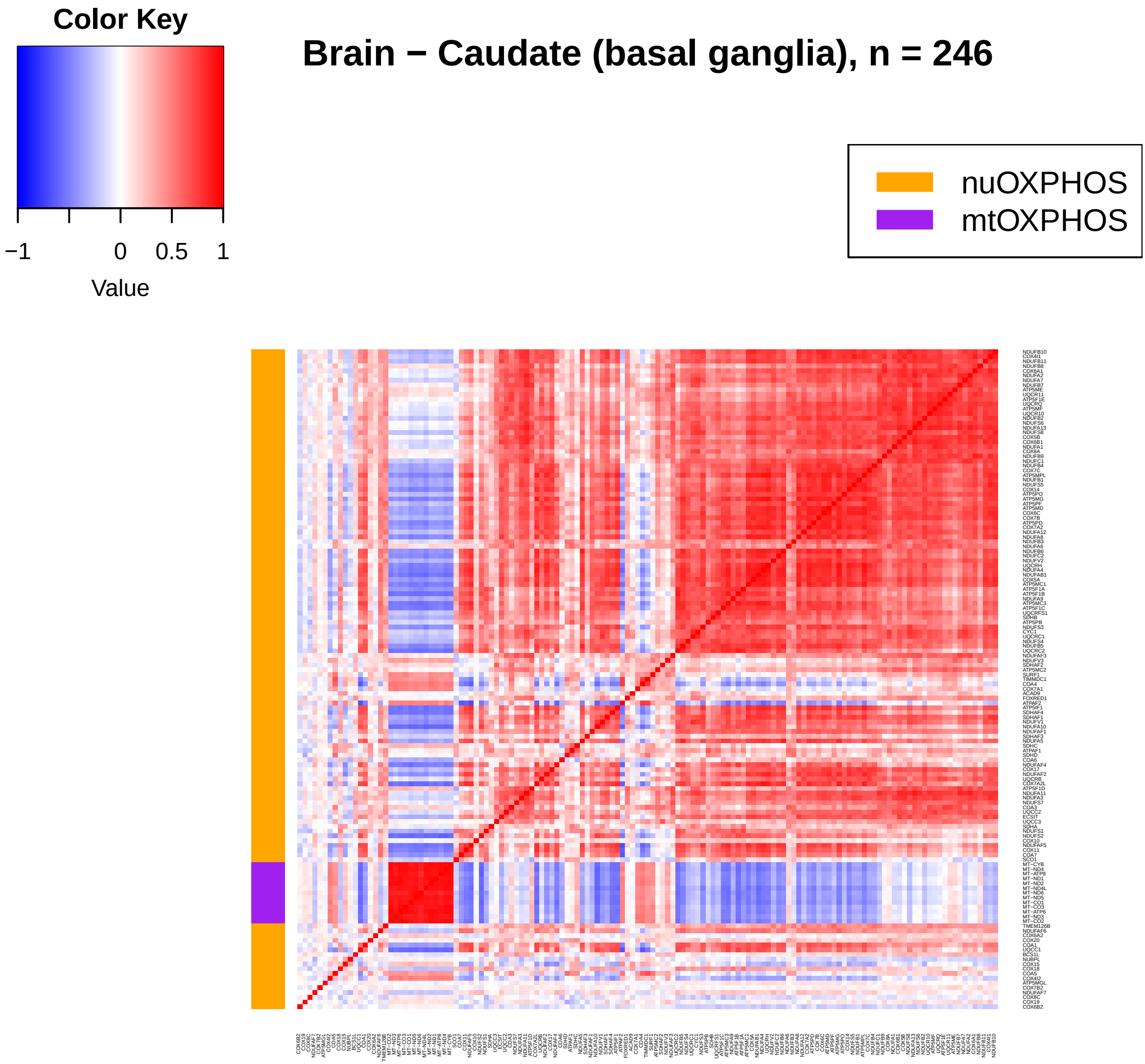
Brain − Caudate (basal ganglia), n = 246
Color Key
−1
0
0.5
1
Value
nuOXPHOS
mtOXPHOS

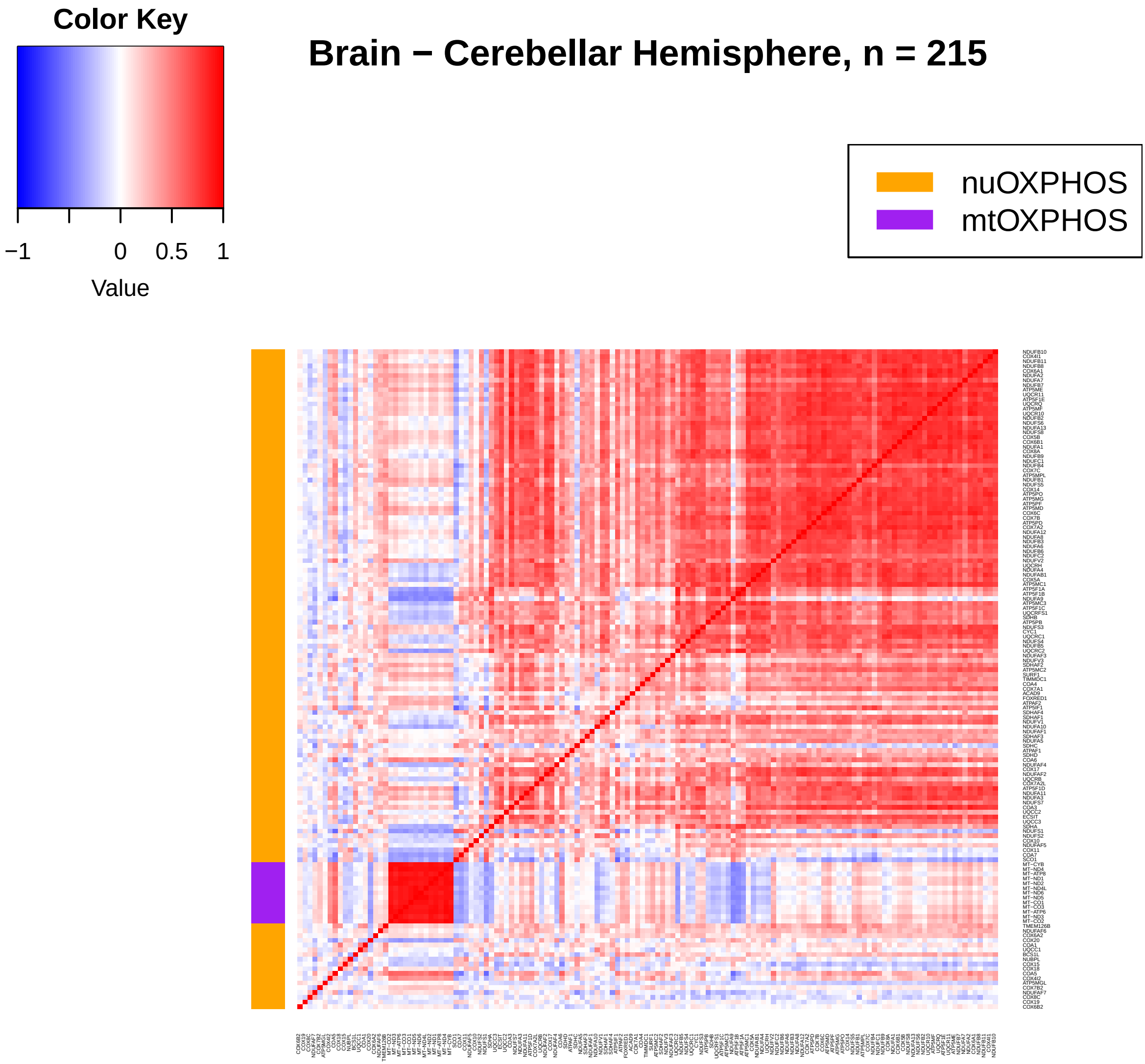
Color Key
−1
0
0.5
1
Value
Brain − Cerebellar Hemisphere, n = 215
nuOXPHOS
mtOXPHOS

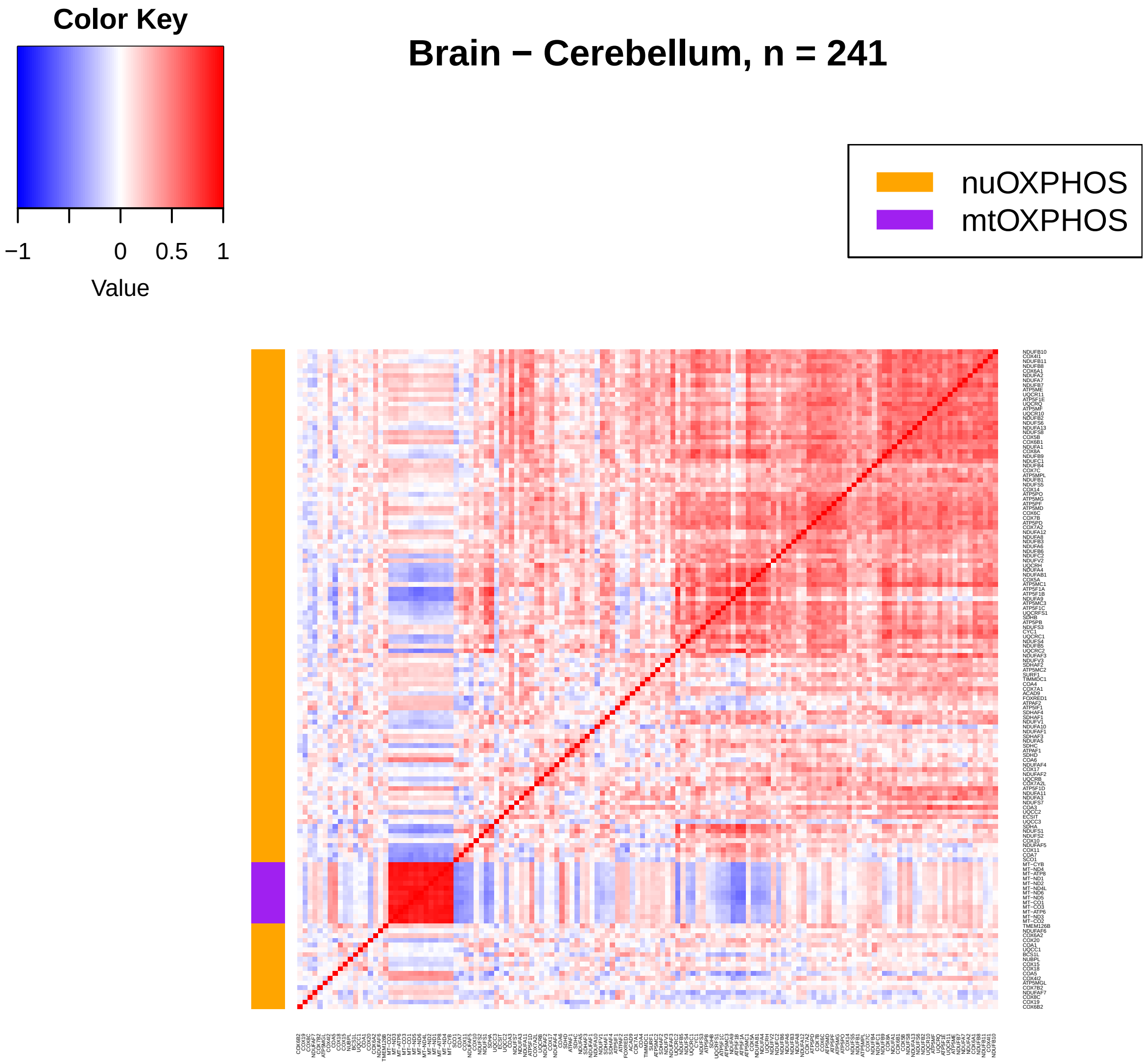
Color Key
−1
0
0.5
1
Value
Brain − Cerebellum, n = 241
nuOXPHOS
mtOXPHOS

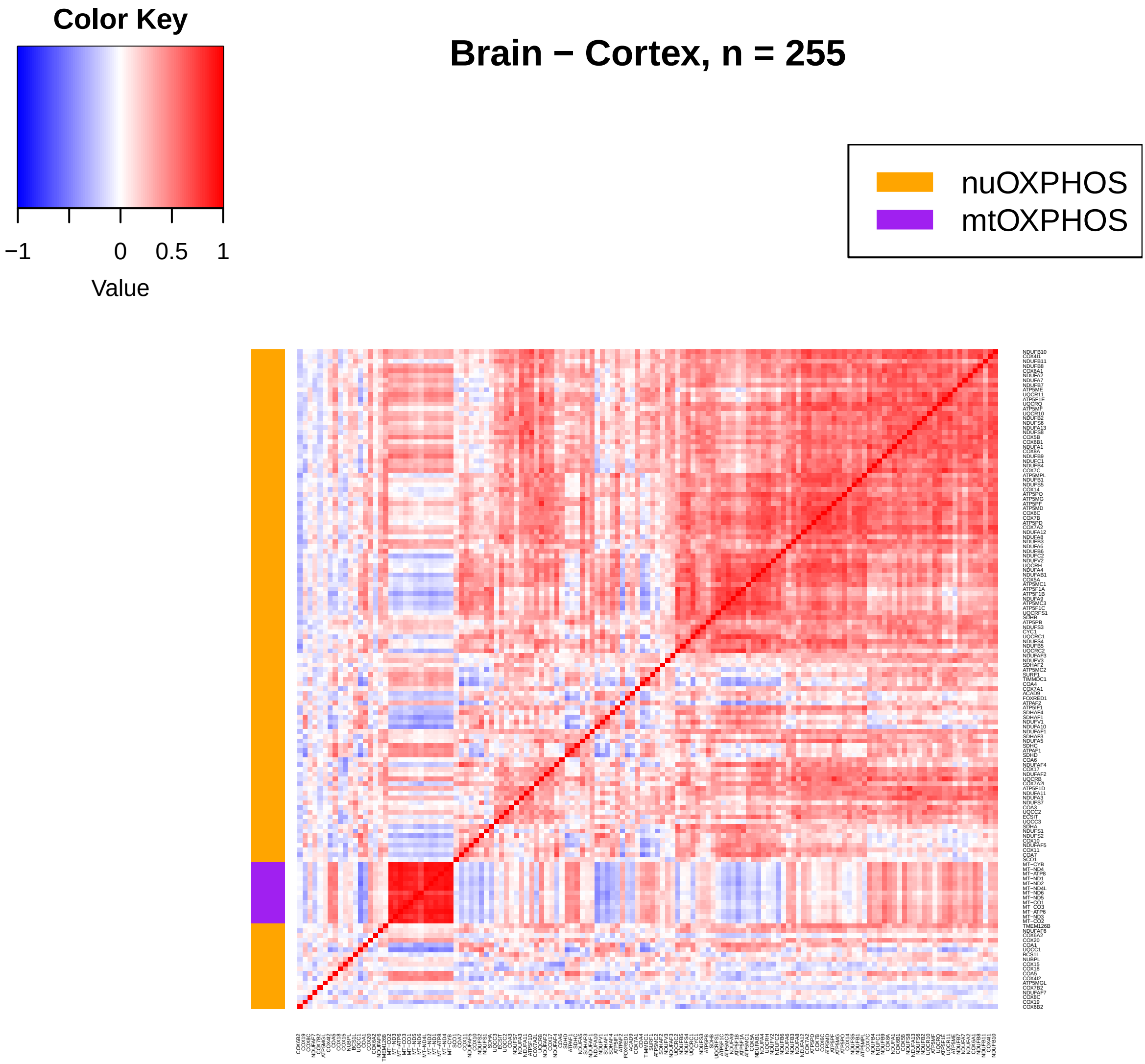
Color Key
−1
0
0.5
1
Value
Brain − Cortex, n = 255
nuOXPHOS
mtOXPHOS

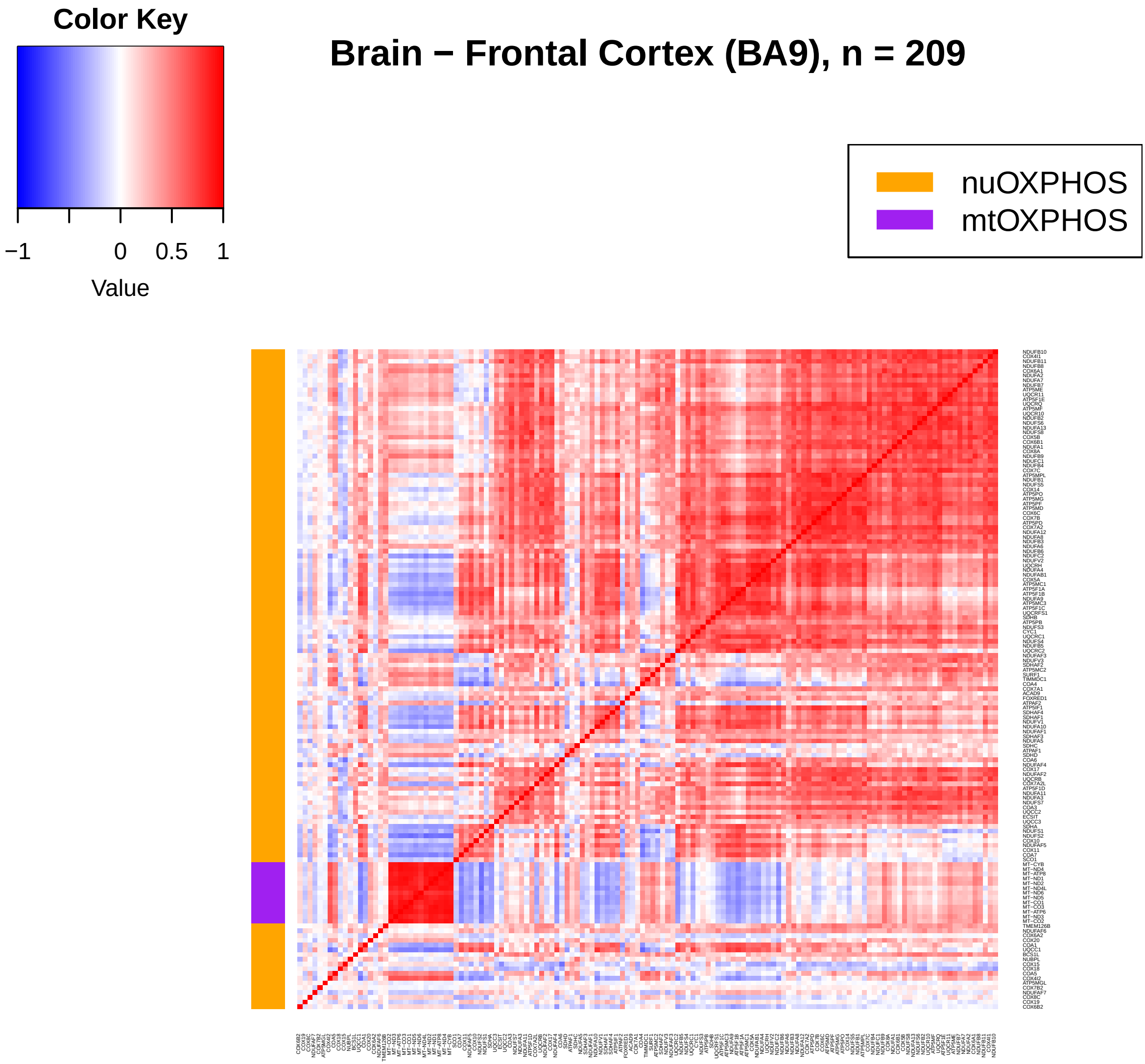
Color Key
−1
0
0.5
1
Value
Brain − Frontal Cortex (BA9), n = 209
nuOXPHOS
mtOXPHOS

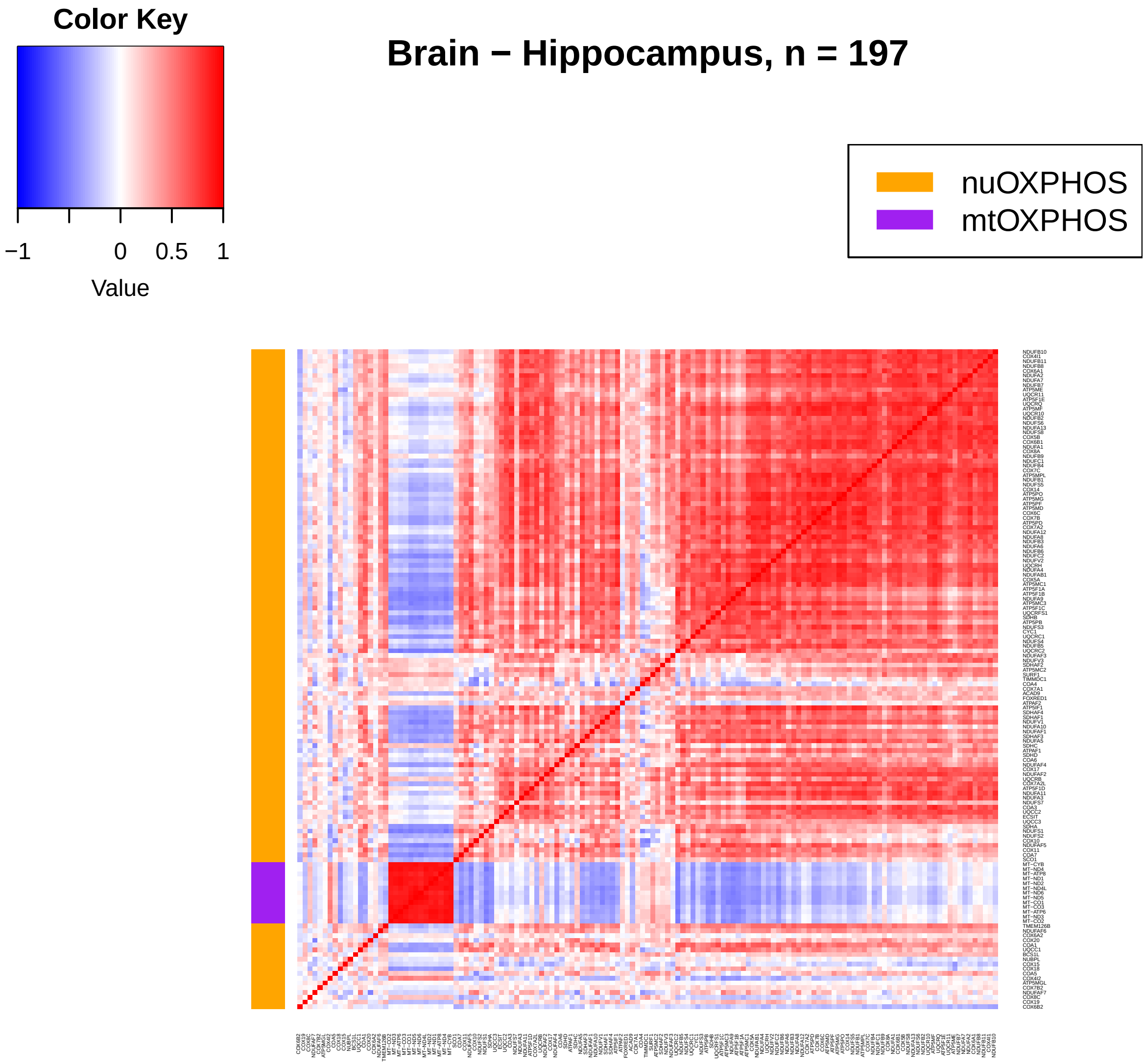
Color Key
−1
0
0.5
1
Value
Brain − Hippocampus, n = 197
nuOXPHOS
mtOXPHOS

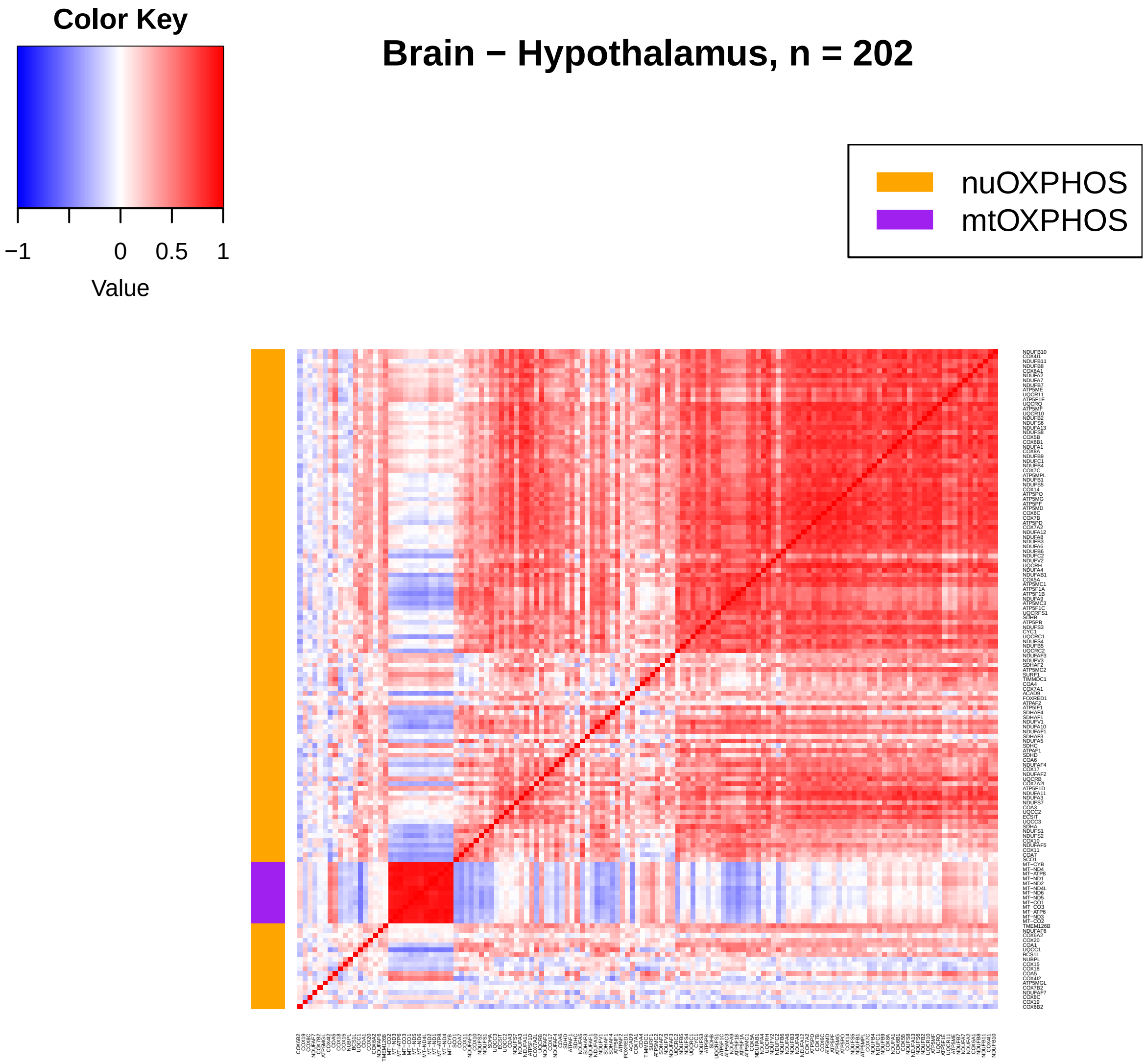
Color Key
−1
0
0.5
1
Value
Brain − Hypothalamus, n = 202
nuOXPHOS
mtOXPHOS

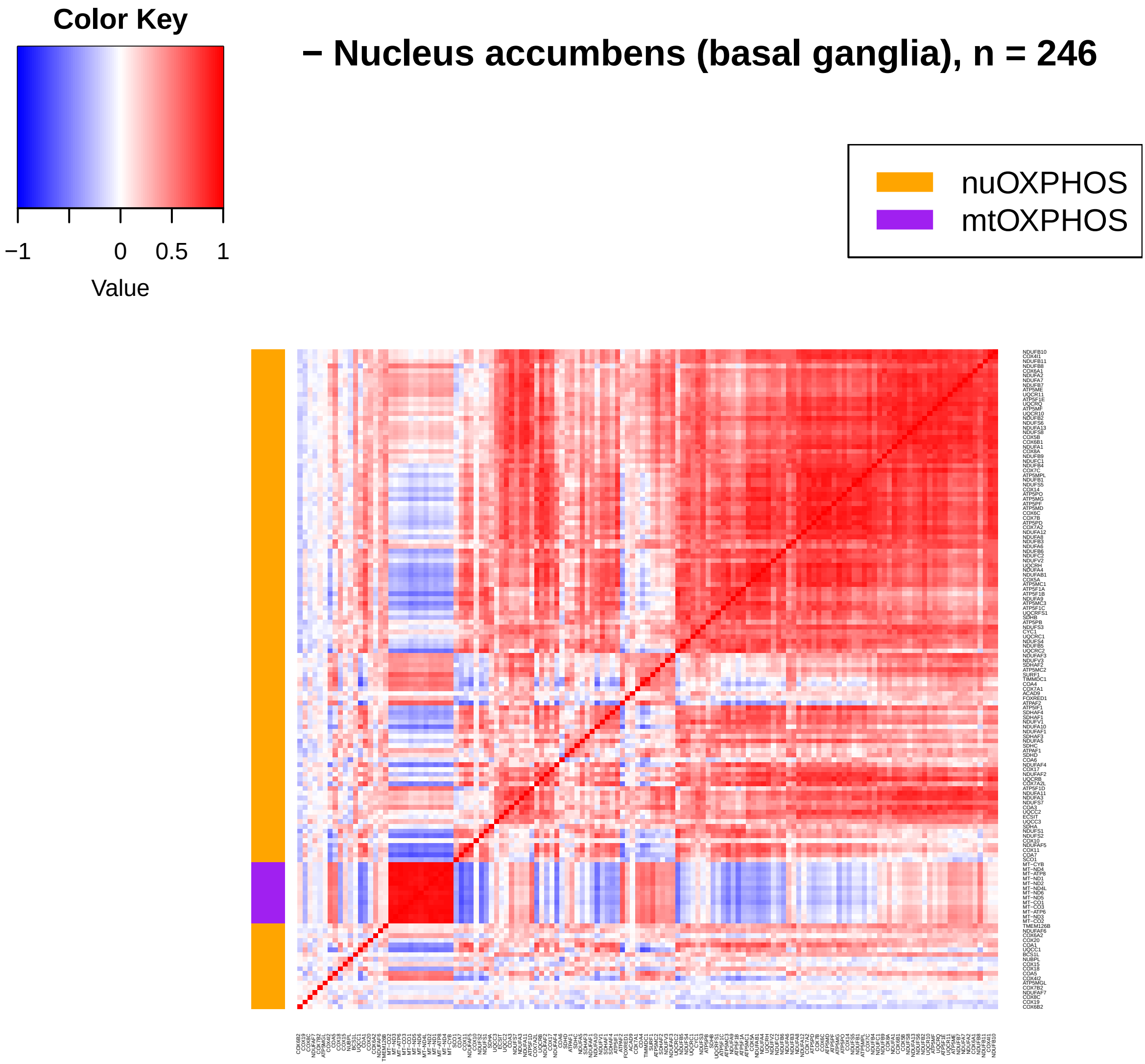
Color Key
−1
0
0.5
1
Value
− Nucleus accumbens (basal ganglia), n = 246
nuOXPHOS
mtOXPHOS

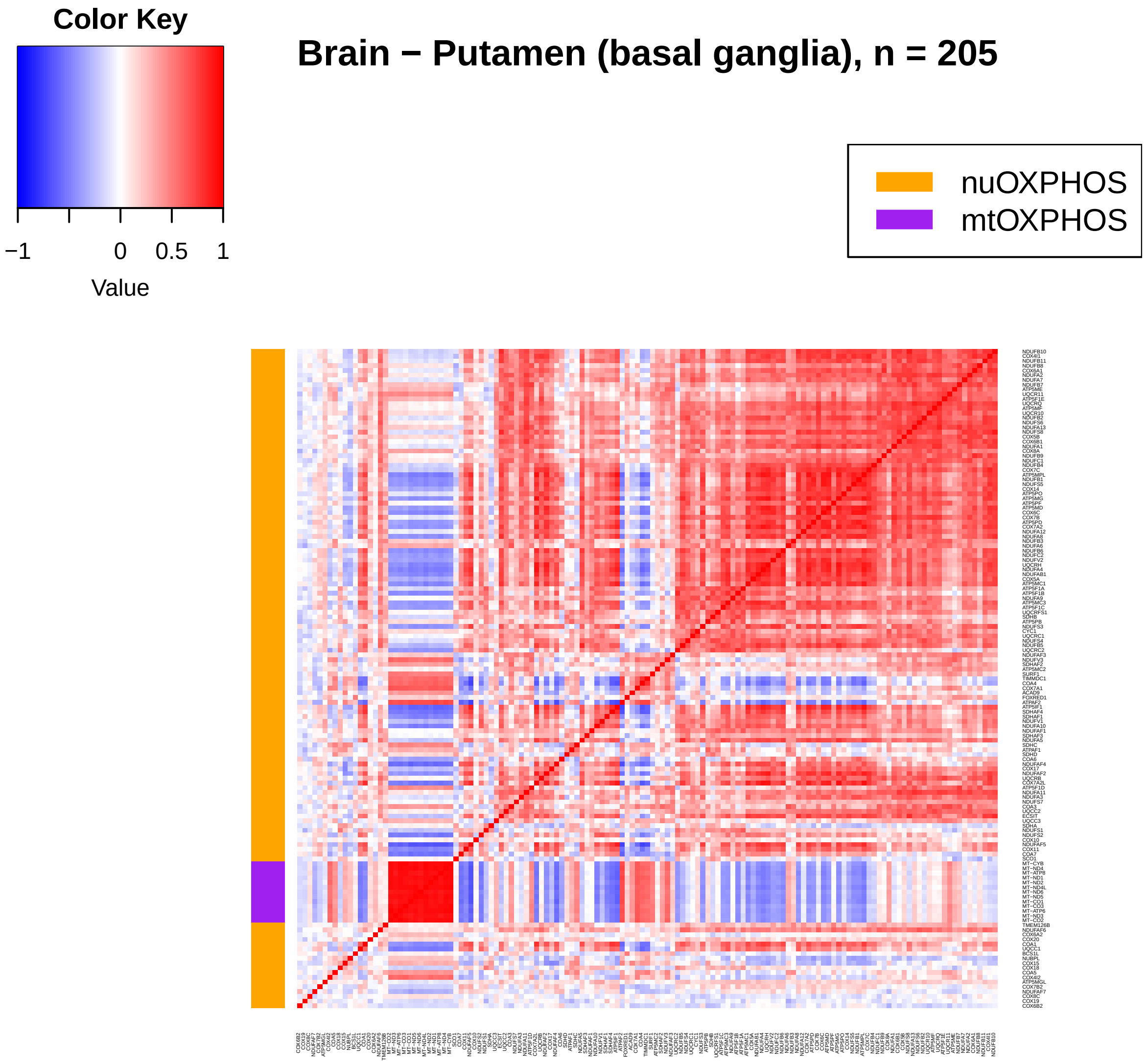
Color Key
−1
0
0.5
1
Value
Brain − Putamen (basal ganglia), n = 205
nuOXPHOS
mtOXPHOS

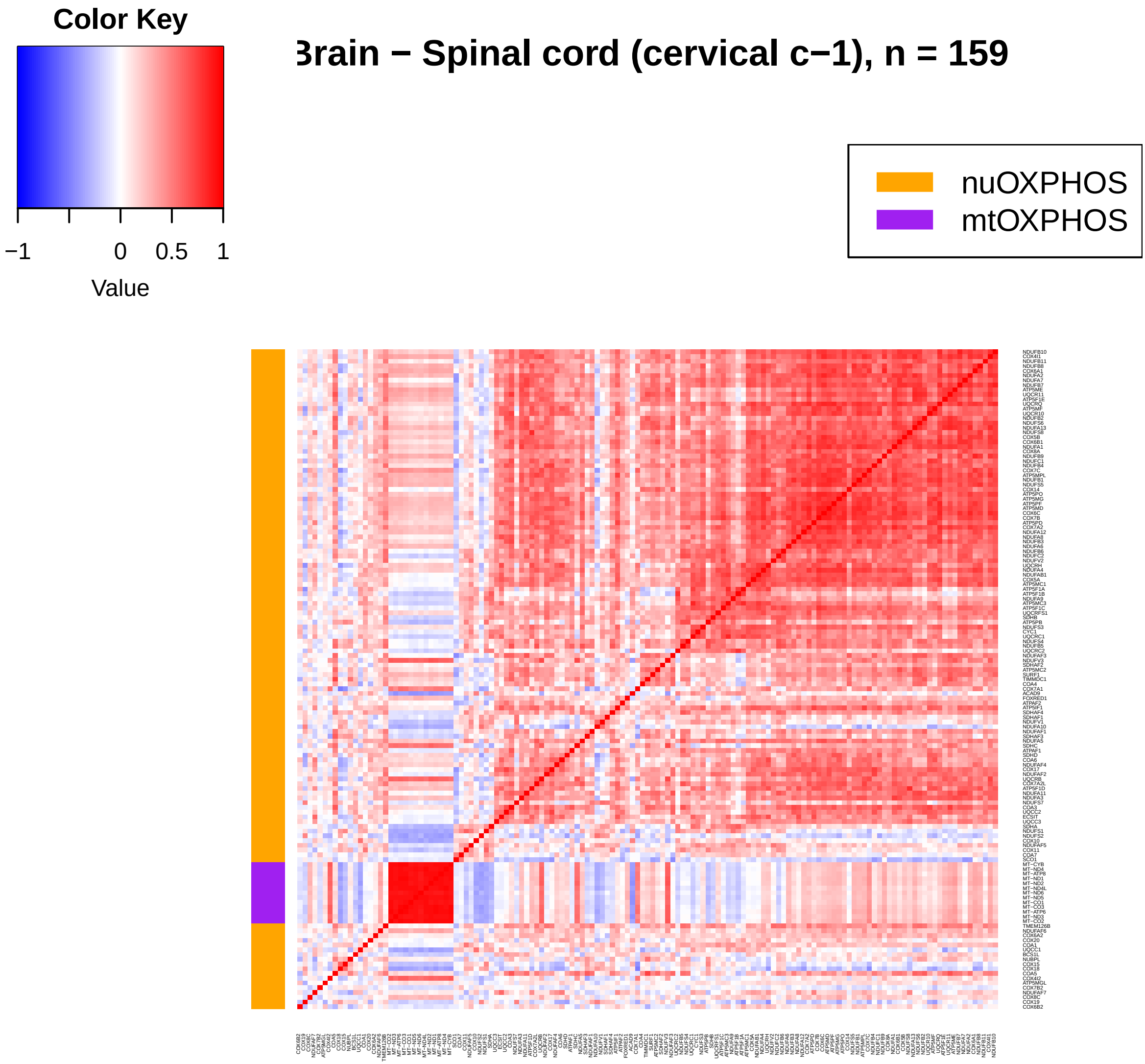
Color Key
−1
0
0.5
1
Value
rain − Spinal cord (cervical c−1), n = 159
nuOXPHOS
mtOXPHOS

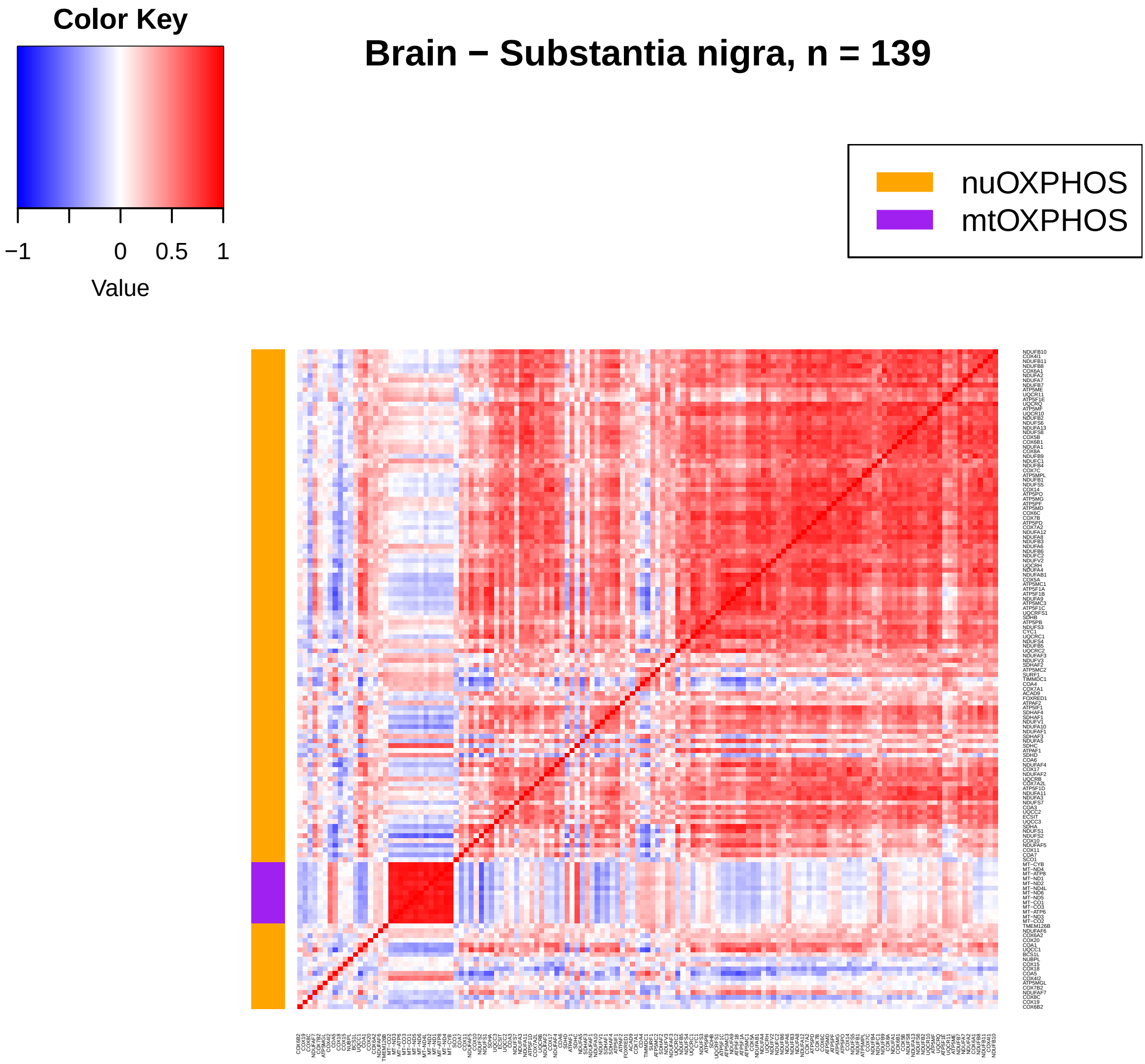
Color Key
−1
0
0.5
1
Value
Brain − Substantia nigra, n = 139
nuOXPHOS
mtOXPHOS

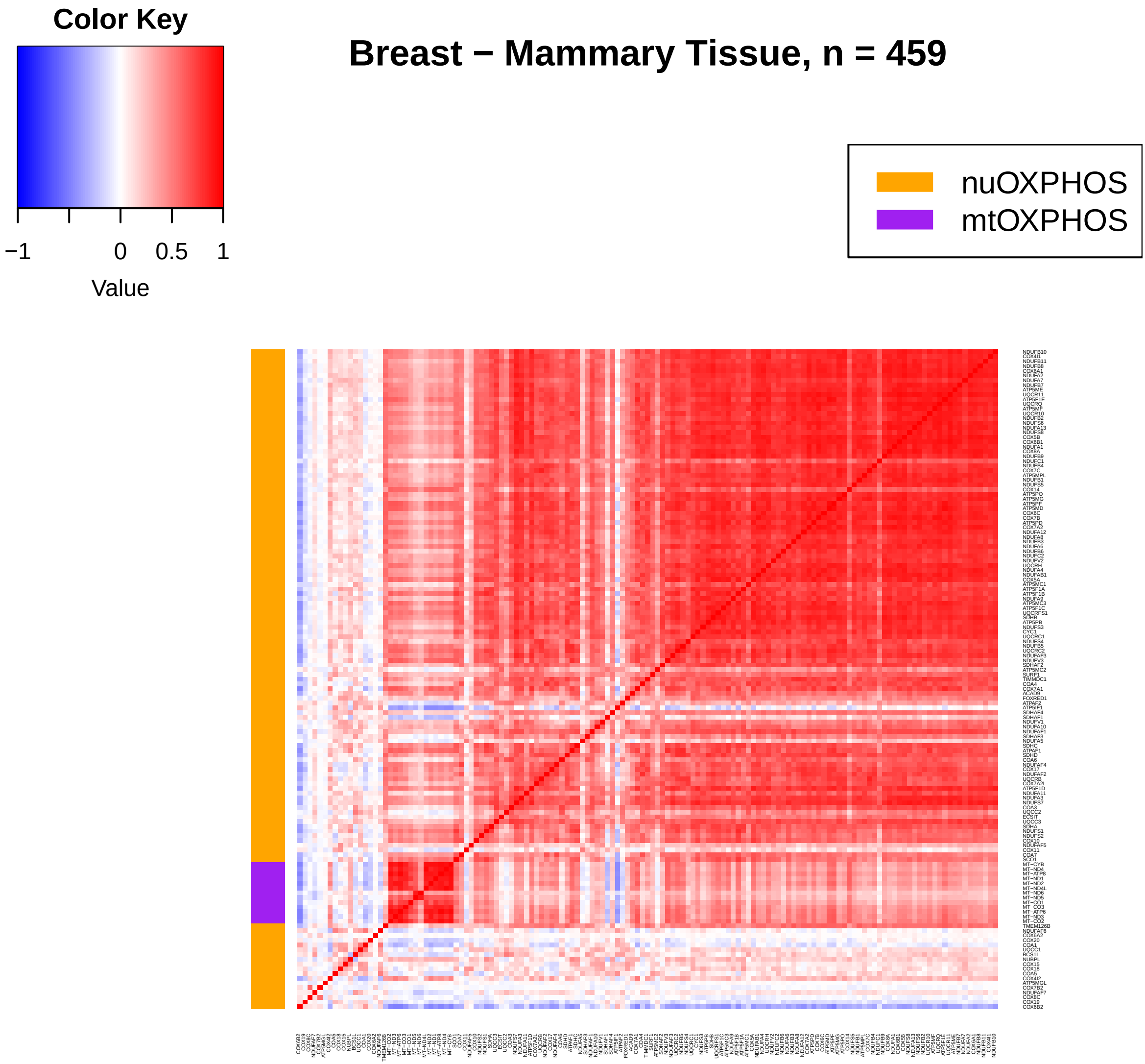
Color Key
−1
0
0.5
1
Value
Breast − Mammary Tissue, n = 459
nuOXPHOS
mtOXPHOS

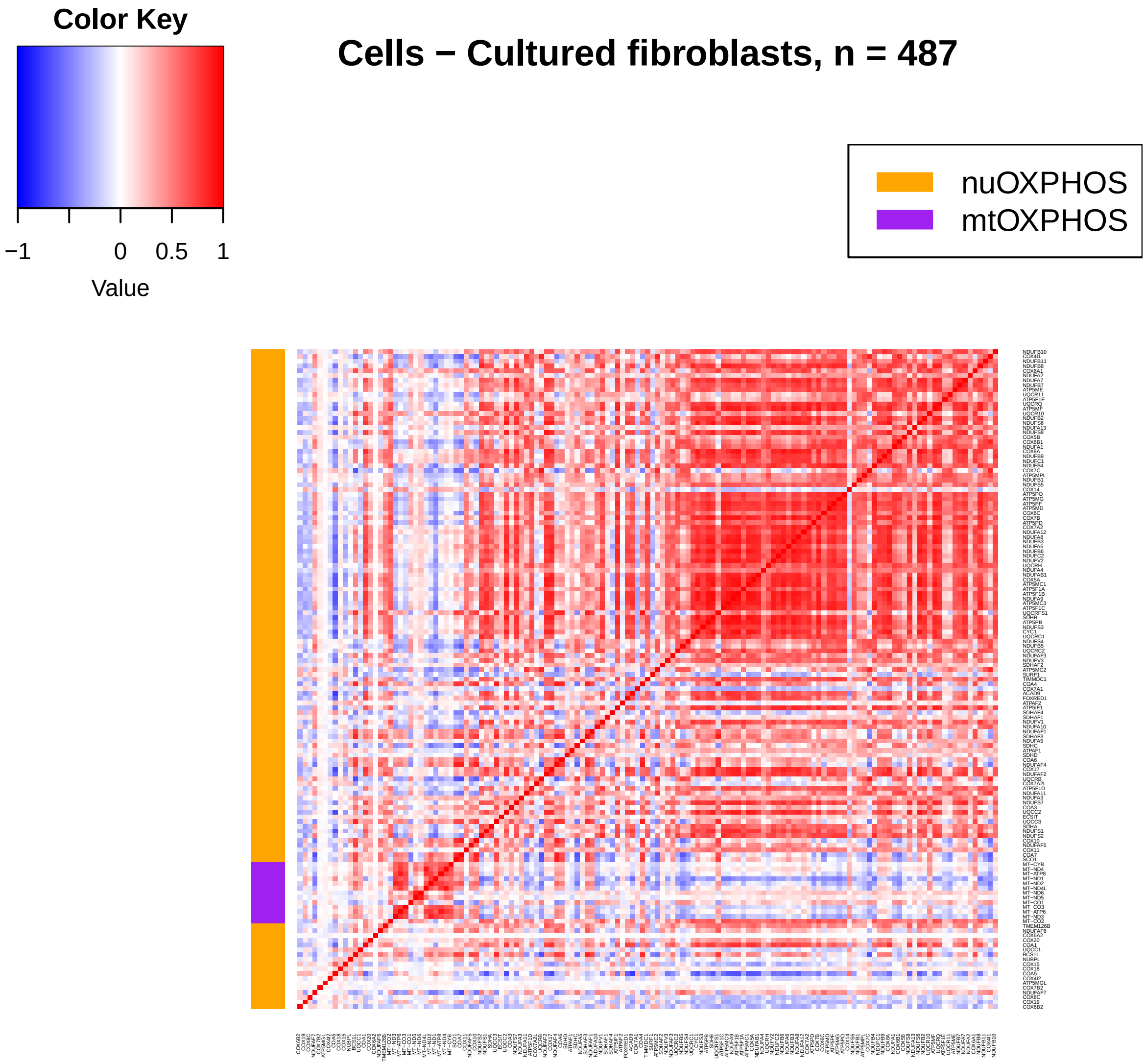
Color Key
−1
0
0.5
1
Value
Cells − Cultured fibroblasts, n = 487
nuOXPHOS
mtOXPHOS

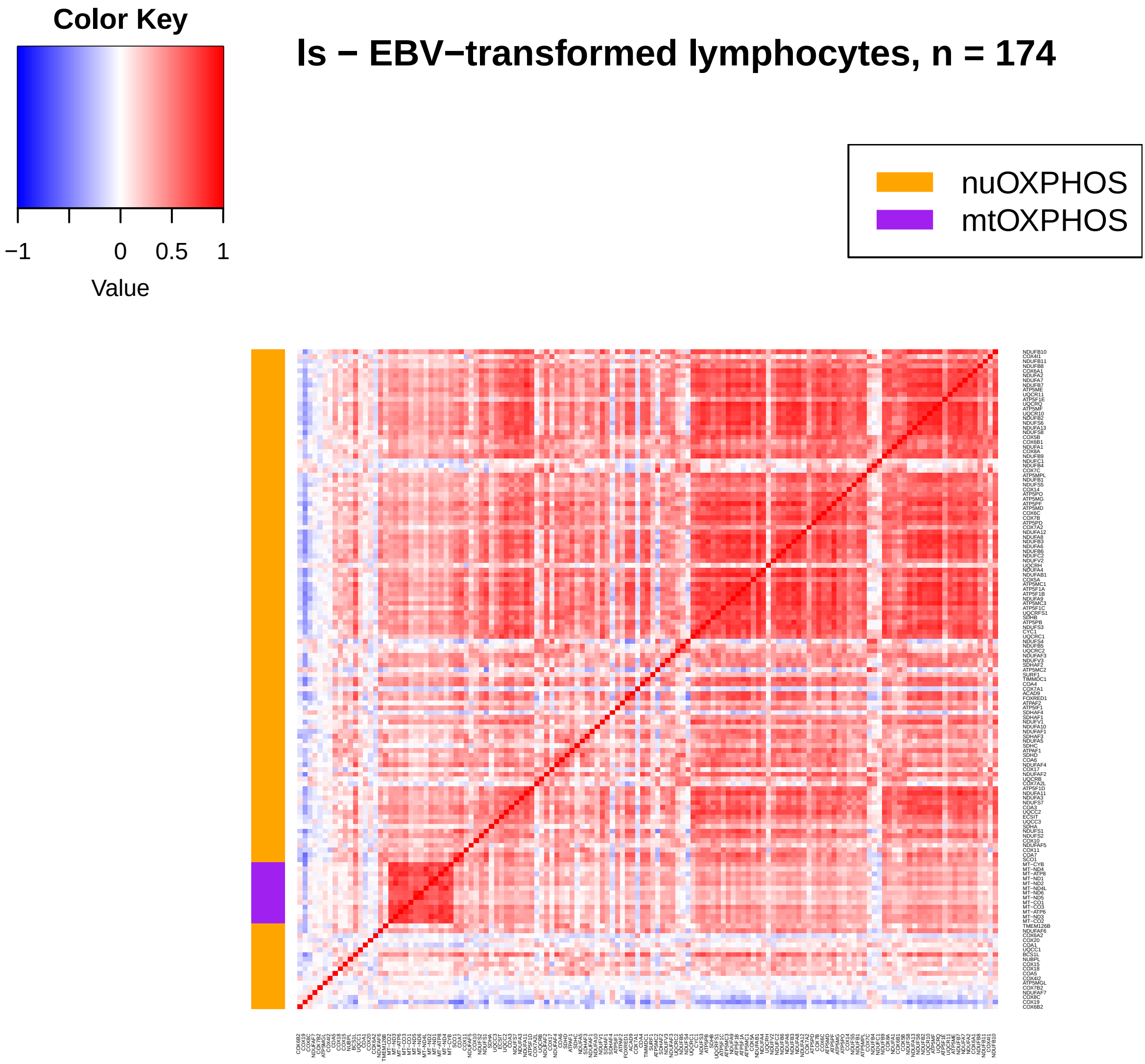
Color Key
−1
0
0.5
1
Value
ls − EBV−transformed lymphocytes, n = 174
nuOXPHOS
mtOXPHOS

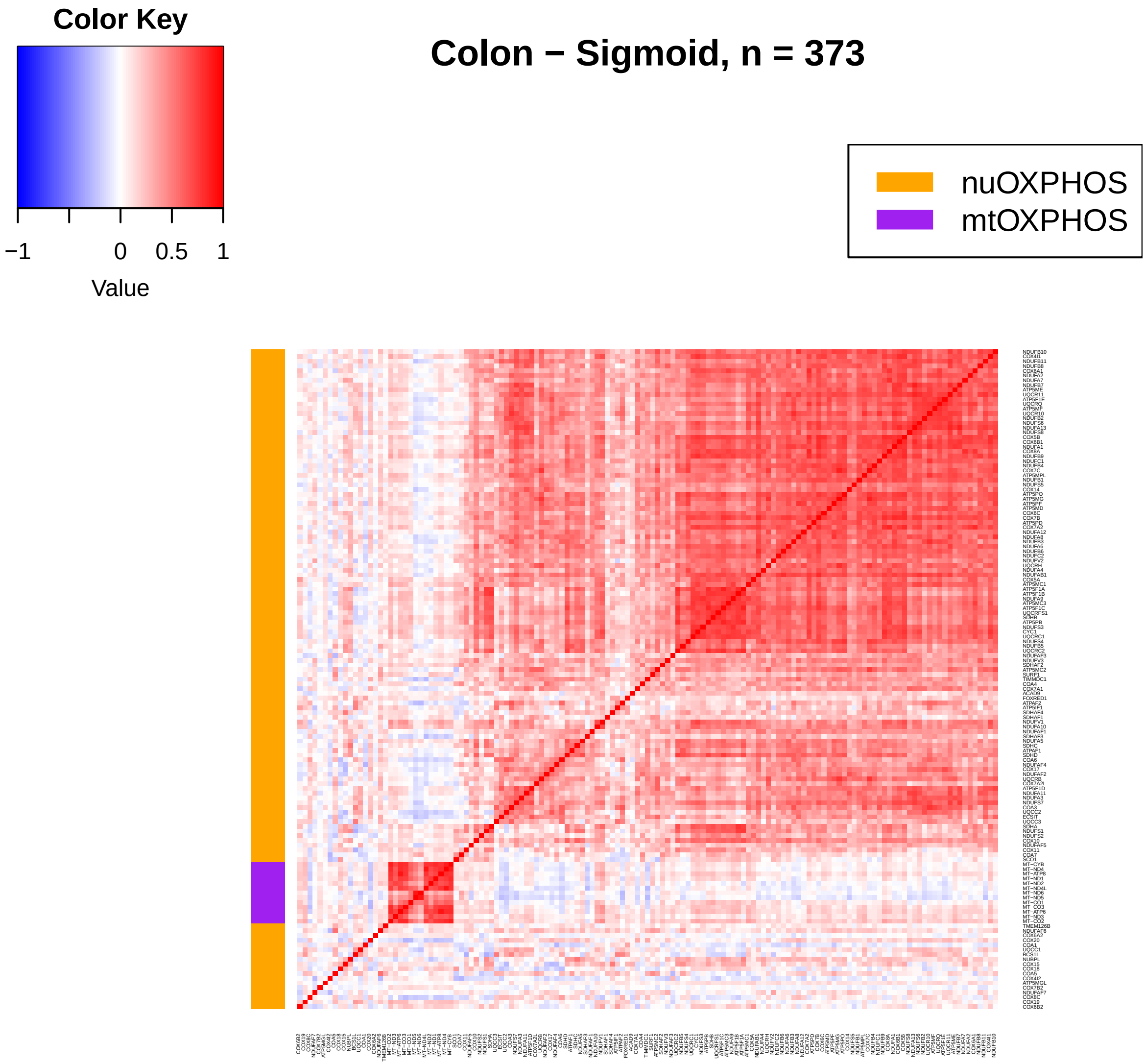
Color Key
−1
0
0.5
1
Value
Colon − Sigmoid, n = 373
nuOXPHOS
mtOXPHOS

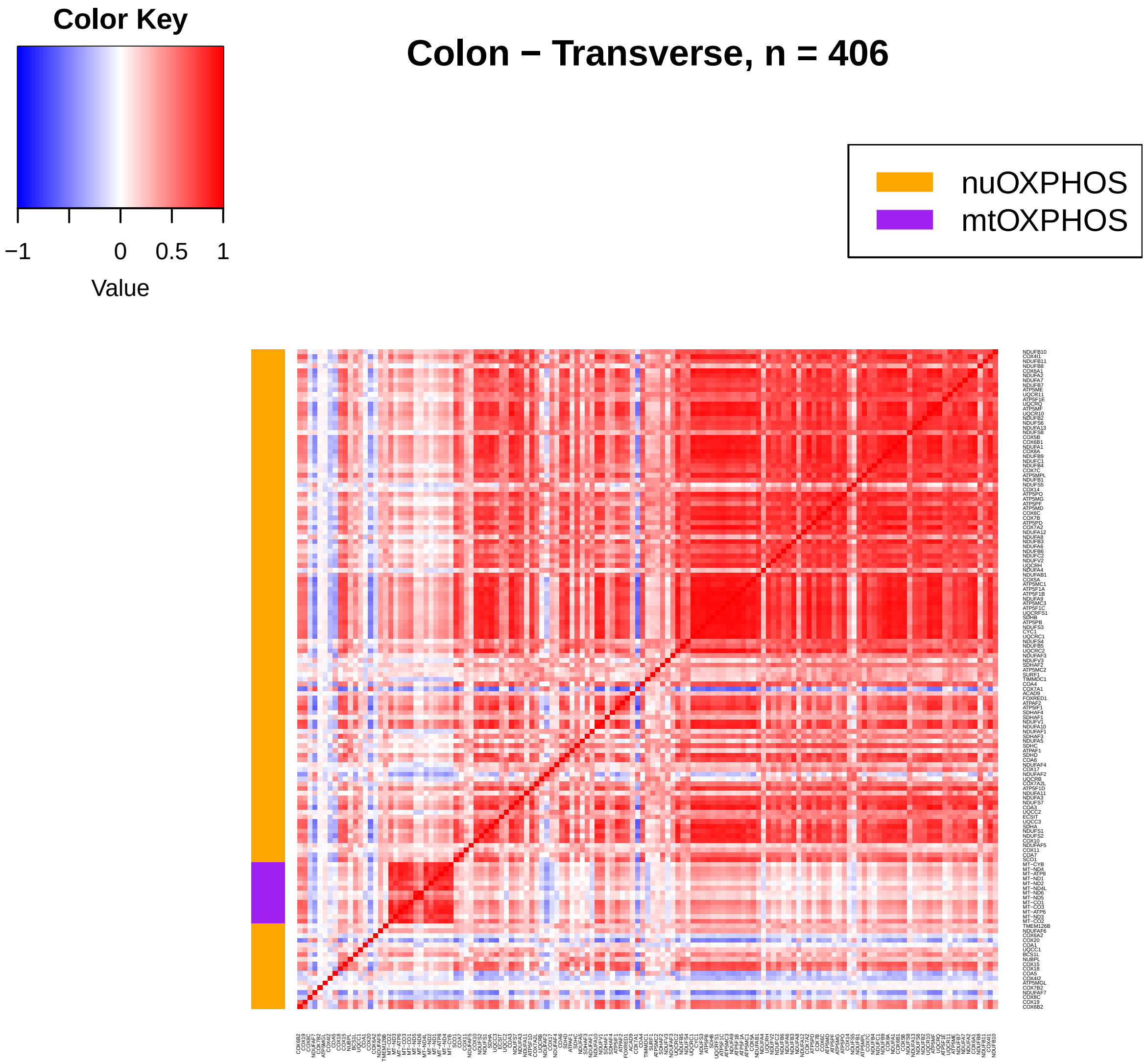
Color Key
−1
0
0.5
1
Value
Colon − Transverse, n = 406
nuOXPHOS
mtOXPHOS

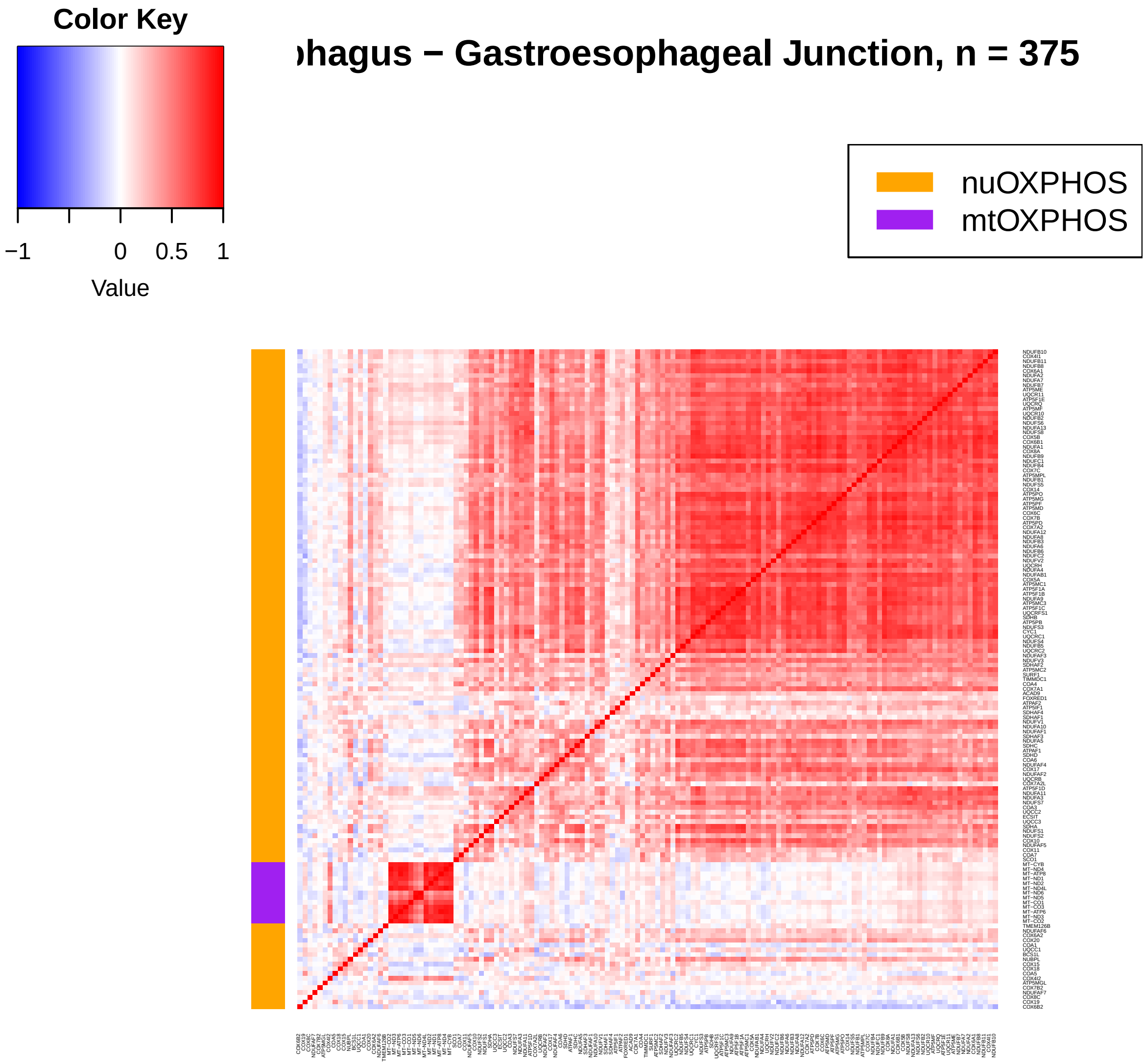
Color Key
−1
0
0.5
1
Value
ohagus − Gastroesophageal Junction, n = 375
nuOXPHOS
mtOXPHOS

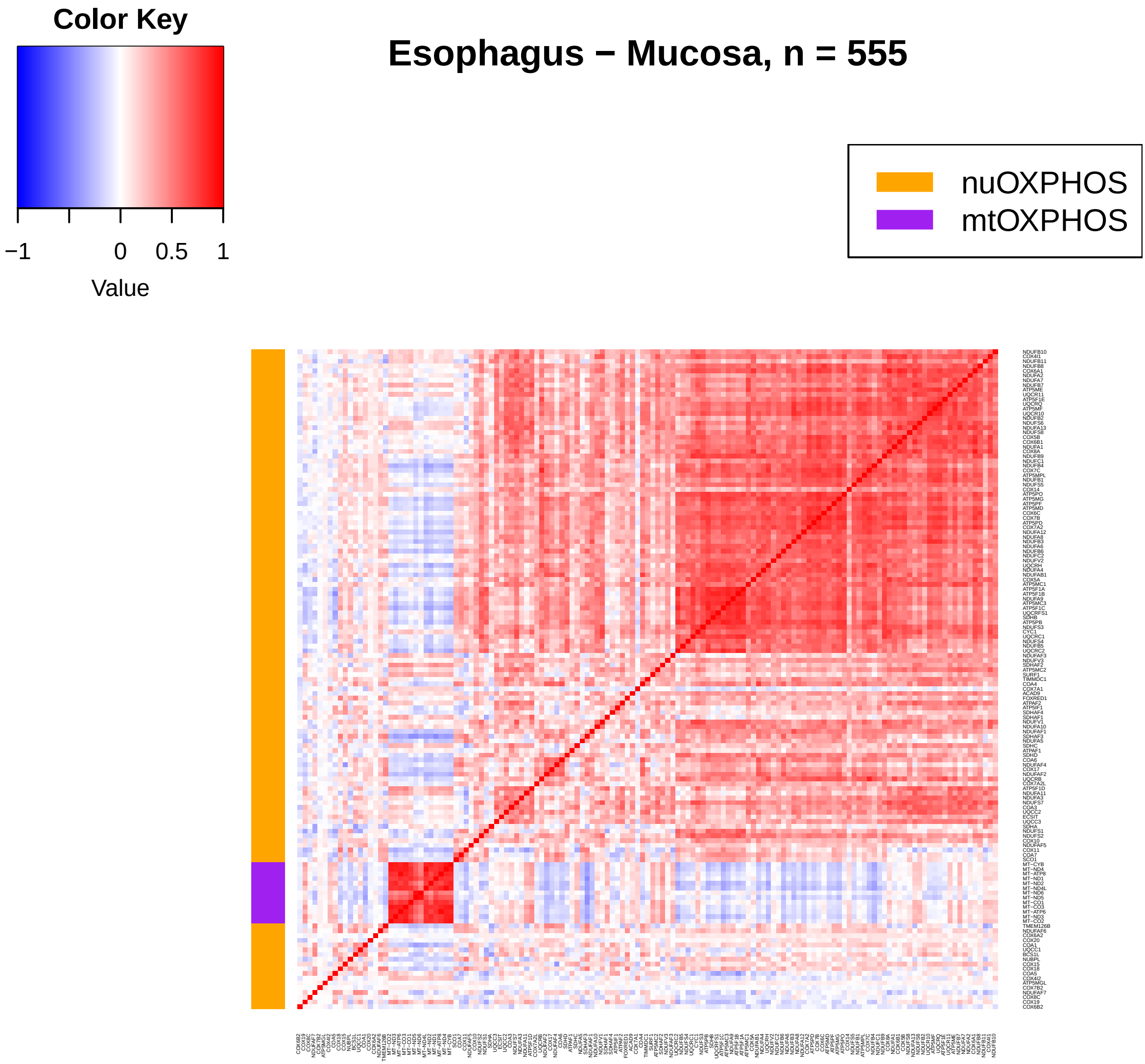
Color Key
−1
0
0.5
1
Value
Esophagus − Mucosa, n = 555
nuOXPHOS
mtOXPHOS

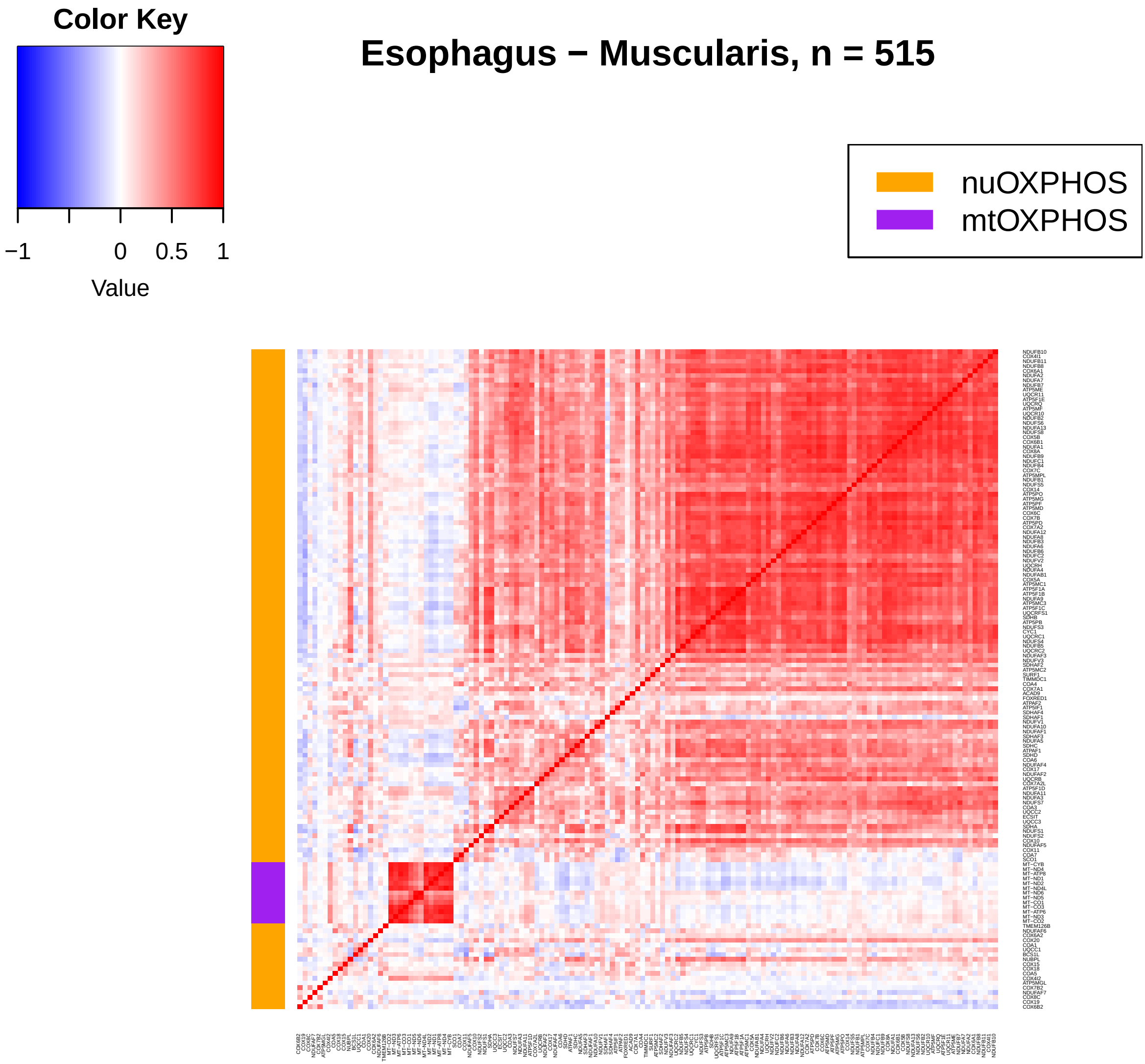
Color Key
−1
0
0.5
1
Value
Esophagus − Muscularis, n = 515
nuOXPHOS
mtOXPHOS

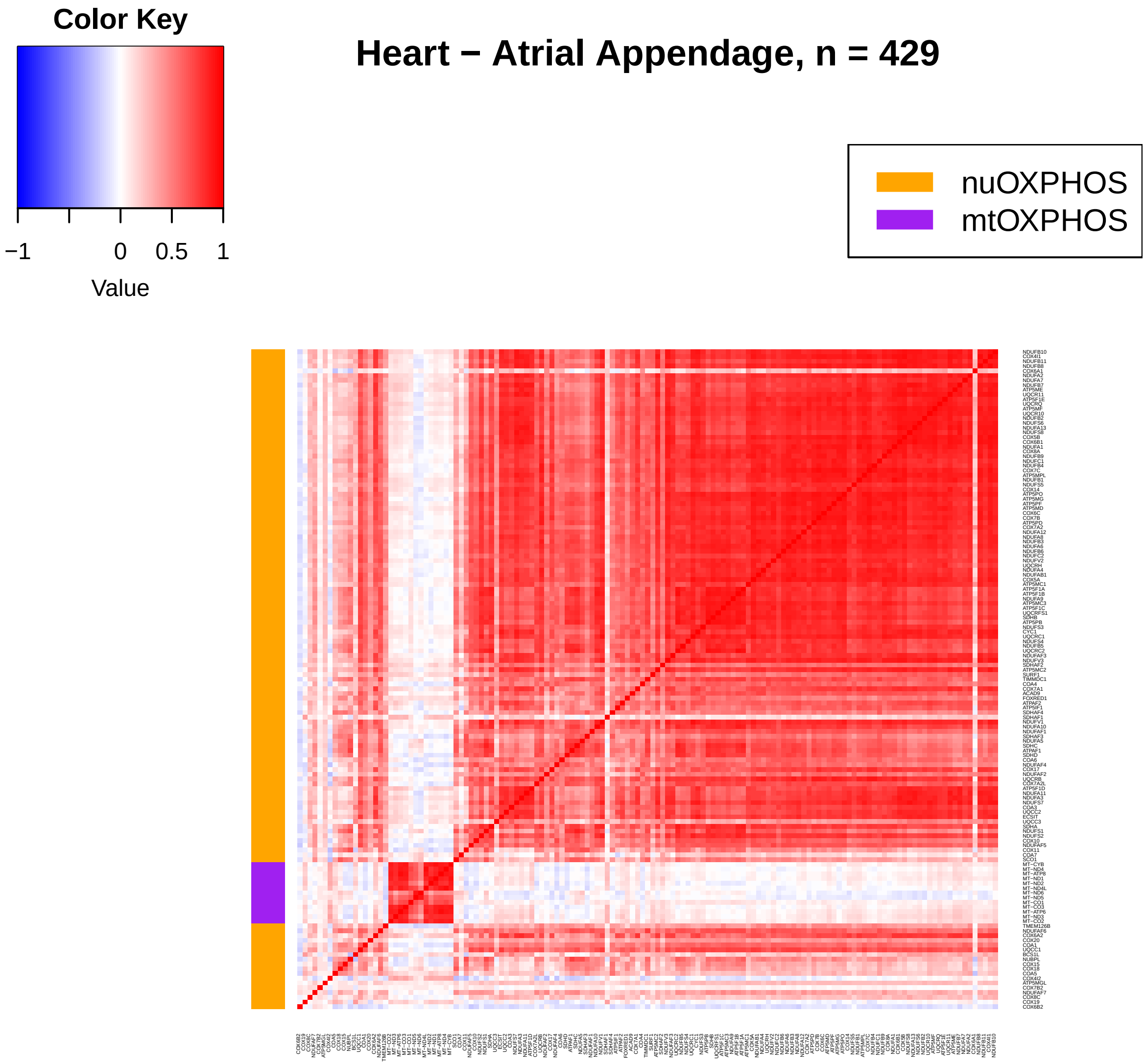
Color Key
−1
0
0.5
1
Value
Heart − Atrial Appendage, n = 429
nuOXPHOS
mtOXPHOS

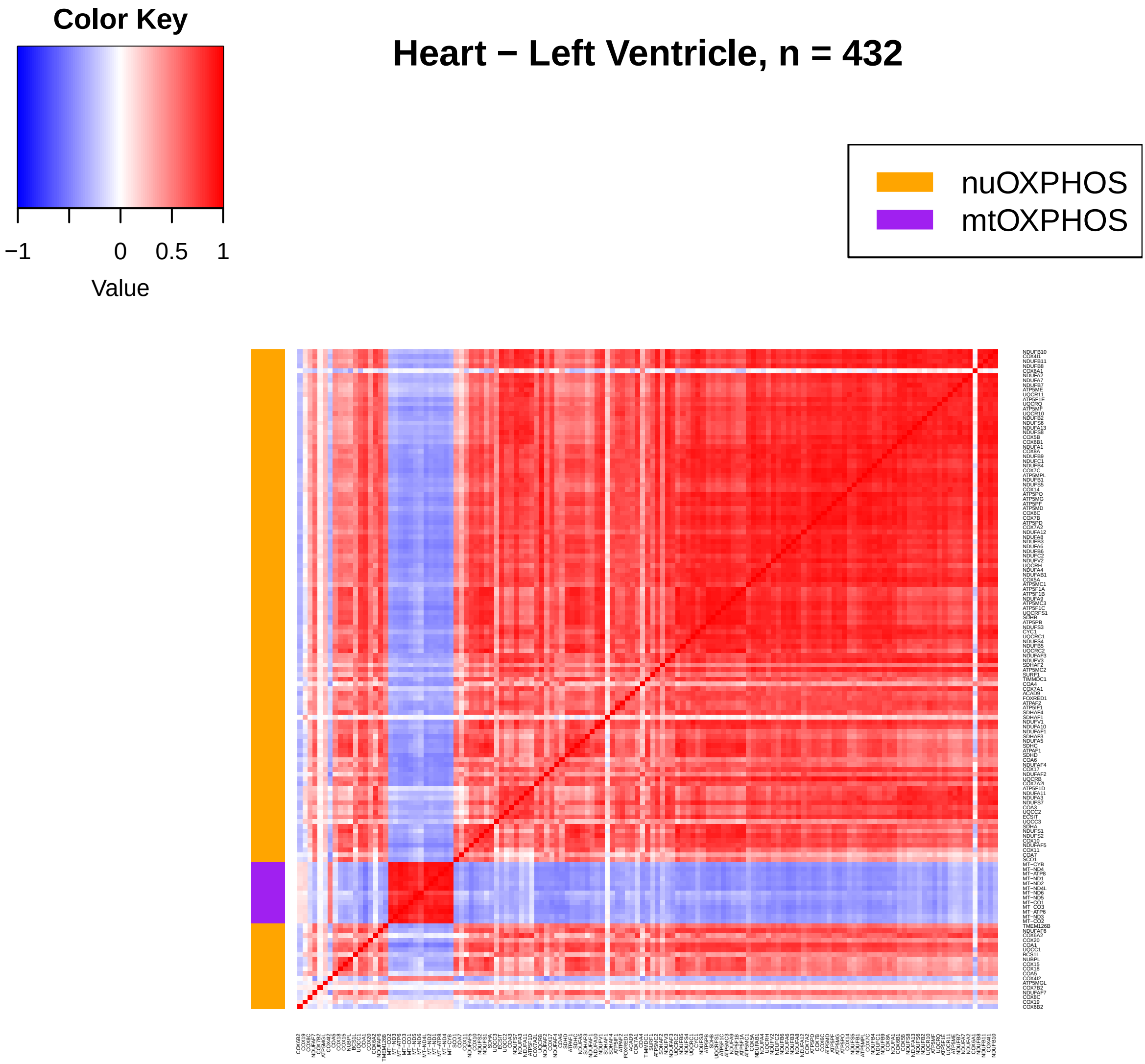
Color Key
−1
0
0.5
1
Value
Heart − Left Ventricle, n = 432
nuOXPHOS
mtOXPHOS

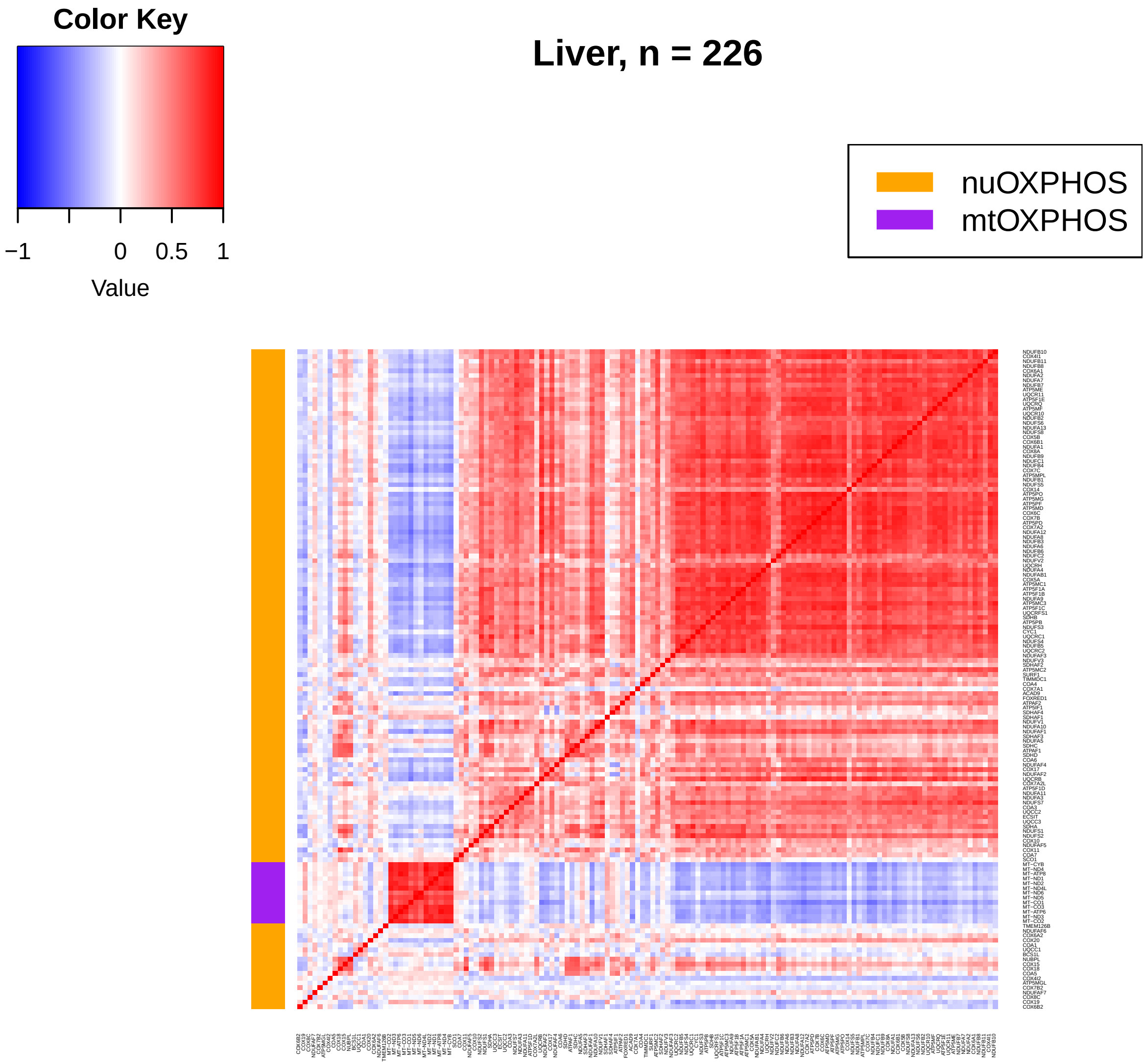
Color Key
−1
0
0.5
1
Value
Liver, n = 226
nuOXPHOS
mtOXPHOS

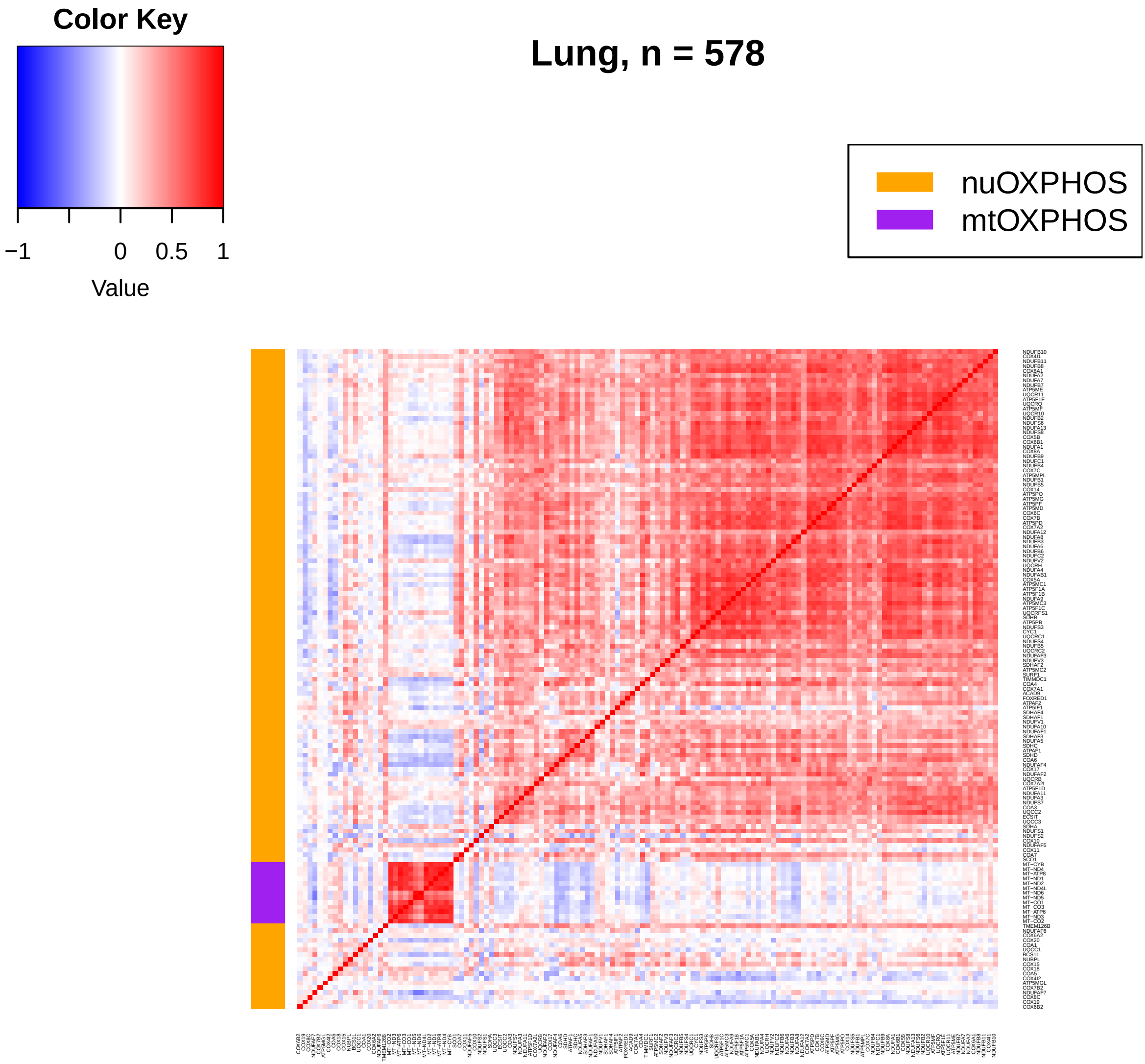
Color Key
−1
0
0.5
1
Value
Lung, n = 578
nuOXPHOS
mtOXPHOS

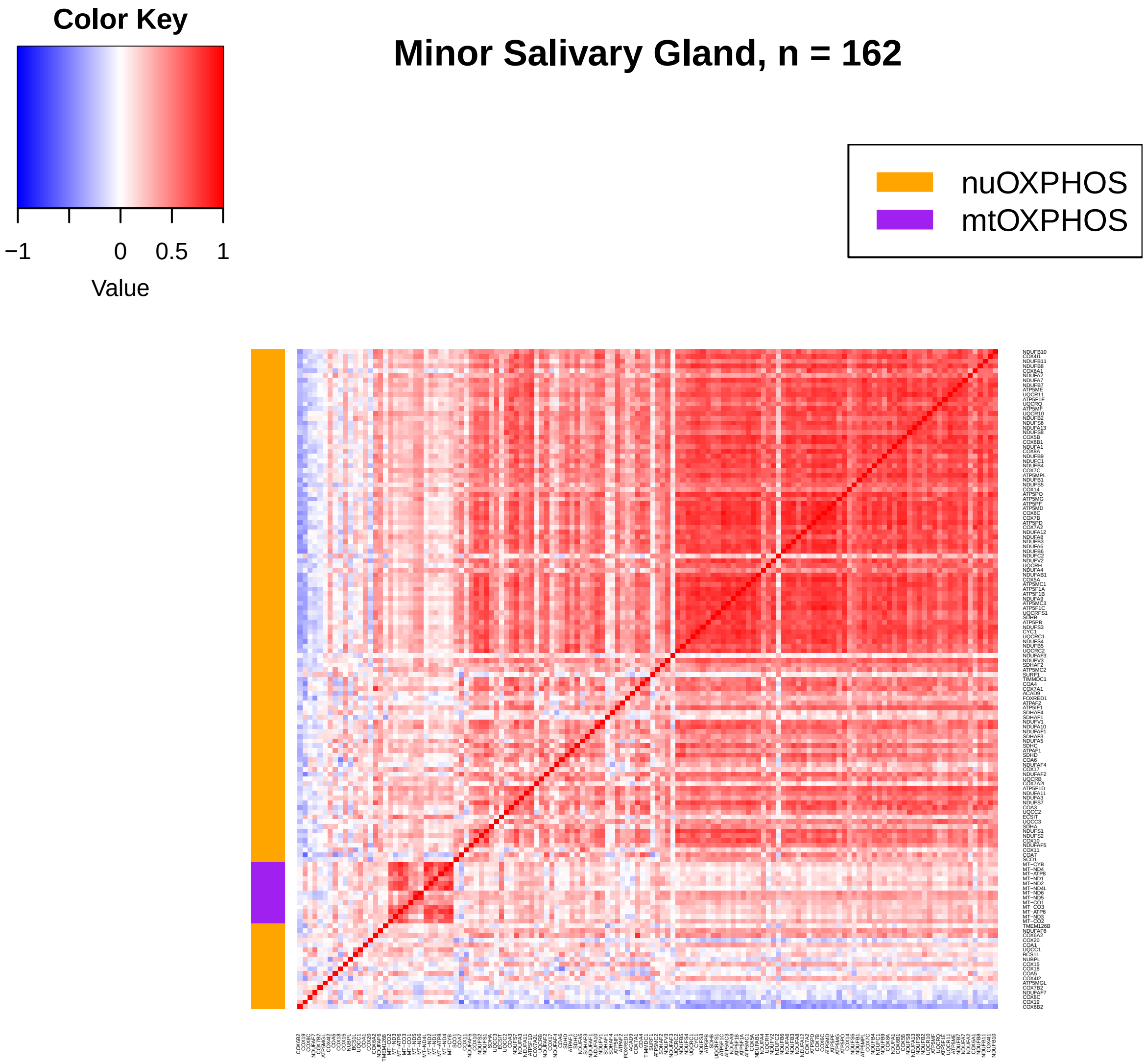
Color Key
−1
0
0.5
1
Value
Minor Salivary Gland, n = 162
nuOXPHOS
mtOXPHOS

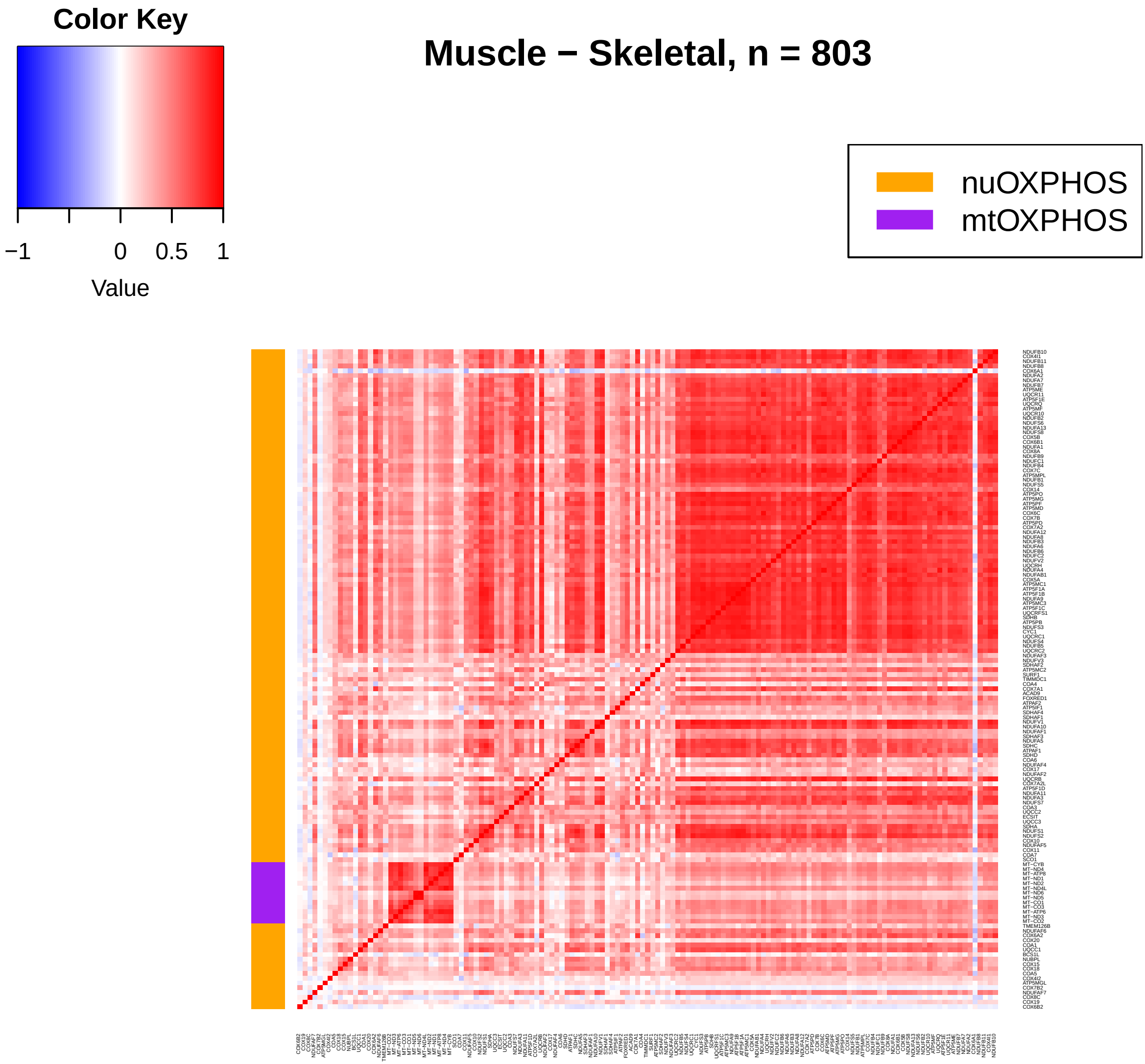
Color Key
−1
0
0.5
1
Value
Muscle − Skeletal, n = 803
nuOXPHOS
mtOXPHOS

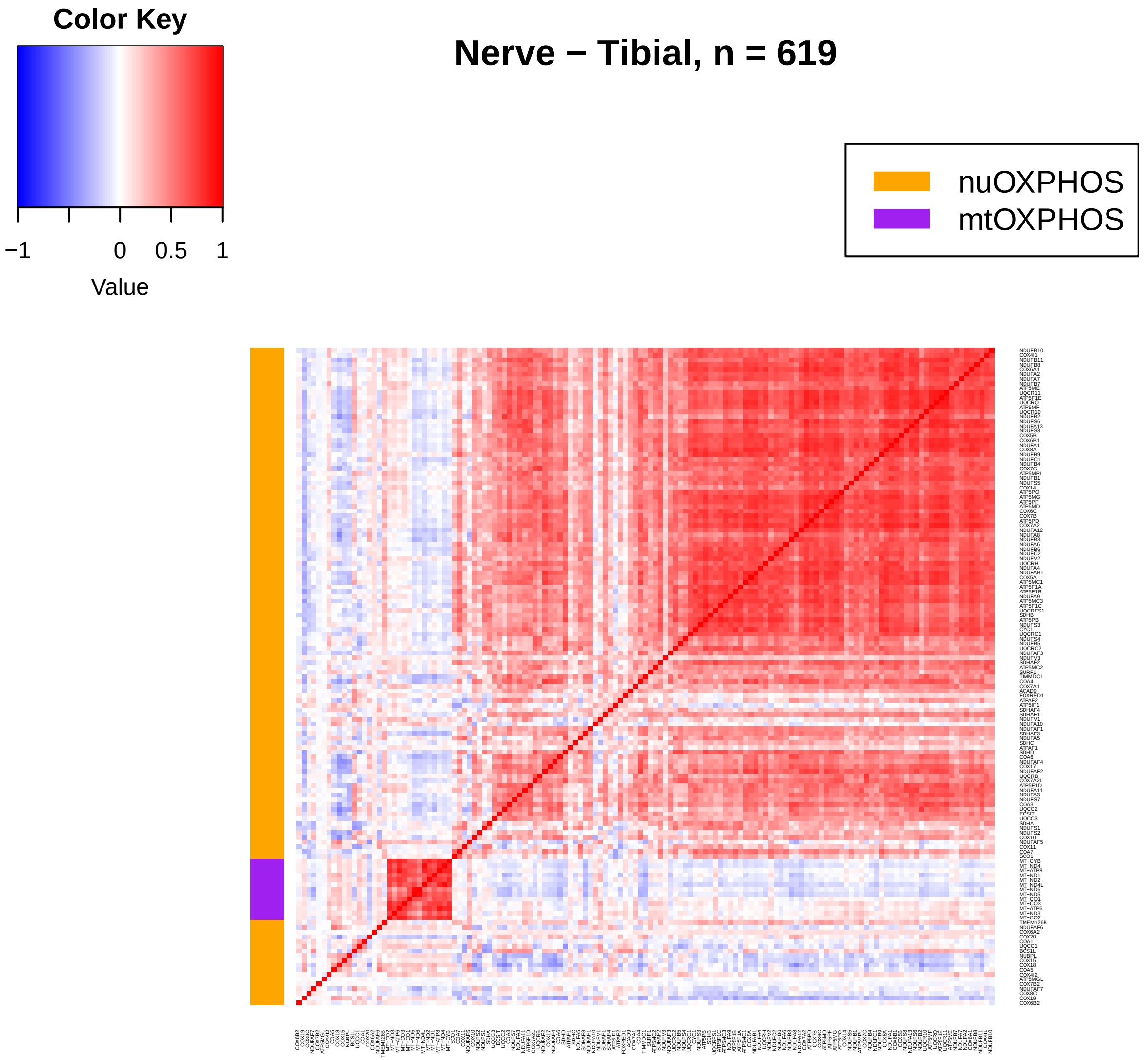
Color Key
−1
0
0.5
1
Value
Nerve − Tibial, n = 619
nuOXPHOS
mtOXPHOS

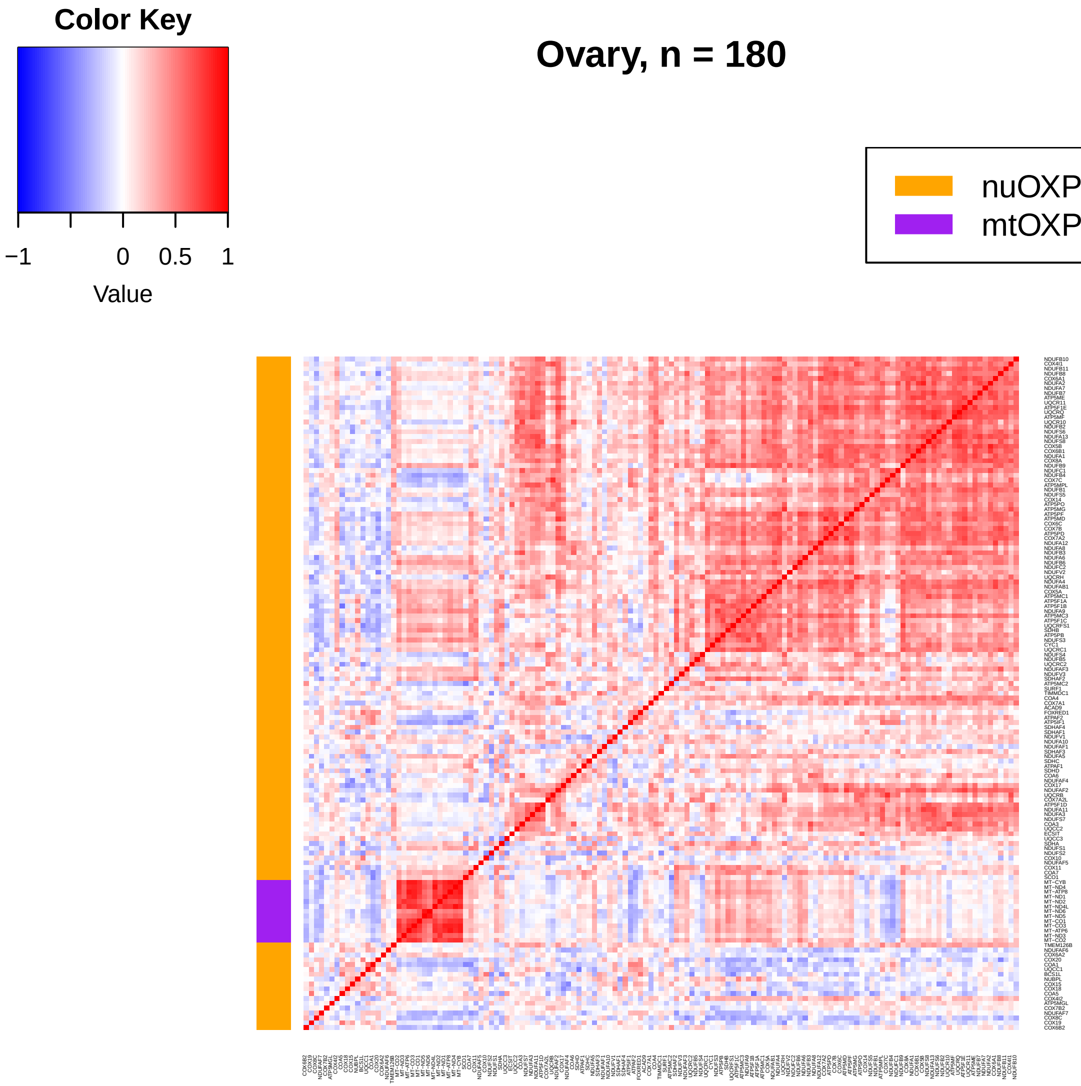
Color Key
−1
0
0.5
1
Value
Ovary, n = 180
nuOXPHOS
mtOXPHOS

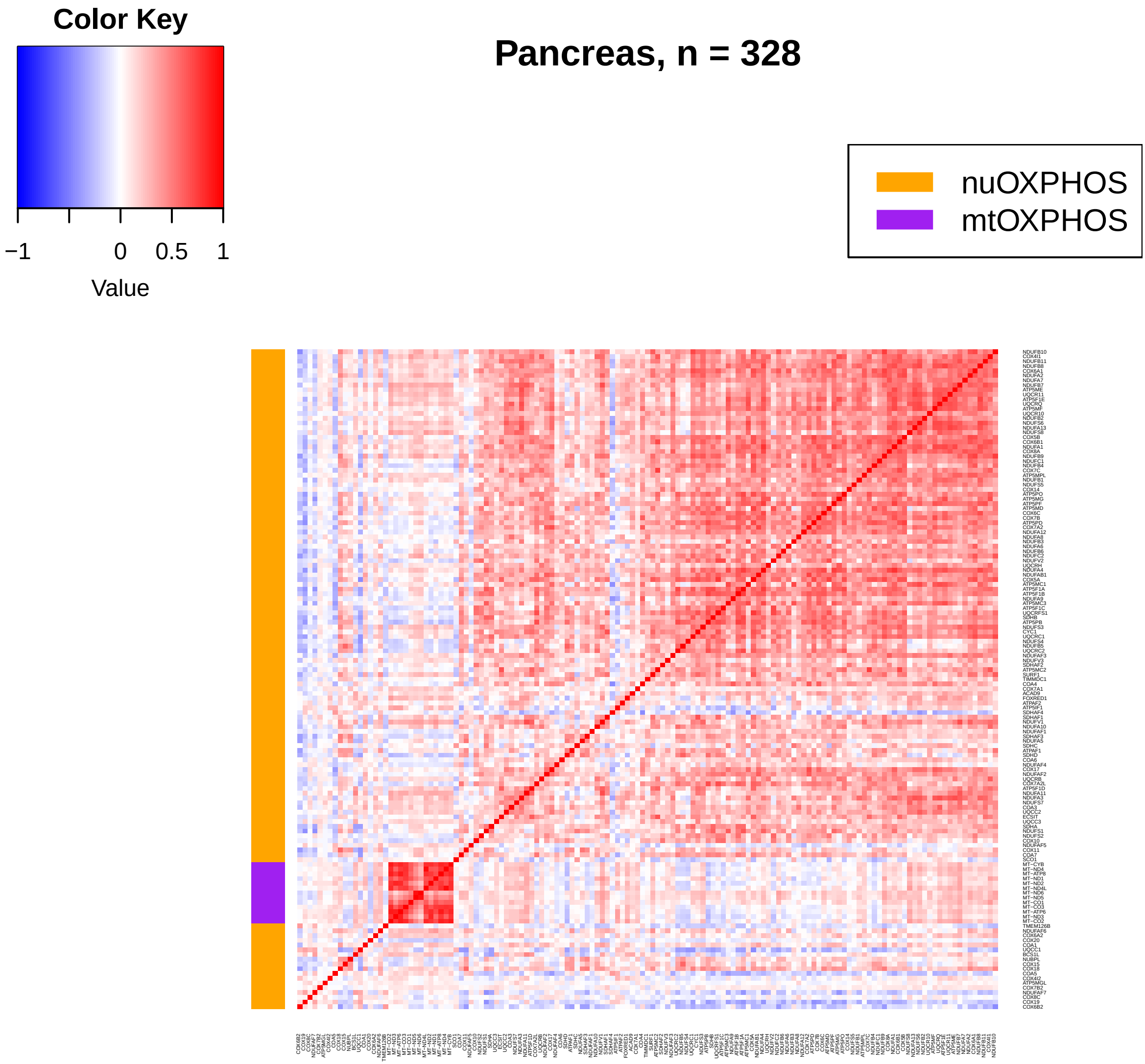
Color Key
−1
0
0.5
1
Value
Pancreas, n = 328
nuOXPHOS
mtOXPHOS

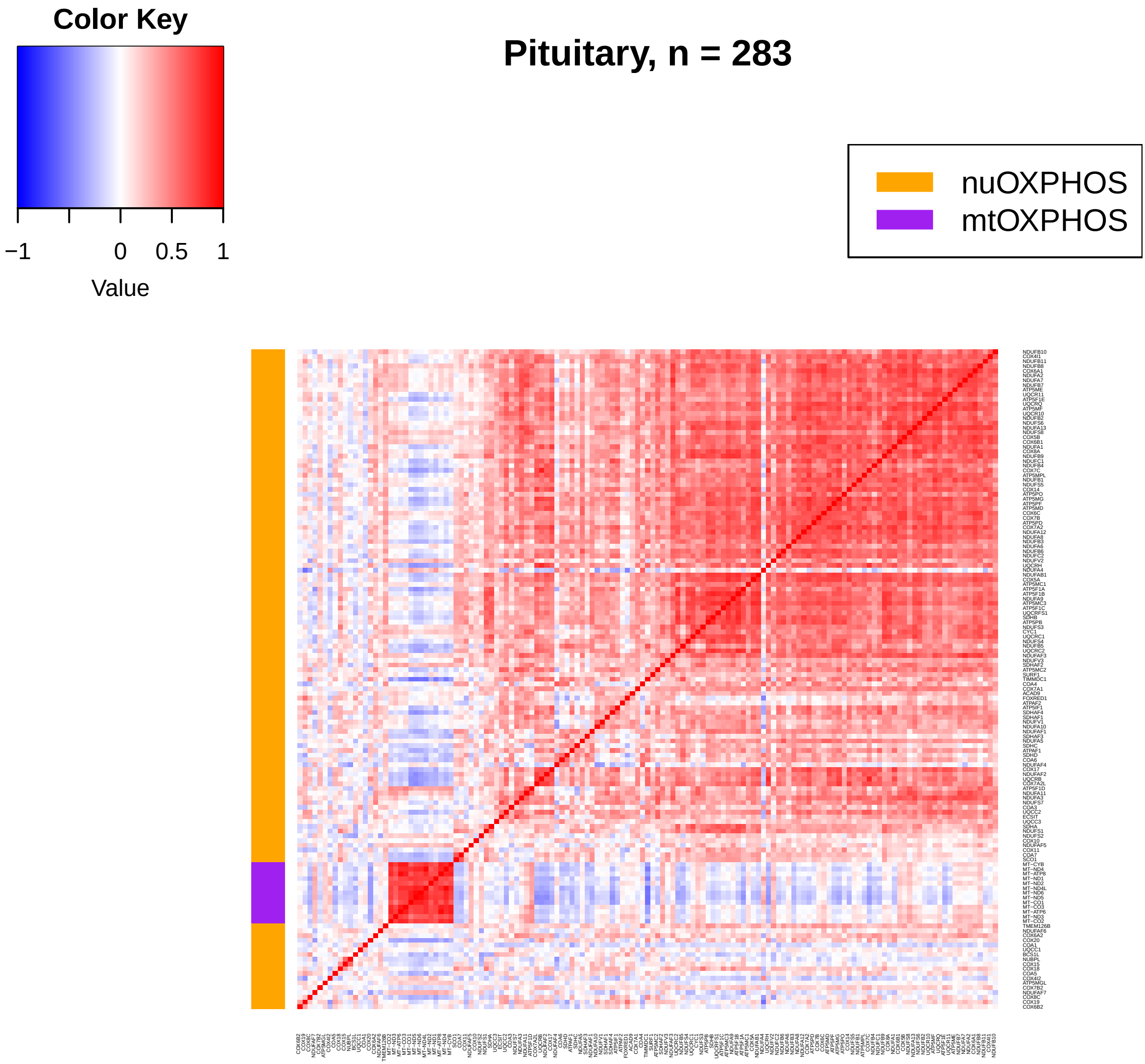
Color Key
−1
0
0.5
1
Value
Pituitary, n = 283
nuOXPHOS
mtOXPHOS

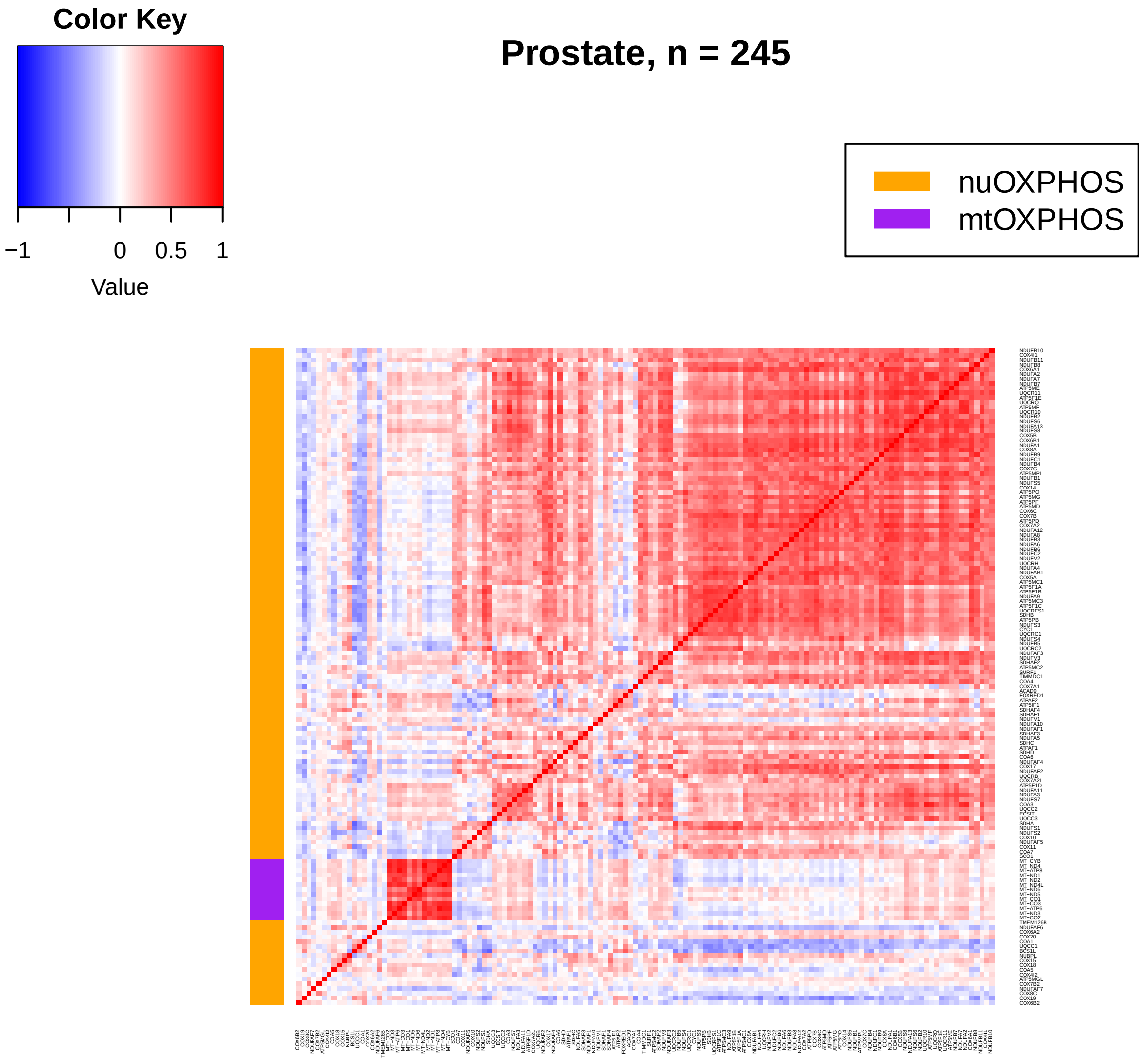
Color Key
−1
0
0.5
1
Value
Prostate, n = 245
nuOXPHOS
mtOXPHOS

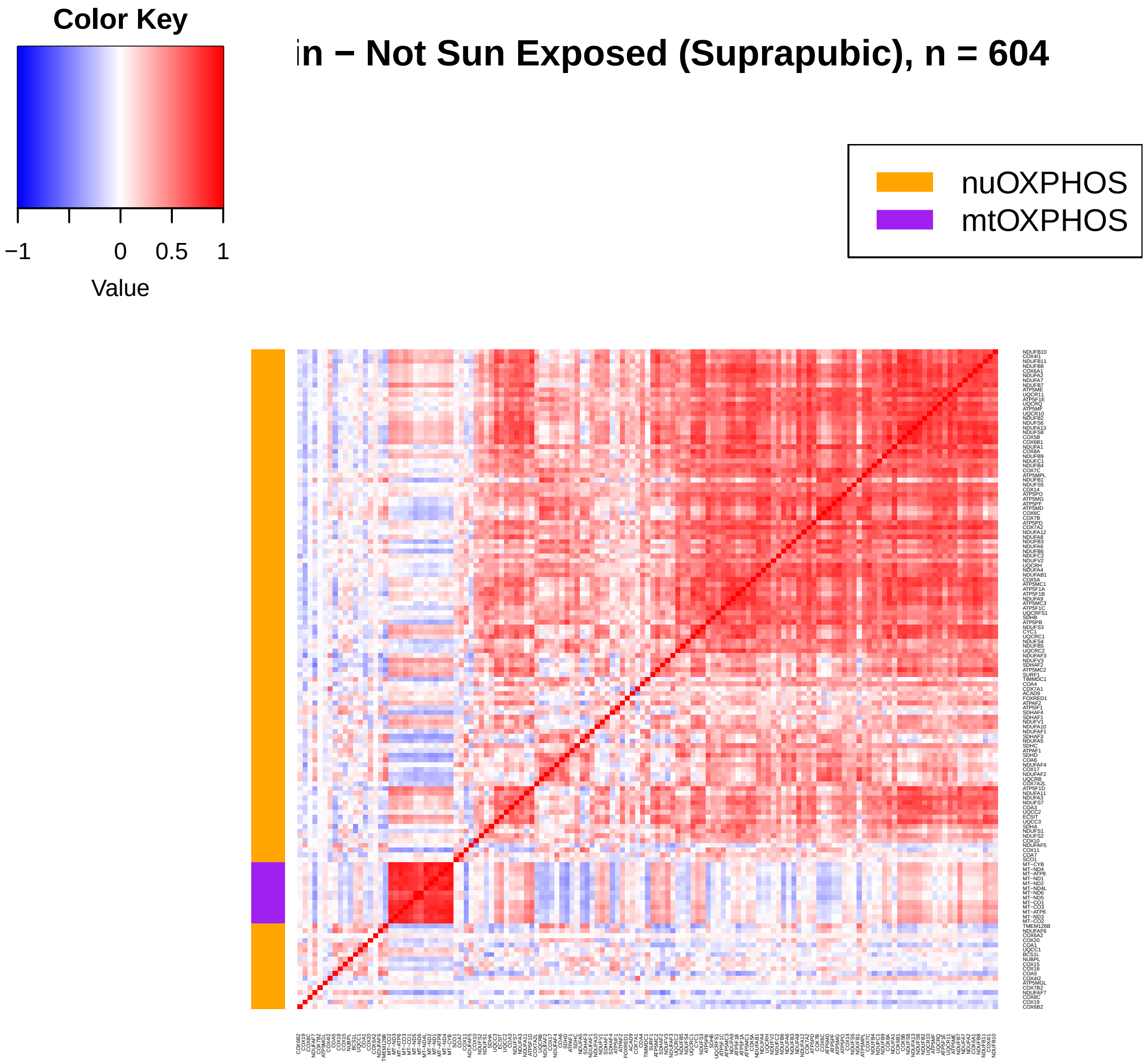
Color Key
−1
0
0.5
1
Value
in − Not Sun Exposed (Suprapubic), n = 604
nuOXPHOS
mtOXPHOS

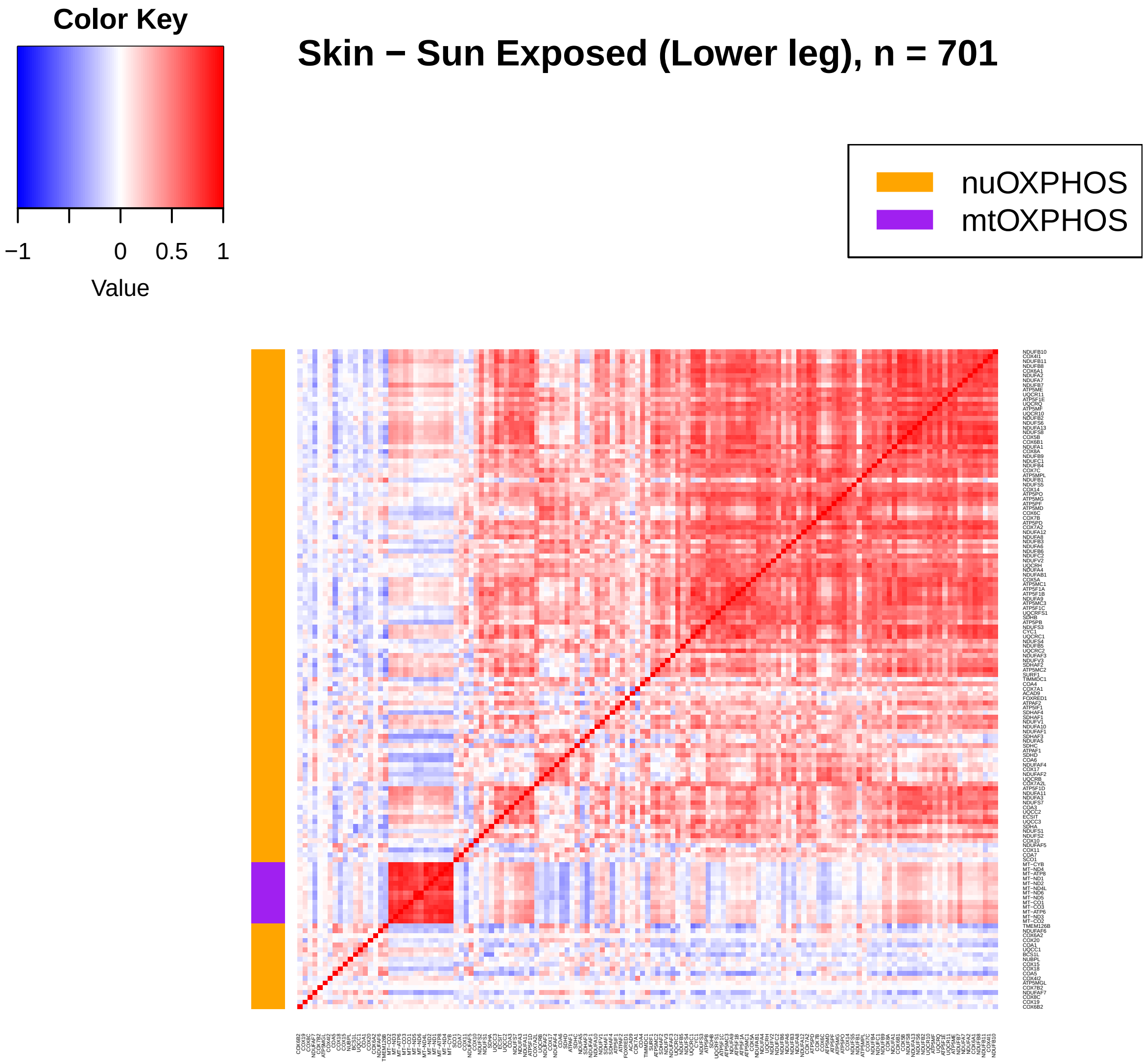
Color Key
−1
0
0.5
1
Value
Skin − Sun Exposed (Lower leg), n = 701
nuOXPHOS
mtOXPHOS

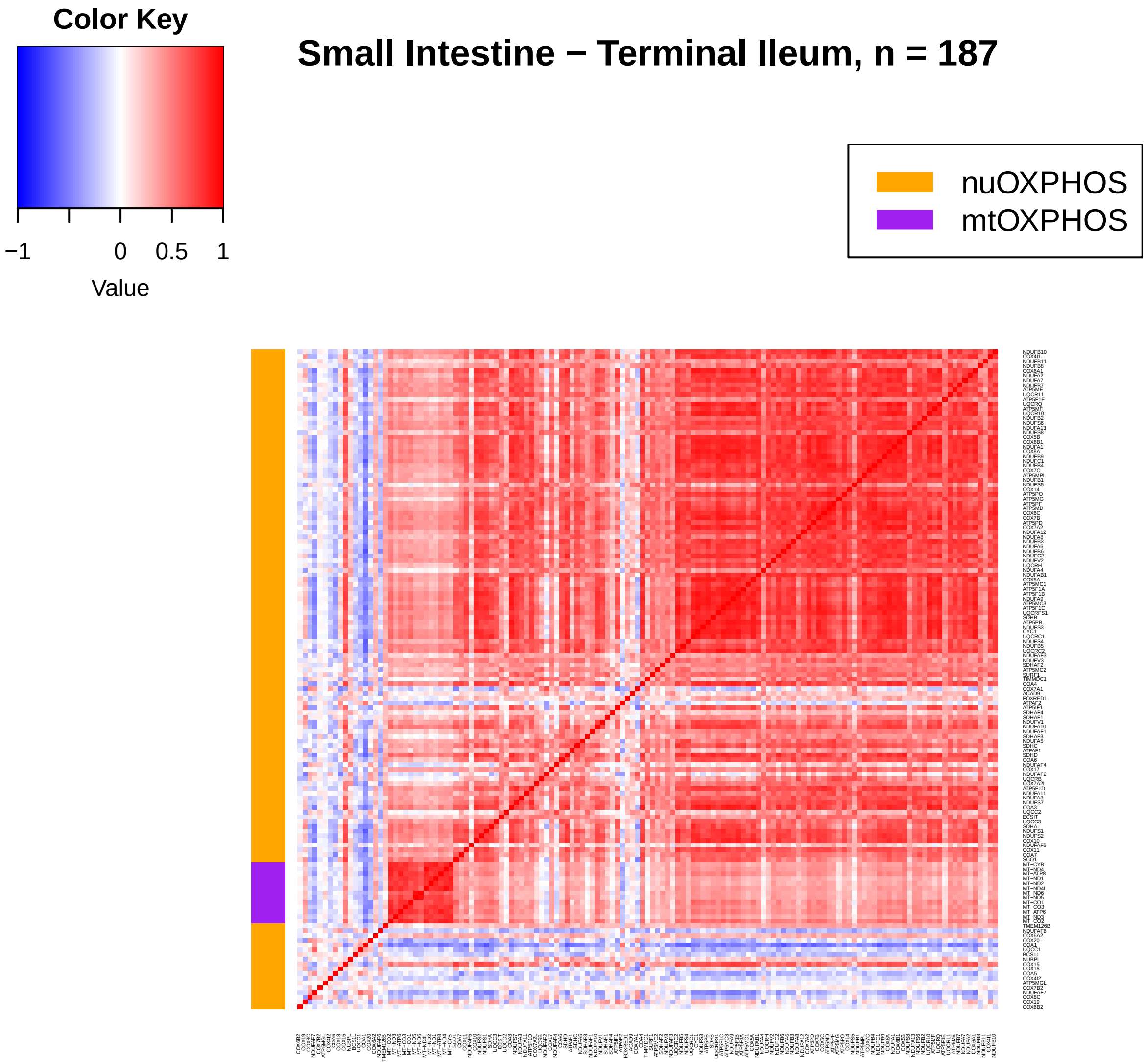
Color Key
−1
0
0.5
1
Value
Small Intestine − Terminal Ileum, n = 187
nuOXPHOS
mtOXPHOS

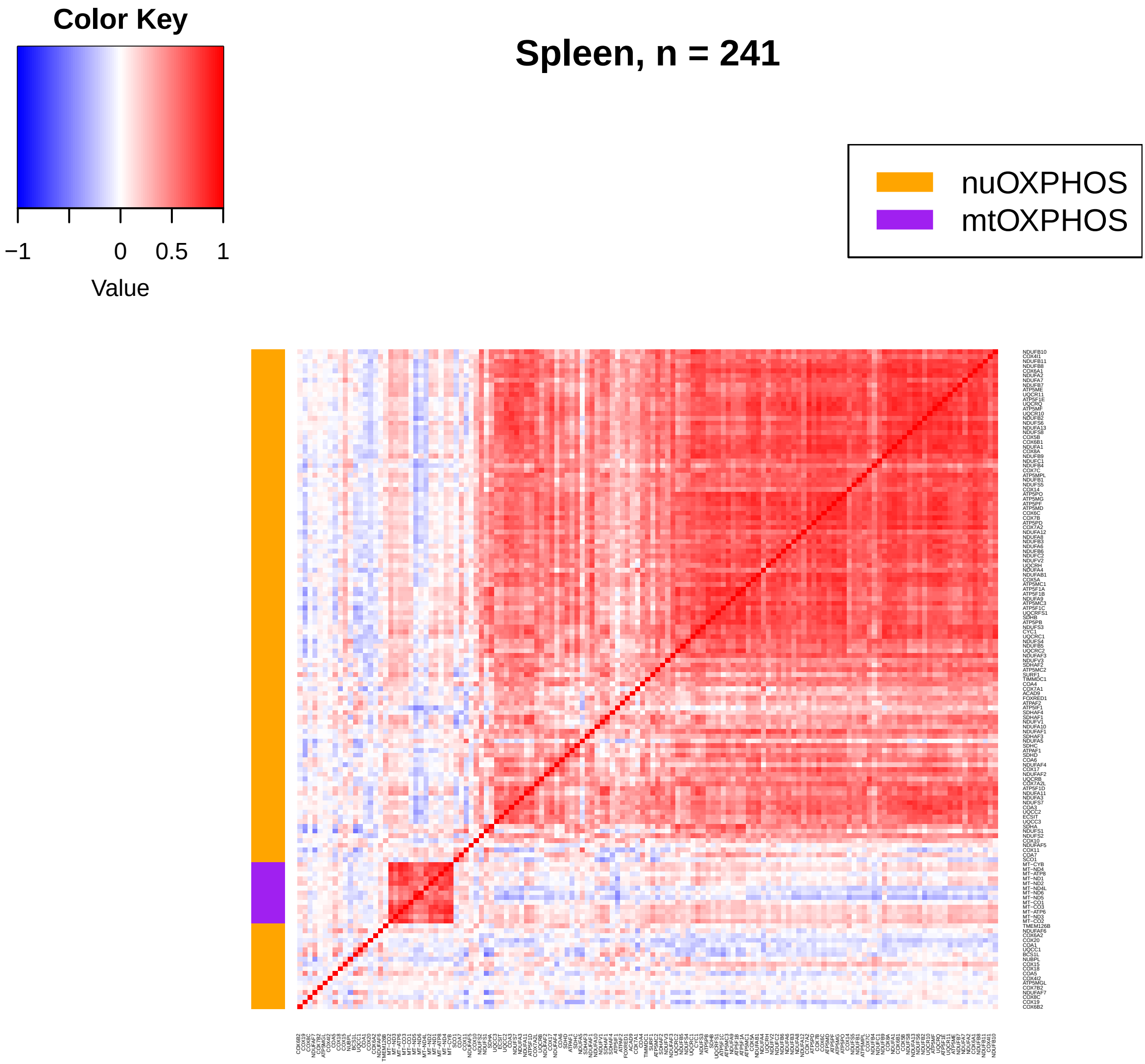
Color Key
−1
0
0.5
1
Value
Spleen, n = 241
nuOXPHOS
mtOXPHOS

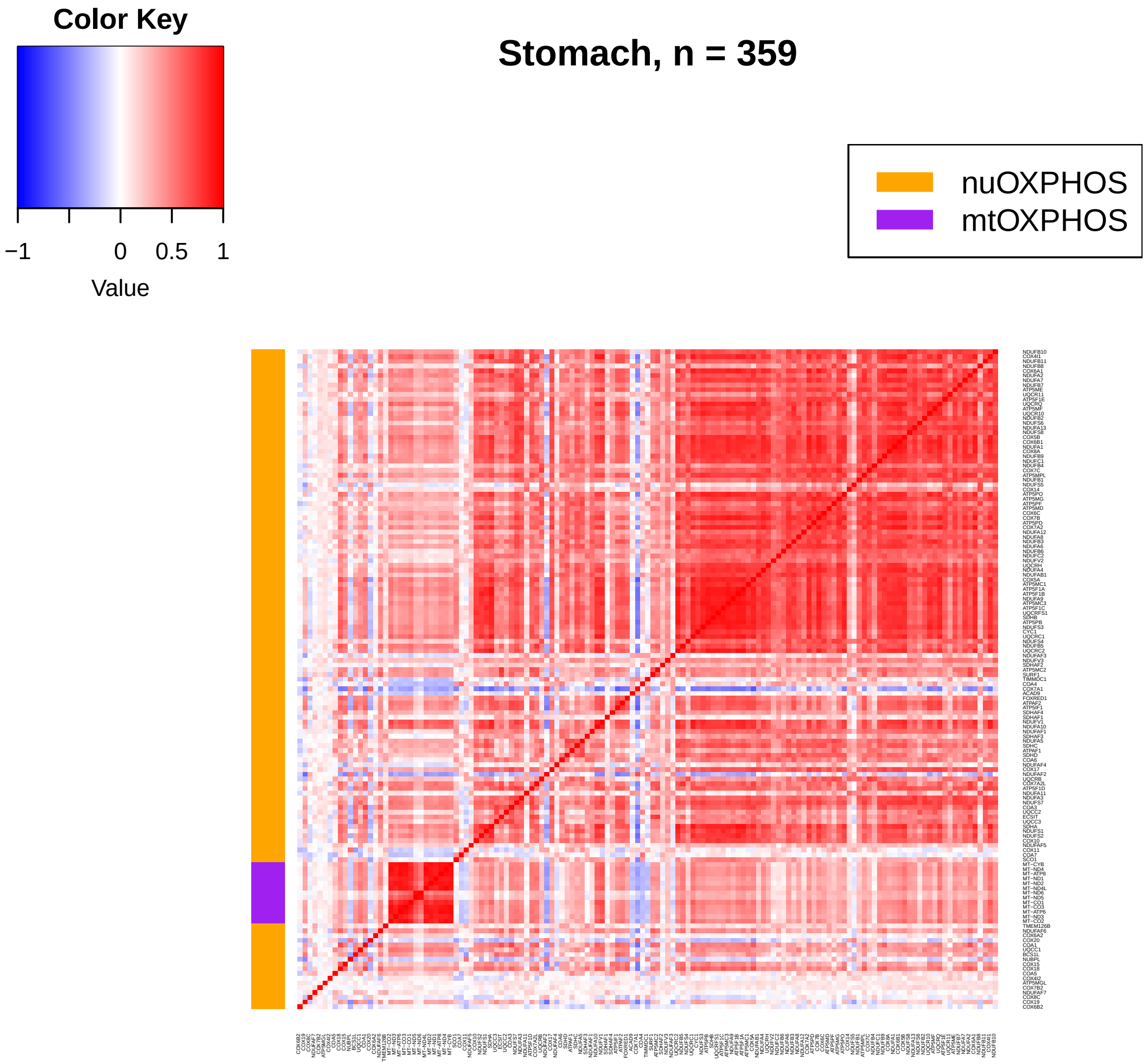
Color Key
−1
0
0.5
1
Value
Stomach, n = 359
nuOXPHOS
mtOXPHOS

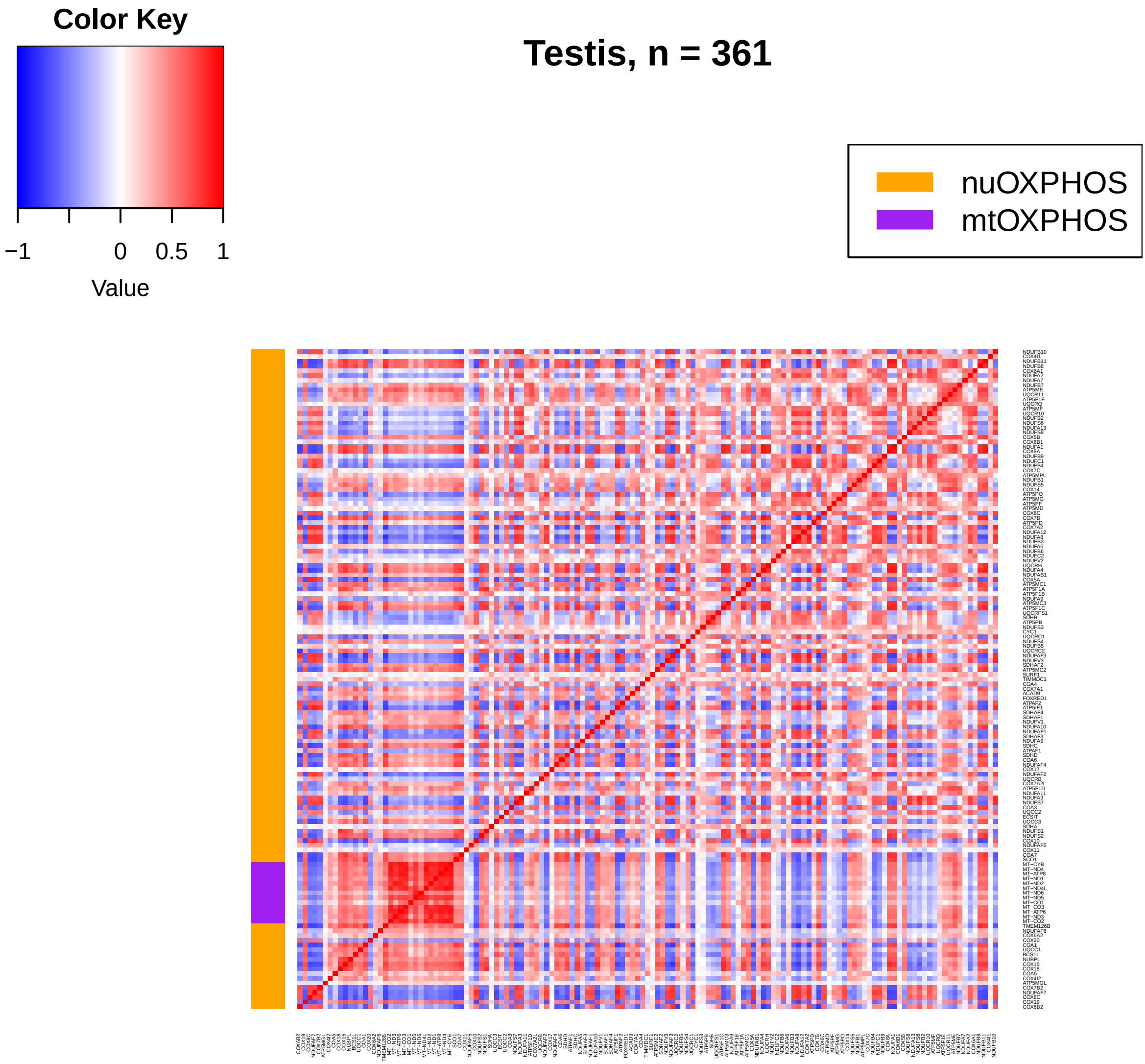
Color Key
−1
0
0.5
1
Value
Testis, n = 361
nuOXPHOS
mtOXPHOS

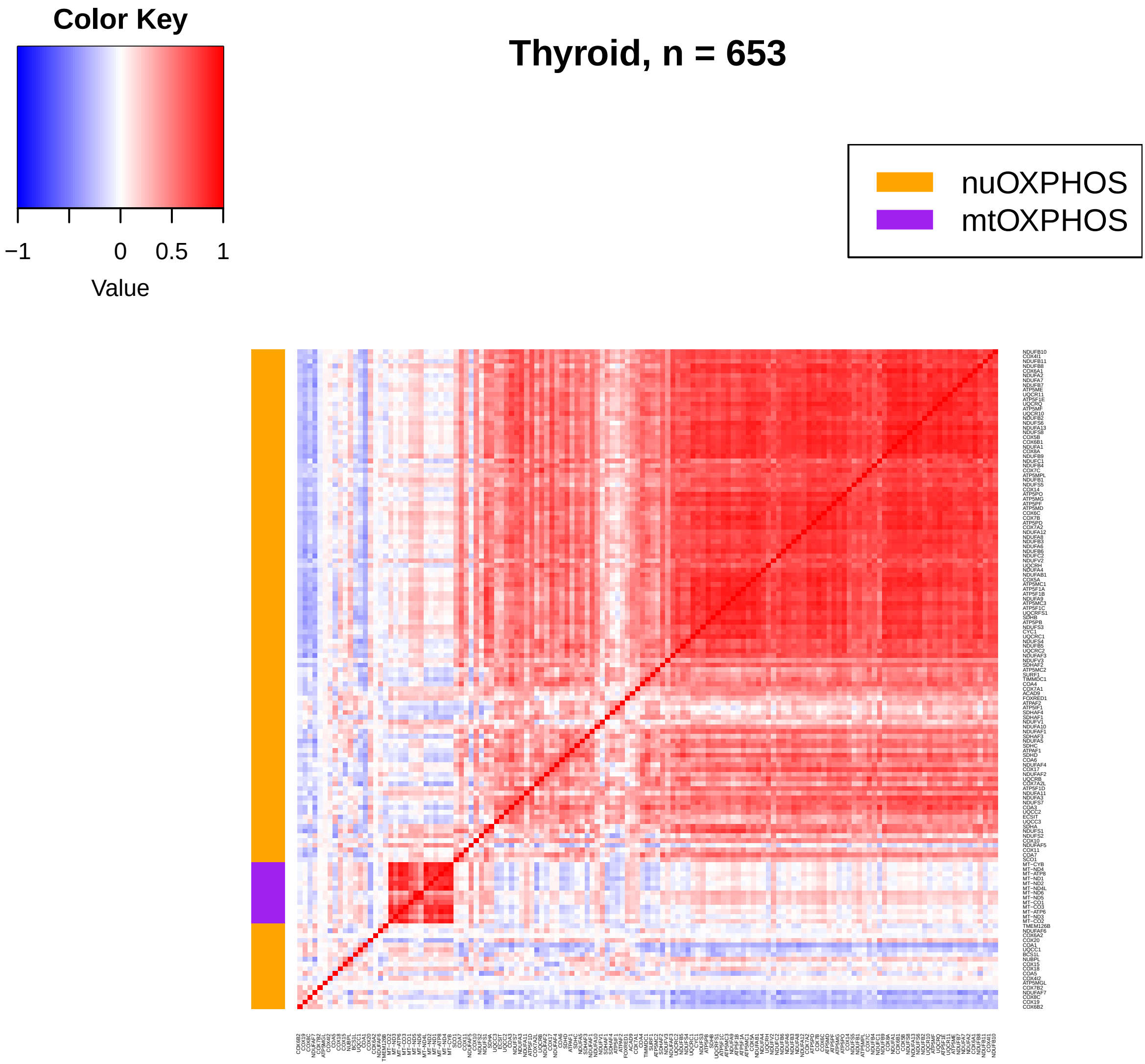
Thyroid, n = 653
Color Key
−1
0
0.5
1
Value
nuOXPHOS
mtOXPHOS

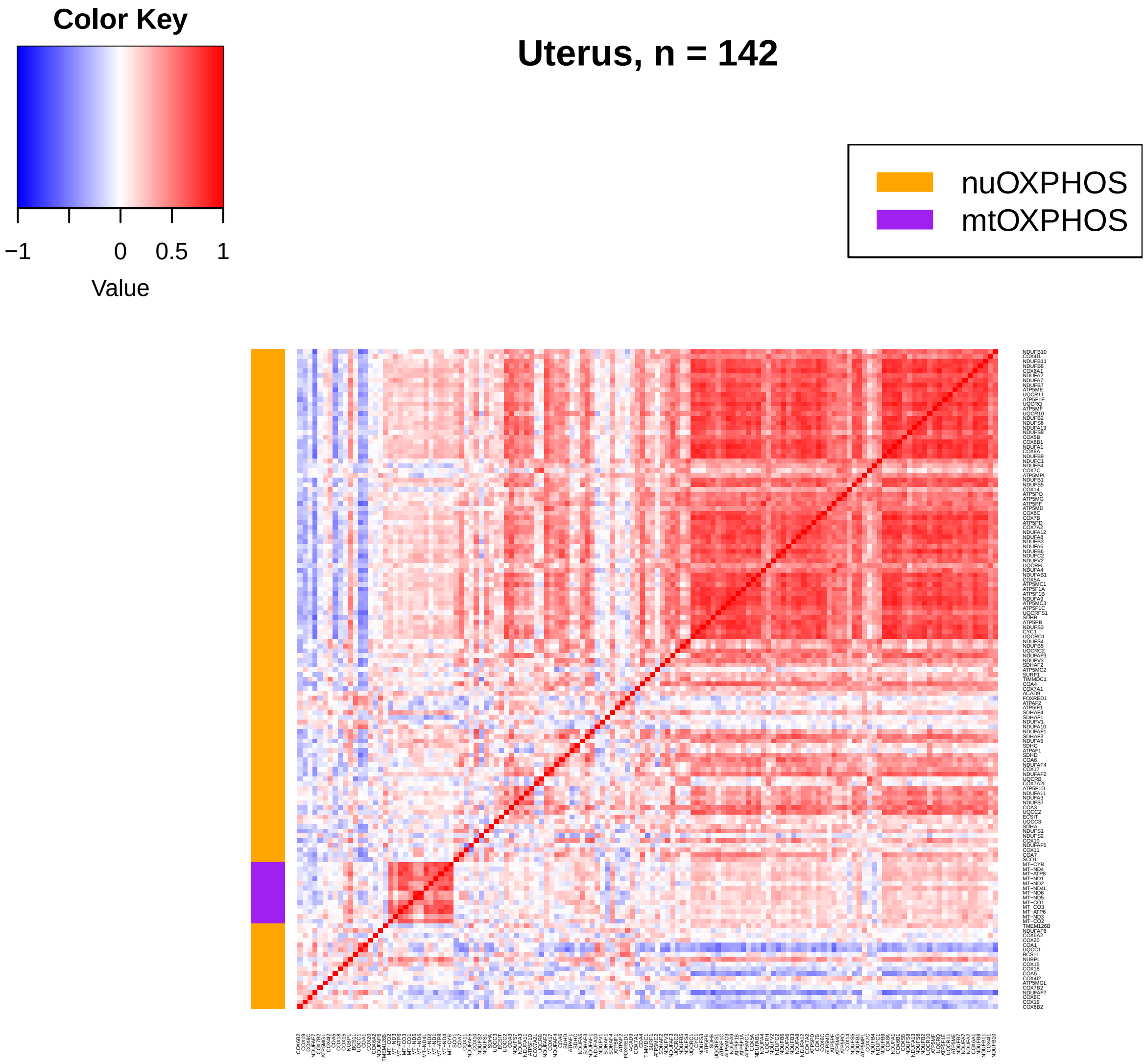
Color Key
−1
0
0.5
1
Value
Uterus, n = 142
nuOXPHOS
mtOXPHOS

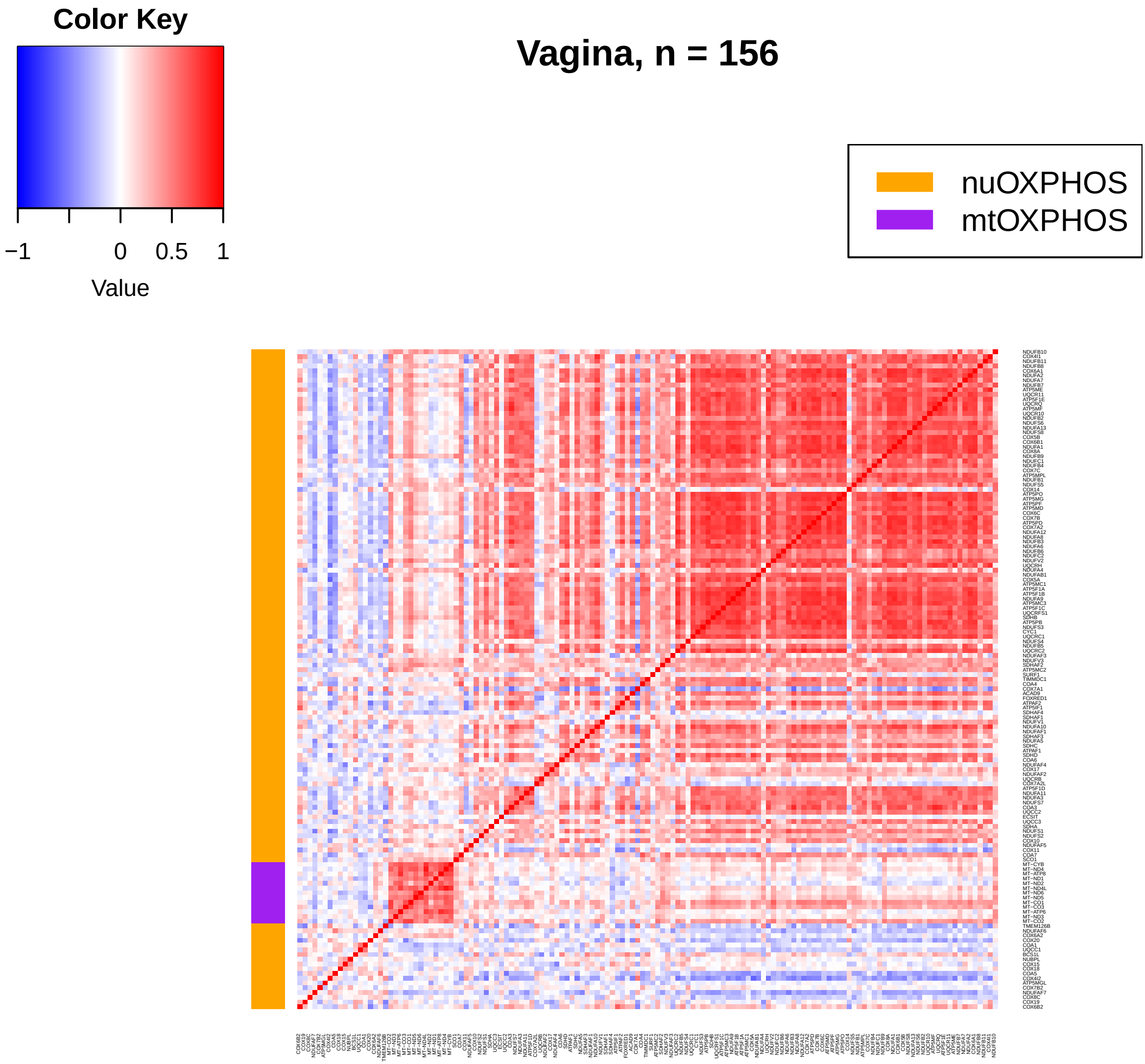
Color Key
−1
0
0.5
1
Value
Vagina, n = 156
nuOXPHOS
mtOXPHOS

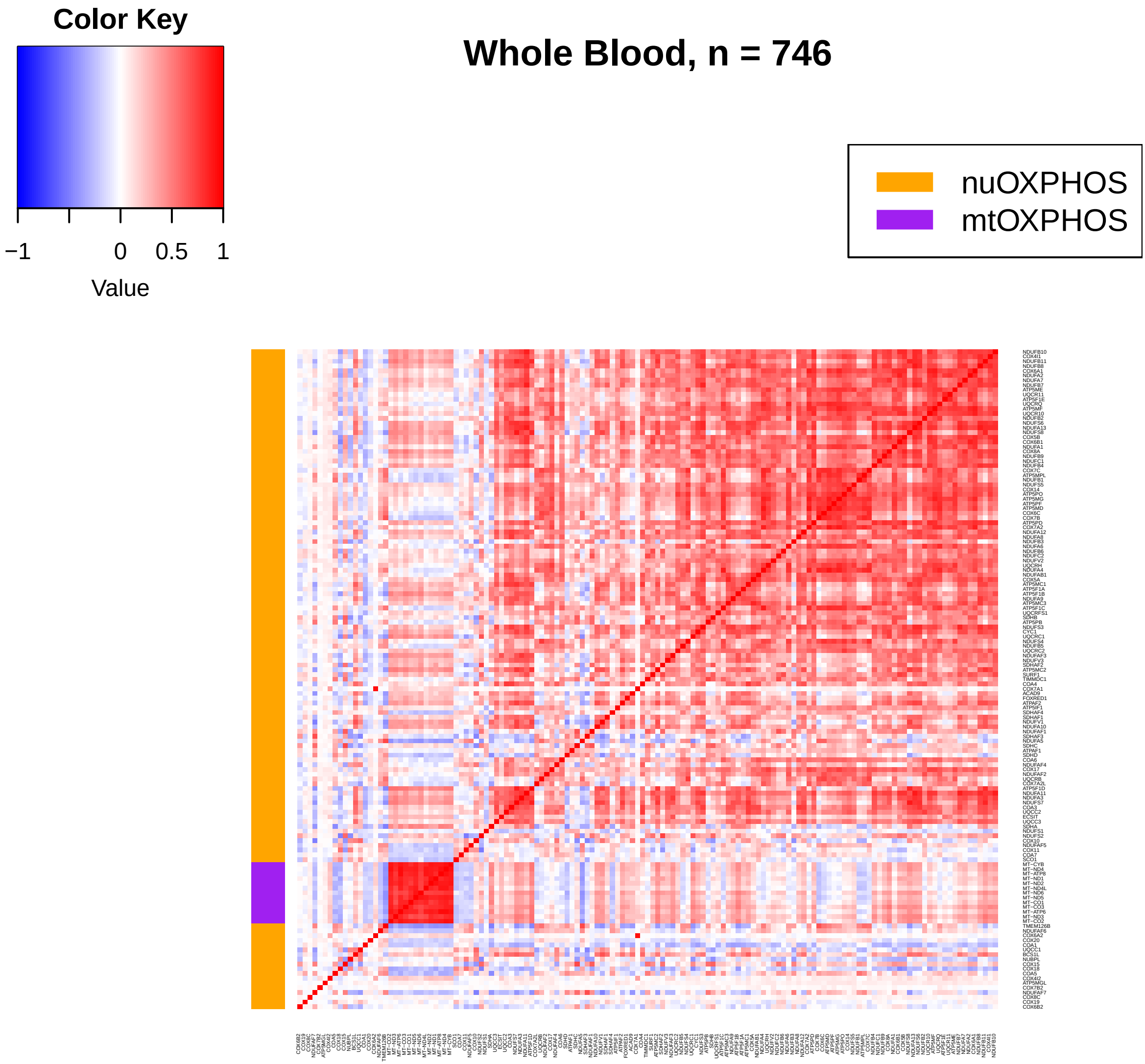
Color Key
−1
0
0.5
1
Value
Whole Blood, n = 746
nuOXPHOS
mtOXPHOS
