## Supplemental tables and scripts for "Malignancy and NF-kB signalling strengthen coordination between the expression of mitochondrial and nuclear-encoded oxidative phosphorylation genes": Supplementary-Dataset-2_MRN-mtOXPHOS-nuOXPHOS-cancer-heatmaps.pdf

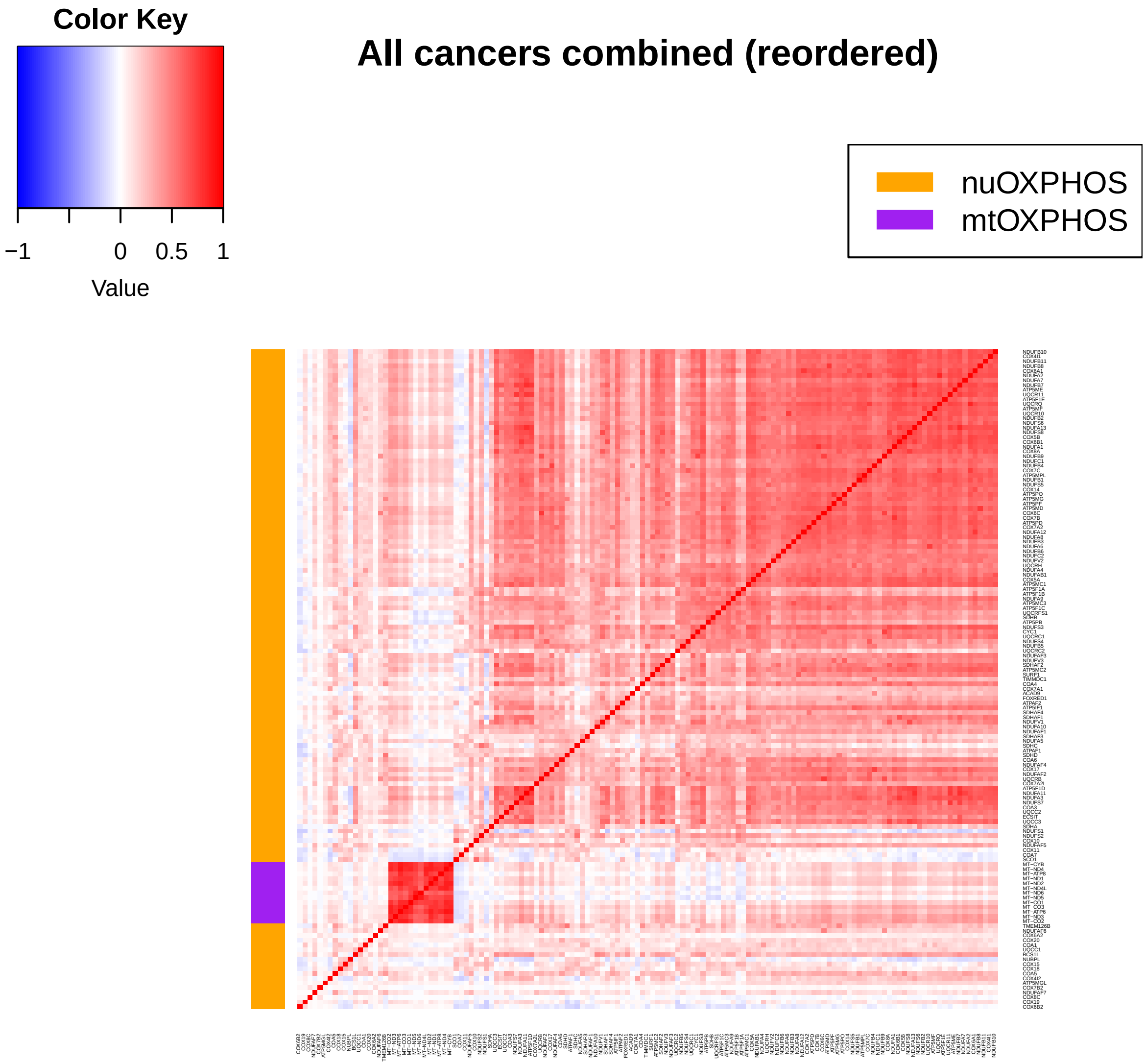
Color Key
−1
0
0.5
1
Value
All cancers combined (reordered)
nuOXPHOS
mtOXPHOS

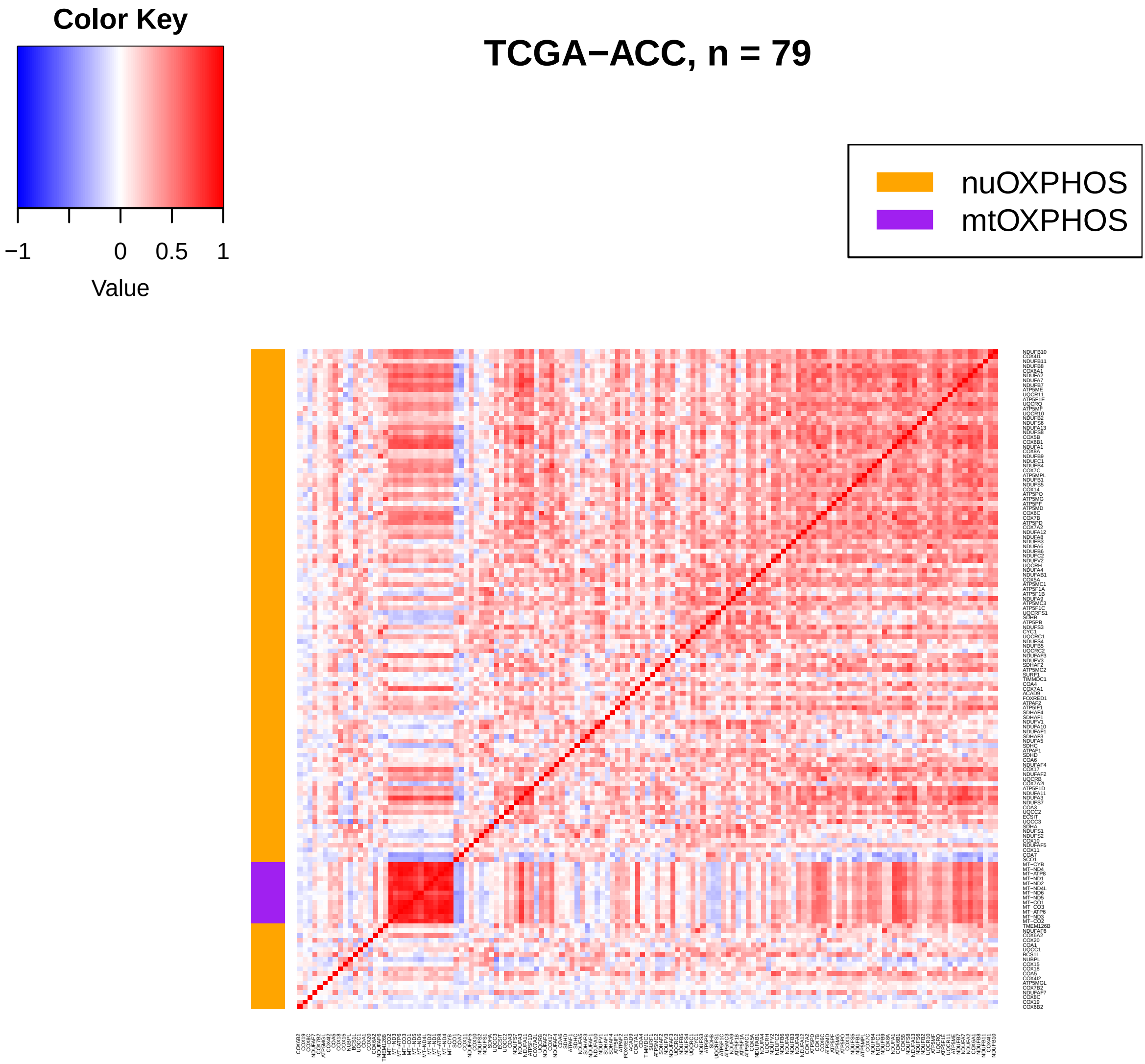
Color Key
−1
0
0.5
1
Value
TCGA−ACC, n = 79
nuOXPHOS
mtOXPHOS

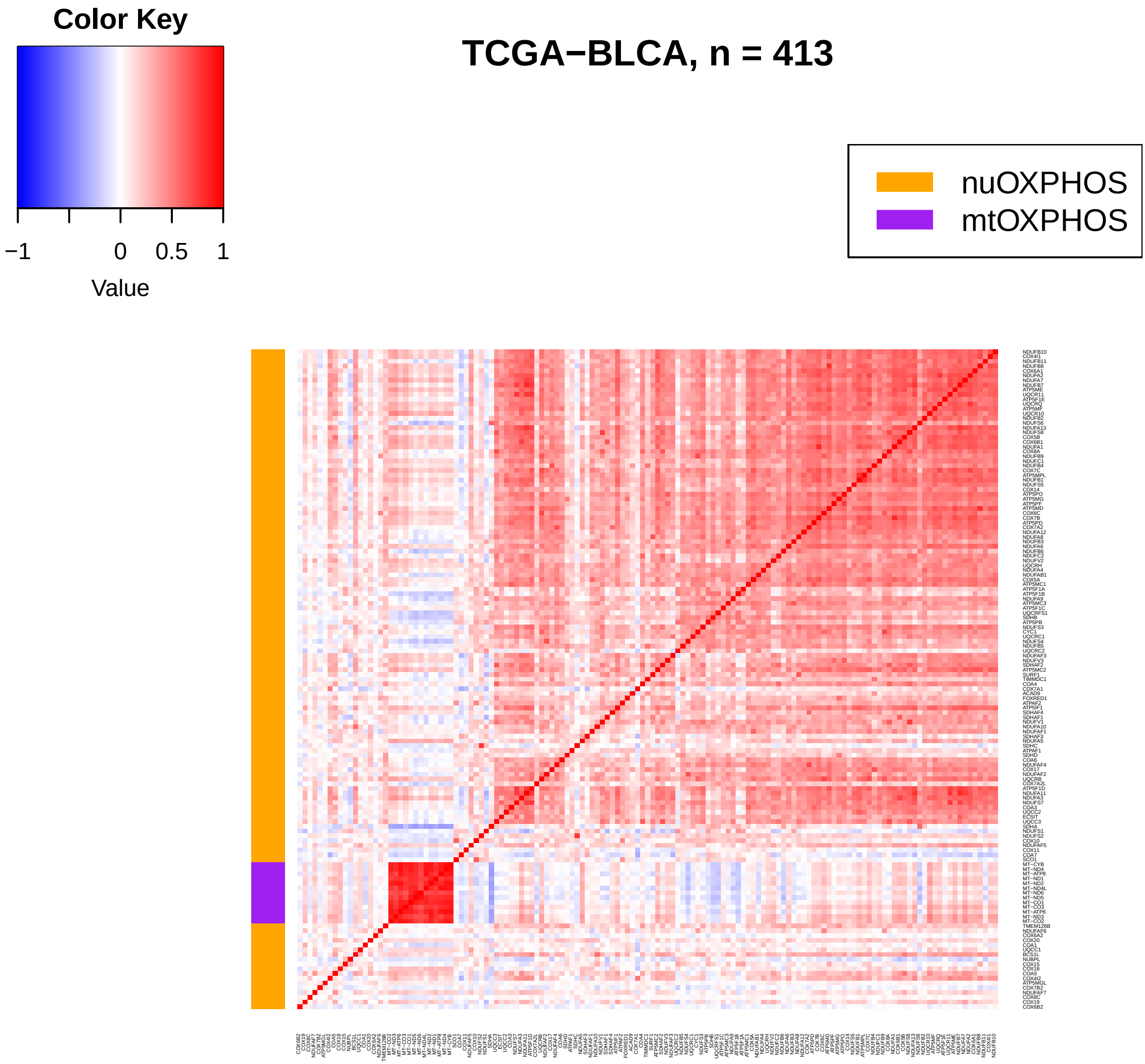

Color Key
−1
0
0.5
1
Value
TCGA−BLCA, n = 413
nuOXPHOS
mtOXPHOS

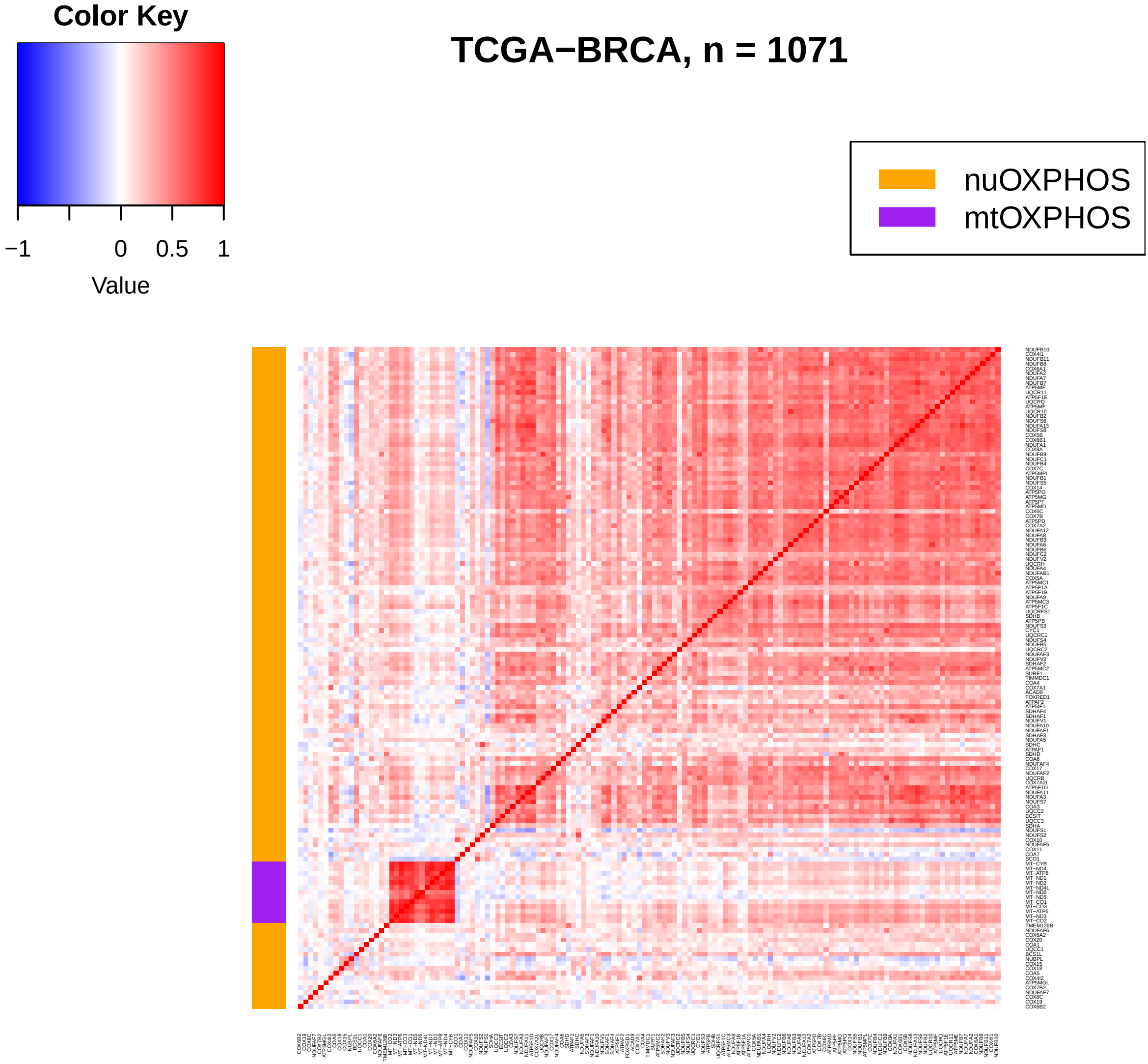
Color Key
−1
0
0.5
1
Value
TCGA−BRCA, n = 1071
nuOXPHOS
mtOXPHOS

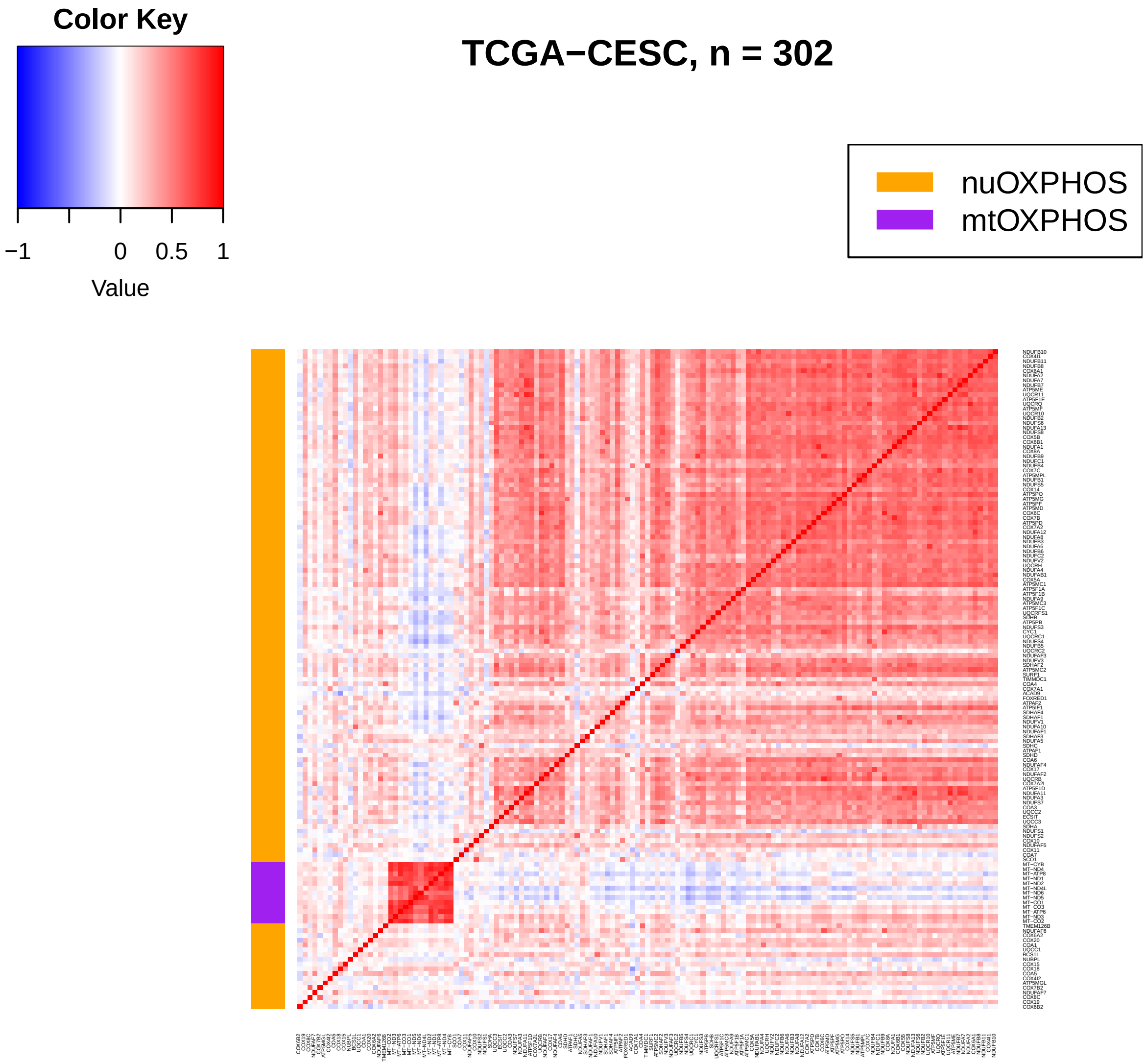

Color Key
−1
0
0.5
1
Value
TCGA−CESC, n = 302
nuOXPHOS
mtOXPHOS

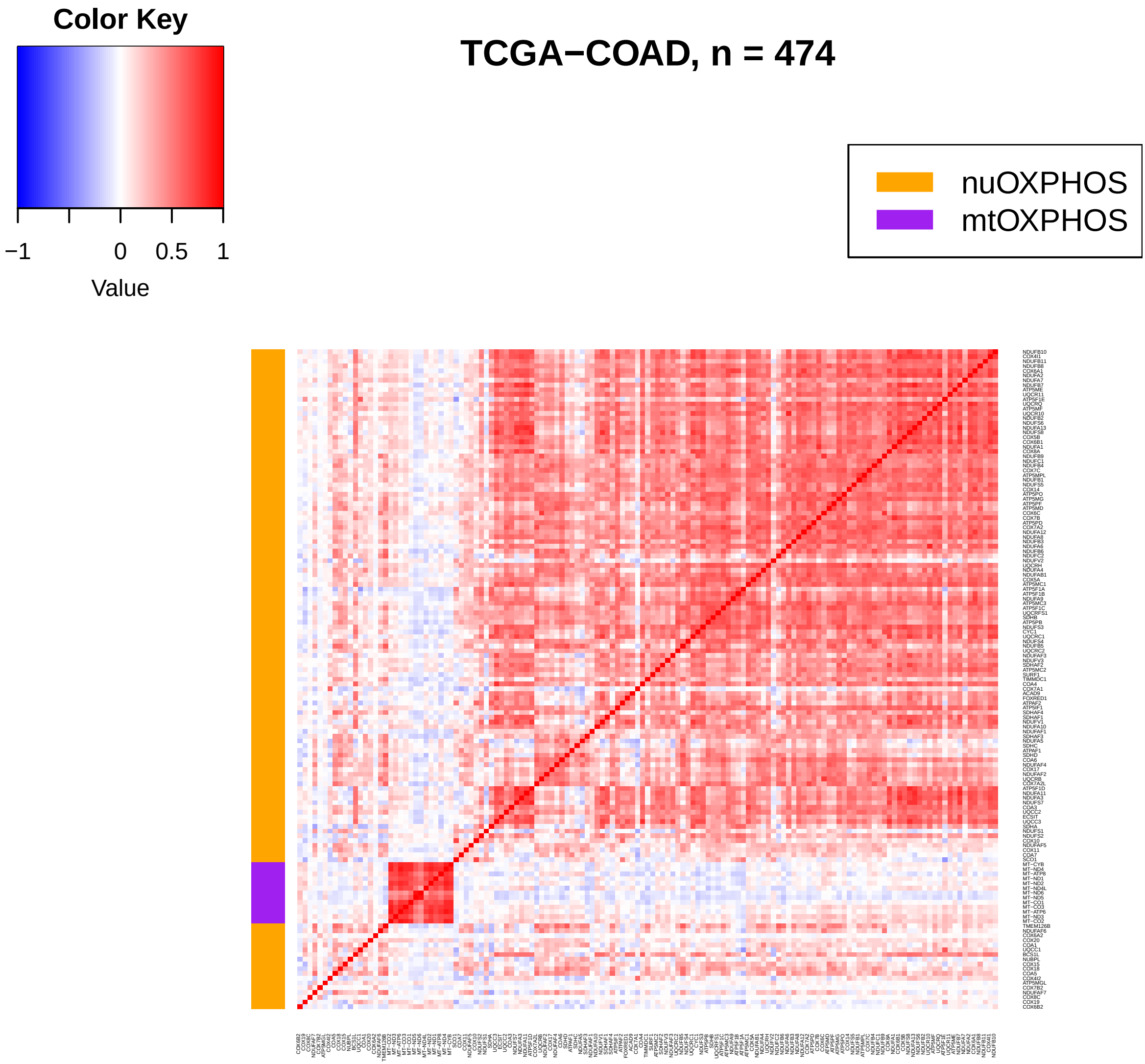
Color Key
−1
0
0.5
1
Value
TCGA−COAD, n = 474
nuOXPHOS
mtOXPHOS

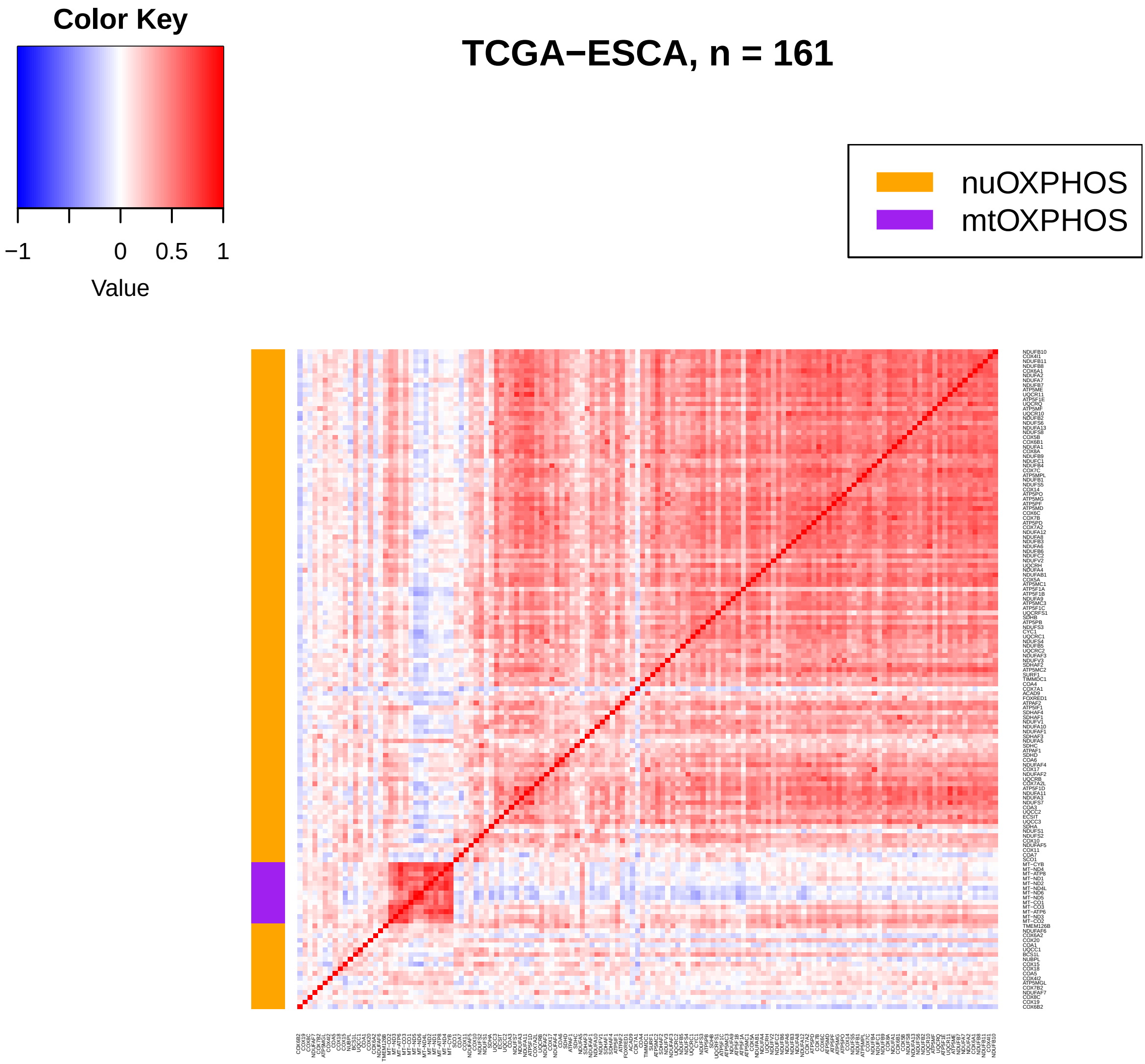
Color Key
−1
0
0.5
1
Value
TCGA−ESCA, n = 161
nuOXPHOS
mtOXPHOS

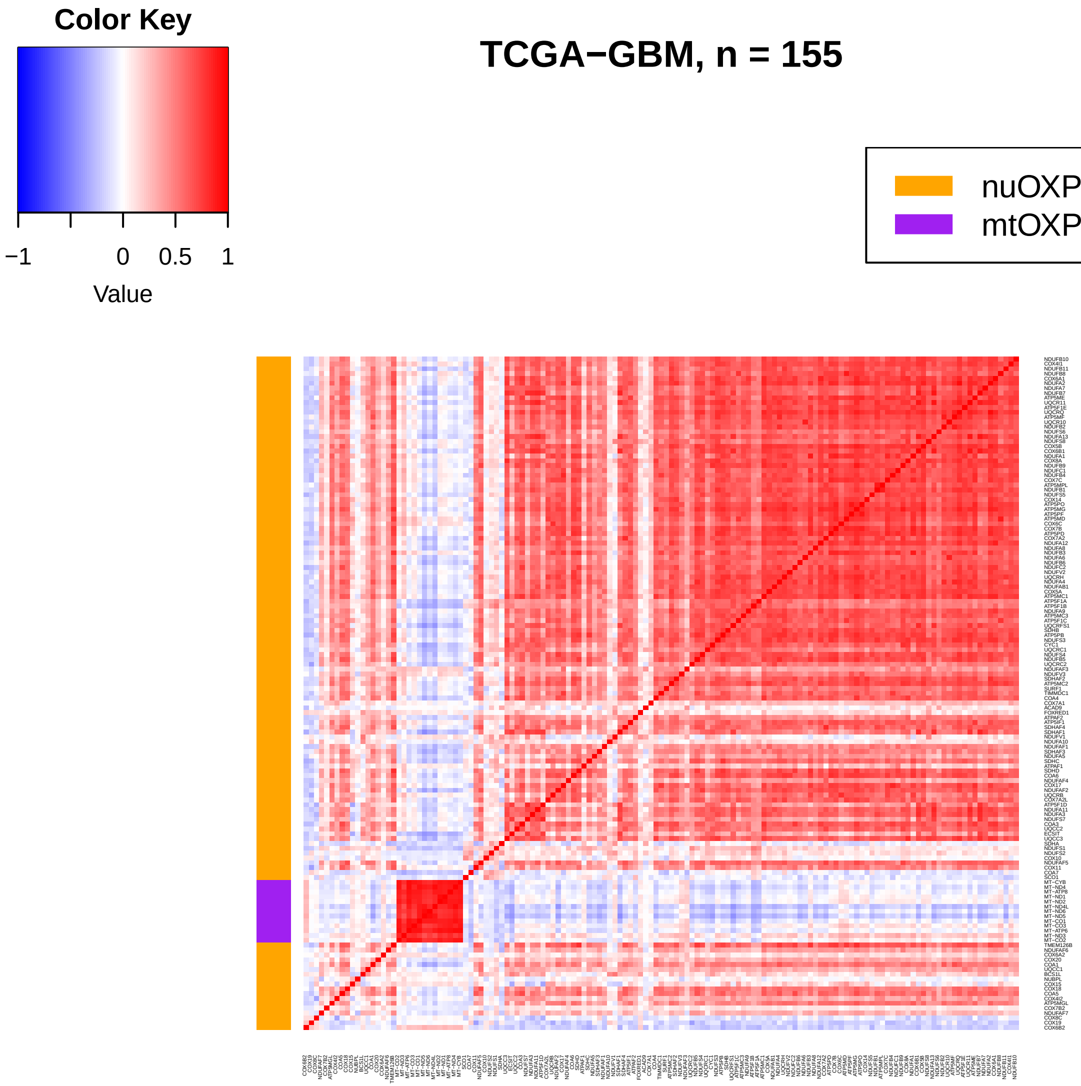
Color Key
−1
0
0.5
1
Value
TCGA−GBM, n = 155
nuOXPHOS
mtOXPHOS

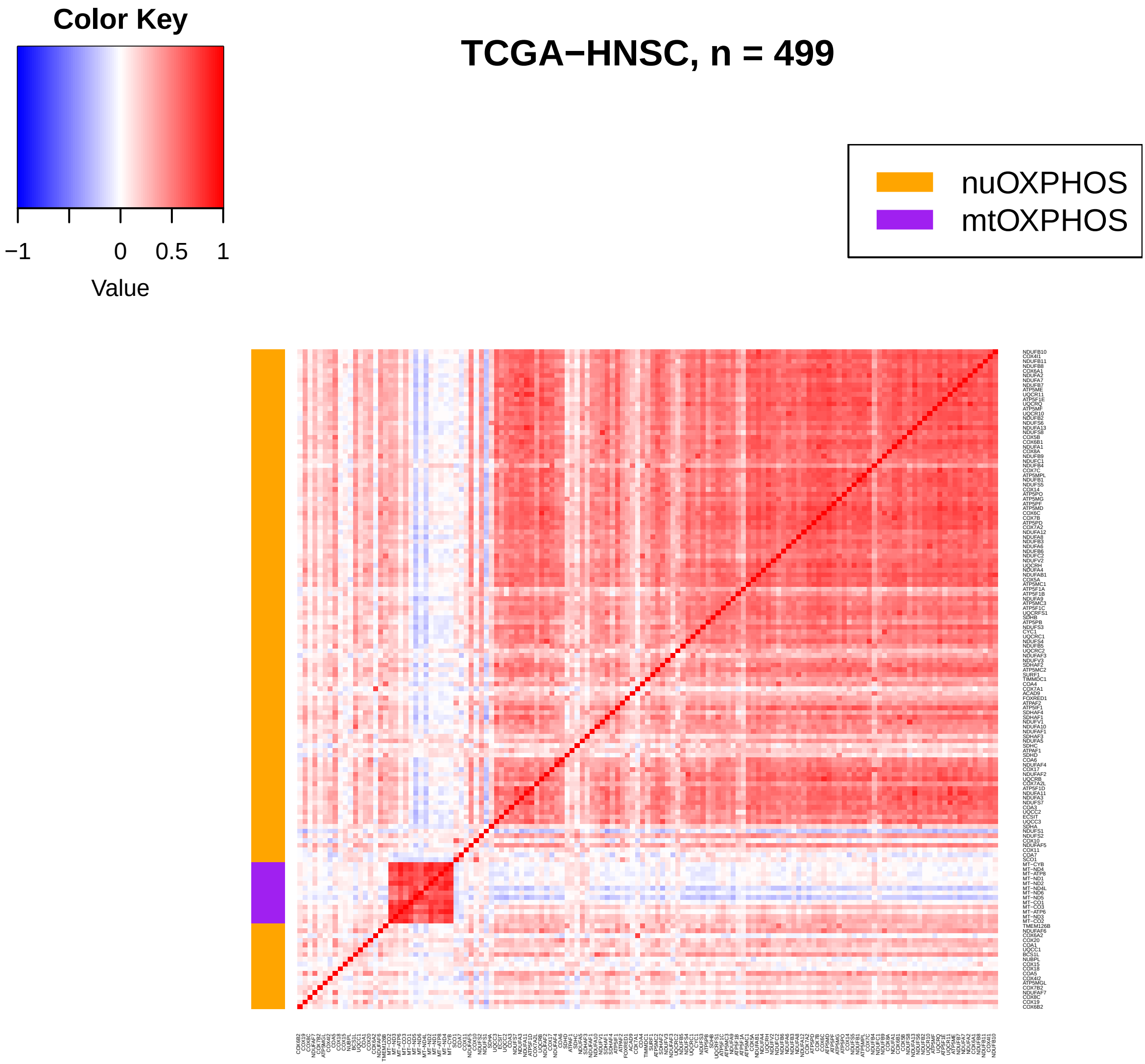
Color Key
−1
0
0.5
1
Value
TCGA−HNSC, n = 499
nuOXPHOS
mtOXPHOS

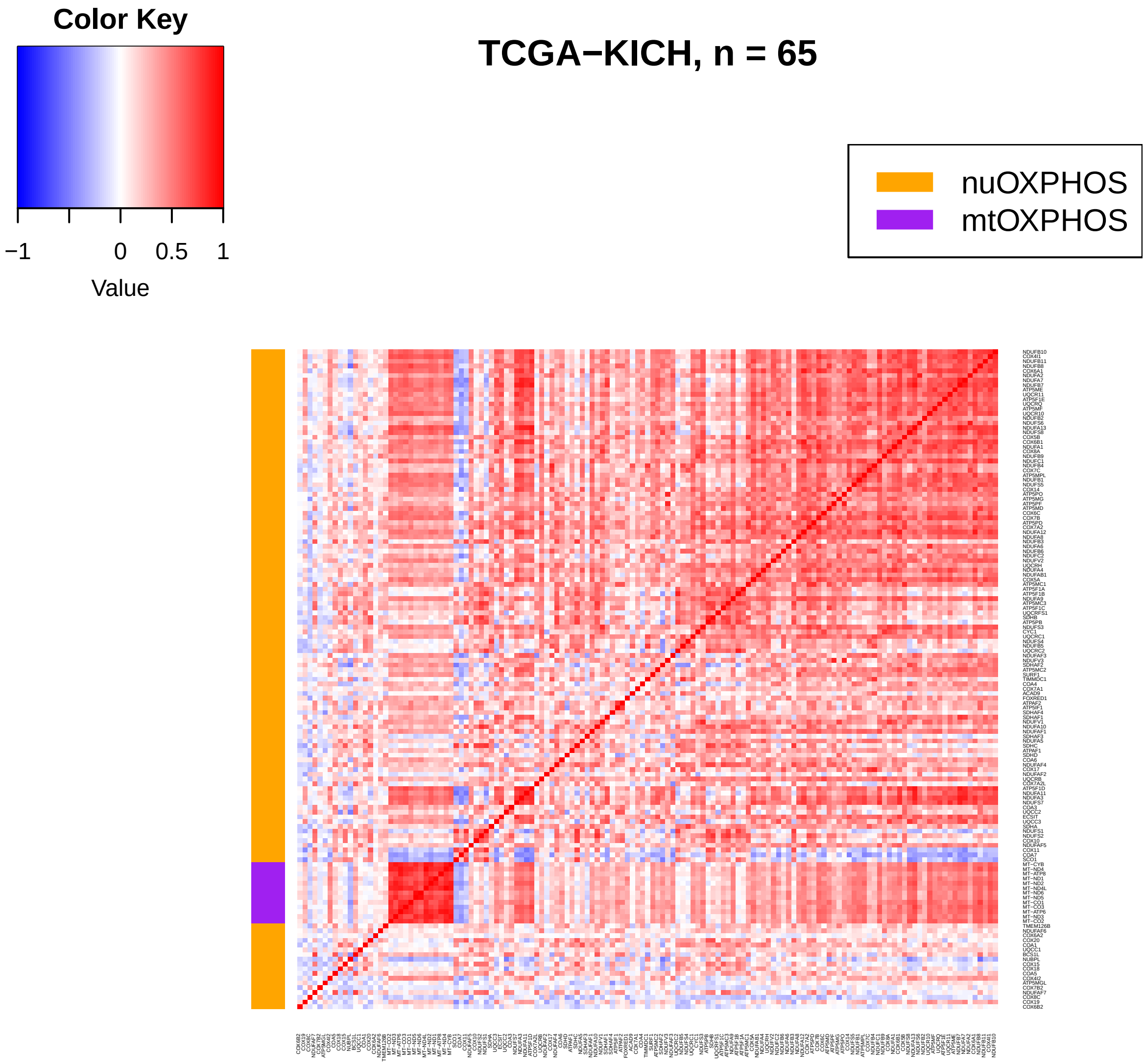

Color Key
−1
0
0.5
1
Value
TCGA−KICH, n = 65
nuOXPHOS
mtOXPHOS

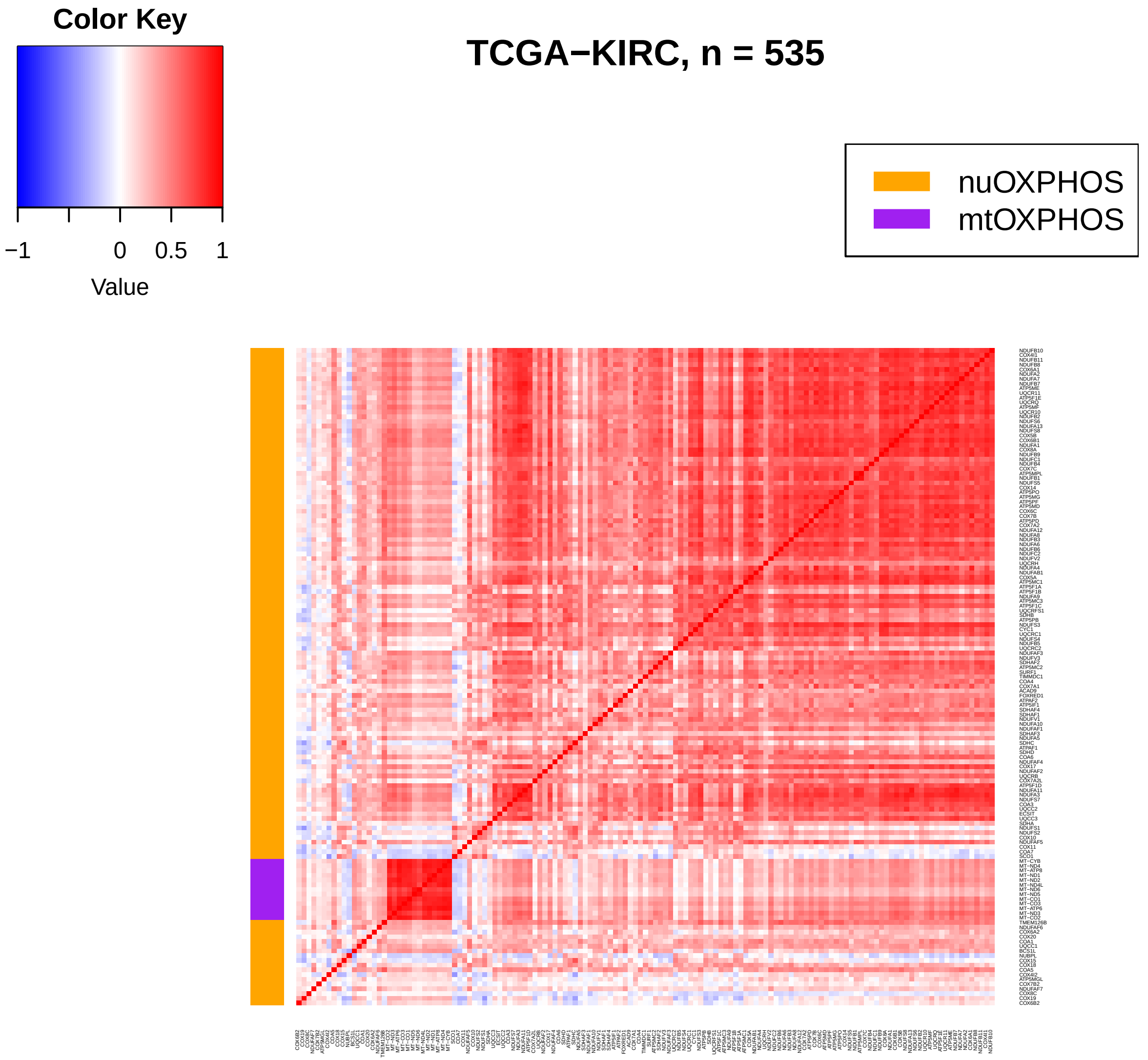
Color Key
−1
0
0.5
1
Value
TCGA−KIRC, n = 535
nuOXPHOS
mtOXPHOS

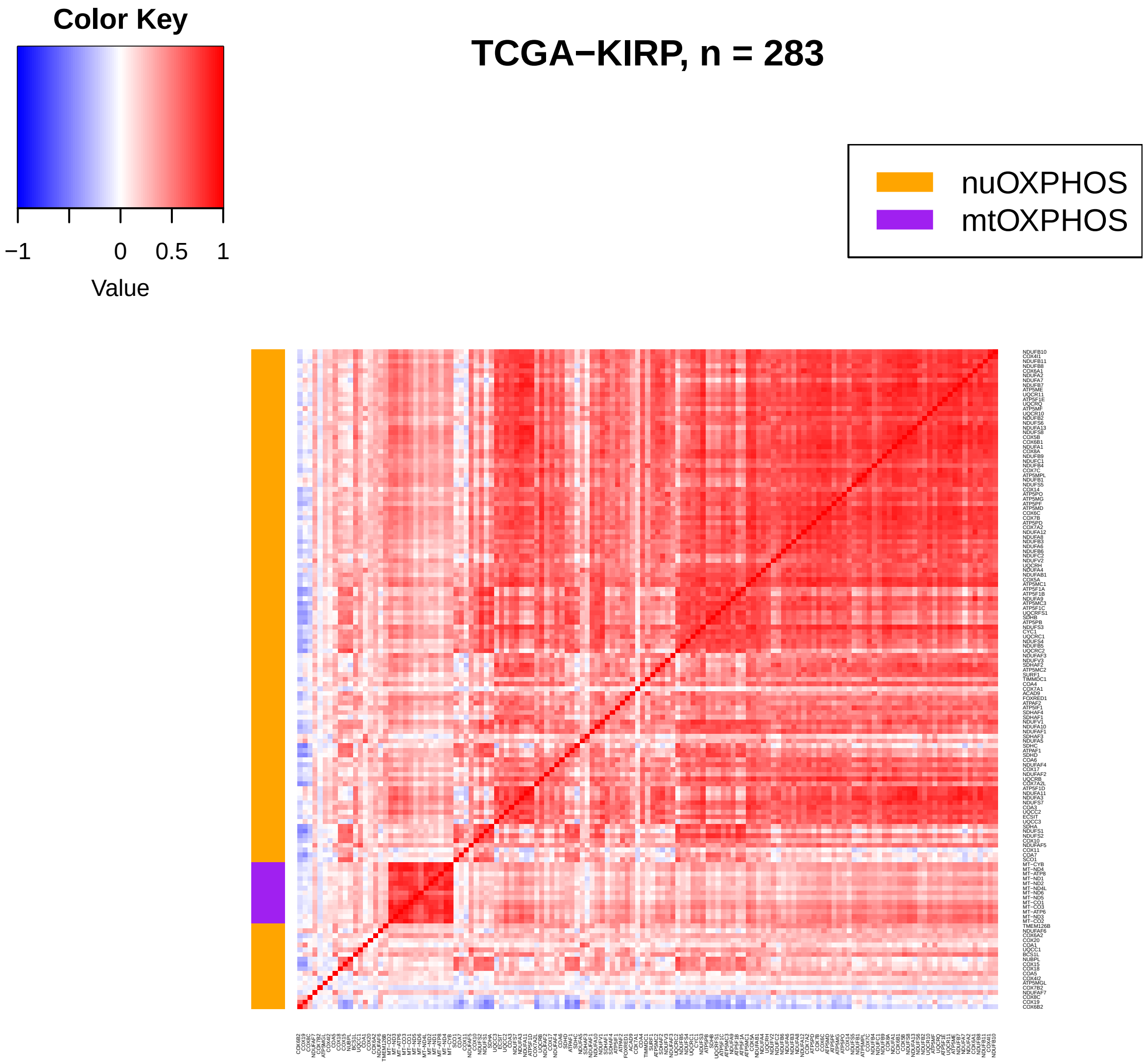

Color Key
−1
0
0.5
1
Value
TCGA−KIRP, n = 283
nuOXPHOS
mtOXPHOS

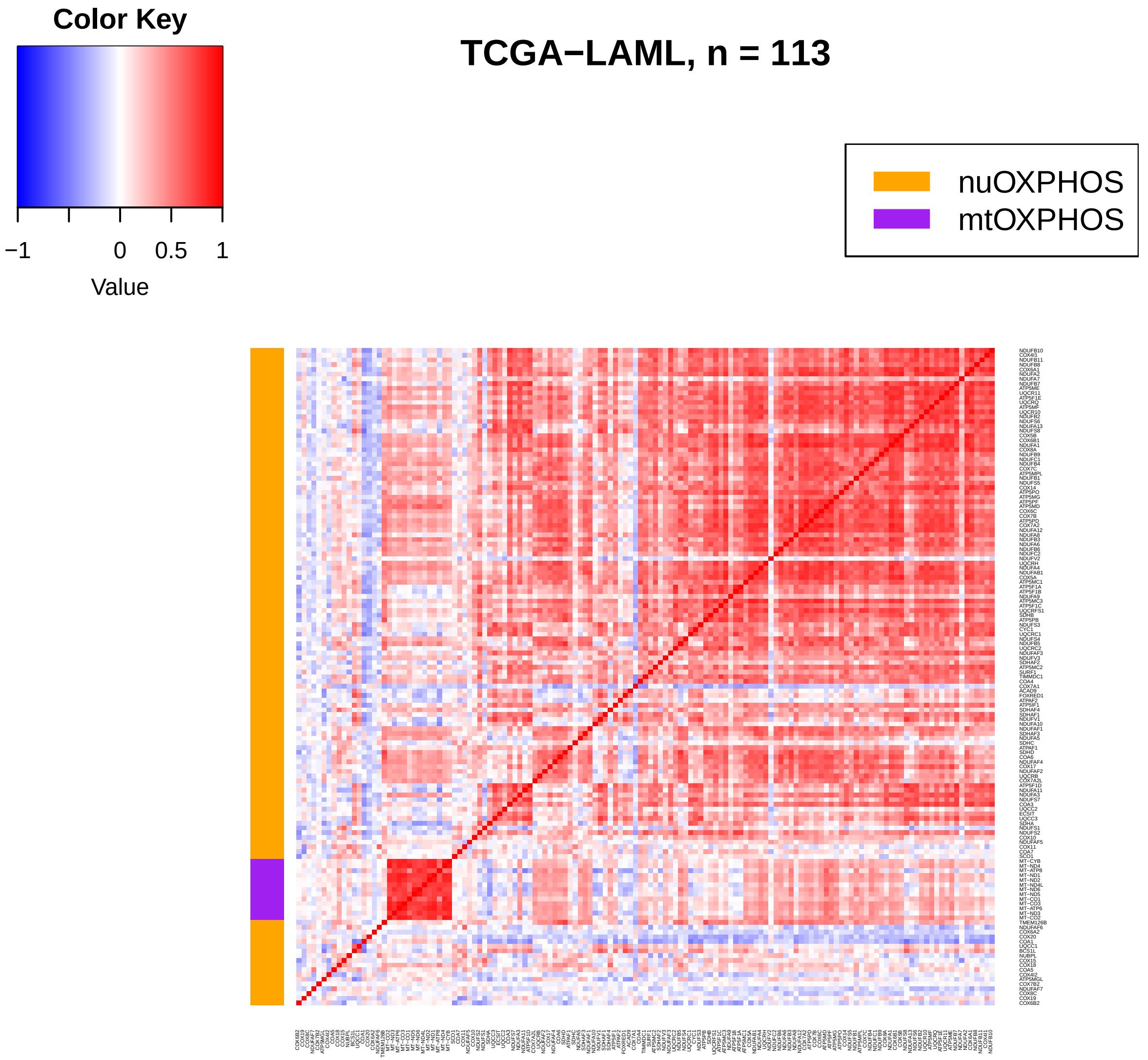
Color Key
−1
0
0.5
1
Value
TCGA−LAML, n = 113
nuOXPHOS
mtOXPHOS

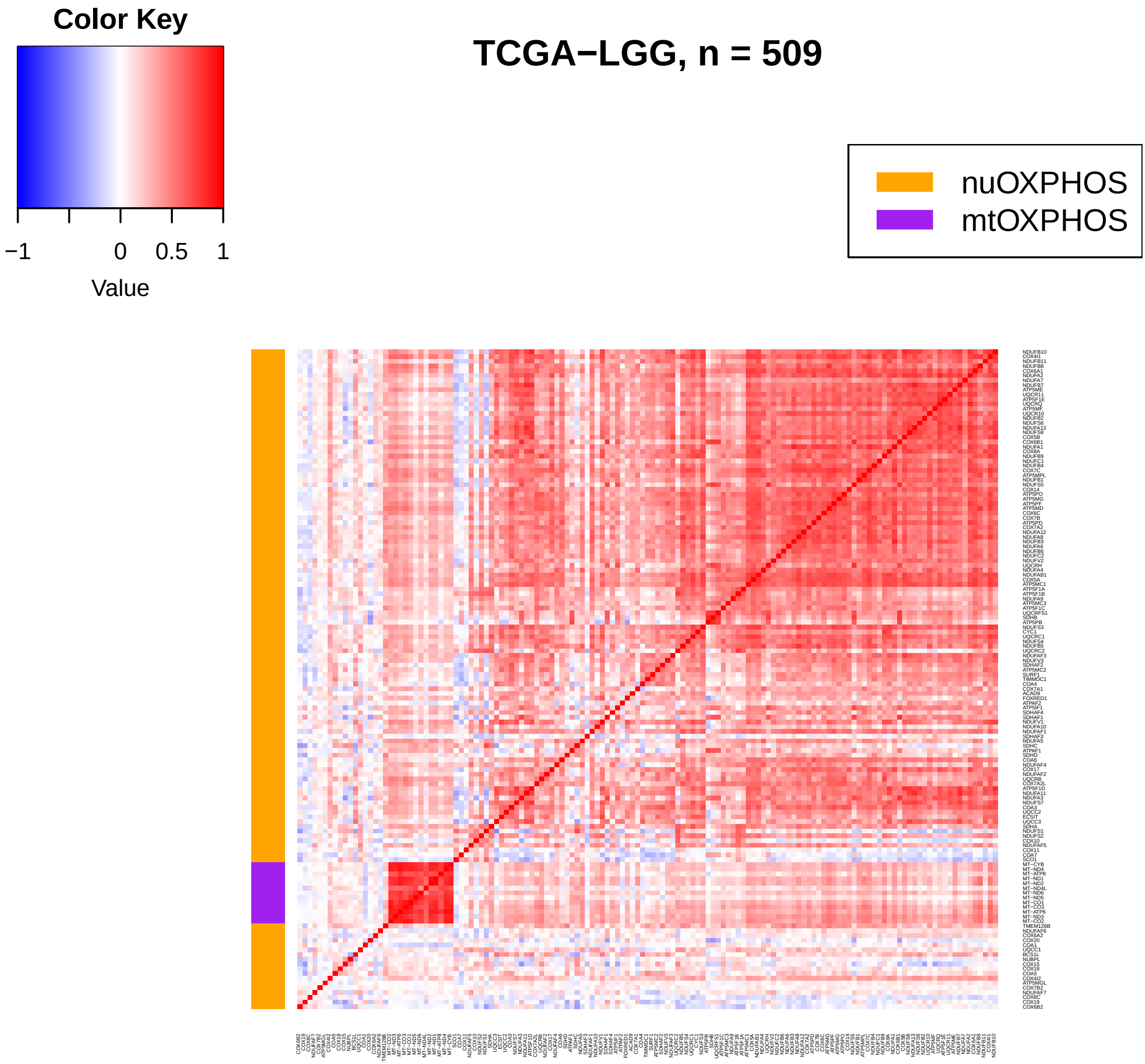
Color Key
−1
0
0.5
1
Value
TCGA−LGG, n = 509
nuOXPHOS
mtOXPHOS

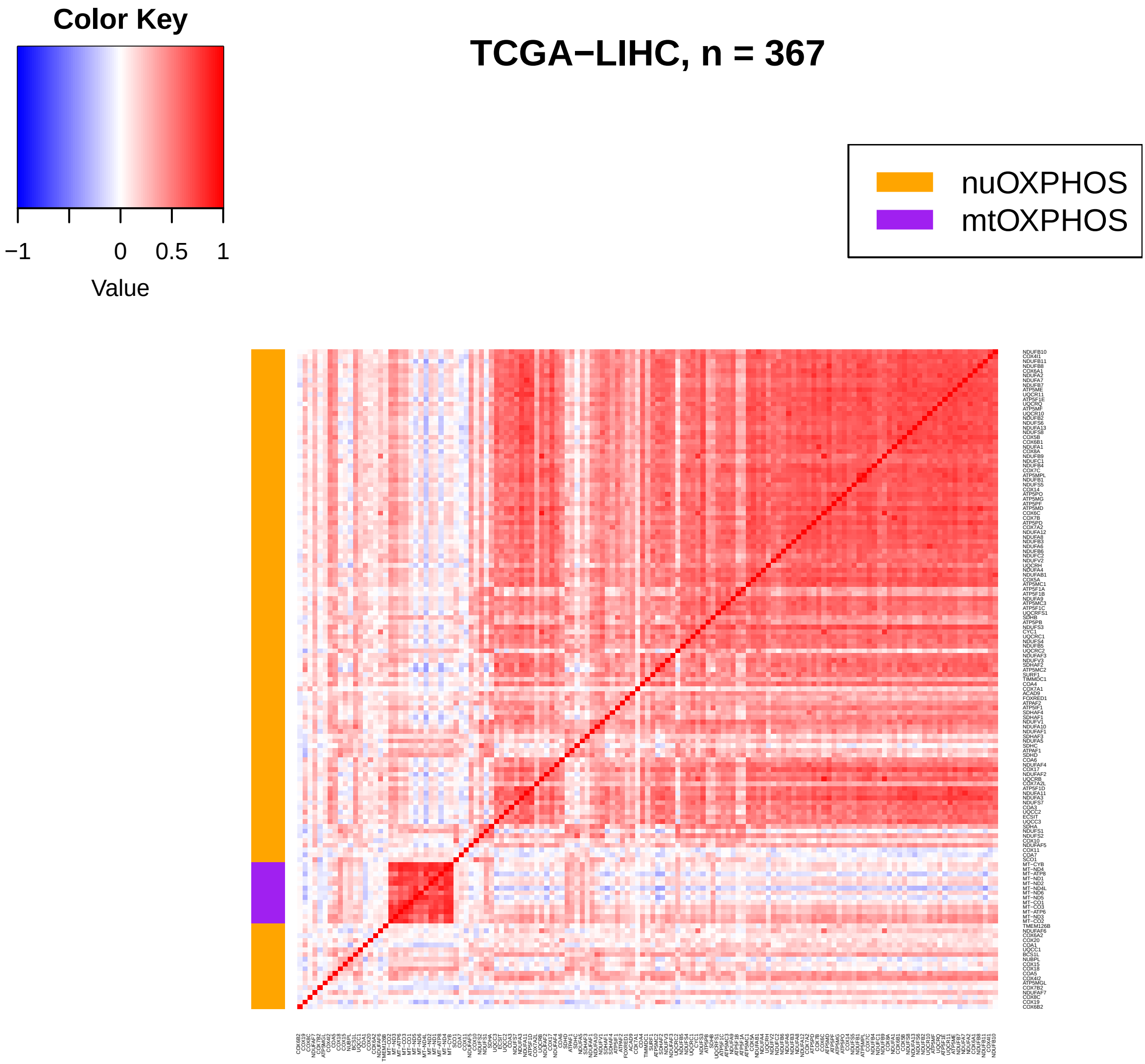
Color Key
−1
0
0.5
1
Value
TCGA−LIHC, n = 367
nuOXPHOS
mtOXPHOS

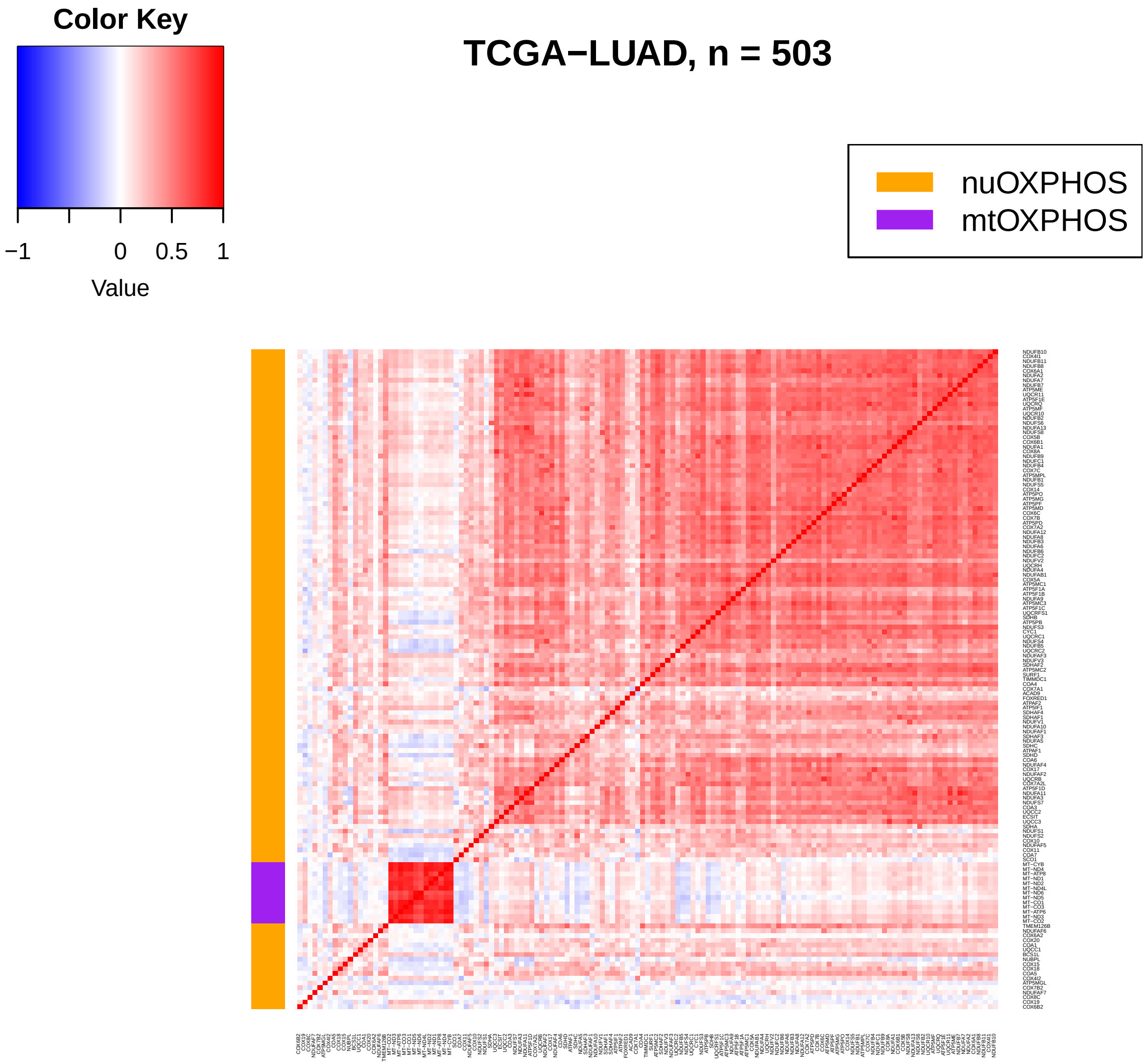

Color Key
−1
0
0.5
1
Value
TCGA−LUAD, n = 503
nuOXPHOS
mtOXPHOS

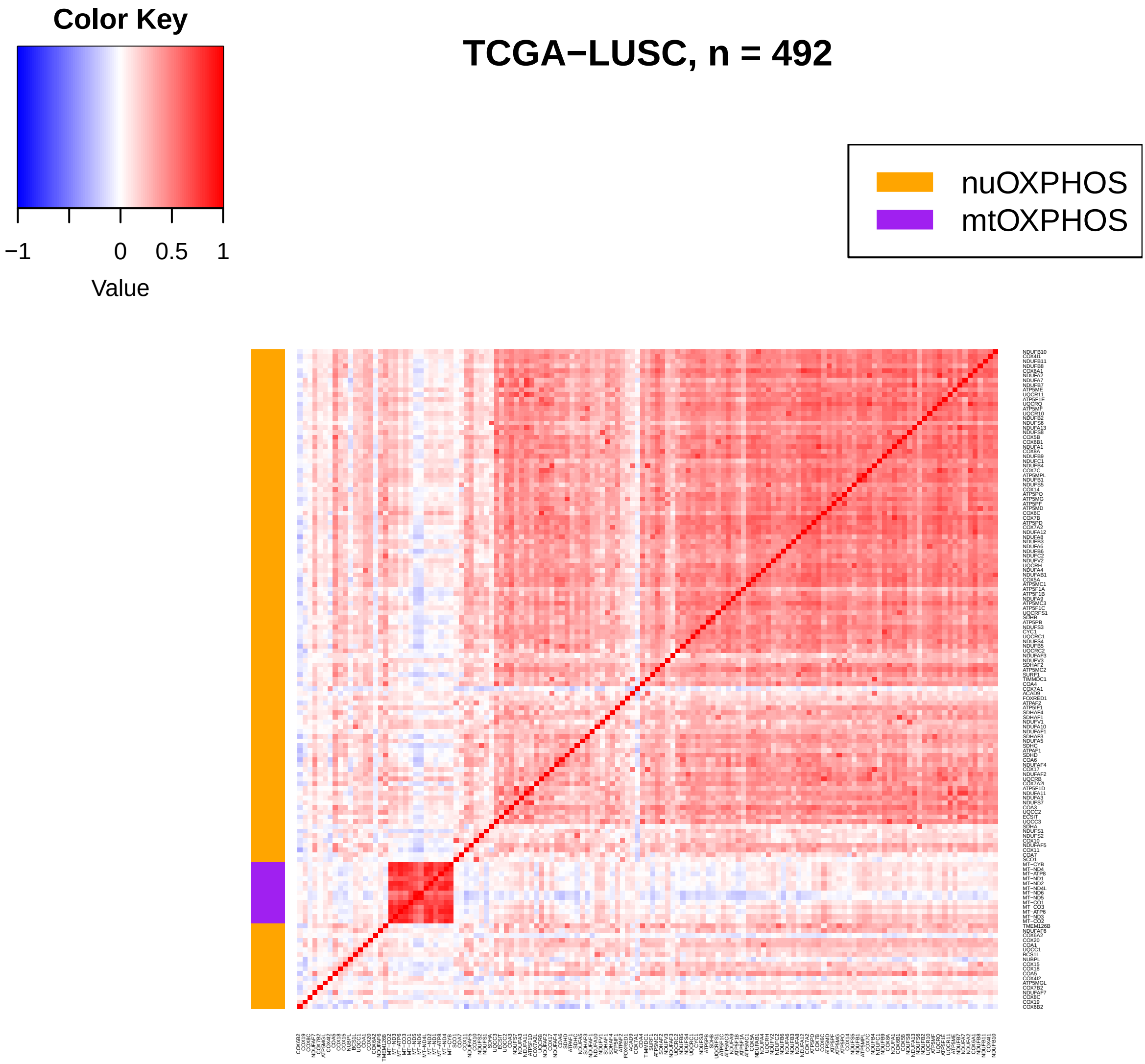
Color Key
−1
0
0.5
1
Value
TCGA−LUSC, n = 492
nuOXPHOS
mtOXPHOS

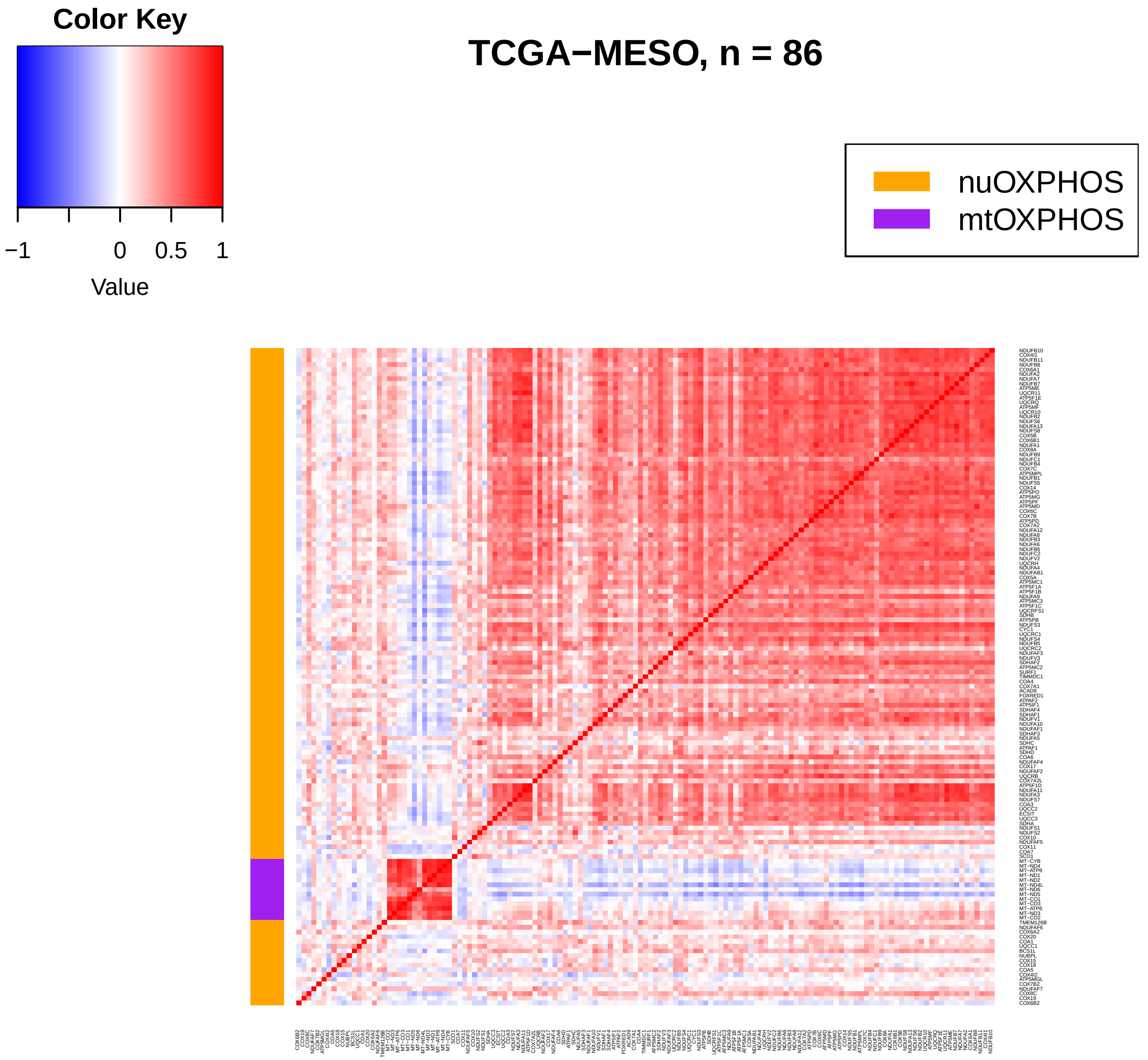

Color Key
−1
0
0.5
1
Value
TCGA−MESO, n = 86
nuOXPHOS
mtOXPHOS

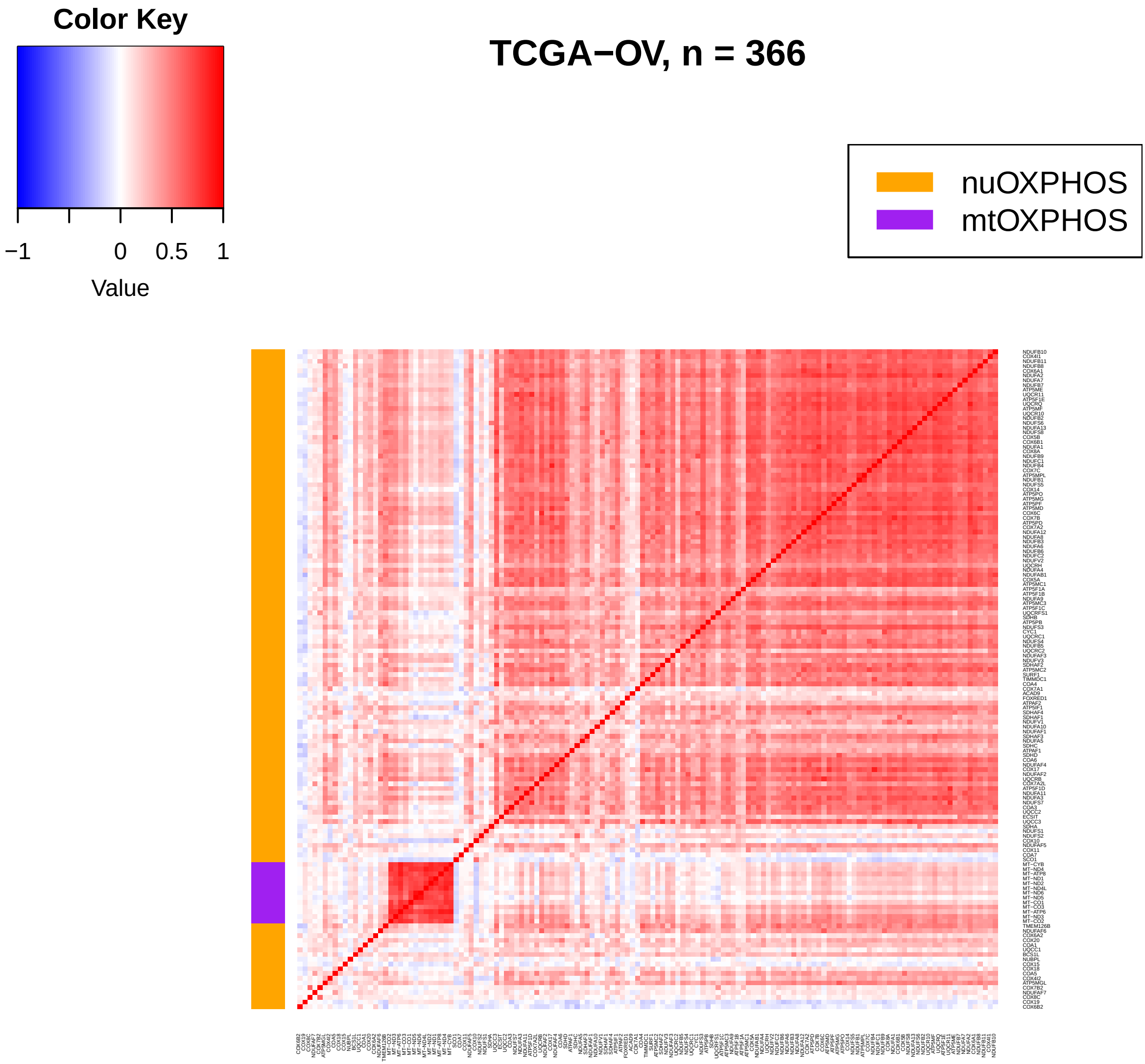

Color Key
−1
0
0.5
1
Value
TCGA−OV, n = 366
nuOXPHOS
mtOXPHOS

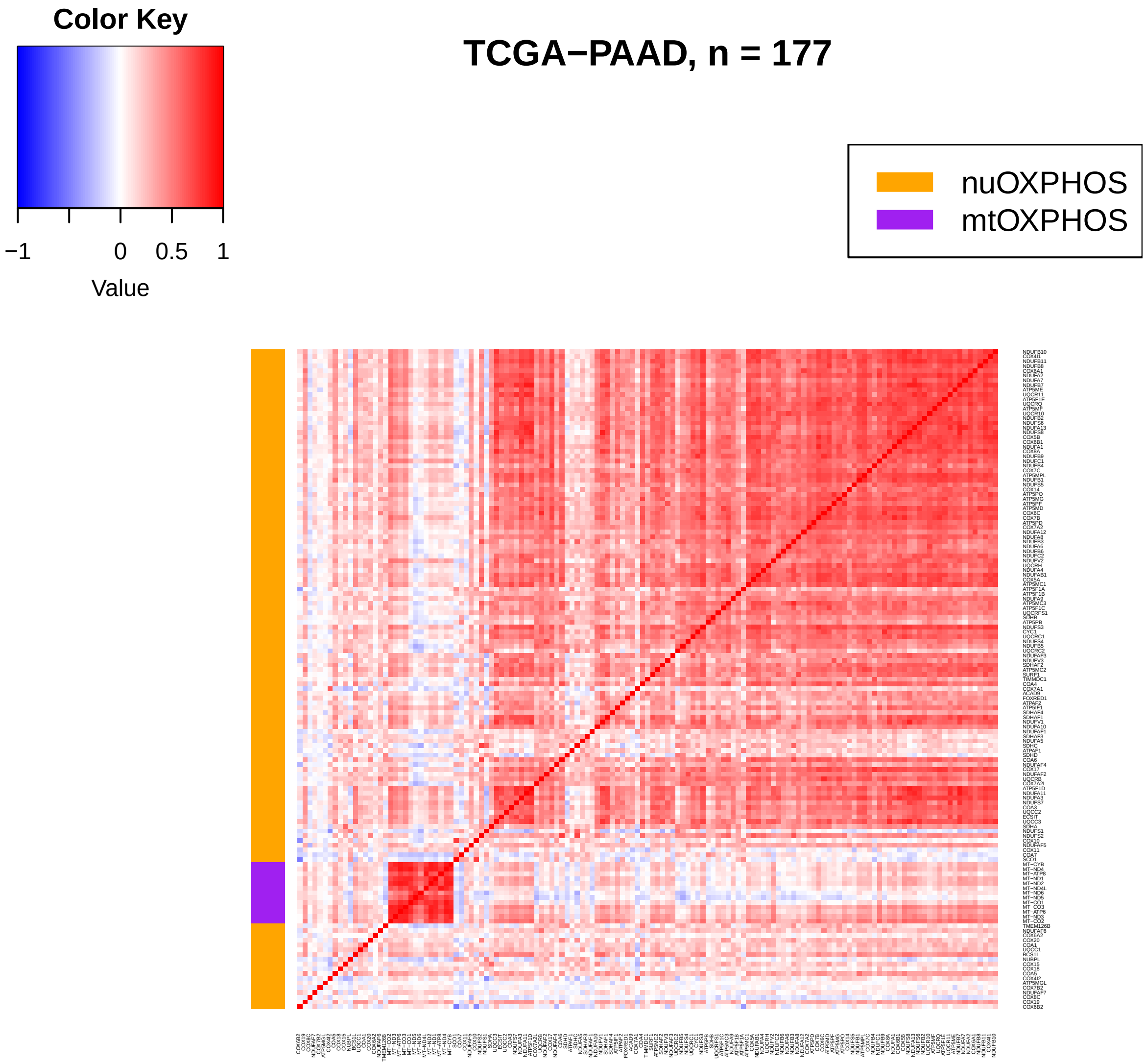
Color Key
−1
0
0.5
1
Value
TCGA−PAAD, n = 177
nuOXPHOS
mtOXPHOS

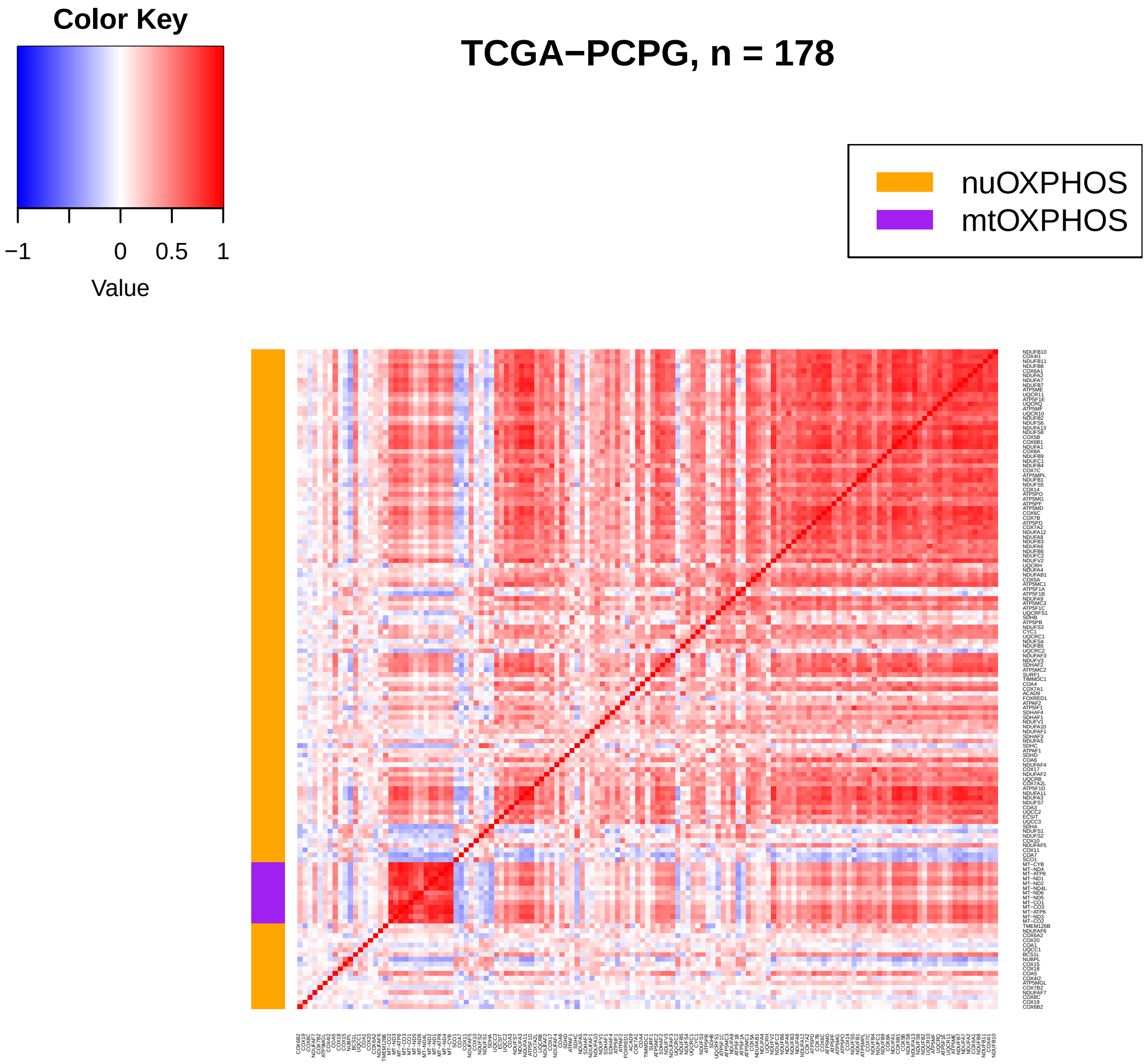

Color Key
−1
0
0.5
1
Value
TCGA−PCPG, n = 178
nuOXPHOS
mtOXPHOS

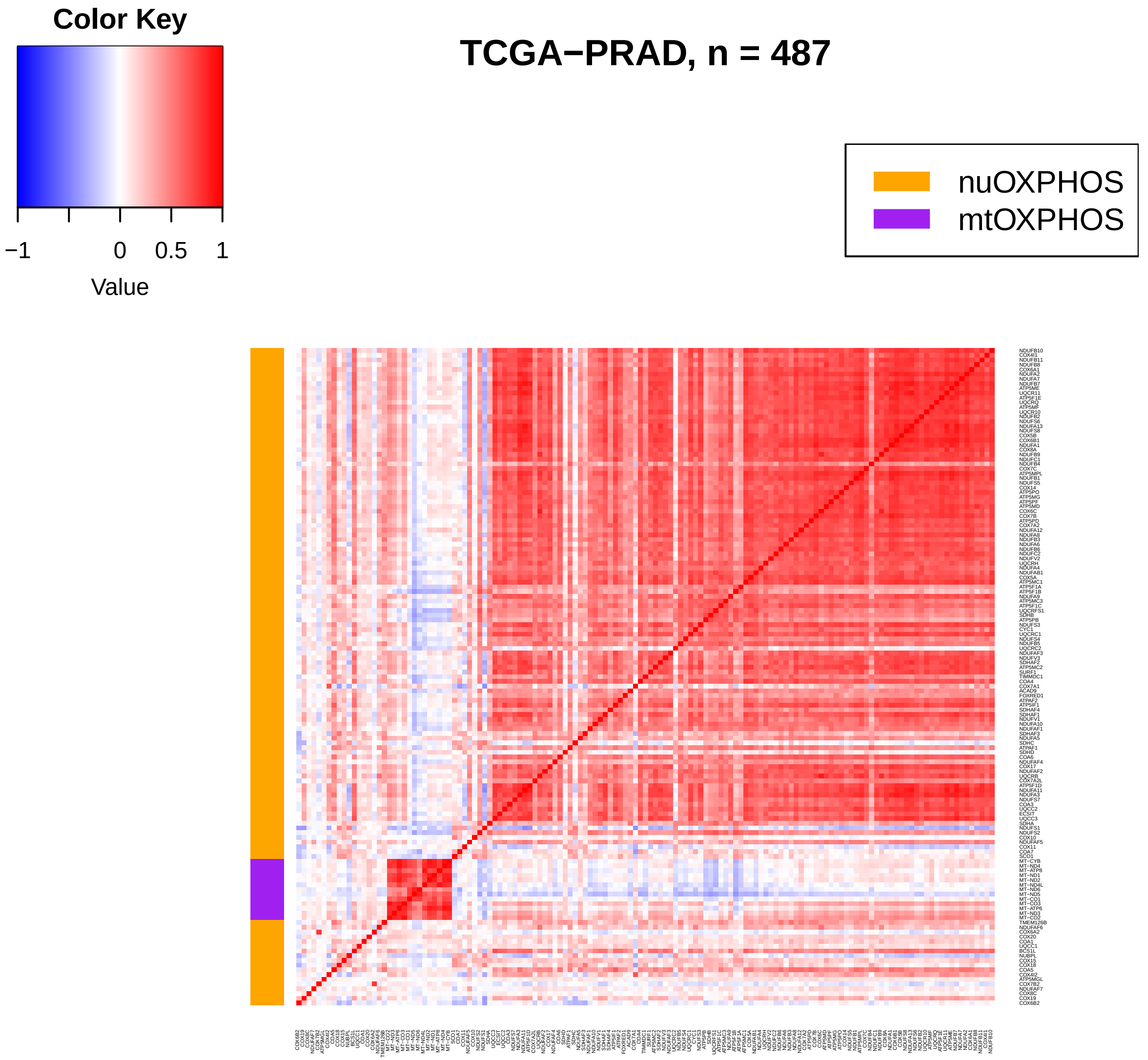
Color Key
−1
0
0.5
1
Value
TCGA−PRAD, n = 487
nuOXPHOS
mtOXPHOS

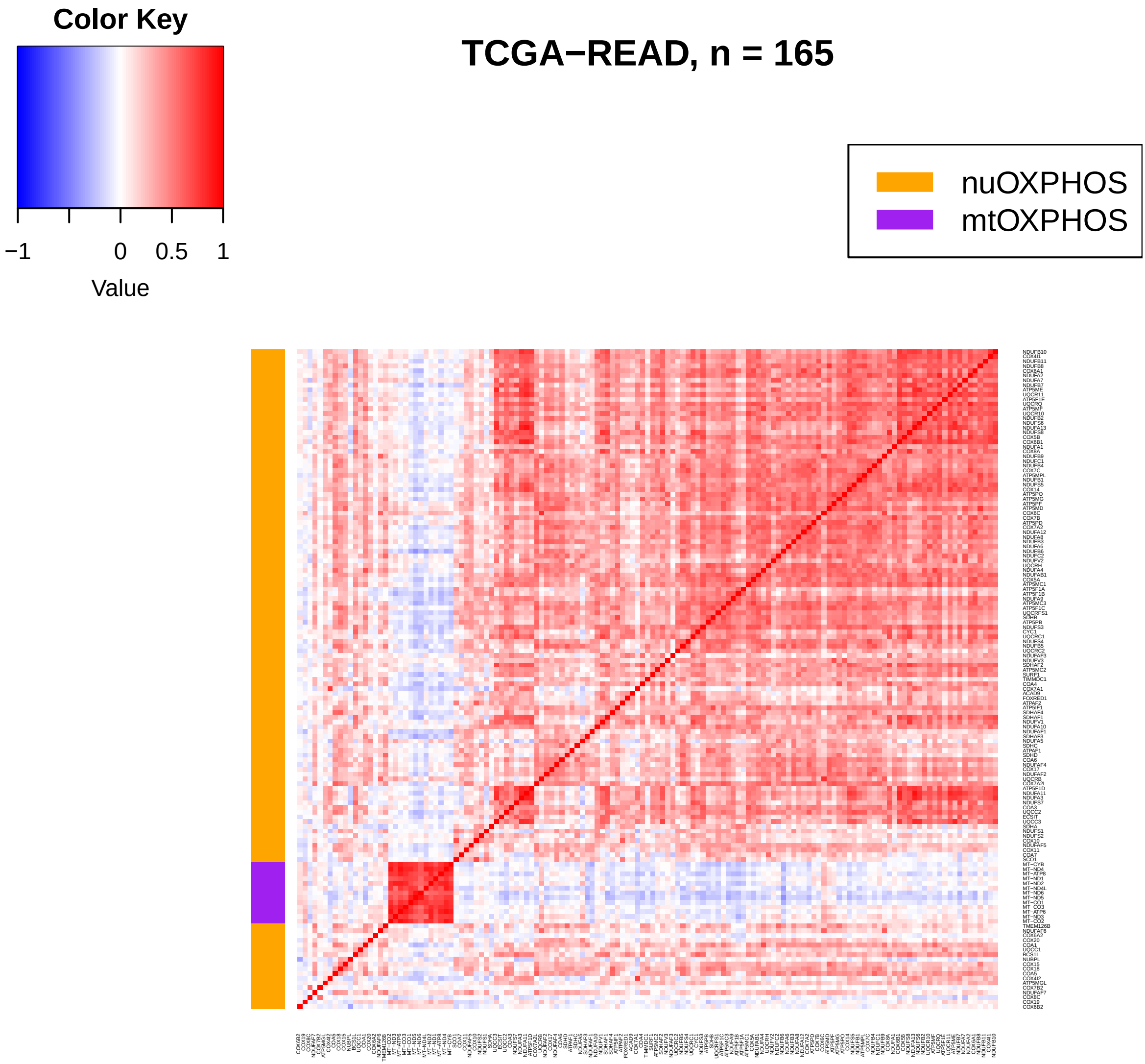
Color Key
−1
0
0.5
1
Value
TCGA−READ, n = 165
nuOXPHOS
mtOXPHOS

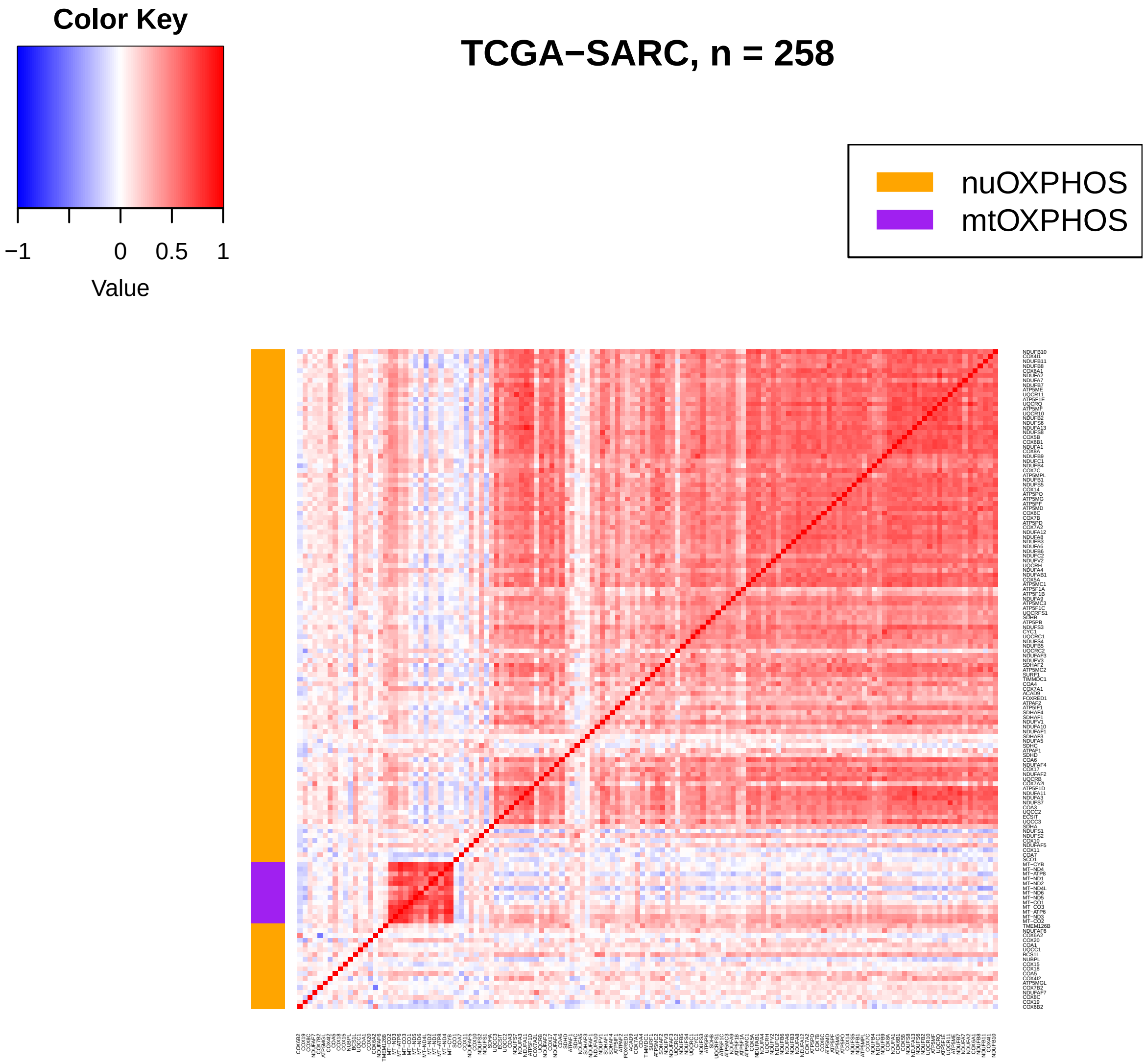

Color Key
−1
0
0.5
1
Value
TCGA−SARC, n = 258
nuOXPHOS
mtOXPHOS

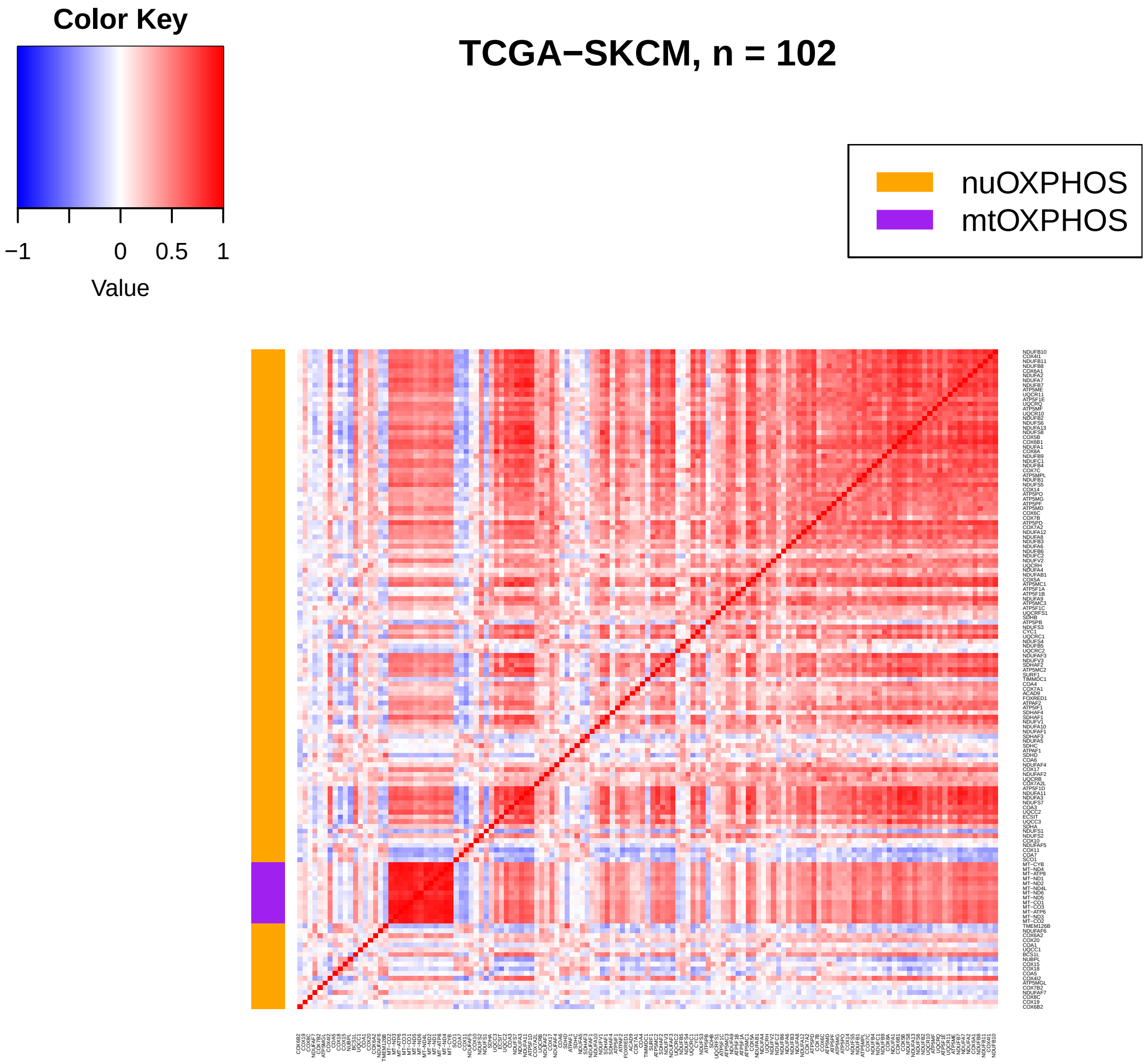
Color Key
−1
0
0.5
1
Value
TCGA−SKCM, n = 102
nuOXPHOS
mtOXPHOS

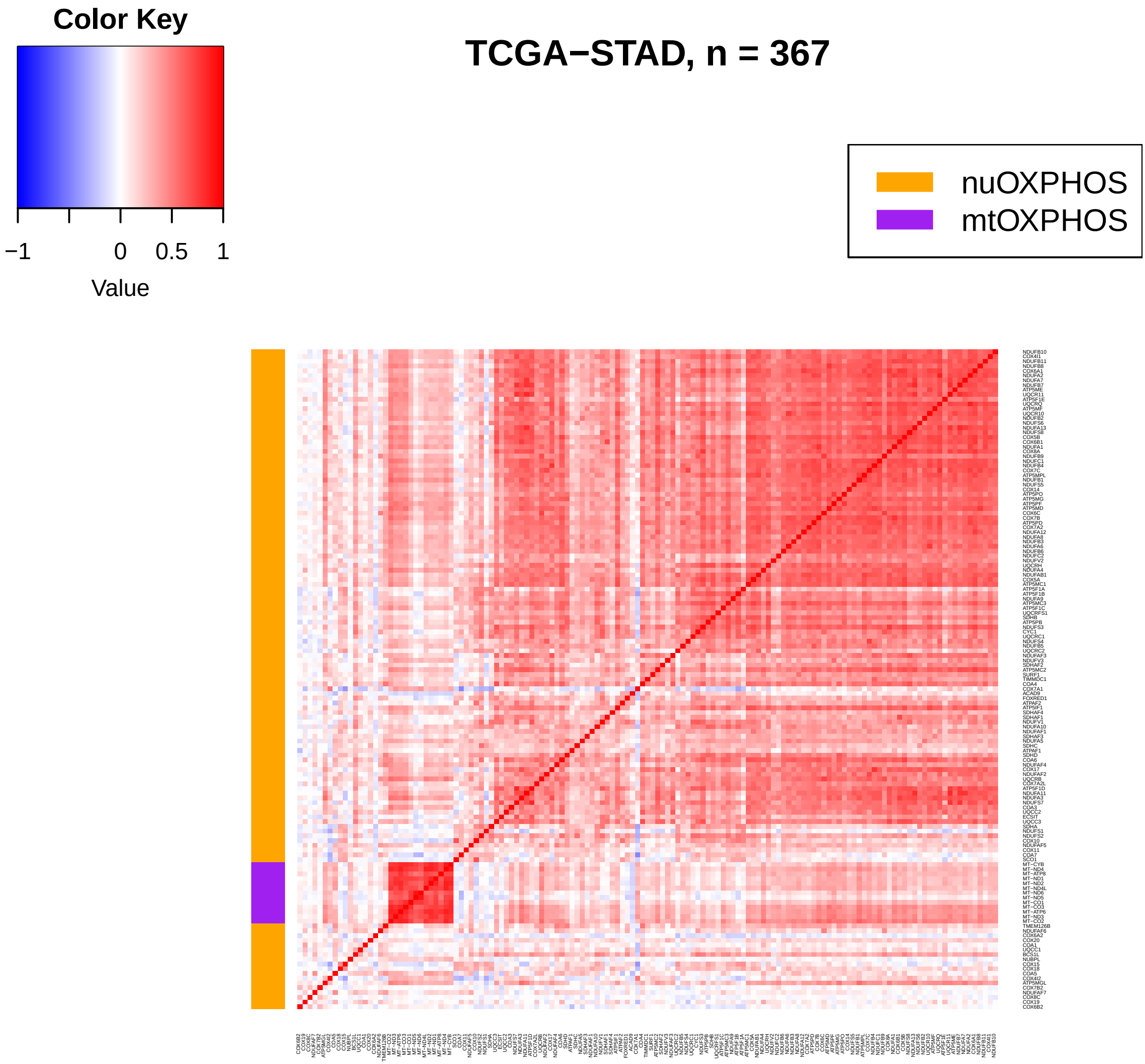
Color Key
−1
0
0.5
1
Value
TCGA−STAD, n = 367
nuOXPHOS
mtOXPHOS

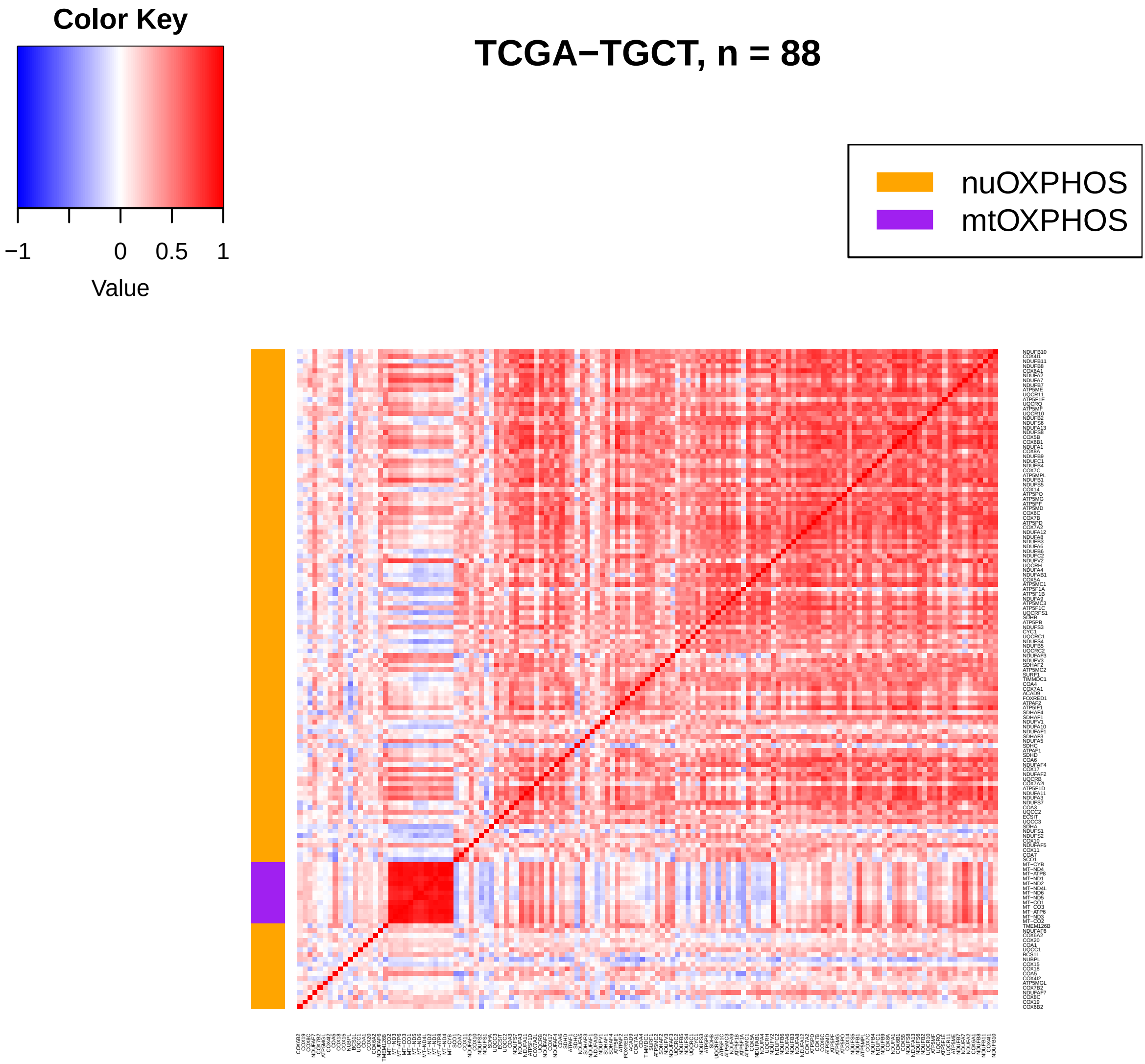
Color Key
−1
0
0.5
1
Value
TCGA−TGCT, n = 88
nuOXPHOS
mtOXPHOS

### TCGA−THCA, n = 502

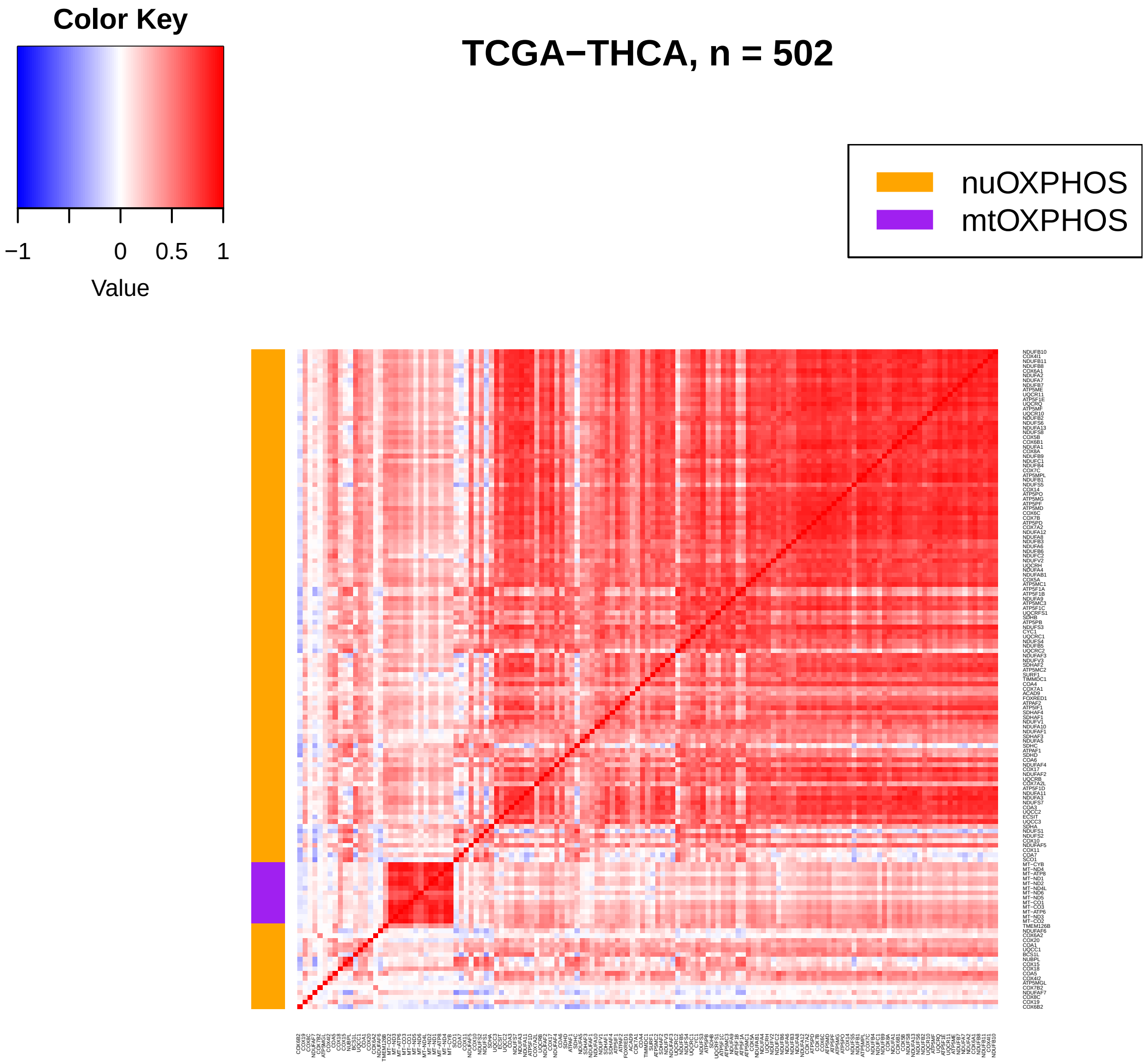

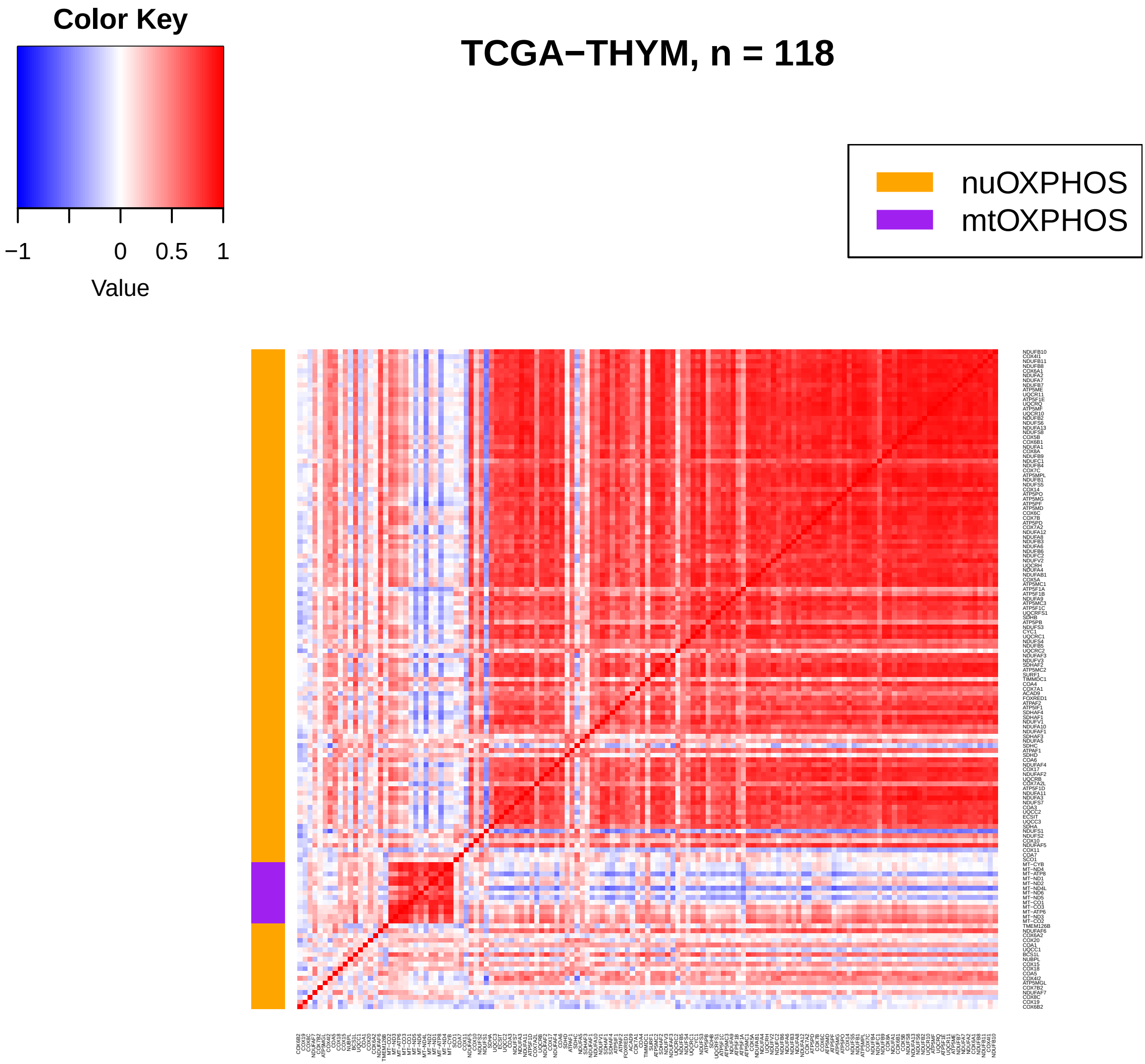
Color Key
−1
0
0.5
1
Value
TCGA−THYM, n = 118
nuOXPHOS
mtOXPHOS

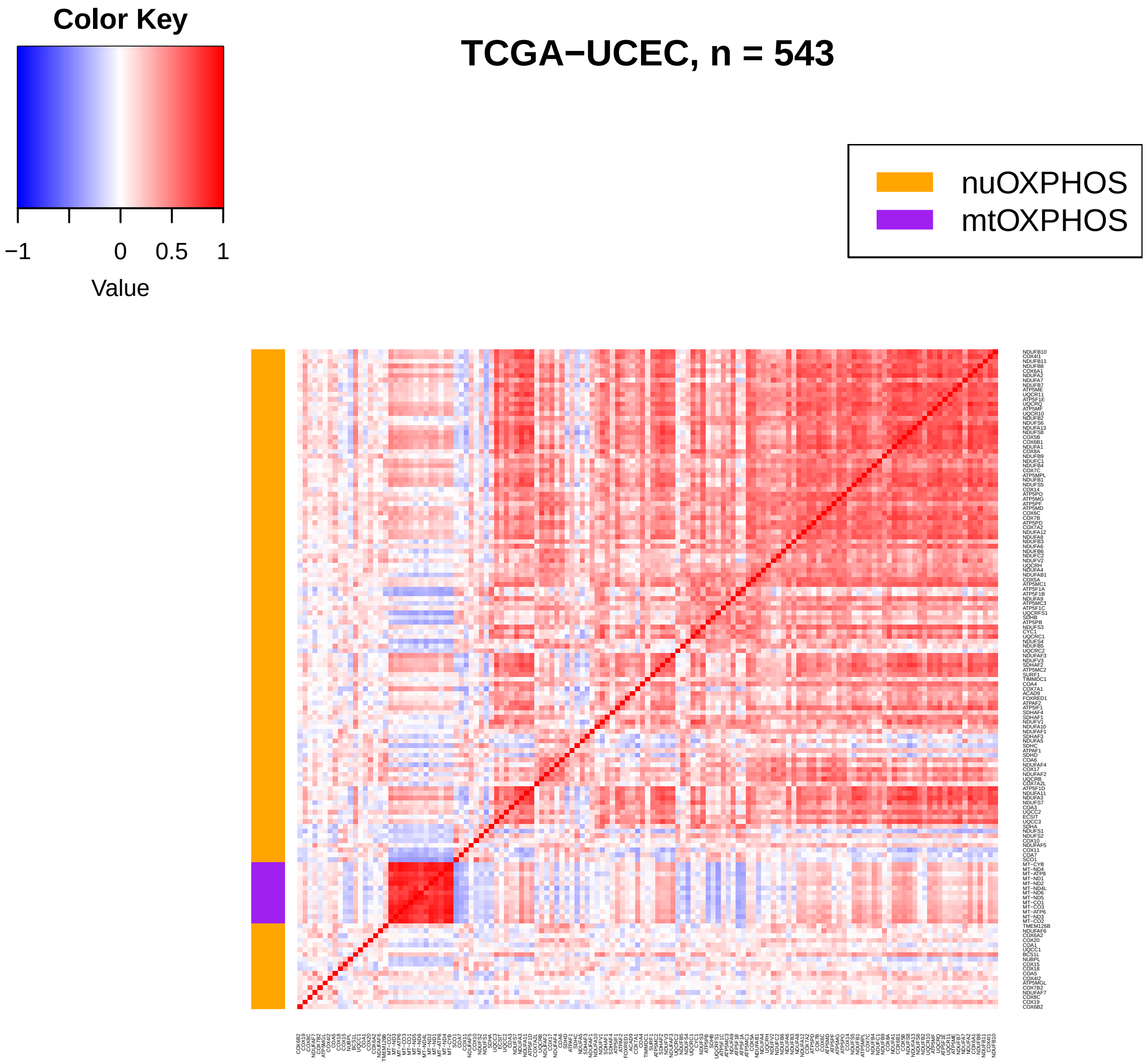
Color Key
−1
0
0.5
1
Value
TCGA−UCEC, n = 543
nuOXPHOS
mtOXPHOS

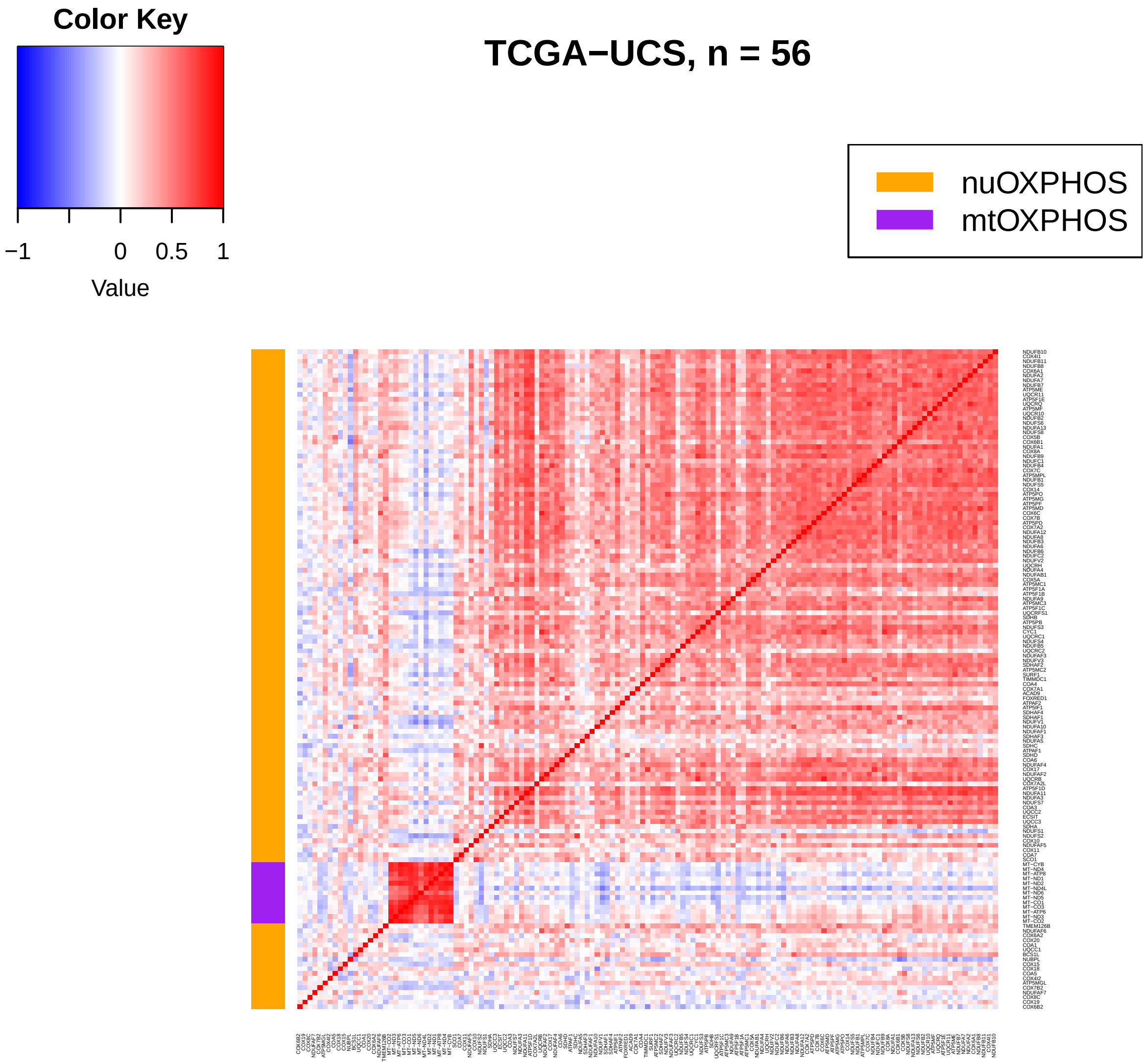

Color Key
−1
0
0.5
1
Value
TCGA−UCS, n = 56
nuOXPHOS
mtOXPHOS

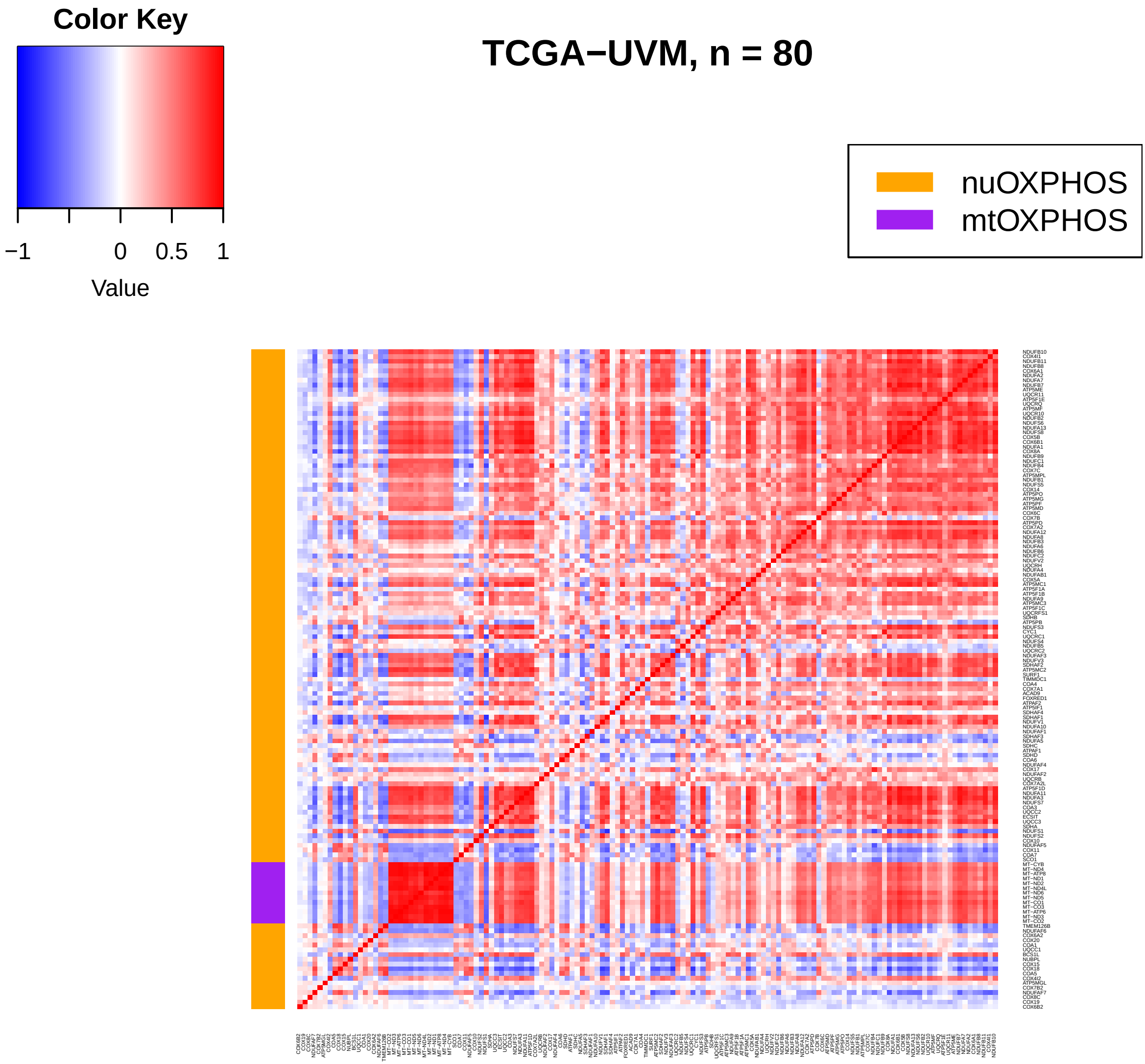
Color Key
−1
0
0.5
1
Value
TCGA−UVM, n = 80
nuOXPHOS
mtOXPHOS
