## Supplemental tables and scripts for "Malignancy and NF-kB signalling strengthen coordination between the expression of mitochondrial and nuclear-encoded oxidative phosphorylation genes": Supplementary-Dataset-4_MRN-mtOXPHOS-nuOXPHOS-matched-normal-heatmaps.pdf

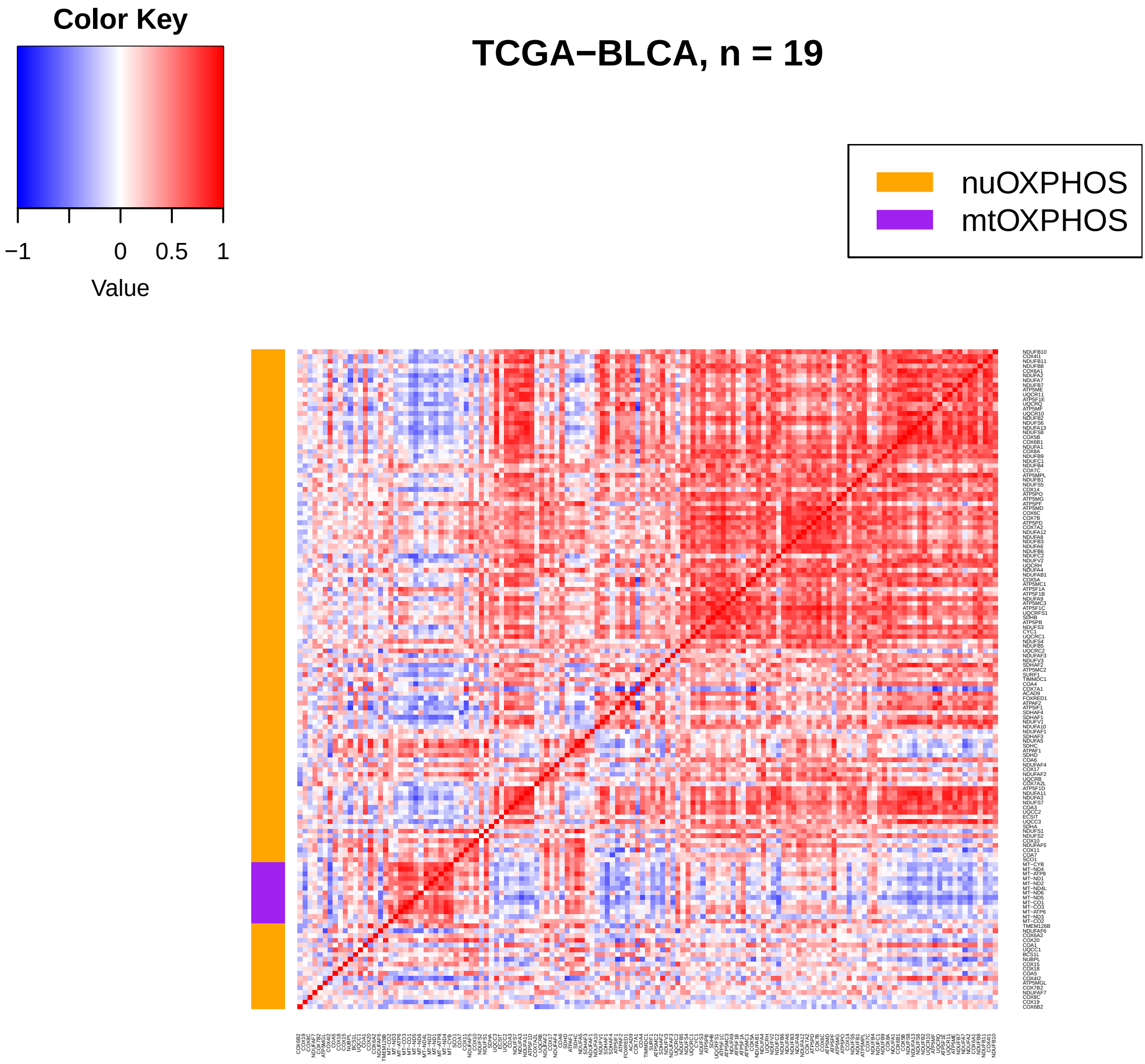
Color Key
−1
0
0.5
1
Value
TCGA−BLCA, n = 19
nuOXPHOS
mtOXPHOS

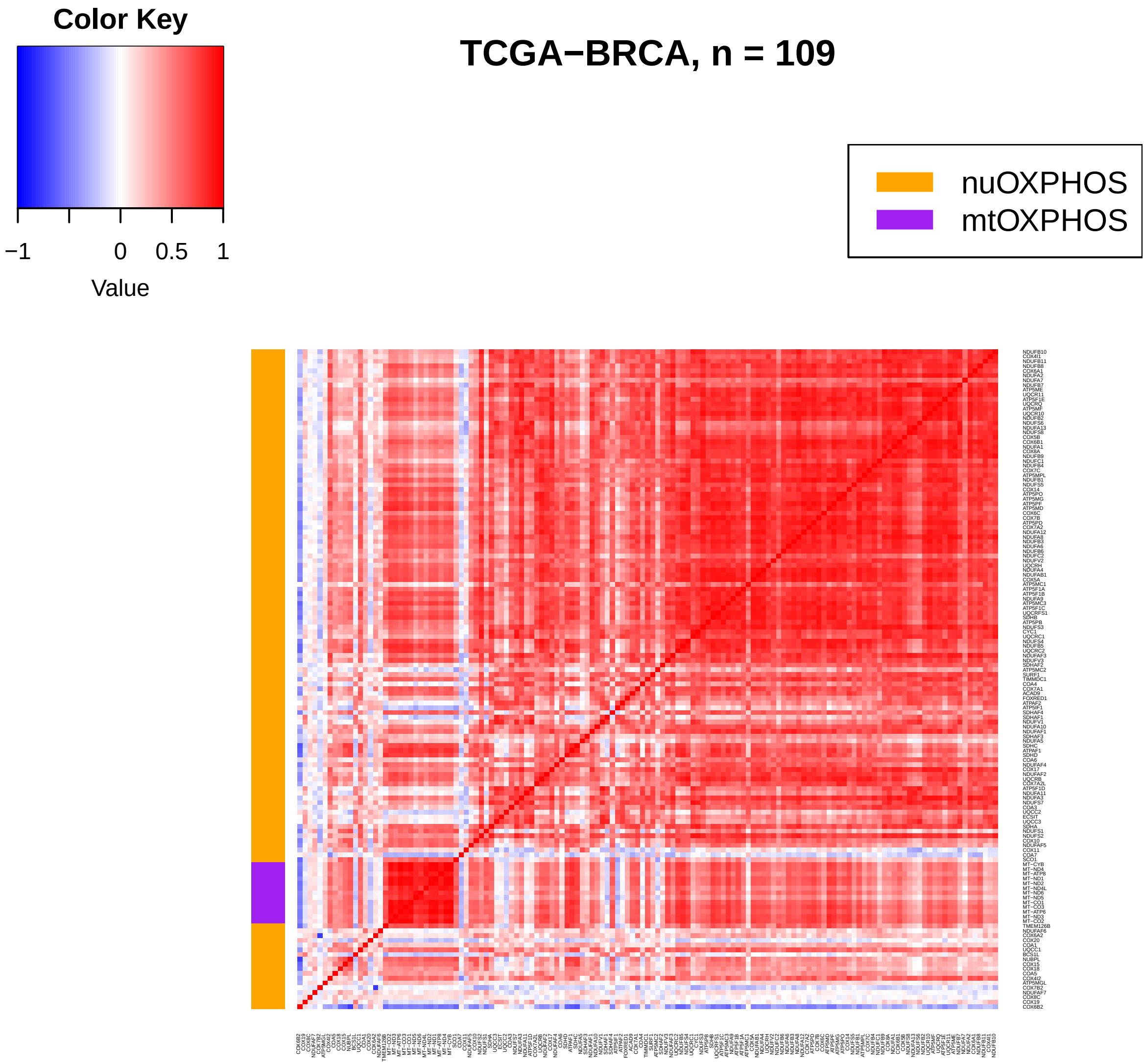

Color Key
−1
0
0.5
1
Value
TCGA−BRCA, n = 109
nuOXPHOS
mtOXPHOS

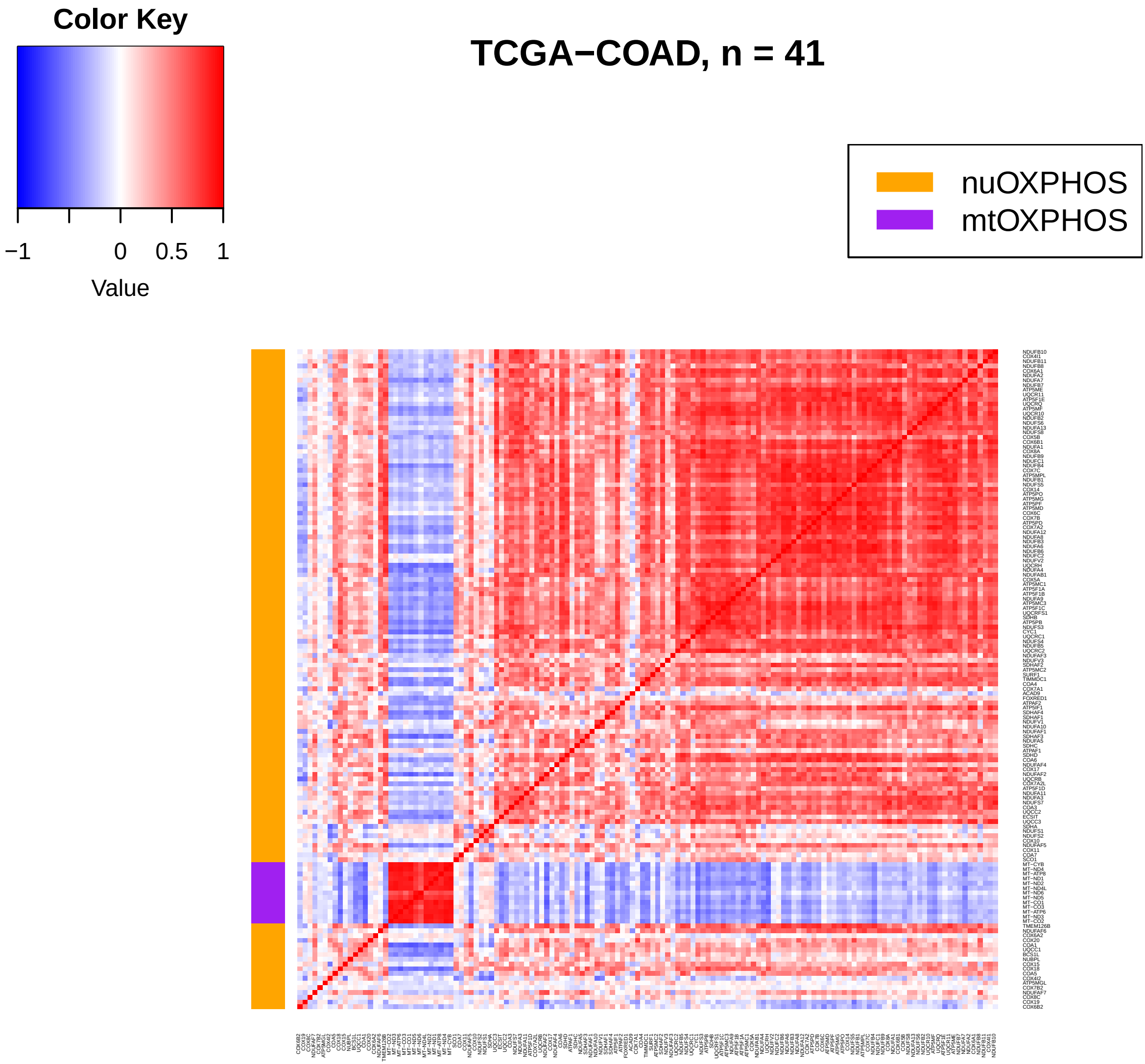
Color Key
−1
0
0.5
1
Value
TCGA−COAD, n = 41
nuOXPHOS
mtOXPHOS

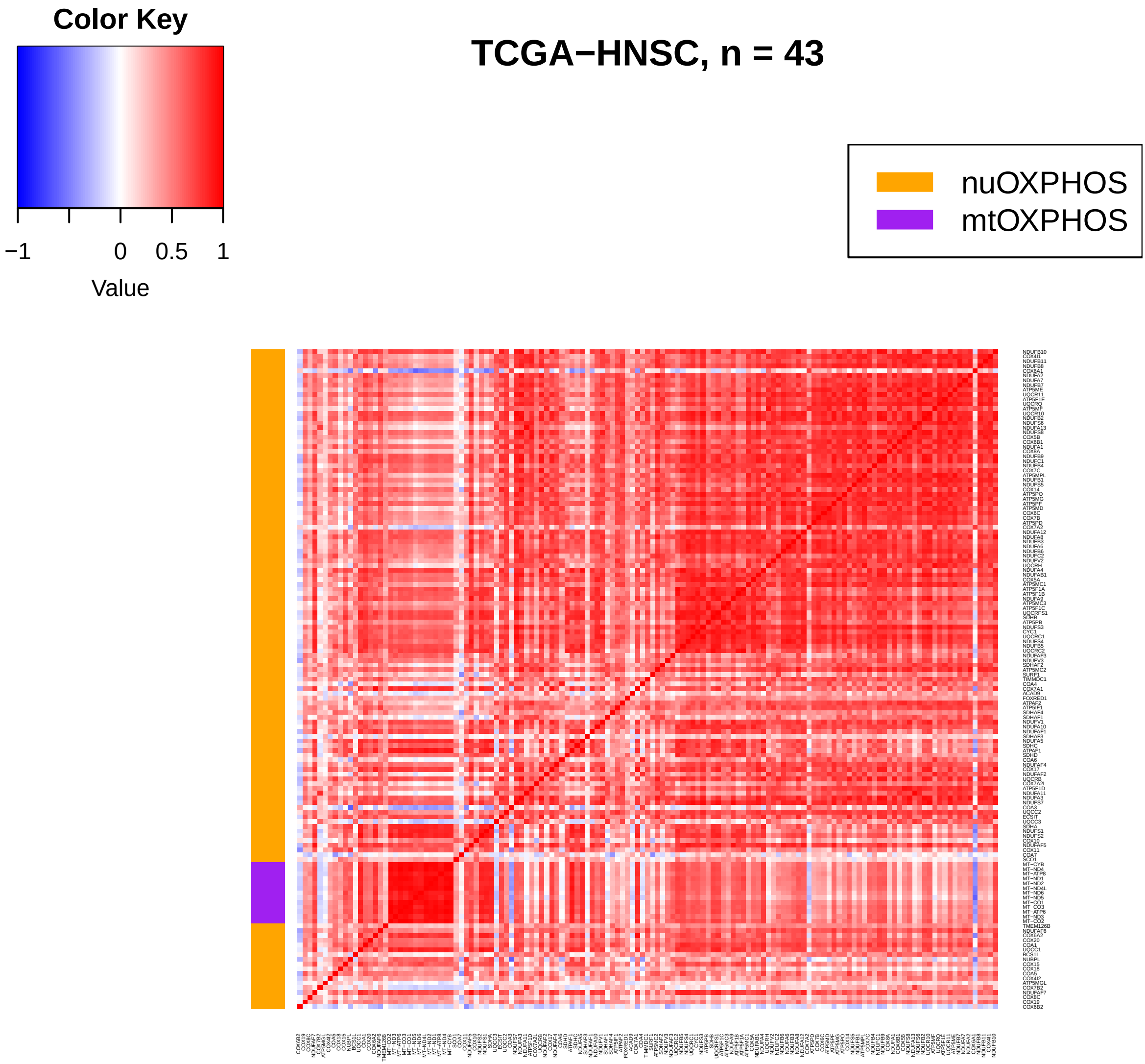
Color Key
−1
0
0.5
1
Value
TCGA−HNSC, n = 43
nuOXPHOS
mtOXPHOS

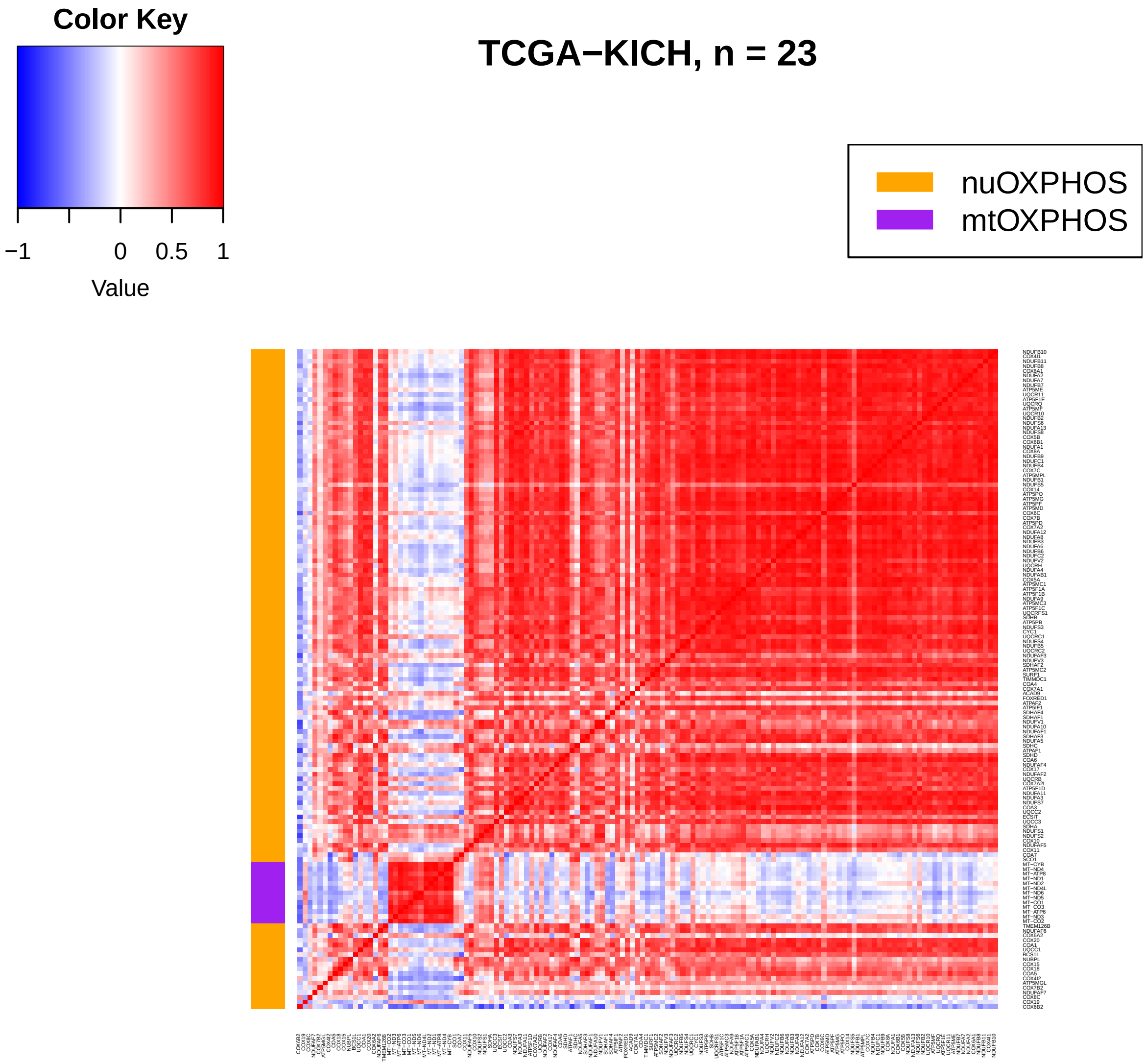

Color Key
−1
0
0.5
1
Value
TCGA−KICH, n = 23
nuOXPHOS
mtOXPHOS

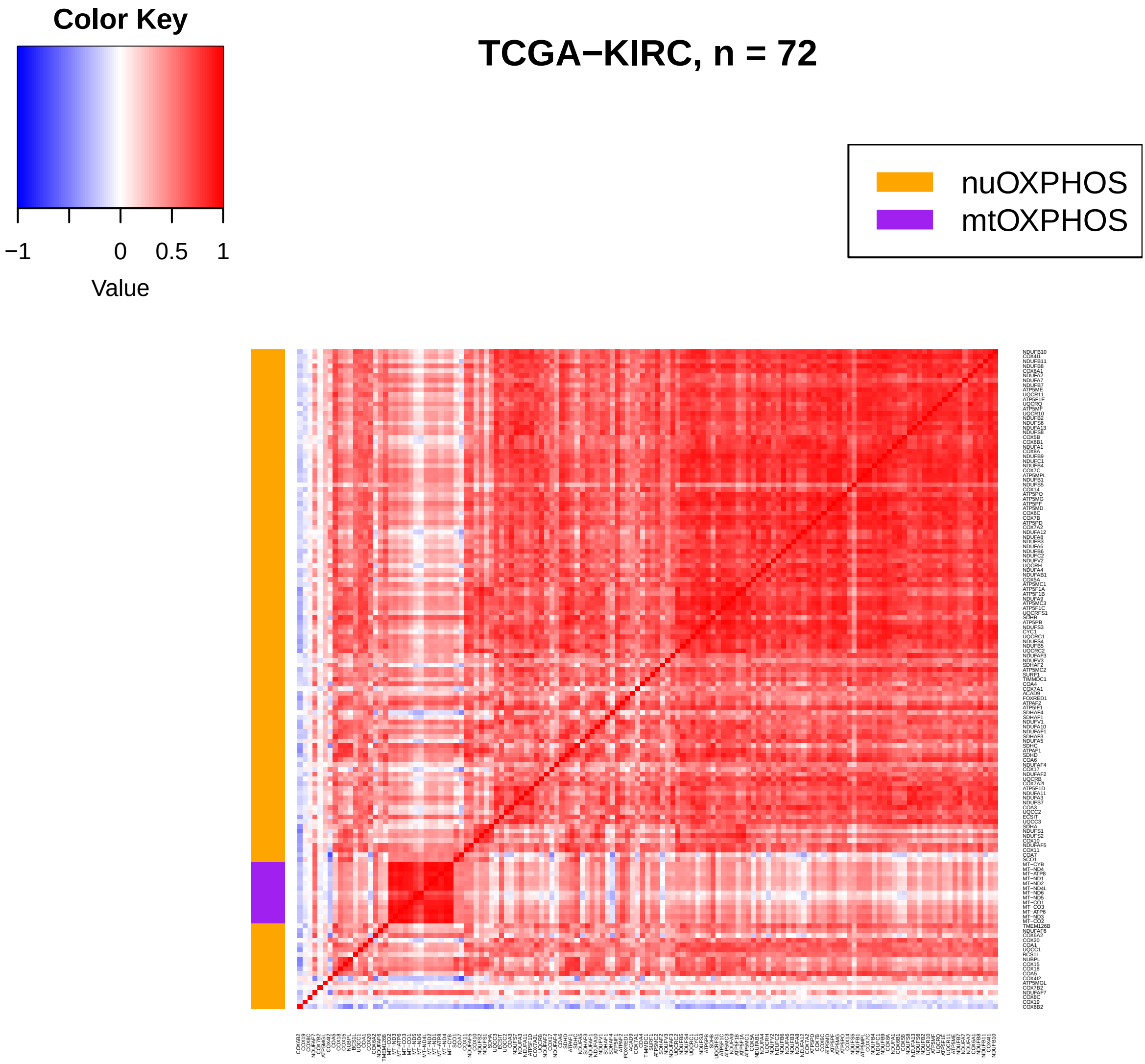

Color Key
−1
0
0.5
1
Value
TCGA−KIRC, n = 72
nuOXPHOS
mtOXPHOS

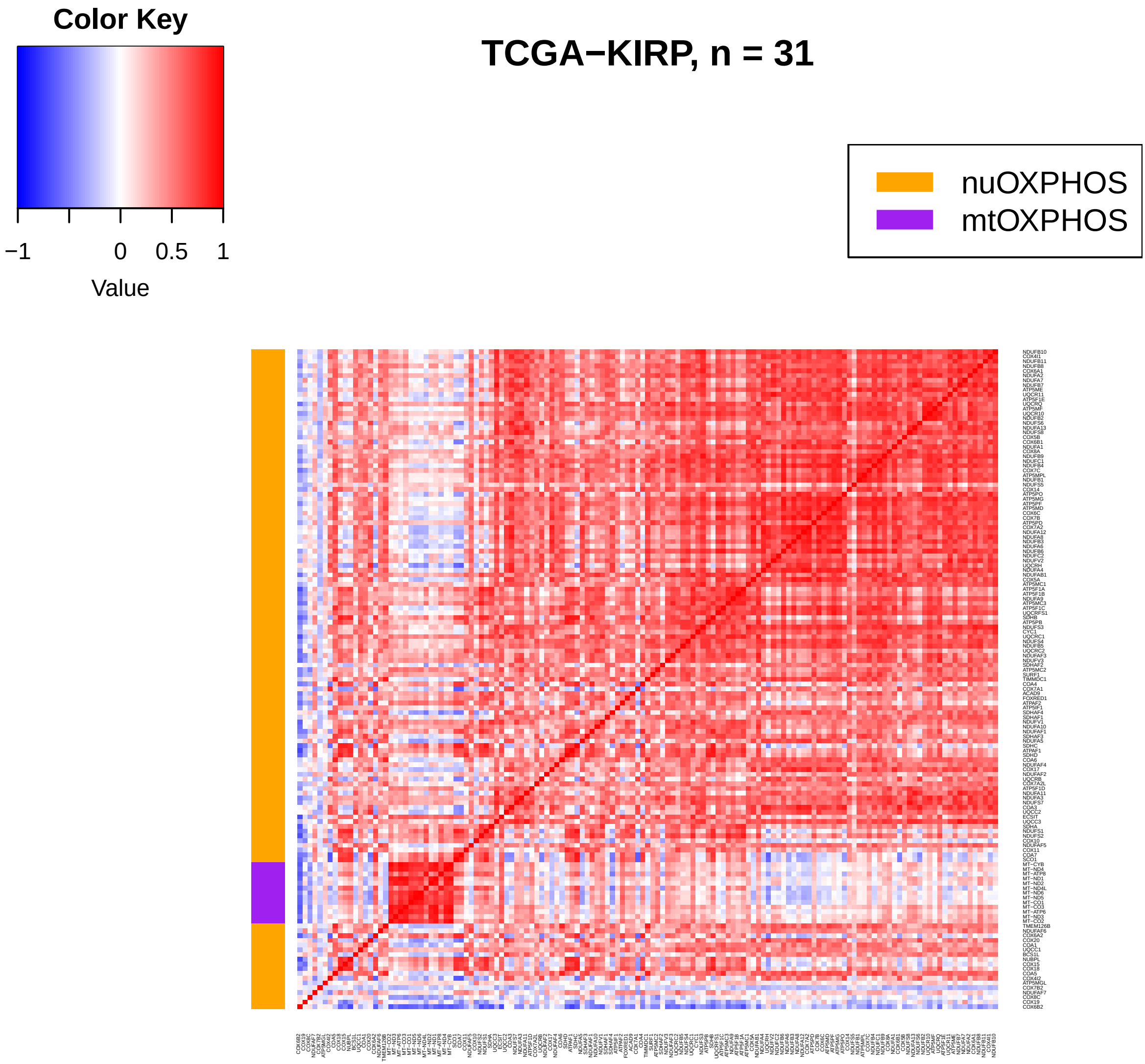
TCGA−KIRP, n = 31
Color Key
−1
0
0.5
1
Value
nuOXPHOS
mtOXPHOS

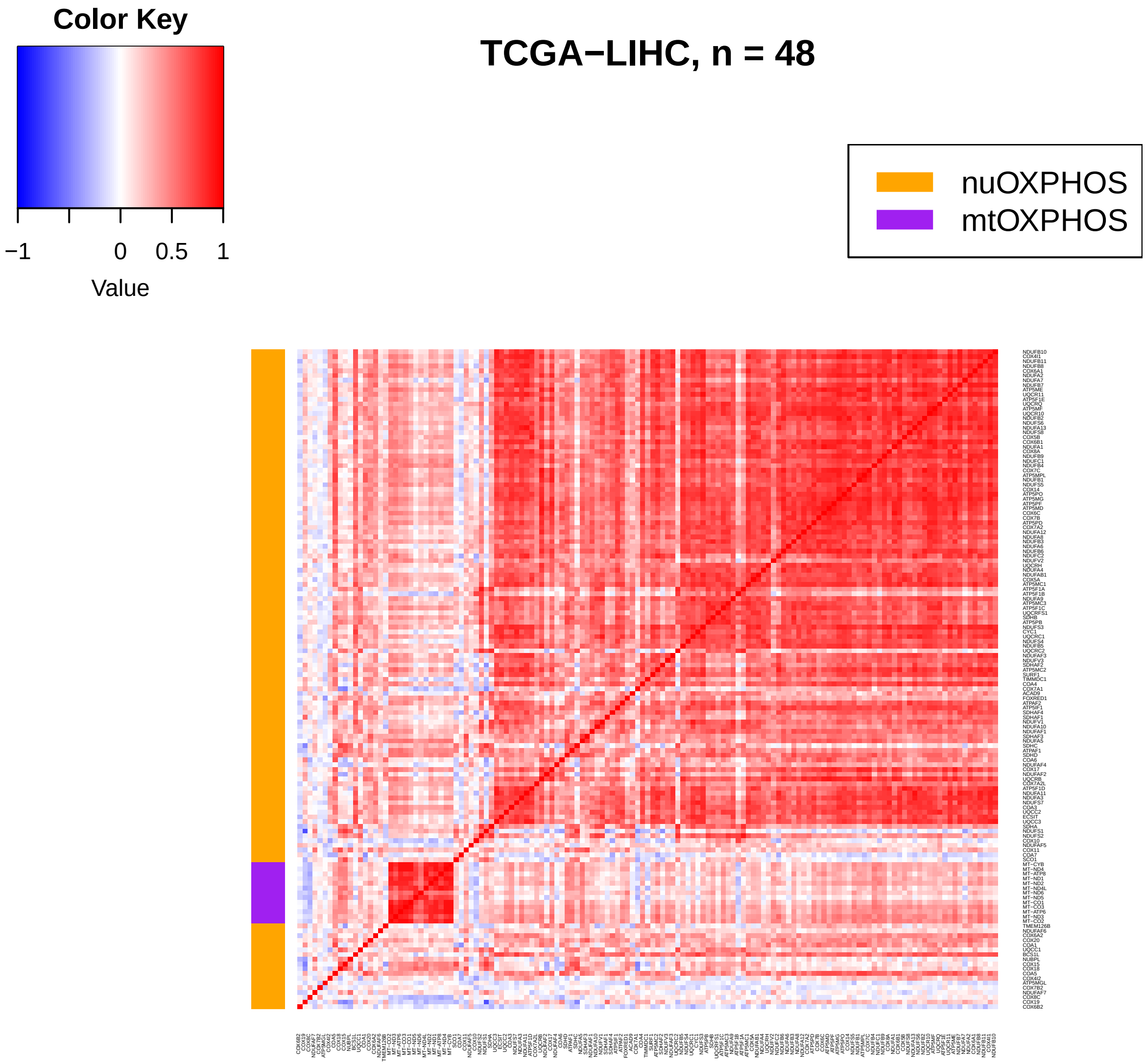
Color Key
−1
0
0.5
1
Value
TCGA−LIHC, n = 48
nuOXPHOS
mtOXPHOS

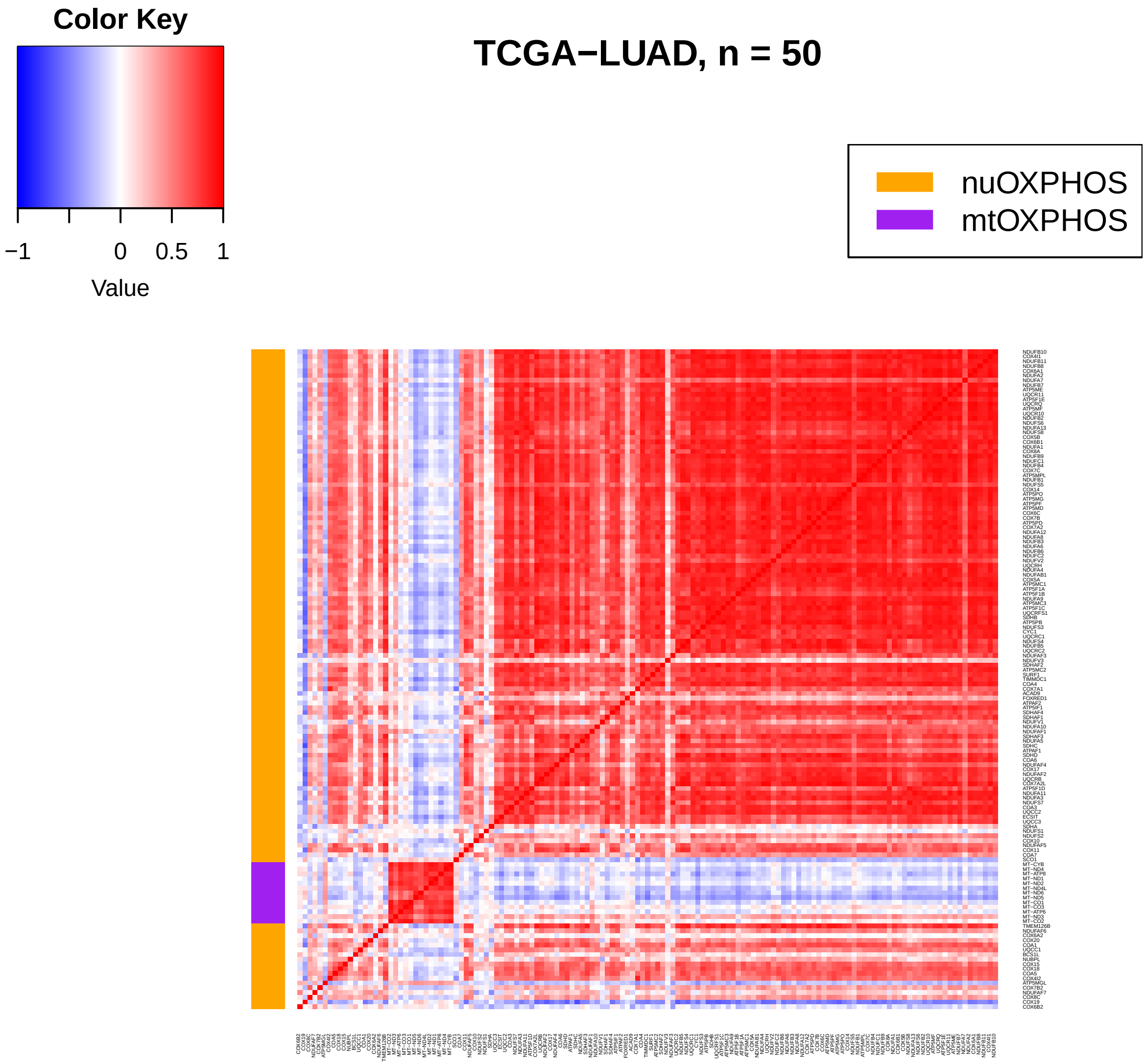
Color Key
−1
0
0.5
1
Value
TCGA−LUAD, n = 50
nuOXPHOS
mtOXPHOS

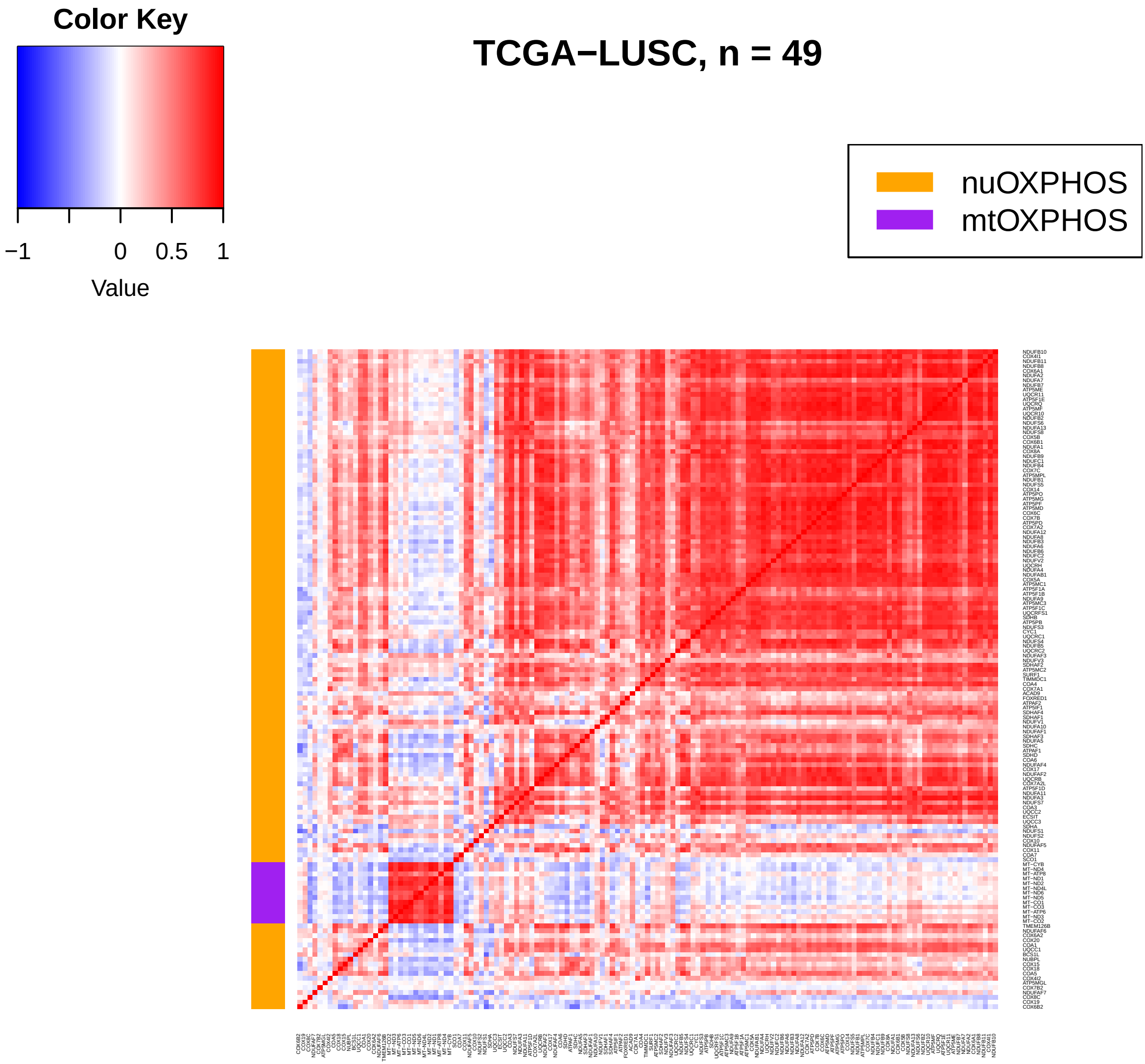

Color Key
−1
0
0.5
1
Value
TCGA−LUSC, n = 49
nuOXPHOS
mtOXPHOS

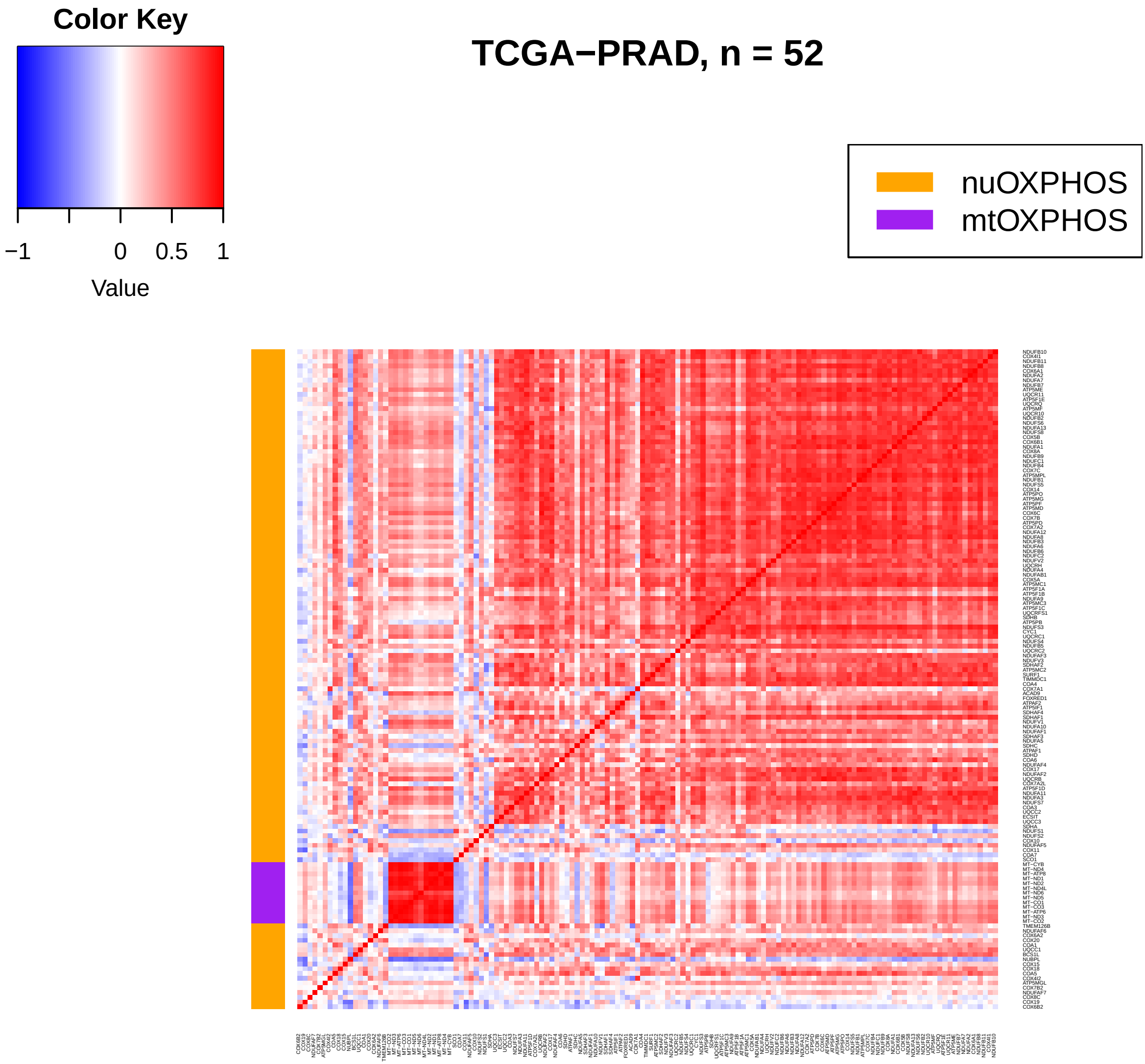
Color Key
−1
0
0.5
1
Value
TCGA−PRAD, n = 52
nuOXPHOS
mtOXPHOS

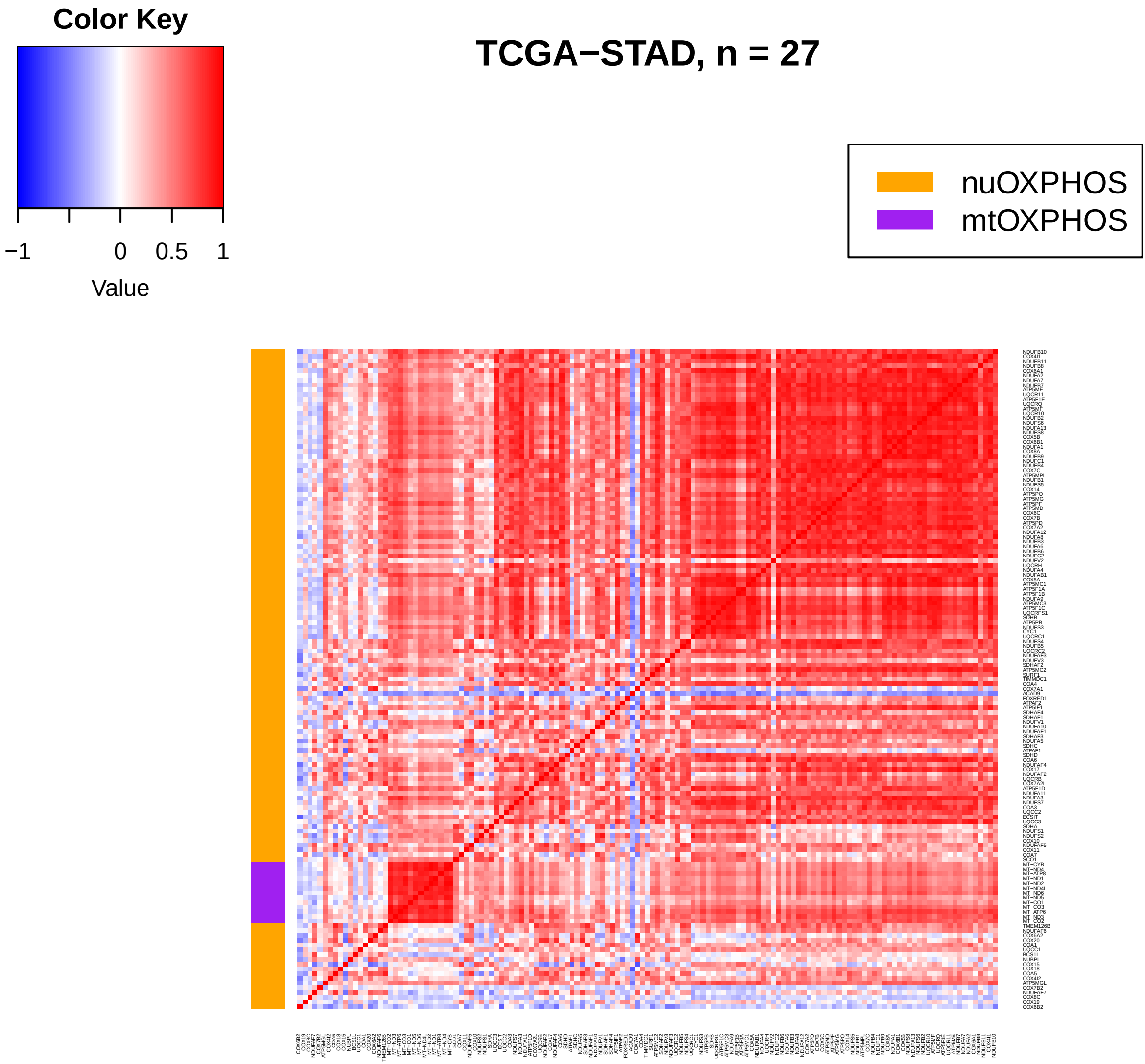
Color Key
−1
0
0.5
1
Value
TCGA−STAD, n = 27
nuOXPHOS
mtOXPHOS

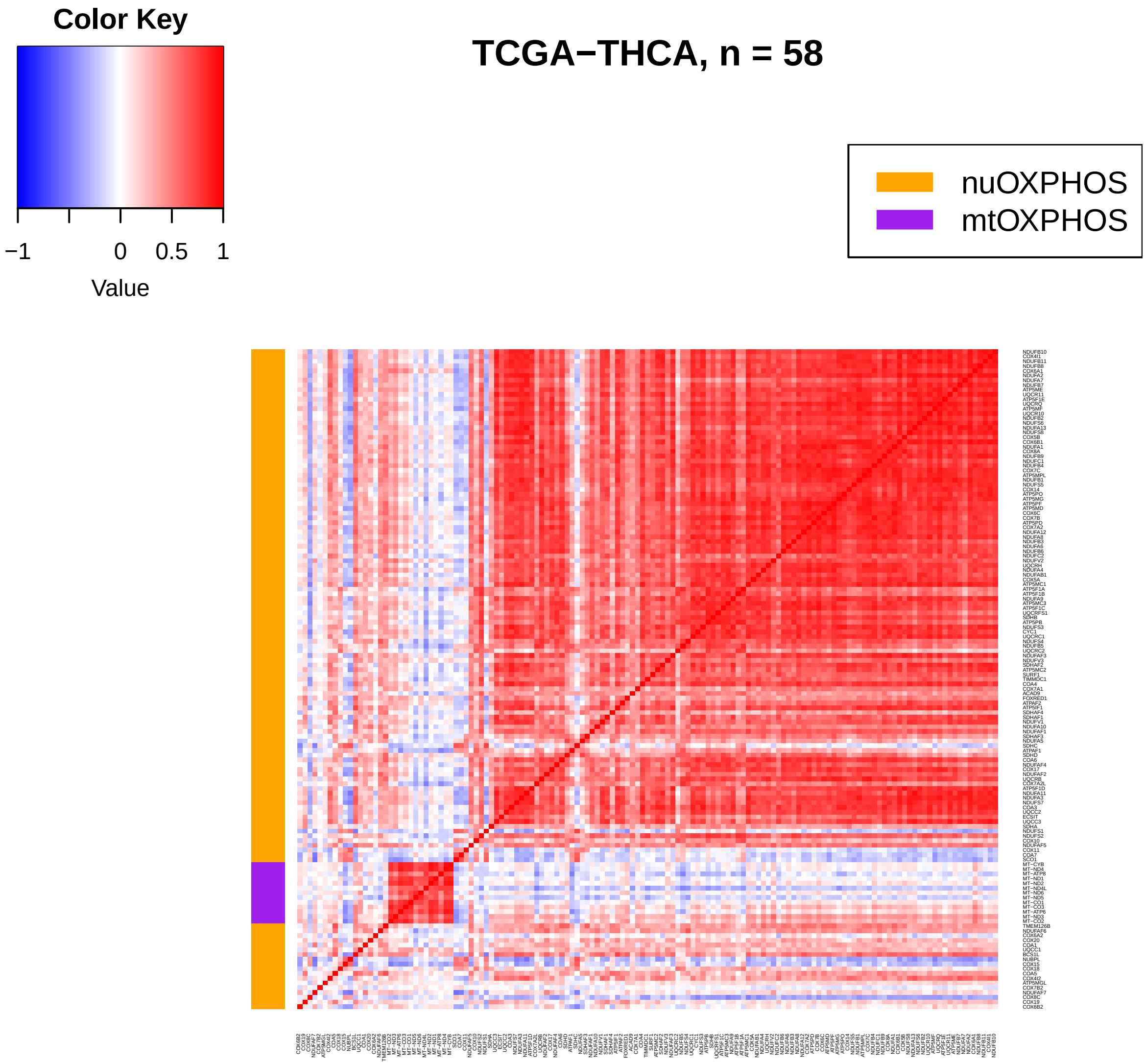

Color Key
−1
0
0.5
1
Value
TCGA−THCA, n = 58
nuOXPHOS
mtOXPHOS

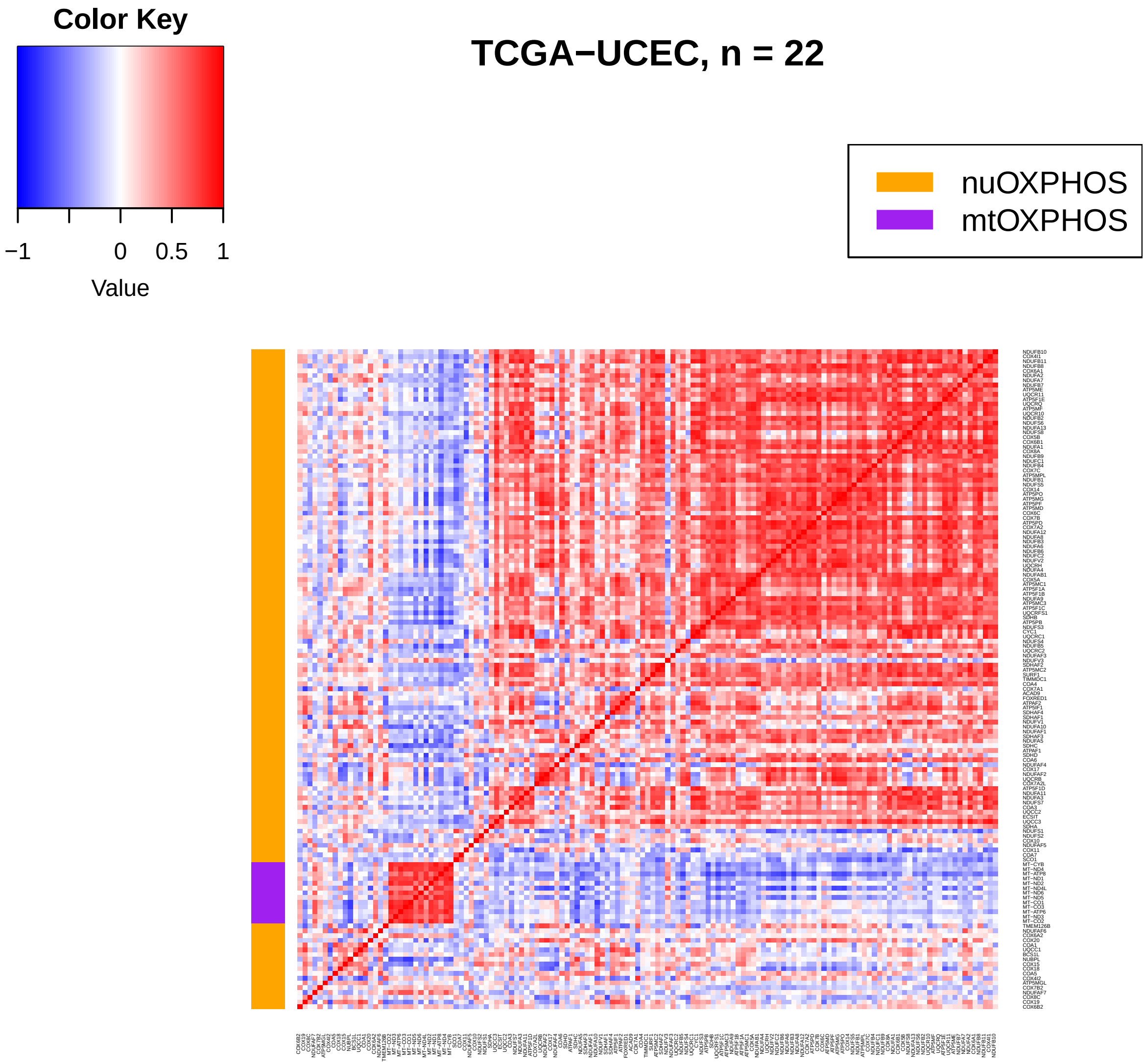
Color Key
−1
0
0.5
1
Value
TCGA−UCEC, n = 22
nuOXPHOS
mtOXPHOS
