## Supplemental tables and scripts for "Malignancy and NF-kB signalling strengthen coordination between the expression of mitochondrial and nuclear-encoded oxidative phosphorylation genes": Supplementary_text_1_code.R

```
#start with empty workspace

rm(list = ls(all = TRUE))

#### set working directory ####

# here create new folder and set working directory within it

dir.create("~/manuscript")
setwd("~/manuscript")

# create subfolders for input, output and graphics

dir.create("input")

# into input folder, add input files 

dir.create("output")
dir.create("graphics")


#### DEFINE FUNCTIONS ####

# apply linear models to counts functions

apply.lm.to.counts <- function(countdata, lookuptable = LUT) {
  
  # create data frame with expression data and explanatory variables (age, sex, death) for linear regression
  # data will be transposed to have genes in columns and sample in rows
  # 3 new columns will be added: age, sex, cause of death for each sample
  temp_df <- as.data.frame(matrix(nrow = ncol(countdata), ncol = nrow(countdata) + 5))
  
  colnames(temp_df) <- c(row.names(countdata), "age_bracket", "sex", "death", "ischemic_time", "batch")
  row.names(temp_df) <- colnames(countdata)
  
  # insert tranposed data into new data frame
  temp_df[, 1:nrow(countdata)] <- t(countdata)
  
  # insert age bracket for each sample
  # use lookup table created above (lookuptable)
  temp_df[, "age_bracket"] <- lookuptable[row.names(temp_df), "age_bracket"]
  
  # for each bracket, replace by midpoint to treat as a numerical explanatory variable, rather than a factor
  temp_df[, "age_bracket"] <- str_replace_all(temp_df$age_bracket, pattern = "20-29", replacement = "25")
  temp_df[, "age_bracket"] <- str_replace_all(temp_df$age_bracket, pattern = "30-39", replacement = "35")
  temp_df[, "age_bracket"] <- str_replace_all(temp_df$age_bracket, pattern = "40-49", replacement = "45")
  temp_df[, "age_bracket"] <- str_replace_all(temp_df$age_bracket, pattern = "50-59", replacement = "55")
  temp_df[, "age_bracket"] <- str_replace_all(temp_df$age_bracket, pattern = "60-69", replacement = "65")
  temp_df[, "age_bracket"] <- str_replace_all(temp_df$age_bracket, pattern = "70-79", replacement = "75")
  
  # make column numeric, not a character string
  temp_df$age_bracket <- as.numeric(temp_df$age_bracket)
  
  # from lookup table, add Sex and Death variables
  temp_df[, "sex"] <- lookuptable[row.names(temp_df), "Sex"]
  temp_df[, "death"] <- lookuptable[row.names(temp_df), "DeathScale"]
  
  # replace death = NA (unknown cause of death) with a 5th factor level
  temp_df[is.na(temp_df$death), "death"] <- 5
  
  temp_df[, "ischemic_time"] <- lookuptable[row.names(temp_df), "IschemicTime"]
  
  temp_df[, "batch"] <- lookuptable[row.names(temp_df), "Batch"]
  
  # check for any missing values and remove entry if so
  # here additional clause to work with GTEX v6, which has 4 brain tissues with no values for ischemic time
  # we aim to check if the tissue has NO values for ischemic time. in which case we will leave it
  # if it has some values, then will remove any entries with missing values
  
  if(sum(!is.na(temp_df$ischemic_time)) > 0){
    
    if(sum(is.na(temp_df$ischemic_time)) > 0){
      
      temp_df <- temp_df[!(is.na(temp_df$ischemic_time)), ]
      
    }
    
  }
  
  # do linear regression across all samples (all tissues combined)
  # perform linear regression with age (bracket midpoint), sex and cause of death
  
  # create dataframe to deposit residuals
  residuals_df <- as.data.frame(matrix(nrow = nrow(temp_df), ncol = ncol(temp_df) - 5))
  row.names(residuals_df) <- row.names(temp_df)
  colnames(residuals_df) <- colnames(temp_df)[1:(ncol(temp_df) - 5)]
  
  # loop over columns (genes); for each gene perform linear regression and deposit residuals in new data frame
  # sex is considered a factor automatically as a character string; as.factor() must be specified for cause of death
  
  if(sum(!is.na(temp_df$ischemic_time)) > 0){
  
  if((length(unique(temp_df$sex)) == 1) & (length(unique(temp_df$batch)) > 1)){
    
    for(j in 1:(ncol(temp_df) - 5)){
      
      residuals_df[, colnames(temp_df)[j]] <- lm(temp_df[,j] ~ age_bracket + ischemic_time + as.factor(death) + batch, data = temp_df)$residuals
      
    }
    
  } 
  
  if((length(unique(temp_df$sex)) > 1) & (length(unique(temp_df$batch)) > 1)){
    
    for(j in 1:(ncol(temp_df) - 5)){
      
      residuals_df[, colnames(temp_df)[j]] <- lm(temp_df[,j] ~ age_bracket + ischemic_time + sex + as.factor(death) + batch, data = temp_df)$residuals
      
    }
  }
  
  if((length(unique(temp_df$sex)) == 1) & (length(unique(temp_df$batch)) == 1)){
    
    for(j in 1:(ncol(temp_df) - 5)){
      
      residuals_df[, colnames(temp_df)[j]] <- lm(temp_df[,j] ~ age_bracket + ischemic_time + as.factor(death), data = temp_df)$residuals
      
    }
    
  }
  
  if((length(unique(temp_df$sex)) > 1) & (length(unique(temp_df$batch)) == 1)){
    
    for(j in 1:(ncol(temp_df) - 5)){
      
      residuals_df[, colnames(temp_df)[j]] <- lm(temp_df[,j] ~ age_bracket + ischemic_time + sex + as.factor(death), data = temp_df)$residuals
      
    }
  }
  }
  
  if(sum(is.na(temp_df$ischemic_time)) > 0){
    
    if((length(unique(temp_df$sex)) == 1) & (length(unique(temp_df$batch)) > 1)){
      
      for(j in 1:(ncol(temp_df) - 5)){
        
        residuals_df[, colnames(temp_df)[j]] <- lm(temp_df[,j] ~ age_bracket + as.factor(death) + batch, data = temp_df)$residuals
        
      }
      
    } 
    
    if((length(unique(temp_df$sex)) > 1) & (length(unique(temp_df$batch)) > 1)){
      
      for(j in 1:(ncol(temp_df) - 5)){
        
        residuals_df[, colnames(temp_df)[j]] <- lm(temp_df[,j] ~ age_bracket + sex + as.factor(death) + batch, data = temp_df)$residuals
        
      }
    }
    
    if((length(unique(temp_df$sex)) == 1) & (length(unique(temp_df$batch)) == 1)){
      
      for(j in 1:(ncol(temp_df) - 5)){
        
        residuals_df[, colnames(temp_df)[j]] <- lm(temp_df[,j] ~ age_bracket + as.factor(death), data = temp_df)$residuals
        
      }
      
    }
    
    if((length(unique(temp_df$sex)) > 1) & (length(unique(temp_df$batch)) == 1)){
      
      for(j in 1:(ncol(temp_df) - 5)){
        
        residuals_df[, colnames(temp_df)[j]] <- lm(temp_df[,j] ~ age_bracket + sex + as.factor(death), data = temp_df)$residuals
        
      }
    }
  }
  
  return(t(residuals_df))
  
}

TCGA.apply.lm.to.counts <- function(countdata, lookuptable = TCGA_LUT) {
  
  # create data frame with expression data and explanatory variables (age, sex, death) for linear regression
  # data will be transposed to have genes in columns and sample in rows
  # 5 new columns will be added for variables of interest
  
  # restrict to samples present in lookuptable 
  tempcountdata <- countdata[, colnames(countdata) %in% lookuptable$SAMPID]
  
  temp_df <- as.data.frame(matrix(nrow = ncol(tempcountdata), ncol = nrow(tempcountdata) + 5))
  
  colnames(temp_df) <- c(row.names(tempcountdata), "race", "gender", "tumour_stage", "sequencing_centre", "days_to_birth")
  row.names(temp_df) <- colnames(tempcountdata)
  
  # insert tranposed data into new data frame
  temp_df[, 1:nrow(tempcountdata)] <- t(tempcountdata)
  
  # from lookup table, add variables
  temp_df[, "days_to_birth"] <- as.numeric(lookuptable[row.names(temp_df), "days_to_birth"])
  temp_df[, "gender"] <- lookuptable[row.names(temp_df), "gender"]
  temp_df[, "race"] <- lookuptable[row.names(temp_df), "race"]
  temp_df[, "tumour_stage"] <- lookuptable[row.names(temp_df), "tumour_stage"]
  temp_df[, "sequencing_centre"] <- lookuptable[row.names(temp_df), "sequencing_centre"]
  
  # check for any missing values and remove entry if so
  temp_df <- temp_df[!(is.na(temp_df$days_to_birth)), ]
  temp_df <- temp_df[!(is.na(temp_df$gender)), ]
  temp_df <- temp_df[!(is.na(temp_df$race)), ]
  temp_df <- temp_df[!(is.na(temp_df$tumour_stage)), ]
  temp_df <- temp_df[!(is.na(temp_df$sequencing_centre)), ]
  
  
  # do linear regression across all samples (all tissues combined)
  # perform linear regression with age (bracket midpoint), sex and cause of death
  
  # create dataframe to deposit residuals
  residuals_df <- as.data.frame(matrix(nrow = nrow(temp_df), ncol = ncol(temp_df) - 5))
  row.names(residuals_df) <- row.names(temp_df)
  colnames(residuals_df) <- colnames(temp_df)[1:(ncol(temp_df) - 5)]
  
  # loop over columns (genes); for each gene perform linear regression and deposit residuals in new data frame
  # sex is considered a factor automatically as a character string; sequencing centre is already considered as factor
  # here we note that in some projects, all sequencing centres are equal. Likewise gender for some cancer tpyes (e.g. ovarian)
  # need conditional to ignore seq centre, gender or tumour_stage if only one level exists among considered data in order to avoid an error
  
  if(length(unique(temp_df$race)) > 1){
    
    if(length(unique(temp_df$tumour_stage)) == 1) {
      
      if((length(unique(temp_df$gender)) == 1) & (length(unique(temp_df$sequencing_centre)) > 1)){
        
        for(j in 1:(ncol(temp_df) - 5)){
          
          residuals_df[, colnames(temp_df)[j]] <- lm(temp_df[, j] ~ days_to_birth + race + sequencing_centre, data = temp_df)$residuals
          
        }
        
      } 
      
      if((length(unique(temp_df$gender)) == 1) & (length(unique(temp_df$sequencing_centre)) == 1)){
        
        for(j in 1:(ncol(temp_df) - 5)){
          
          residuals_df[, colnames(temp_df)[j]] <- lm(temp_df[,j] ~ days_to_birth + race, data = temp_df)$residuals
          
        }
      }
      
      if((length(unique(temp_df$gender)) > 1) & (length(unique(temp_df$sequencing_centre)) > 1)){
        
        for(j in 1:(ncol(temp_df) - 5)){
          
          residuals_df[, colnames(temp_df)[j]] <- lm(temp_df[,j] ~ days_to_birth + race + gender + sequencing_centre, data = temp_df)$residuals
          
        }
      }
      
      if((length(unique(temp_df$gender)) > 1) & (length(unique(temp_df$sequencing_centre)) == 1)){
        
        for(j in 1:(ncol(temp_df) - 5)){
          
          residuals_df[, colnames(temp_df)[j]] <- lm(temp_df[,j] ~ days_to_birth + race + gender, data = temp_df)$residuals
          
        }
      }
    }
    
    if(length(unique(temp_df$tumour_stage)) > 1) {
      
      if((length(unique(temp_df$gender)) == 1) & (length(unique(temp_df$sequencing_centre)) > 1)){
        
        for(j in 1:(ncol(temp_df) - 5)){
          
          residuals_df[, colnames(temp_df)[j]] <- lm(temp_df[, j] ~ days_to_birth + race + sequencing_centre + tumour_stage, data = temp_df)$residuals
          
        }
        
      } 
      
      if((length(unique(temp_df$gender)) == 1) & (length(unique(temp_df$sequencing_centre)) == 1)){
        
        for(j in 1:(ncol(temp_df) - 5)){
          
          residuals_df[, colnames(temp_df)[j]] <- lm(temp_df[,j] ~ days_to_birth + race + tumour_stage, data = temp_df)$residuals
          
        }
      }
      
      if((length(unique(temp_df$gender)) > 1) & (length(unique(temp_df$sequencing_centre)) > 1)){
        
        for(j in 1:(ncol(temp_df) - 5)){
          
          residuals_df[, colnames(temp_df)[j]] <- lm(temp_df[,j] ~ days_to_birth + race + gender + sequencing_centre + tumour_stage, data = temp_df)$residuals
          
        }
      }
      
      if((length(unique(temp_df$gender)) > 1) & (length(unique(temp_df$sequencing_centre)) == 1)){
        
        for(j in 1:(ncol(temp_df) - 5)){
          
          residuals_df[, colnames(temp_df)[j]] <- lm(temp_df[,j] ~ days_to_birth + race + gender + tumour_stage, data = temp_df)$residuals
          
        }
      }
    }
  }
  
  if(length(unique(temp_df$race) == 1)){
    
    if(length(unique(temp_df$tumour_stage)) == 1) {
      
      if((length(unique(temp_df$gender)) == 1) & (length(unique(temp_df$sequencing_centre)) > 1)){
        
        for(j in 1:(ncol(temp_df) - 5)){
          
          residuals_df[, colnames(temp_df)[j]] <- lm(temp_df[, j] ~ days_to_birth + sequencing_centre, data = temp_df)$residuals
          
        }
        
      } 
      
      if((length(unique(temp_df$gender)) == 1) & (length(unique(temp_df$sequencing_centre)) == 1)){
        
        for(j in 1:(ncol(temp_df) - 5)){
          
          residuals_df[, colnames(temp_df)[j]] <- lm(temp_df[,j] ~ days_to_birth, data = temp_df)$residuals
          
        }
      }
      
      if((length(unique(temp_df$gender)) > 1) & (length(unique(temp_df$sequencing_centre)) > 1)){
        
        for(j in 1:(ncol(temp_df) - 5)){
          
          residuals_df[, colnames(temp_df)[j]] <- lm(temp_df[,j] ~ days_to_birth + gender + sequencing_centre, data = temp_df)$residuals
          
        }
      }
      
      if((length(unique(temp_df$gender)) > 1) & (length(unique(temp_df$sequencing_centre)) == 1)){
        
        for(j in 1:(ncol(temp_df) - 5)){
          
          residuals_df[, colnames(temp_df)[j]] <- lm(temp_df[,j] ~ days_to_birth + gender, data = temp_df)$residuals
          
        }
      }
    }
    
    if(length(unique(temp_df$tumour_stage)) > 1) {
      
      if((length(unique(temp_df$gender)) == 1) & (length(unique(temp_df$sequencing_centre)) > 1)){
        
        for(j in 1:(ncol(temp_df) - 5)){
          
          residuals_df[, colnames(temp_df)[j]] <- lm(temp_df[, j] ~ days_to_birth + sequencing_centre + tumour_stage, data = temp_df)$residuals
          
        }
        
      } 
      
      if((length(unique(temp_df$gender)) == 1) & (length(unique(temp_df$sequencing_centre)) == 1)){
        
        for(j in 1:(ncol(temp_df) - 5)){
          
          residuals_df[, colnames(temp_df)[j]] <- lm(temp_df[,j] ~ days_to_birth + tumour_stage, data = temp_df)$residuals
          
        }
      }
      
      if((length(unique(temp_df$gender)) > 1) & (length(unique(temp_df$sequencing_centre)) > 1)){
        
        for(j in 1:(ncol(temp_df) - 5)){
          
          residuals_df[, colnames(temp_df)[j]] <- lm(temp_df[,j] ~ days_to_birth + gender + sequencing_centre + tumour_stage, data = temp_df)$residuals
          
        }
      }
      
      if((length(unique(temp_df$gender)) > 1) & (length(unique(temp_df$sequencing_centre)) == 1)){
        
        for(j in 1:(ncol(temp_df) - 5)){
          
          residuals_df[, colnames(temp_df)[j]] <- lm(temp_df[,j] ~ days_to_birth + gender + tumour_stage, data = temp_df)$residuals
          
        }
      }
    }
  }
  
  
  return(t(residuals_df))
  
}

# map2color function

map2color <- function(x, 
                      pal,
                      limits = NULL){
  
  if(is.null(limits)){
    limits = range(x)
  }
  
  pal[findInterval(x, seq(limits[1], limits[2], length.out = length(pal) + 1), all.inside = TRUE)]
  
}

#find.all.correlations function

find.all.correlations <- function(expression_df,
                                  method = "spearman",
                                  stat = "rho",
                                  add_crumb = FALSE) {
  
  if(!require(gtools)){
    install.packages(gtools)
    library(gtools)
    message("Installing required package 'gtools'")
  }
  
  uniquepairs <- combinations(length(row.names(expression_df)), 2, row.names(expression_df))
  
  output_mat <- matrix(nrow = nrow(uniquepairs), ncol = 4)
  colnames(output_mat) <- c("Gene1", "Gene2", paste(stat), "p_value")
  
  b4 <- Sys.time()
  
  for(i in 1:nrow(uniquepairs)){
    
    x <- unlist(expression_df[uniquepairs[i, 1], ])
    y <- unlist(expression_df[uniquepairs[i, 2], ])
    
    if(isTRUE(add_crumb)){
      
      # this step prevents artifactual positive correlations caused by many 0,0 values
      
      x <- x + rnorm(x, 1)
      y <- y + rnorm(y, 1)
      
    }
    
    tempcorr <- cor.test(x, y, method = method, exact = FALSE)
    
    output_mat[i, paste(stat)]      <- tempcorr$estimate
    output_mat[i, "p_value"]  <- tempcorr$p.value
    
    # give progress
    
    if (i %% 1000 == 0) {
      
      print(paste("Combination", i, sep = " "))
      
      time_elapsed <- Sys.time() - b4
      print(paste("Time elapsed:", round(as.numeric(time_elapsed), digits = 2), attr(time_elapsed, "units"), sep = " "))
      
      timeperiteration <- time_elapsed / i
      
      remaining <- nrow(uniquepairs) - i
      print(paste("Estimated time remaining:", round((as.numeric(timeperiteration, units = "hours") * remaining), digits = 2), "hours", sep = " "))
      
    }
  }
  
  output_df <- as.data.frame(output_mat)
  output_df[, 1:2] <- uniquepairs
  
  return(output_df)
  
}

# build.matrix.4.cluster  function

build.matrix.4.cluster <- function(corr_df, stat = "rho", plotorder = NULL){
  
  corr_mat <- matrix(nrow = length(unique(c(corr_df$Gene1, corr_df$Gene2))), ncol = length(unique(c(corr_df$Gene1, corr_df$Gene2))))
  
  #This loop pulls out the correlations for each pair and sticks it in the matrix
  
  if(is.null(plotorder)) {
    
    row.names(corr_mat) <- unique(c(corr_df$Gene1, corr_df$Gene2))
    colnames(corr_mat) <- unique(c(corr_df$Gene1, corr_df$Gene2))
    
    for (i in 1:nrow(corr_df)) {
      
      genex <- corr_df[i, 1] 
      geney <- corr_df[i, 2]
      
      corr_mat[paste(genex), paste(geney)] <- corr_df[i, paste(stat)]
      corr_mat[paste(geney), paste(genex)] <- corr_df[i, paste(stat)]
      
      if (i%%1000 == 0) {
        message(paste("Combination", i, "of", nrow(corr_df), sep = " "))
      }
    }
  } else {
    
    row.names(corr_mat) <- rev(plotorder)
    colnames(corr_mat) <- plotorder
    
    for (i in 1:nrow(corr_df)) {
      
      genex <- corr_df[i, 1] 
      geney <- corr_df[i, 2]
      
      corr_mat[paste(genex), paste(geney)] <- corr_df[i, paste(stat)]
      corr_mat[paste(geney), paste(genex)] <- corr_df[i, paste(stat)]
      
      if (i%%1000 == 0) {
        message(paste("Combination", i, "of", nrow(corr_df), sep = " "))
      }
    }
  }
  
  corr_mat[is.na(corr_mat)] <-  1
  
  return(corr_mat)
  
}

# fpkm2tpm function

fpkm2tpm <- function(fpkm) {
  
  tpm_df <- as.data.frame(matrix(nrow = nrow(fpkm), ncol = ncol(fpkm)))
  row.names(tpm_df) <- row.names(fpkm)
  colnames(tpm_df) <- colnames(fpkm)
  
  for (i in 1:ncol(fpkm)){
    
    coltpm <- (fpkm[, i] / sum(fpkm[, i], na.rm = TRUE)) * 1e6
    coltpm[which(is.na(coltpm))] <- 0
    
    tpm_df[,i] <- coltpm
    
  }
  
  return(tpm_df)
  
}

#### LOAD PACKAGES & FUNCTIONS ####

## First specify the packages of interest
packages = c("CePa", 
             "stringr", 
             "matrixStats", 
             "biomaRt", 
             "ggplot2", 
             "dplyr", 
             "reshape2", 
             "qpdf", 
             "gplots", 
             "edgeR",
             "diptest",
             "enrichR",
             "openxlsx",
             "ProliferativeIndex",
             "gtools",
             "infer")

## Now load or install&load all
package.check <- lapply(
  packages,
  FUN = function(x) {
    if (!require(x, character.only = TRUE)) {
      install.packages(x, dependencies = TRUE)
      library(x, character.only = TRUE)
    }
  }
)

# Here define packges which need to be loaded through biocmanager

biocmanager_packages <- c("DESeq2",
                          "SummarizedExperiment",
                          "TCGAbiolinks",
                          "recount")

package.check <- lapply(
  biocmanager_packages,
  FUN = function(x) {
    if (!require(x, character.only = TRUE)) {
      
      if (!requireNamespace("BiocManager", quietly = TRUE)){
        install.packages("BiocManager")
      }
      
      BiocManager::install(x, dependencies = TRUE)
      
      library(x, character.only = TRUE)
      
    }
  }
)

#### LOAD INPUT DATA ####

# read in expression data from GTEX v8 downloaded directly as transcript per million (TPM) values
# file accessed through GTEX data portal: [https://gtexportal.org/home/datasets]
# file can be downloaded here: [https://storage.googleapis.com/gtex_analysis_v8/rna_seq_data/GTEx_Analysis_2017-06-05_v8_RNASeQCv1.1.9_gene_tpm.gct.gz]
TPMdata <- read.gct("input/GTEx_Analysis_2017-06-05_v8_RNASeQCv1.1.9_gene_tpm.gct")
row.names(TPMdata) <- str_remove(row.names(TPMdata), pattern = "\\..*$")

# save as RDS for rapid loading later if code is run in separate sessions
saveRDS(TPMdata, "output/TPMdata.rds")
# TPMdata <- readRDS("output/TPMdata.rds")

# read in expression data from GTEX v8 downloaded as read counts
# file accessed through GTEX data portal: [https://gtexportal.org/home/datasets]
# file can be downloaded here: [https://storage.googleapis.com/gtex_analysis_v8/rna_seq_data/GTEx_Analysis_2017-06-05_v8_RNASeQCv1.1.9_gene_reads.gct.gz]
countsdata <- read.gct("input/GTEx_Analysis_2017-06-05_v8_RNASeQCv1.1.9_gene_reads.gct")
row.names(countsdata) <- str_remove(row.names(countsdata), pattern = "\\..*$")

# save as RDS for rapid loading later if code is run in separate sessions
saveRDS(countsdata, "output/countsdata.rds")
# countsdata <- readRDS("output/countsdata.rds")

# read in countsdata from GTEX v6p
# file accessed through GTEX data portal: [https://gtexportal.org/home/datasets]
# file can be downloaded here: [https://storage.googleapis.com/gtex_analysis_v6p/rna_seq_data/GTEx_Analysis_v6p_RNA-seq_RNA-SeQCv1.1.8_gene_reads.gct.gz]
v6pcountsdata <- read.gct("input/GTEx_Analysis_v6p_RNA-seq_RNA-SeQCv1.1.8_gene_reads.gct")
row.names(v6pcountsdata) <- str_remove(row.names(v6pcountsdata), pattern = "\\..*$")

# read in rpkm data from GTEX v6p
# file accessed through GTEX data portal: [https://gtexportal.org/home/datasets]
# file can be downloaded here: [https://storage.googleapis.com/gtex_analysis_v6p/rna_seq_data/GTEx_Analysis_v6p_RNA-seq_RNA-SeQCv1.1.8_gene_rpkm.gct.gz]
v6prpkmdata <- read.gct("input/GTEx_Analysis_v6p_RNA-seq_RNA-SeQCv1.1.8_gene_rpkm.gct")
row.names(v6prpkmdata) <- str_remove(row.names(v6prpkmdata), pattern = "\\..*$")

# convert rpkm to TPM data
v6TPMdata <- fpkm2tpm(v6prpkmdata)
v6TPMdata <- as.matrix(v6TPMdata)

# read in QTL quantile normalised files
# data accessed through GTEX data portal: [https://gtexportal.org/home/datasets]
# data downloaded here: [https://storage.googleapis.com/gtex_analysis_v8/single_tissue_qtl_data/GTEx_Analysis_v8_eQTL_expression_matrices.tar]
# BED files sorted into input folder 'GTEX_bed')

QTLfilelist <- list.files("input/GTEX_bed/")

GTEXQTLdatalist <- list()

for(i in 1:length(QTLfilelist)){

  A <- as.data.frame(read.table(paste0("input/GTEX_bed/", QTLfilelist[i]), sep = "\t", stringsAsFactors = F))
  row.names(A) <- str_remove(string = A[,4], pattern = "\\.[0-9]+$")
  A <- A[, 5:ncol(A)]
  GTEXQTLdatalist[[i]] <- A
  print(i)
  names(GTEXQTLdatalist)[i] <- str_remove(QTLfilelist[i], pattern = ".v8.normalized_expression.bed")
}

# read in OXPHOS gene names
# OXPHOS gene names provided as spreadsheet Table S6
OXPHOS_names <- read.table(file = "input/OXPHOS-genes-names.txt", sep = "\t", header = TRUE)

# save mtOXPHOS and nuOXPHOS ensembl gene ids as objects for use downstream
mtOXPHOS <- OXPHOS_names[OXPHOS_names$Sourcelist == "mtOXPHOS", "ensembl_gene_id"]
nuOXPHOS <- OXPHOS_names[OXPHOS_names$Sourcelist == "nuOXPHOS", "ensembl_gene_id"]

# retrieve all mitochondrial genes (including RNA genes) from ENSEMBL biomart using biomaRt package

# set mart as ensembl human genes
ensembl <- useMart(biomart = "ensembl", dataset = "hsapiens_gene_ensembl")

# retrieve ensembl gene id and HGNC gene symbols for genes on mitochondrial DNA from biomart
mitogenes <- getBM(mart = ensembl,
                   attributes = c("ensembl_gene_id", "hgnc_symbol"),
                   filters = "chromosome_name",
                   values = "MT")

# for later, create list of nuclear genes restricted to those that are expressed in most tissues
# our criterion will be median TPM > 5 across all samples
# and genes not mitochondrially encoded or nuOXPHOS

nuclear_gene_expressed <- row.names(TPMdata[matrixStats::rowMedians(TPMdata) > 5, ])

nuclear_gene_expressed <- nuclear_gene_expressed[!(nuclear_gene_expressed %in% mitogenes$ensembl_gene_id)]

nuclear_gene_expressed <- nuclear_gene_expressed[!(nuclear_gene_expressed %in% OXPHOS_names$ensembl_gene_id)]

saveRDS(nuclear_gene_expressed, "output/GTEX_nuclear_gene_expressed.rds")
# nuclear_gene_expressed <- readRDS("output/GTEX_nuclear_gene_expressed.rds")

#### QUERY DATA WITH TCGA BIOLINKS ####

#project names downloaded from GDC Data Portal: [https://portal.gdc.cancer.gov/projects]

# change file name to match
proj_table <- read.table("input/projects-table.2020-12-03.tsv", sep="\t", header=TRUE)
proj_names <- proj_table$Project
proj_names <- proj_names[1:67]

# restrict to only TCGA projects
proj_names <- proj_names[str_detect(proj_names, pattern = "^TCGA-")]

dir.create("output/fulldata-counts")

# Query platform 
TCGA_counts_list <- list()

for (i in 1:length(proj_names)){
  
  favfile <- paste0("output/fulldata-counts/", proj_names[i], "-counts.rds")
  
  if (file.exists(favfile)) {
    
    message(paste0("We got this one (", proj_names[i], ") already"))
    next
  } else {
    
    tryCatch(
      expr = {
        query <- GDCquery(project = paste(proj_names[i]),
                          data.category = "Transcriptome Profiling",
                          data.type = "Gene Expression Quantification",
                          workflow.type = "HTSeq - Counts",
                          experimental.strategy = "RNA-Seq",
                          legacy = FALSE)
        
        GDCdownload(query, method = "api", files.per.chunk = 10)
        
        tempdata <- GDCprepare(query, summarizedExperiment = FALSE)
        
        tempdata_mat <- as.matrix(tempdata[,2:ncol(tempdata)])
        row.names(tempdata_mat) = sub("\\.[0-9]+$", "", unlist(tempdata[,1]))
        
        saveRDS(tempdata_mat, file = paste0("output/fulldata-counts/", proj_names[i] ,"-counts.rds"))
        
        TCGA_counts_list[[i]] <- tempdata_mat
        
        rm(query)
        rm(tempdata)
        rm(tempmatrix)
        rm(tempfav)
        
        gc()
        
      }, error = function(e) {
        message(paste("An error occurred for project ", proj_names[i], sep=""))
        print(e)
        
        rm(query)
        rm(tempdata)
        
        gc()
        
      }
    )
  }
}


# if count files already downloaded and saved as RDS #

# TCGA_counts_list <- list()

# TCGAcountsfilelist <- list.files(path = "output/fulldata-counts/")
#for(i in 1:length(TCGAcountsfilelist)){
  
#TCGA_counts_list[[i]] <- readRDS(paste0("output/fulldata-counts/", TCGAcountsfilelist[i]))
#}

names(TCGA_counts_list) <- str_remove(TCGAcountsfilelist, pattern = "-counts.rds")

saveRDS(TCGA_counts_list, "output/TCGA_counts_list.rds")
#  TCGA_counts_list <-readRDS("output/TCGA_counts_list.rds")

# download TCGA RPKM data
dir.create("output/fulldata-FPKM")

# Query platform 
TCGA_FPKM_list <- list()

for (i in 1:length(proj_names)){
  
  favfile <- paste0("output/fulldata-FPKM/", proj_names[i], "-FPKM.rds")
  
  if (file.exists(favfile)) {
    
    message(paste0("We got this one (", proj_names[i], ") already"))
    next
  } else {
    
    tryCatch(
      expr = {
        query <- GDCquery(project = paste(proj_names[i]),
                          data.category = "Transcriptome Profiling",
                          data.type = "Gene Expression Quantification",
                          workflow.type = "HTSeq - FPKM",
                          experimental.strategy = "RNA-Seq",
                          legacy = FALSE)
        
        GDCdownload(query, method = "api", files.per.chunk = 10)
        
        tempdata <- GDCprepare(query, summarizedExperiment = FALSE)
        
        tempdata_mat <- as.matrix(tempdata[,2:ncol(tempdata)])
        row.names(tempdata_mat) = sub("\\.[0-9]+$", "", unlist(tempdata[,1]))
        
        saveRDS(tempdata_mat, file = paste0("output/fulldata-FPKM/", proj_names[i] ,"-FPKM.rds"))
        
        TCGA_FPKM_list[[i]] <- tempdata_mat
        
        rm(query)
        rm(tempdata)
        rm(tempmatrix)
        rm(tempfav)
        
        gc()
        
      }, error = function(e) {
        message(paste("An error occurred for project ", proj_names[i], sep=""))
        print(e)
        
        rm(query)
        rm(tempdata)
        
        gc()
        
      }
    )
  }
}

## if FPKM already downloaded ##

# TCGA_FPKM_list <- list()

# TCGAFPKMfilelist <- list.files(path = "output/fulldata-FPKM/")

#for(i in 1:length(TCGAFPKMfilelist)){
  
#TCGA_FPKM_list[[i]] <- as.data.frame(readRDS(paste0("output/fulldata-FPKM/", TCGAFPKMfilelist[i])))
#}

names(TCGA_FPKM_list) <- str_remove(str_replace_all(TCGAFPKMfilelist, pattern = "-", replacement = "_"), pattern = ".rds")

# remove ENSEMBL gene transcript ID version
TCGA_FPKM_list <- lapply(TCGA_FPKM_list, function(x){
  row.names(x) <- str_remove(x$X1, pattern = "\\..*$")
  x <- x[, 2:ncol(x)]
})

saveRDS(TCGA_FPKM_list, "output/TCGA_FPKM_list.rds")
# TCGA_FPKM_list <- readRDS("output/TCGA_FPKM_list.rds")

# convert TCGA FPKM data to TPM

TCGA_TPM_list <- list()

TCGAFPKMfilelist <- list.files(path = "output/fulldata-FPKM/")

TCGA_TPM_data <- merge(fpkm2tpm(TCGA_FPKM_list[[1]]), fpkm2tpm(TCGA_FPKM_list[[2]]), by = 0)
row.names(TCGA_TPM_data) <- TCGA_TPM_data$Row.names
TCGA_TPM_data <- TCGA_TPM_data[, 2:ncol(TCGA_TPM_data)]

TCGA_TPM_list[[1]] <- fpkm2tpm(TCGA_FPKM_list[[1]])
TCGA_TPM_list[[2]] <- fpkm2tpm(TCGA_FPKM_list[[2]])

for(i in 3:length(TCGA_FPKM_list)){
  
  TPMconvert <- fpkm2tpm(TCGA_FPKM_list[[i]])
  TCGA_TPM_data <- merge(TCGA_TPM_data, TPMconvert, by = 0)
  row.names(TCGA_TPM_data) <- TCGA_TPM_data$Row.names
  TCGA_TPM_data <- TCGA_TPM_data[, 2:ncol(TCGA_TPM_data)]
  
  TCGA_TPM_list[[i]] <- TPMconvert
  
}

names(TCGA_TPM_list) <- names(TCGA_FPKM_list)

saveRDS(TCGA_TPM_list, "output/TCGA_TPM_list.rds")
saveRDS(TCGA_TPM_data, "output/TCGA_TPM_data.rds")
# TCGA_TPM_data <- readRDS("output/TCGA_TPM_data.rds")

#### GTEX LOOKUP TABLES ####

# create lookup table for GTEX v8
# load sample annotation and phenotype files and make lookup table for samples and tissues for later use

# file accessed through GTEX data portal: [https://gtexportal.org/home/datasets]
# file can be downloaded here: [https://storage.googleapis.com/gtex_analysis_v8/annotations/GTEx_Analysis_v8_Annotations_SampleAttributesDS.txt]
sampleattributes <- read.table("input/GTEx_Analysis_v8_Annotations_SampleAttributesDS.txt", sep = "\t", header = TRUE, quote = "")
row.names(sampleattributes) <- sampleattributes$SAMPID

# file accessed through GTEX data portal: [https://gtexportal.org/home/datasets]
# file can be downloaded here: [https://storage.googleapis.com/gtex_analysis_v8/annotations/GTEx_Analysis_v8_Annotations_SubjectPhenotypesDS.txt]
subjectphenotypes <- read.table("input/GTEx_Analysis_v8_Annotations_SubjectPhenotypesDS.txt", sep = "\t", header = TRUE, quote = "")
row.names(subjectphenotypes) <- subjectphenotypes$SUBJID

# Create new data frame in which to build our lookup table for the samples present in the expression data table
LUT <- as.data.frame(matrix(nrow = ncol(TPMdata), ncol = 2))
colnames(LUT) <- c("SUBJID", "SAMPID")

# for the purposes of constructing the lookup table, will use column names from the TPM data
# this code recreates the SUBJID from the column/sample names in the RPKMtable
# first splitting by the separator (in this case, ".")
sampid_split_list <- str_split(colnames(TPMdata), pattern = "\\.")

# then pasting back the first two fields with a "-" separator to get the subject ID, as SUBJID appears in sample annotation file.
SUBJID_vec <- c()

for(i in 1:length(sampid_split_list)){
  SUBJID_vec[i] <- paste(sampid_split_list[[i]][1:2], collapse = "-")  
}

# fill SUBJID column with SUBJIDs recreated from RPKM file
LUT$SUBJID <- SUBJID_vec

# fill SAMPID column with sample IDs from RPKM file
LUT$SAMPID <- colnames(TPMdata)

# add age bracket as column to look up table
LUT[, "age_bracket"] <- subjectphenotypes[LUT$SUBJID, "AGE"]

# set row names as sample IDs to allow for easy subsetting later
rownames(LUT) <- LUT$SAMPID

# add tissue as column to look up table (in sample attributes SMTSD column)
# note str_replace command is needed here to match the sampleattributes file as SAMPID from TPM table has "." separator, not "-"
LUT[, "Tissue"] <- sampleattributes[str_replace_all(LUT$SAMPID, pattern = "\\.", replacement = "-"), "SMTSD"]

# add gender as column to look up table 
LUT[, "Sex"] <- subjectphenotypes[LUT$SUBJID, "SEX"]

# replace gender codes (0,1) with appropriate strings
LUT[, "Sex"] <- str_replace_all(as.character(LUT$Sex), pattern = "1", replacement = "MALE")
LUT[, "Sex"] <- str_replace_all(as.character(LUT$Sex), pattern = "2", replacement = "FEMALE")

# add Hardy deathscale ('cause of death') to lookup table
LUT[, "DeathScale"] <- subjectphenotypes[LUT$SUBJID, "DTHHRDY" ]

# add sample ischemic time to lookup table
LUT[, "IschemicTime"] <- sampleattributes[str_replace_all(LUT$SAMPID, pattern = "\\.", replacement = "-"), "SMTSISCH"]

# add sample sequencing batch to lookup table
LUT[, "Batch"] <- sampleattributes[str_replace_all(LUT$SAMPID, pattern = "\\.", replacement = "-"), "SMGEBTCH"]

# save the lookup table in the output folder
saveRDS(LUT, file = "output/GTEX-version8-sampleID-LUT.rds")
# LUT <- readRDS("output/GTEX-version8-sampleID-LUT.rds")

# Make vector of tissue names to use downstream
tissues <- unique(LUT$Tissue)

# Here we will restrict all analysis to tissues with at least 100 samples

# This results in the exclusion of samples from 6 tissues:
# Bladder
# Cervix - Ectocervix
# Cervix - Endocervix
# Fallopian Tube
# Kidney - Cortex
# Kidney - Medulla

# 48 of 54 tissues remain in the analysis
tissues <- names(table(LUT$Tissue))[table(LUT$Tissue) > 100]
saveRDS(tissues, file = "output/GTEX-tissues.rds")

# make lookup table for V6
# load sample annotation and phenotype files for version 6 and make lookup table

# file accessed through GTEX data portal: [https://gtexportal.org/home/datasets]
# file can be downloaded here: [https://storage.googleapis.com/gtex_analysis_v6p/annotations/GTEx_Data_V6_Annotations_SampleAttributesDS.txt]
v6sampleattributes <- read.table("input/GTEx_Data_V6_Annotations_SampleAttributesDS.txt", sep = "\t", header = TRUE, quote = "")
row.names(v6sampleattributes) <- v6sampleattributes$SAMPID

# file accessed through GTEX data portal: [https://gtexportal.org/home/datasets]
# file can be downloaded here: [https://storage.googleapis.com/gtex_analysis_v6p/annotations/GTEx_Data_V6_Annotations_SubjectPhenotypesDS.txt]
v6subjectphenotypes <- read.table("input/GTEx_Data_V6_Annotations_SubjectPhenotypesDS.txt", sep = "\t", header = TRUE, quote = "")
row.names(v6subjectphenotypes) <- v6subjectphenotypes$SUBJID

# this code recreates the SUBJID from the column/sample names in the RPKMtable
# first splitting by the separator (in this case, ".")
sampid_split_list <- str_split(colnames(v6prpkmdata), pattern = "\\.")

# then pasting back the first two fields with a "-" separator, as SUBJID appears in sample annotation file.
SUBJID_vec <- c()

for(i in 1:length(sampid_split_list)){
  SUBJID_vec[i] <- paste(sampid_split_list[[i]][1:2], collapse = "-")  
}

# Now create new data frame to construct our lookup table for the samples present in the RPKM table
v6LUT <- as.data.frame(matrix(nrow = ncol(v6prpkmdata), ncol = 2))
colnames(v6LUT) <- c("SUBJID", "SAMPID")

# fill SUBJID column with SUBJIDs recreated from RPKM file
v6LUT$SUBJID <- SUBJID_vec

# fill SAMPID column with sample IDs from RPKM file
v6LUT$SAMPID <- colnames(v6prpkmdata)

# add age bracket as column to look up table
v6LUT[, "age_bracket"] <- v6subjectphenotypes[v6LUT$SUBJID, "AGE"]

# set row names as sample IDs to allow for easy subsetting later
rownames(v6LUT) <- v6LUT$SAMPID

# add tissue as column to look up table
# note str_replace command is needed to match the v6sampleattributes file as SAMPID from RPKM table has "." separator, not "-"
v6LUT[, "Tissue"] <- v6sampleattributes[str_replace_all(v6LUT$SAMPID, pattern = "\\.", replacement = "-"), "SMTSD"]

# add gender as column to look up table 
v6LUT[, "Sex"] <- v6subjectphenotypes[v6LUT$SUBJID, "GENDER"]

# replace gender codes with strings to use as factors in linear regression
v6LUT[, "Sex"] <- str_replace_all(as.character(v6LUT$Sex), pattern = "1", replacement = "MALE")
v6LUT[, "Sex"] <- str_replace_all(as.character(v6LUT$Sex), pattern = "2", replacement = "FEMALE")

# add Hardy deathscale ('cause of death') to lookup table
v6LUT[, "DeathScale"] <- v6subjectphenotypes[v6LUT$SUBJID, "DTHHRDY" ]

# add sample ischemic time to lookup table
v6LUT[, "IschemicTime"] <- v6sampleattributes[str_replace_all(v6LUT$SAMPID, pattern = "\\.", replacement = "-"), "SMTSISCH"]

# add sample sequencing batch to lookup table
v6LUT[, "Batch"] <- v6sampleattributes[str_replace_all(v6LUT$SAMPID, pattern = "\\.", replacement = "-"), "SMGEBTCH"]

# the lookup table is now completed.
v6tissues <- names(table(v6LUT$Tissue))[table(v6LUT$Tissue) > 100]

saveRDS(tissues, file = "output/GTEX_V6_LUT.rds")

#### TCGA LOOKUP TABLES ####

# download TCGA metadata using 'recount' package
TCGA_metadata <- all_metadata(subset = "tcga", verbose = TRUE) 

# compile fields of interest into data frame
metadata_for_lm <- data.frame(cbind(TCGA_metadata$gdc_cases.samples.portions.analytes.aliquots.submitter_id,
                                    TCGA_metadata$gdc_cases.project.project_id,
                                    TCGA_metadata$gdc_cases.samples.sample_type,
                                    TCGA_metadata$gdc_cases.demographic.race,
                                    TCGA_metadata$gdc_cases.demographic.gender,
                                    TCGA_metadata$gdc_cases.diagnoses.tumor_stage,
                                    TCGA_metadata$gdc_cases.diagnoses.days_to_birth
))

# name columns
colnames(metadata_for_lm) <- c("SAMPID",
                               "cancer",
                               "sample_type",
                               "race",
                               "gender",
                               "tumour_stage",
                               "days_to_birth"
)

# Add field from barcode for sequencing centre
metadata_for_lm[, "sequencing_centre"] <- as.factor(str_extract(string = metadata_for_lm$SAMPID, pattern = "[:digit:]{2}$"))

# simplify tumour stage classification into stage 0, 1, 2, 3 or 4 (or not reported)
metadata_for_lm$tumour_stage <- str_replace_all(metadata_for_lm$tumour_stage, pattern = "stage i$", replacement = "stage_1")
metadata_for_lm$tumour_stage <- str_replace_all(metadata_for_lm$tumour_stage, pattern = "stage ia$", replacement = "stage_1")
metadata_for_lm$tumour_stage <- str_replace_all(metadata_for_lm$tumour_stage, pattern = "stage ib$", replacement = "stage_1")

metadata_for_lm$tumour_stage <- str_replace_all(metadata_for_lm$tumour_stage, pattern = "stage ii$", replacement = "stage_2")
metadata_for_lm$tumour_stage <- str_replace_all(metadata_for_lm$tumour_stage, pattern = "stage iia$", replacement = "stage_2")
metadata_for_lm$tumour_stage <- str_replace_all(metadata_for_lm$tumour_stage, pattern = "stage iib$", replacement = "stage_2")
metadata_for_lm$tumour_stage <- str_replace_all(metadata_for_lm$tumour_stage, pattern = "stage iic$", replacement = "stage_2")

metadata_for_lm$tumour_stage <- str_replace_all(metadata_for_lm$tumour_stage, pattern = "stage iii$", replacement = "stage_3")
metadata_for_lm$tumour_stage <- str_replace_all(metadata_for_lm$tumour_stage, pattern = "stage iiia$", replacement = "stage_3")
metadata_for_lm$tumour_stage <- str_replace_all(metadata_for_lm$tumour_stage, pattern = "stage iiib$", replacement = "stage_3")
metadata_for_lm$tumour_stage <- str_replace_all(metadata_for_lm$tumour_stage, pattern = "stage iiic$", replacement = "stage_3")

metadata_for_lm$tumour_stage <- str_replace_all(metadata_for_lm$tumour_stage, pattern = "stage iv$", replacement = "stage_4")
metadata_for_lm$tumour_stage <- str_replace_all(metadata_for_lm$tumour_stage, pattern = "stage iva$", replacement = "stage_4")
metadata_for_lm$tumour_stage <- str_replace_all(metadata_for_lm$tumour_stage, pattern = "stage ivb$", replacement = "stage_4")
metadata_for_lm$tumour_stage <- str_replace_all(metadata_for_lm$tumour_stage, pattern = "stage ivc$", replacement = "stage_4")

#Keep only primary tumour samples; exclude stages i/ii nos, is, stage 0 and stage x; latter excludes only 84 samples across all TCGA
TCGA_LUT <- metadata_for_lm[metadata_for_lm$sample_type %in% c("Primary Tumor","Primary Blood Derived Cancer - Peripheral Blood"), ]
TCGA_LUT <- TCGA_LUT[!(TCGA_LUT$tumour_stage %in% c("i/ii nos", "is", "stage x")), ]

# remove duplicate entries (duplicated sample ID)
TCGA_LUT <- TCGA_LUT[!(duplicated(TCGA_LUT$SAMPID)),]

# remove entries where multiple samples are derived from one donor. keep only donors that provide one cancer sample.
TGA_LUT <- TCGA_LUT[(!duplicated(str_extract(string = TCGA_LUT$SAMPID, pattern = "^TCGA-[:alnum:]{2}-[:alnum:]{4}")) & !duplicated(str_extract(string = TCGA_LUT$SAMPID, pattern = "^TCGA-[:alnum:]{2}-[:alnum:]{4}"), fromLast = TRUE)), ]

# set SAMPID as row names
row.names(TCGA_LUT) <- TCGA_LUT$SAMPID

saveRDS(TCGA_LUT, "output/TCGA_LUT.rds")
# TCGA_LUT <- readRDS("output/TCGA_LUT.rds")

cancertypes <- str_sort(unique(TCGA_LUT$cancer))
saveRDS(cancertypes, "output/TCGA_cancertypes.rds")

#### PERFORM NORMALISATIONS ####

# perform MRN normalisation by tissue
# NB in this script the MRN normalisation is referred to as 'MOR' (median of ratios) instead of MRN (median ratio normalisation) as in the accompanying manuscript
# MRN/MOR normalisation uses 'DESeq2' package

MOR_tissue_list <- list()

for (i in 1:length(tissues)){
  
  # retrieve sample IDs for this tissue from the lookup table
  sampleIDs <- LUT[LUT$Tissue == tissues[i], "SAMPID"]
  
  # subset the dataframe of raw counts by sample ids corresponding to tissue i
  tissuecounts <- countsdata[, colnames(countsdata) %in% sampleIDs]

  # DESeq2 wants a colData object. Not actually used for the normalisation. Here we can use the sample IDs with tissue.
  col_tissue <- LUT[colnames(tissuecounts), "Tissue"]
  col_tissue <- as.matrix(col_tissue)
  
  rownames(col_tissue) <- colnames(tissuecounts)
  colnames(col_tissue) <- c("Tissue")
  
  # need to create a DESeq2 object. Design set to ~1 allows for use of estimateSizeFactors
  tempdds <- DESeqDataSetFromMatrix(countData = tissuecounts, colData = col_tissue, design = ~ 1)
  
  # this function estimates the scaling factors from the samples from the median of ratios wrt to the geometric mean for each gene across samples
  tempdds <- estimateSizeFactors(tempdds)
  
  # put the counts normalised by the scaling factors in a new object
  tissuenormalised_counts <- counts(tempdds, normalized = TRUE)
  
  # put object in list
  MOR_tissue_list[[i]] <- tissuenormalised_counts
  
}

# set the names in the list to be the tissue names
names(MOR_tissue_list) <- tissues

# save the tissue MOR-normalised values in the output folder
saveRDS(MOR_tissue_list, "output/MOR-normalisation-by-tissue-list.rds")
# MOR_tissue_list <- readRDS("output/MOR-normalisation-by-tissue-list.rds")

# perform MOR normalisation by tissue for GTEX v6p

v6MOR_tissue_list <- list()

for (i in 1:length(v6tissues)){
  
  # retrieve sample IDs for this tissue from the lookup table
  sampleIDs <- v6LUT[v6LUT$Tissue == v6tissues[i], "SAMPID"]
  
  # subset the dataframe of raw counts by sample ids corresponding to tissue i
  tissuecounts <- v6pcountsdata[, colnames(v6pcountsdata) %in% sampleIDs]
  
  # DESeq2 wants a colData object. Not actually used for the normalisation. Here we can use the sample IDs with tissue.
  col_tissue <- v6LUT[colnames(tissuecounts), "Tissue"]
  col_tissue <- as.matrix(col_tissue)
  
  rownames(col_tissue) <- colnames(tissuecounts)
  colnames(col_tissue) <- c("Tissue")
  
  # need to create a DESeq2 object. Design set to ~1 allows for use of estimateSizeFactors
  tempdds <- DESeqDataSetFromMatrix(countData = tissuecounts, colData = col_tissue, design = ~ 1)
  
  # this function estimates the scaling factors from the samples from the median of ratios wrt to the geometric mean for each gene across samples
  tempdds <- estimateSizeFactors(tempdds)
  
  # put the counts normalised by the scaling factors in a new object
  tissuenormalised_counts <- counts(tempdds, normalized = TRUE)
  
  # put object in list
  v6MOR_tissue_list[[i]] <- tissuenormalised_counts
  
}

# set the names in the list to be the tissue names
names(v6MOR_tissue_list) <- v6tissues

# perform MOR normalisation across all tissues combined, rather than individually (we will use this later for correlating to NFKB/immune cell fraction/Proliferative Index)

# DESeq2 wants a colData object. Not actually used for the normalisation. Here we can use the sample IDs with tissue.
col_data <- LUT[colnames(countsdata), "Tissue"]
col_data <- as.matrix(col_data)

rownames(col_data) <- colnames(countsdata)
colnames(col_data) <- c("Tissue")

# Normalise reads by scaling factors using DESeq2

# need to create a DESeq2 object. Design set to ~1 allows for use of estimateSizeFactors
dds <- DESeqDataSetFromMatrix(countData = countsdata, colData = col_data, design = ~ 1)

# this function estimates the scaling factors from the samples from the median of ratios wrt to the geometric mean for each gene across samples
dds <- estimateSizeFactors(dds)

# save the counts normalised by the scaling factors in a new object
normalised_counts <- counts(dds, normalized = TRUE)

saveRDS(normalised_counts, "output/MOR-normalisation-across-tissues.rds")

# perform TMM normalisation by tissue
# TMM normalisation uses 'edgeR' package

TMM_tissue_list <- list()

for (i in 1:length(tissues)){

  # retrieve sample IDs for this tissue from the lookup table
  sampleIDs <-  LUT[LUT$Tissue == tissues[i], "SAMPID"]

  # subset the dataframe of raw counts by sample ids corresponding to tissue i
  tissuecounts <- countsdata[, sampleIDs]

  # the edgeR package wants to convert the counts to a DGE object
  temp_dge <- DGEList(tissuecounts)
  
  # to calculate effective library size for normalisation, we use the library size multiplied by the normalisation factor
  # these values correspond closely to the DESeq SizeFactors
  calc_temp <- calcNormFactors(temp_dge,
                               method = "TMM")
  
  effectivelibrarysize_temp <- calc_temp$samples$lib.size * calc_temp$samples$norm.factors

  # to normalise, divide counts by effectivelibrarysize
  # as we want to divide across the samples, the matrix is first transposed before division, and then again afterwards
  TMMtissuecounts <- t(t(tissuecounts) / effectivelibrarysize_temp) * 10e6

  # put object in list
  TMM_tissue_list[[i]] <- TMMtissuecounts
  
}

# set names for tissue matrices in list
names(TMM_tissue_list) <- tissues

# save the list in output folder
saveRDS(TMM_tissue_list, "output/TMM-normalisation-by-tissue-list.rds")
# TMM_tissue_list <- readRDS("output/TMM-normalisation-by-tissue-list.rds")

# perform UQ normalisation by tissue
# UQ normalisation performed with 'edgeR' package

UQ_tissue_list <- list()

for (i in 1:length(tissues)){
  
  # retrieve sample IDs for this tissue from the lookup table
  sampleIDs <-  LUT[LUT$Tissue == tissues[i], "SAMPID"]
  
  # subset the dataframe of raw counts by sample ids corresponding to tissue i
  tissuecounts <- countsdata[, sampleIDs]
  
  # the edgeR package wants to convert the counts to a DGE object
  temp_dge <- DGEList(tissuecounts)
  
  # to calculate effective library size for normalisation, we use the library size multiplied by the normalisation factor
  # these values correspond closely to the DESeq SizeFactors
  calc_temp <- calcNormFactors(temp_dge,
                               method = "upperquartile")
  
  effectivelibrarysize_temp <- calc_temp$samples$lib.size * calc_temp$samples$norm.factors
  
  # to normalise, divide counts by effectivelibrarysize
  # as we want to divide across the samples, the matrix is first transposed before division, and then again afterwards
  UQtissuecounts <- t(t(tissuecounts) / effectivelibrarysize_temp) * 10e6
  
  # put object in list
  UQ_tissue_list[[i]] <- UQtissuecounts
  
}

# set names for tissue matrices in list
names(UQ_tissue_list) <- tissues

# save the list in output folder
saveRDS(UQ_tissue_list, "output/UQ-normalisation-by-tissue-list.rds")

# perform MOR normalisation by cancer type for TCGA

TCGA_MOR_list <- list()

for (i in 1:length(cancertypes)){

  # retrieve sample IDs for this tissue from the lookup table. use metadata_for_lm to not restrict to cancer samples; include normal samples.
  sampleIDs <- metadata_for_lm[metadata_for_lm$cancer == cancertypes[i], "SAMPID"]
  
  # subset the dataframe of raw counts by sample ids corresponding to tissue i
  cancercounts <- TCGA_counts_list[[paste(cancertypes[i])]][, colnames(TCGA_counts_list[[paste(cancertypes[i])]]) %in% sampleIDs]
  
  # DESeq2 wants a colData object. Not actually used for the normalisation. Here we can use the sample IDs with tissue.
  col_cancer <- metadata_for_lm[colnames(cancercounts), "cancer"]
  col_cancer <- as.matrix(col_cancer)
  
  rownames(col_cancer) <- colnames(cancercounts)
  colnames(col_cancer) <- c("cancer")
  
  # need to create a DESeq2 object. Design set to ~1 allows for use of estimateSizeFactors
  tempdds <- DESeqDataSetFromMatrix(countData = cancercounts, colData = col_cancer, design = ~ 1)
  
  # this function estimates the scaling factors from the samples from the median of ratios wrt to the geometric mean for each gene across samples
  tempdds <- estimateSizeFactors(tempdds)
  
  # put the counts normalised by the scaling factors in a new object
  cancernormalised_counts <- counts(tempdds, normalized = TRUE)
  
  # put object in list
  TCGA_MOR_list[[i]] <- cancernormalised_counts
  
}

# set the names in the list to be the tissue names
names(TCGA_MOR_list) <- cancertypes

# save the list in output folder
saveRDS(TCGA_MOR_list, "output/TCGA_MOR_list.rds")
# TCGA_MOR_list <- readRDS("output/TCGA_MOR_list.rds")

# perform MOR normalisation across all cancers (for later correlation with NFKB/Proliferative Index)

# DESeq2 wants a colData object. Not actually used for the normalisation. Here we can use the sample IDs with tissue.
col_data <- TCGA_LUT[unlist(sapply(TCGA_counts_list, colnames)), "cancer"]

col_data <- as.matrix(col_data)

row.names(col_data) <- unlist(sapply(TCGA_counts_list, colnames))
colnames(col_data) <- c("cancer")

TCGA_counts_df <- do.call(cbind, TCGA_counts_list)

col_data <- col_data[colnames(TCGA_counts_df), ]

col_data <- col_data[!is.na(col_data)]
col_data <- data.frame(col_data)
TCGA_counts_df <- TCGA_counts_df[, colnames(TCGA_counts_df) %in% row.names(col_data)]

# Normalise reads by scaling factors using DESeq2

# need to create a DESeq2 object. Design set to ~1 allows for use of estimateSizeFactors
dds <- DESeqDataSetFromMatrix(countData = TCGA_counts_df, colData = col_data, design = ~ 1)

# this function estimates the scaling factors from the samples from the median of ratios wrt to the geometric mean for each gene across samples
dds <- estimateSizeFactors(dds)

# save the counts normalised by the scaling factors in a new object
normalised_counts <- counts(dds, normalized = TRUE)

# save it. this is for all genes. note this normalisation across all samples from all tissues
saveRDS(normalised_counts, "output/TCGA-MOR-normalisation-across-cancers.rds")

# perform TMM normalisation by cancertype

TCGA_TMM_list <- list()

for (i in 1:length(cancertypes)){
  
  # retrieve sample IDs for this tissue from the lookup table
  sampleIDs <- metadata_for_lm[metadata_for_lm$cancer == cancertypes[i], "SAMPID"]
  
  # subset the dataframe of raw counts by sample ids corresponding to tissue i
  cancercounts <- TCGA_counts_list[[paste(cancertypes[i])]][, colnames(TCGA_counts_list[[paste(cancertypes[i])]]) %in% sampleIDs]
  
  # the edgeR package wants to convert the counts to a DGE object
  temp_dge <- DGEList(cancercounts)
  
  # to calculate effective library size for normalisation, we use the library size multiplied by the normalisation factor
  # these values correspond closely to the DESeq SizeFactors
  calc_temp <- calcNormFactors(temp_dge,
                               method = "TMM")
  
  effectivelibrarysize_temp <- calc_temp$samples$lib.size * calc_temp$samples$norm.factors
  
  # to normalise, divide counts by effectivelibrarysize
  # as we want to divide across the samples, the matrix is first transposed before division, and then again afterwards
  TMMcancercounts <- t(t(cancercounts) / effectivelibrarysize_temp) * 10e6
  
  # put object in list
  TCGA_TMM_list[[i]] <- TMMcancercounts
  
}

# set names for tissue matrices in list
names(TCGA_TMM_list) <- cancertypes

# save the list in output folder
saveRDS(TCGA_TMM_list, "output/TCGA_TMM_list.rds")

# perform UQ normalsation by cancertype

TCGA_UQ_list <- list()

for (i in 1:length(cancertypes)){
  
  # retrieve sample IDs for this tissue from the lookup table
  sampleIDs <- metadata_for_lm[metadata_for_lm$cancer == cancertypes[i], "SAMPID"]
  
  # subset the dataframe of raw counts by sample ids corresponding to tissue i
  cancercounts <- TCGA_counts_list[[paste(cancertypes[i])]][, colnames(TCGA_counts_list[[paste(cancertypes[i])]]) %in% sampleIDs]
  
  # the edgeR package wants to convert the counts to a DGE object
  temp_dge <- DGEList(cancercounts)
  
  # to calculate effective library size for normalisation, we use the library size multiplied by the normalisation factor
  # these values correspond closely to the DESeq SizeFactors
  calc_temp <- calcNormFactors(temp_dge,
                               method = "upperquartile")
  
  effectivelibrarysize_temp <- calc_temp$samples$lib.size * calc_temp$samples$norm.factors
  
  # to normalise, divide counts by effectivelibrarysize
  # as we want to divide across the samples, the matrix is first transposed before division, and then again afterwards
  UQcancercounts <- t(t(cancercounts) / effectivelibrarysize_temp) * 10e6
  
  # put object in list
  TCGA_UQ_list[[i]] <- UQcancercounts
  
}

# set names for tissue matrices in list
names(TCGA_UQ_list) <- cancertypes

# save the list in output folder
saveRDS(TCGA_UQ_list, "output/TCGA_UQ_list.rds")

#### VIOLIN PLOT OF MITOCHONDRIAL EXPRESSION ACROSS TISSUES ####
# this corresponds to Fig 1b, Fig S1a

mito_expr_long <- data.frame(matrix(nrow = 0, ncol = 2))
colnames(mito_expr_long) <- c("mito_total", "tissue")

for(i in 1:length(tissues)){
  
  sampleIDs <- LUT[LUT$Tissue == tissues[i], "SAMPID"]
  
  tempdata <- TPMdata[mitogenes$ensembl_gene_id, sampleIDs]
  
  mitosums <- colSums(tempdata)
  
  temp_df <- data.frame(matrix(nrow = length(mitosums), ncol = 2))
  colnames(temp_df) <- c("mito_total", "tissue")
  
  temp_df[, "mito_total"] <- mitosums
  temp_df[, "tissue"] <- tissues[i]
  
  mito_expr_long <- rbind(mito_expr_long, temp_df)
  
}

mito_expr_long$mito_total <- mito_expr_long$mito_total / 1e4

# Here replace tissue names merely to shorten long names for plotting
tissue_select_vec <- c("Adipose \\- Subcutaneous",
                       "Adipose \\- Visceral \\(Omentum\\)",
                       "Adrenal Gland",
                       "Artery \\- Aorta",
                       "Artery \\- Coronary",
                       "Artery  \\- Tibial",
                       "Brain \\- Amygdala",
                       "Brain \\- Anterior cingulate cortex \\(BA24\\)",
                       "Brain \\- Caudate \\(basal ganglia\\)",
                       "Brain \\- Cerebellar Hemisphere",
                       "Brain \\- Cerebellum",
                       "Brain \\- Cortex",
                       "Brain \\- Frontal Cortex \\(BA9\\)",
                       "Brain \\- Hippocampus",
                       "Brain \\- Hypothalamus",
                       "Brain \\- Nucleus accumbens \\(basal ganglia\\)",
                       "Brain \\- Putamen \\(basal ganglia\\)",
                       "Brain \\- Spinal cord \\(cervical c\\-1\\)",
                       "Brain \\- Substantia nigra",
                       "Breast \\- Mammary Tissue",
                       "Cells \\- Cultured fibroblasts",
                       "Cells \\- EBV-transformed lymphocytes",
                       "Colon \\- Sigmoid",
                       "Colon \\- Transverse",
                       "Esophagus \\- Gastroesophageal Junction",
                       "Esophagus \\- Mucosa",
                       "Esophagus \\- Muscularis",
                       "Heart \\- Atrial Appendage",
                       "Heart \\- Left Ventricle",
                       "Liver",
                       "Lung",
                       "Minor Salivary Gland",
                       "Muscle \\- Skeletal",
                       "Nerve \\- Tibial",
                       "Ovary",
                       "Pancreas",
                       "Pituitary",
                       "Prostate",
                       "Skin \\- Not Sun Exposed \\(Suprapubic\\)",
                       "Skin \\- Sun Exposed \\(Lower leg\\)",
                       "Small Intestine \\- Terminal Ileum",
                       "Spleen",
                       "Stomach",
                       "Testis",
                       "Thyroid",
                       "Uterus",
                       "Vagina",
                       "Whole Blood")

tissue_replace_vec <- c("Adipose (Subcut.)",
                        "Adipose (Visc.)",
                        "Adrenal Gland",
                        "Artery (Aorta)",
                        "Artery (Coronary)",
                        "Artery (Tibial)",
                        "Brain (Amygdala)",
                        "Brain (Ant.cing. cortex)",
                        "Brain (Caudate)",
                        "Brain (Cereb. Hemsph.)",
                        "Brain (Cerebellum)",
                        "Brain (Cortex)",
                        "Brain (Frontal Cortex)",
                        "Brain (Hippocampus)",
                        "Brain (Hypothalamus)",
                        "Brain (Nucl. acc.)",
                        "Brain (Putamen)",
                        "Brain (Spinal cord)",
                        "Brain (Subst. nigra)",
                        "Breast (Mammary)",
                        "Cultured fibroblasts",
                        "EBV-transf. lymphocytes",
                        "Colon (Sigmoid)",
                        "Colon (Transverse)",
                        "Esophagus (Gastr. Junc.)",
                        "Esophagus (Mucosa)",
                        "Esophagus (Muscularis)",
                        "Heart (Atrial Appendage)",
                        "Heart (Left Ventricle)",
                        "Liver",
                        "Lung",
                        "Min. Saliv. Gland",
                        "Muscle (Skeletal)",
                        "Nerve (Tibial)",
                        "Ovary",
                        "Pancreas",
                        "Pituitary",
                        "Prostate",
                        "Skin (Suprapubic)",
                        "Skin (Lower leg)",
                        "Small Intestine",
                        "Spleen",
                        "Stomach",
                        "Testis",
                        "Thyroid",
                        "Uterus",
                        "Vagina",
                        "Whole Blood")

for (i in 1:length(tissue_select_vec)){
  mito_expr_long$tissue <- str_replace_all(mito_expr_long$tissue, 
                                           pattern = tissue_select_vec[i], 
                                           replacement = tissue_replace_vec[i])
}

mito_expr_long_unchanged <- mito_expr_long

mito_average <- aggregate(mito_total ~ tissue, 
                          mito_expr_long, 
                          mean)

tissues_mito_order <- mito_average[order(mito_average$mito_total, decreasing = FALSE), "tissue"]

mito_expr_long$tissue <- factor(mito_expr_long$tissue, levels = tissues_mito_order)

# plot for all of the tissues (Fig S1b)

pdf("graphics/mt-expression-violin-all-tissues.pdf", width = 8, height = 3)

ggplot(data = mito_expr_long, aes(x = tissue, 
                                  y = mito_total, 
                                  fill = tissue)) + 
  geom_violin(show.legend = FALSE, 
              scale = 1, 
              position = position_dodge(1), 
              lwd = 0.2) + 
  ylab("% mitochondrial transcripts") +
  xlab("GTEX Tissue") +
  theme_classic() +
        theme(axis.text.x = element_text(angle =30, 
                                         vjust = 1, 
                                         hjust=1, 
                                         family = "sans", 
                                         color = "black", 
                                         size = 5),
              axis.text.y = element_blank(),
              axis.title.x = element_blank(),
              axis.title.y = element_blank(),
              axis.line = element_line(size = 0.25),
              axis.ticks = element_line(size = 0.25),
              plot.margin = unit(c(0, 0, 0, 1.5), "cm"))

dev.off()

# for a selection (Fig 1a)

pdf("graphics/mt-expression-violin-selection-of-tissues.pdf", width = 5, height = 3)

ggplot(data = mito_expr_long[mito_expr_long$tissue %in% c("Whole Blood",
                                                          "Pancreas",
                                                          "Uterus",
                                                          "Spleen",
                                                          "Lung",
                                                          "Pituitary",
                                                          "Skin (Suprapubic)",
                                                          "Breast (Mammary)",
                                                          "Prostate",
                                                          "Stomach",
                                                          "Muscle (Skeletal)",
                                                          "Liver",
                                                          "Brain (Spinal Cord)",
                                                          "Brain (Cortex)",
                                                          "Heart (Left Ventricle)",
                                                          "Brain (Putamen)"), ], 
       aes(x = tissue, 
           y = mito_total, 
           fill = tissue)) + 
  geom_violin(show.legend = FALSE, 
              scale = 1, 
              position = position_dodge(1), 
              lwd = 0.2) + 
  theme_classic() +
  theme(axis.text.x = element_text(angle = 30, 
                                   vjust = 1, 
                                   hjust=1, 
                                   family = "sans", 
                                   color = "black", 
                                   size = 8),
        axis.text.y = element_blank(),
        axis.title.x = element_blank(),
        axis.title.y = element_blank(),
        axis.line = element_line(size = 0.25),
        axis.ticks = element_line(size = 0.25),
        plot.margin = unit(c(0, 0, 0, 1.5), "cm"))

dev.off()

#### VIOLIN PLOT OF MITOCHONDRIAL EXPRESSION ACROSS CANCERS ####
# corresponds to Fig 3a

cancer_mito_expr_long <- data.frame(matrix(nrow = 0, ncol = 2))
colnames(cancer_mito_expr_long) <- c("mito_total", "cancer")

for(i in 1:length(cancertypes)){
  
  sampleIDs <- TCGA_LUT[TCGA_LUT$cancer == cancertypes[i], "SAMPID"]
  
  tempdata <- TCGA_TPM_data[mitogenes$ensembl_gene_id, colnames(TCGA_TPM_data) %in% sampleIDs]
  
  mitosums <- colSums(tempdata)
  
  temp_df <- data.frame(matrix(nrow = length(mitosums), ncol = 2))
  colnames(temp_df) <- c("mito_total", "cancer")
  
  temp_df[, "mito_total"] <- mitosums
  temp_df[, "cancer"] <- cancertypes[i]
  
  cancer_mito_expr_long <- rbind(cancer_mito_expr_long, temp_df)
  
}

cancer_mito_expr_long$mito_total <- cancer_mito_expr_long$mito_total / 1e4

cancer_mito_expr_long_unchanged <- cancer_mito_expr_long

cancer_mito_average <- aggregate(mito_total ~ cancer, cancer_mito_expr_long, mean)

cancer_mito_order <- cancer_mito_average[order(cancer_mito_average$mito_total, decreasing = FALSE), "cancer"]
cancer_mito_order <- str_remove(cancer_mito_order, "^TCGA-")

cancer_mito_expr_long$cancer <- str_remove(cancer_mito_expr_long$cancer, "^TCGA-")
cancer_mito_expr_long$cancer <- factor(cancer_mito_expr_long$cancer, levels = cancer_mito_order)

# plot for all of the cancers

pdf("graphics/TCGA-mt-expression-violin-all-cancers.pdf", width = 4.75, height = 2.75)

ggplot(data = cancer_mito_expr_long, aes(x = cancer, y = mito_total, fill = cancer)) + 
  geom_violin(show.legend = FALSE, scale = "area", position = position_dodge(1), lwd = 0.2) + 
  theme_classic() +
  theme(axis.text.x = element_text(angle = 90, vjust = 1, hjust=1, family = "sans", color = "black", size = 6),
        axis.text.y = element_blank(),
        axis.title.x = element_blank(),
        axis.title.y = element_blank(),
        axis.line = element_line(size = 0.25),
        axis.ticks = element_line(size = 0.25))+
  coord_cartesian(ylim = c(0, 95.5))

dev.off()

#### PLOT TISSUE MTOXPHOS VS NUOXPHOS EXPRESSION ####
# correspond to Fig 2c

# use MOR normalisation

total_mtOXPHOS_expression <- sapply(MOR_tissue_list, function(x){
  colSums(x[mtOXPHOS, ])
})

total_nuOXPHOS_expression <- sapply(MOR_tissue_list, function(x){
  colSums(x[nuOXPHOS, ])
})

testforplot <- cbind(sapply(total_mtOXPHOS_expression, mean), sapply(total_nuOXPHOS_expression, mean))
testforplot <- as.data.frame(testforplot)
testforplot[, "brain"] <- str_detect(row.names(testforplot), pattern = "^Brain")
testforplot$brain <- ifelse(testforplot$brain, "brain", "not")
colnames(testforplot) <- c("mtOXPHOS", "nuOXPHOS", "brain")

ln_reg_mtOXPHOSnuOXPHOS <- lm(nuOXPHOS ~ mtOXPHOS, data = testforplot)
sum_ln_reg_mtOXPHOSnuOXPHOS <- summary(ln_reg_mtOXPHOSnuOXPHOS)

spearcor_mtOXPHOSnuOXPHOS <- cor.test(testforplot$nuOXPHOS, testforplot$mtOXPHOS)

pdf("graphics/mtOXPHOS-nuOXPHOS-by-tissue.pdf",
    height = 2.2,
    width = 2.2)

ggplot(data = testforplot, aes(x = nuOXPHOS, y = mtOXPHOS))+
  geom_point(aes(color = brain), size = 0.6)+
  scale_color_manual(values = c("brain" = "red", "not" = "black"))+
  theme_classic() +
  geom_smooth(method = "lm", formula = y~ x, se = FALSE, size = 0.25) +
  theme(panel.border = element_rect(colour = "black", fill = NA, size = 0.25),
        axis.title.x = element_blank(),
        axis.line = element_line(size = 0.25),
        axis.ticks = element_line(size = 0.25),
        axis.title.y = element_blank(),
        axis.text.x = element_blank(),
        axis.text.y = element_blank(),
        legend.position = "none"
  ) + 
  coord_cartesian(xlim = c(0, 1200000),
                  ylim = c(0, 32000000))+
  geom_text(x = 750000, y = 30000000, label = deparse(bquote(R^2 == .(round(sum_ln_reg_mtOXPHOSnuOXPHOS$r.squared, 2)))), parse = TRUE, size = 2.3)+
  geom_text(x = 768000, y = 27500000, label = deparse(bquote(rho == .(round(as.numeric(spearcor_mtOXPHOSnuOXPHOS$estimate), 2)))), parse = TRUE, size = 2.3)

dev.off()

#### HEATMAPS FOR MTOXPHOS-NUOXPHOS COMBINING TISSUES ####
# corresponds to Fig 1a (TPM), Fig 2a (MRN/MOR), Fig S5a (TMM) 

# for TPM put the samples into a list
TPM_tissue_list <- list()

for(i in 1:length(tissues)){

  sampleIDs <- LUT[LUT$Tissue == tissues[i], "SAMPID"]
  tissueTPM <- TPMdata[, sampleIDs]
  row.names(tissueTPM) <- str_remove_all(row.names(tissueTPM), pattern = "\\..*$")
  TPM_tissue_list[[i]] <- tissueTPM
  names(TPM_tissue_list)[i] <- tissues[i]
  
}

saveRDS(TPM_tissue_list, "output/TPM_tissue_list.rds")

# here perform linear model corrections with all samples for each tissue for the OXPHOS genes and save in a list
MOR_OXPHOS_lmresiduals_list <- lapply(MOR_tissue_list, function(x){apply.lm.to.counts(x[c(mtOXPHOS, nuOXPHOS),])})
TMM_OXPHOS_lmresiduals_list <- lapply(TMM_tissue_list, function(x){apply.lm.to.counts(x[c(mtOXPHOS, nuOXPHOS),])})
TPM_OXPHOS_lmresiduals_list <- lapply(TPM_tissue_list, function(x){apply.lm.to.counts(x[c(mtOXPHOS, nuOXPHOS),])})
UQ_OXPHOS_lmresiduals_list <- lapply(UQ_tissue_list, function(x){apply.lm.to.counts(x[c(mtOXPHOS, nuOXPHOS),])})

# here put code in a function which can be applied to both lists of normalised values
make.combined.tissue.mtOXPHOS.nuOXPHOS.heatmaps <- function(tissuelist, 
                                                            iterations, 
                                                            graphics_subfolder = "", 
                                                            filename = "combined_heatmap",
                                                            additionalplotorder = NULL,
                                                            orderedfilename = "combined_heatmap_ordered"){

# create graphics subfolder, if it doesn't exist, to deposit plots
dir.create(paste0("graphics/", graphics_subfolder), showWarnings = FALSE)
  
# set colours for main plot (mainpal) and sidebar (sidepal)
mainpal <- (colorRampPalette(c("blue", "white", "red"))(100))
sidepal <- c("orange", "purple")

# make a vector with the sidebar information (nuOXPHOS or mtOXPHOS)
sidebar_vec <- OXPHOS_names[, "Sourcelist"]
  
# replace the values in the sidebar vector with colours specified by sidepal
frequencies <- table(sidebar_vec)[order(table(sidebar_vec), decreasing = TRUE)]
categories <- unique(names(frequencies))
  for(j in 1:length(categories)){
    sidebar_vec <- replace(sidebar_vec, which(sidebar_vec == categories[j]), sidepal[j])
  }
  
# assign name information to the sidebar_vec
names(sidebar_vec) <- OXPHOS_names[, "hgnc_symbol"]
  
# create list to deposit iterations
alltissues_list <- list()

for (i in 1:iterations) {

tissuesamples_rankspercent_df <- as.data.frame(matrix(nrow = nrow(OXPHOS_names), ncol = 0))
row.names(tissuesamples_rankspercent_df) <- OXPHOS_names$ensembl_gene_id

message(paste0("Starting iteration #", i))

for(j in 1:length(tissuelist)){

  tissuecounts <- tissuelist[[j]]
  
  # take 100 random samples from tissue and restrict to OXPHOS genes
  samplecounts <- tissuecounts[c(mtOXPHOS, nuOXPHOS), sample(colnames(tissuecounts), 100)]
  
  # rank the samples
  tempdata_ranks <- matrix(nrow = nrow(samplecounts), ncol = ncol(samplecounts))
  row.names(tempdata_ranks) <- row.names(samplecounts)
  colnames(tempdata_ranks) <- colnames(samplecounts)
  
  for (k in 1:nrow(samplecounts)){
    tempdata_ranks[k, ] <- rank(samplecounts[k, ])
  } 

  # here we add the ranks for this tissue to those previously calculated
  tissuesamples_rankspercent_df <- merge(tissuesamples_rankspercent_df, tempdata_ranks, all.x = TRUE, all.y = FALSE, by = "row.names")
  row.names(tissuesamples_rankspercent_df) <- tissuesamples_rankspercent_df[, "Row.names"]
  tissuesamples_rankspercent_df <- tissuesamples_rankspercent_df[, 2:ncol(tissuesamples_rankspercent_df)]
  
}

combinecorr <-find.all.correlations(tissuesamples_rankspercent_df, method = "spearman")

alltissues_list[[i]] <- combinecorr

}

iterations_matrix <- sapply(1:nrow(combinecorr), function(i){
  sapply(alltissues_list, function(x){x[i, 3]})
})

alliterations_mediancorr <- cbind(combinecorr[, 1:2], rowMedians(t(iterations_matrix)))
colnames(alliterations_mediancorr)[3] <- "MedianSpearman"

# build matrix for plotting heatmap
alliterations_clustermat <- build.matrix.4.cluster(corr_df = alliterations_mediancorr, 
                                                   stat = "MedianSpearman")

# rename rows and columns so gene symbols appear in plot
row.names(alliterations_clustermat) <- OXPHOS_names[row.names(alliterations_clustermat), "hgnc_symbol"]
colnames(alliterations_clustermat) <- OXPHOS_names[colnames(alliterations_clustermat), "hgnc_symbol"]

# ensures correct order of sidebar_vec
temptissue_sidebar_vec <- sidebar_vec[row.names(alliterations_clustermat)]

pdf(paste0(file = "graphics/", graphics_subfolder, "/", filename, ".pdf"))

heatmap <- heatmap.2(alliterations_clustermat,
          main = "All tissues combined",
          symkey = FALSE,
          symbreaks = TRUE,
          breaks = NULL,
          density.info = "none",
          trace = "none",
          margins = c(12,9),
          col = mainpal,
          dendrogram = "none",
          cexRow = 0.2,
          cexCol = 0.2,
          RowSideColors = temptissue_sidebar_vec)

par(lend = 1)           
legend("topright",      
       legend = categories,
       col = unique(names(table(sidebar_vec)[order(table(sidebar_vec), decreasing = TRUE)])),
       lty= 1,
       lwd = 10)

dev.off()

# save plot order according to dendogram produced with this method
plotorder <- row.names(alliterations_clustermat)[heatmap$rowInd]

functionoutput <- list()

functionoutput[["iterations"]] <- alltissues_list
functionoutput[["median_matrix"]] <- alliterations_clustermat
functionoutput[["heatmap_plot_order"]] <- plotorder

# if desired, produce ADDITIONAL plot ordered by a predefined order supplied in the arguments
if(!(is.null(additionalplotorder))){
    
    # build matrix for plotting heatmap
    alliterations_clustermat <- build.matrix.4.cluster(corr_df = alliterations_mediancorr, 
                                                       stat = "MedianSpearman",
                                                       plotorder = additionalplotorder)
  
  # rename rows and columns so gene symbols appear in plot
  row.names(alliterations_clustermat) <- OXPHOS_names[row.names(alliterations_clustermat), "hgnc_symbol"]
  colnames(alliterations_clustermat) <- OXPHOS_names[colnames(alliterations_clustermat), "hgnc_symbol"]
  
  # ensures correct order of sidebar_vec
  temptissue_sidebar_vec <- sidebar_vec[row.names(alliterations_clustermat)]
  
  pdf(paste0(file = "graphics/", graphics_subfolder, "/", orderedfilename, ".pdf"))
  
  heatmap <- heatmap.2(alliterations_clustermat,
                       main = "All tissues combined (reordered)",
                       symkey = FALSE,
                       symbreaks = TRUE,
                       breaks = NULL,
                       density.info = "none",
                       trace = "none",
                       margins = c(12,9),
                       col = mainpal,
                       Rowv = FALSE,
                       Colv = "Rowv",
                       dendrogram = "none",
                       cexRow = 0.2,
                       cexCol = 0.2,
                       RowSideColors = temptissue_sidebar_vec)
  
  par(lend = 1)           
  legend("topright",      
         legend = categories,
         col = unique(names(table(sidebar_vec)[order(table(sidebar_vec), decreasing = TRUE)])),
         lty= 1,
         lwd = 10)
  
  dev.off()
  
}

return(functionoutput)

}

MOR_OXPHOS_spearman_alltissues_combined <- make.combined.tissue.mtOXPHOS.nuOXPHOS.heatmaps(MOR_OXPHOS_lmresiduals_list,
                                                                                           iterations = 100,
                                                                                           graphics_subfolder = "tissues-combined",
                                                                                           filename = "MOR_OXPHOS_spearman_alltissues_combined")

saveRDS(MOR_OXPHOS_spearman_alltissues_combined, "output/MOR_OXPHOS_spearman_alltissues_combined.rds")
# MOR_OXPHOS_spearman_alltissues_combined <- readRDS("output/MOR_OXPHOS_spearman_alltissues_combined.rds")

# thjs plot order will be used to define the order of genes in all heatmaps published in this study
MOR_OXPHOS_spearman_plot_order <- OXPHOS_names[match(MOR_OXPHOS_spearman_alltissues_combined$heatmap_plot_order, OXPHOS_names$hgnc_symbol), "ensembl_gene_id"]

# extract mean correlation of mtOXPHOS-nuOXPHOS gene pairs
MORnumtOXPHOScorr <- mean(sapply(MOR_OXPHOS_spearman_alltissues_combined$iterations, function(x){median(x[(x[, "Gene1"] %in% mtOXPHOS|x[, "Gene2"] %in% mtOXPHOS) 
                                                                                                          & 
                                                                                                            (!(x[, "Gene1"] %in% mtOXPHOS)|!(x[, "Gene2"] %in% mtOXPHOS)), "rho"])}))

# extract mean correlation within mtOXPHOS genes (quoted in main text)
MORmtOXPHOSinternalcorr <- mean(sapply(MOR_OXPHOS_spearman_alltissues_combined$iterations, function(x){median(x[(x[, "Gene1"] %in% mtOXPHOS & x[, "Gene2"] %in% mtOXPHOS), "rho"])}))

# extract mean correlation within nuOXPHOS genes (quoted in main text)
MORnuOXPHOSinternalcorr <- mean(sapply(MOR_OXPHOS_spearman_alltissues_combined$iterations, function(x){median(x[(x[, "Gene1"] %in% nuOXPHOS & x[, "Gene2"] %in% nuOXPHOS), "rho"])}))

# repeat with TMM normalisation
TMM_OXPHOS_spearman_alltissues_combined <- make.combined.tissue.mtOXPHOS.nuOXPHOS.heatmaps(TMM_OXPHOS_lmresiduals_list,
                                                                                           iterations = 100,
                                                                                           graphics_subfolder = "tissues-combined",
                                                                                           filename = "TMM_OXPHOS_spearman_alltissues_combined",
                                                                                           additionalplotorder = MOR_OXPHOS_spearman_plot_order,
                                                                                           orderedfilename = "TMM_OXPHOS_spearman_alltissues_combined_MORorder")

saveRDS(TMM_OXPHOS_spearman_alltissues_combined, "output/TMM_OXPHOS_spearman_alltissues_combined.rds")
TMM_OXPHOS_spearman_plot_order <- OXPHOS_names[match(TMM_OXPHOS_spearman_alltissues_combined$heatmap_plot_order, OXPHOS_names$hgnc_symbol), "ensembl_gene_id"]

TPM_OXPHOS_spearman_alltissues_combined <- make.combined.tissue.mtOXPHOS.nuOXPHOS.heatmaps(TPM_OXPHOS_lmresiduals_list,
                                                                                           iterations = 100,
                                                                                           graphics_subfolder = "tissues-combined",
                                                                                           filename = "TPM_OXPHOS_spearman_alltissues_combined",
                                                                                           additionalplotorder = MOR_OXPHOS_spearman_plot_order,
                                                                                           orderedfilename = "TPM_OXPHOS_spearman_alltissues_combined_MORorder")

saveRDS(TPM_OXPHOS_spearman_alltissues_combined, "output/TPM_OXPHOS_spearman_alltissues_combined.rds")
TPM_OXPHOS_spearman_plot_order <- OXPHOS_names[match(TPM_OXPHOS_spearman_alltissues_combined$heatmap_plot_order, OXPHOS_names$hgnc_symbol), "ensembl_gene_id"]

# repeat with TPM normalisation
TPMnumtOXPHOScorr <- mean(sapply(TPM_OXPHOS_spearman_alltissues_combined$iterations, function(x){median(x[(x[, "Gene1"] %in% mtOXPHOS|x[, "Gene2"] %in% mtOXPHOS) 
                                                               & 
                                                                 (!(x[, "Gene1"] %in% mtOXPHOS)|!(x[, "Gene2"] %in% mtOXPHOS)), "rho"])}))

# extract mean mtOXPHOS correlation (quoted in main text)
TPMmtOXPHOSinternalcorr <- mean(sapply(TPM_OXPHOS_spearman_alltissues_combined$iterations, function(x){median(x[(x[, "Gene1"] %in% mtOXPHOS & x[, "Gene2"] %in% mtOXPHOS), "rho"])}))

# extract mean nuOXPHOS correlation (quoted in main text)
TPMnuOXPHOSinternalcorr <- mean(sapply(TPM_OXPHOS_spearman_alltissues_combined$iterations, function(x){median(x[(x[, "Gene1"] %in% nuOXPHOS & x[, "Gene2"] %in% nuOXPHOS), "rho"])}))

# repeat for UQ normalisation (not presented in manuscript)
UQ_OXPHOS_spearman_alltissues_combined <- make.combined.tissue.mtOXPHOS.nuOXPHOS.heatmaps(UQ_OXPHOS_lmresiduals_list,
                                                                                           iterations = 100,
                                                                                           graphics_subfolder = "tissues-combined",
                                                                                           filename = "UQ_OXPHOS_spearman_alltissues_combined",
                                                                                           additionalplotorder = MOR_OXPHOS_spearman_plot_order,
                                                                                           orderedfilename = "UQ_OXPHOS_spearman_alltissues_combined_MORorder")

saveRDS(UQ_OXPHOS_spearman_alltissues_combined, "output/UQ_OXPHOS_spearman_alltissues_combined.rds")
UQ_OXPHOS_spearman_plot_order <- OXPHOS_names[match(UQ_OXPHOS_spearman_alltissues_combined$heatmap_plot_order, OXPHOS_names$hgnc_symbol), "ensembl_gene_id"]

#### DISTRIBUTION FOR ALL TISSUES COMBINED WITH mtOXPHOS AND RANDOM NUCLEAR FACTORS ####
# correspond to Fig 2b (GTEX MRN)

# here put code in a function which can be applied to both lists of normalised values
generate.distribution.mtOXPHOS.random.medians <- function(tissuelist, 
                                                            iterations, 
                                                            method = c("spearman", "pearson"),
                                                            stat = c("rho", "r"))
                                                            {

  method <- match.arg(method)
  stat <- match.arg(stat)
  
  # create list to deposit iterations
  alltissues_list <- list()

  for (i in 1:iterations) {
    
    message(paste0("Starting iteration #", i))
    
    # take nuclear sample of 126 non-mito genes (same size as nuOXPHOS group)
    nuclear_sample <- sample(nuclear_gene_expressed, size = 126)

    randomnuclearsample_residuals <- lapply(tissuelist, function(x){
      apply.lm.to.counts(x[c(mtOXPHOS, nuclear_sample), ])
    })

    tissuesamples_rankspercent_df <- data.frame(matrix(nrow = length(c(mtOXPHOS, nuclear_sample)), ncol = 0))
    row.names(tissuesamples_rankspercent_df) <- c(mtOXPHOS, nuclear_sample)
    
    for(j in 1:length(randomnuclearsample_residuals)){

      # take 100 random samples from the tissue
      samplecounts <- randomnuclearsample_residuals[[j]][, sample(colnames(randomnuclearsample_residuals[[j]]), 100)]
      
      # rank the samples
      tempdata_ranks <- matrix(nrow = nrow(samplecounts), ncol = ncol(samplecounts))
      row.names(tempdata_ranks) <- row.names(samplecounts)
      colnames(tempdata_ranks) <- colnames(samplecounts)
      
      for (k in 1:nrow(samplecounts)){
        tempdata_ranks[k, ] <- rank(samplecounts[k, ])
      } 
      
      # here we add the ranks for this tissue to those previously calculated
      tissuesamples_rankspercent_df <- merge(tissuesamples_rankspercent_df, tempdata_ranks, all.x = TRUE, all.y = FALSE, by = "row.names")
      row.names(tissuesamples_rankspercent_df) <- tissuesamples_rankspercent_df[, "Row.names"]
      tissuesamples_rankspercent_df <- tissuesamples_rankspercent_df[, 2:ncol(tissuesamples_rankspercent_df)]
      
    }
    
    combinecorr <- find.all.correlations(tissuesamples_rankspercent_df, 
                                         method = method,
                                         stat = stat)
    
    alltissues_list[[i]] <- combinecorr
    
  } # end of looping over iterations

  if(method == "spearman"){
    average <- function(x){
      median(x)
    }
  }
  
  if(method == "pearson"){
    average <- function(x){
      mean(x)
    }
  }
  
 iteration_avg <- sapply(alltissues_list, function(x){average(x[(x[, "Gene1"] %in% mtOXPHOS|x[, "Gene2"] %in% mtOXPHOS) 
                                                                   & 
                                                                     (!(x[, "Gene1"] %in% mtOXPHOS)|!(x[, "Gene2"] %in% mtOXPHOS)), paste(stat)])})

 functionoutput <- list()
 
 functionoutput[["iterations"]] <- alltissues_list
 functionoutput[["mtOXPHOS_random_avg"]] <- iteration_avg
  
 return(functionoutput)
 
  }
  
MOR_alltissues_mtOXPHOSrandomdist_spearman <- generate.distribution.mtOXPHOS.random.medians(tissuelist = MOR_tissue_list,
                                              iterations = 100,
                                              method = "spearman",
                                              stat = "rho")

saveRDS(MOR_alltissues_mtOXPHOSrandomdist_spearman, "output/MOR_alltissues_mtOXPHOSrandomdist_spearman.rds")
#MOR_alltissues_mtOXPHOSrandomdist_spearman <- readRDS("output/MOR_alltissues_mtOXPHOSrandomdist_spearman.rds")
#MOR_OXPHOS_spearman_alltissues_combined <- readRDS("output/MOR_OXPHOS_spearman_alltissues_combined.rds")

# perform Wilcox/t test on medians from mt-nuOXPHOS tissues combined vs medians from mtOXPHOS-random nuclear combined

MOR_mtOXPHOSnuOXPHOSorrandom_iterations <- data.frame(cbind(sapply(MOR_OXPHOS_spearman_alltissues_combined$iterations, function(x){median(x[(x[, "Gene1"] %in% mtOXPHOS|x[, "Gene2"] %in% mtOXPHOS) 
                                                                                      & 
                                                                                        (!(x[, "Gene1"] %in% mtOXPHOS)|!(x[, "Gene2"] %in% mtOXPHOS)), "rho"])})),
                                                      MOR_alltissues_mtOXPHOSrandomdist_spearman$mtOXPHOS_random_avg)

colnames(MOR_mtOXPHOSnuOXPHOSorrandom_iterations) <- c("mtOXPHOSnuOXPHOS", "mtOXPHOSrandomnuclear")

MOR_mtOXPHOSnuOXPHOS_wilcoxtest_spearman <- wilcox.test(MOR_mtOXPHOSnuOXPHOSorrandom_iterations$mtOXPHOSnuOXPHOS,
                                                        MOR_mtOXPHOSnuOXPHOSorrandom_iterations$mtOXPHOSrandomuclear)
MOR_mtOXPHOSnuOXPHOS_ttest_spearman <- t.test(MOR_mtOXPHOSnuOXPHOSorrandom_iterations$mtOXPHOSnuOXPHOS,
                                              MOR_mtOXPHOSnuOXPHOSorrandom_iterations$mtOXPHOSrandomnuclear)

GTEXforplottogether <- as.data.frame(cbind(rowMeans(t(MOR_mtOXPHOSnuOXPHOSorrandom_iterations)),
                                           rowSds(t(MOR_mtOXPHOSnuOXPHOSorrandom_iterations))))
colnames(GTEXforplottogether) <- c("mean", "sd")
GTEXforplottogether[, "group"] <- c("mtOXPHOS - nuOXPHOS",
                                    "mtOXPHOS - random nuclear")
GTEXforplottogether$group <- relevel(factor(GTEXforplottogether$group),
                                     "mtOXPHOS - random nuclear")

# plot a boxplot of the two distributions
pdf("graphics/mtOXPHOS_randomnuclear/MOR_mtOXPHOSrandom_tissuecombined_boxplot.pdf",
    width = 2.2,
    height = 2.2)

ggplot(data = GTEXforplottogether, aes(x = group,
                                       y = mean))+
  geom_col(fill = c("magenta", "grey"),
               lwd = 0.2, size = 0.25, col = "black")+
  geom_hline(yintercept = 0,
             linetype = "dashed",
             color = "grey",
             size = 0.25)+
  geom_errorbar(aes(x = group,
                    ymin = mean - sd,
                    ymax = mean +sd),
                width = 0.5,
                size = 0.25)+
  theme_classic() + 
  coord_cartesian(ylim = c(-0.2, 0.2)) +
  theme(panel.border = element_rect(colour = "black",
                                    fill = NA,
                                    size = 0.25),
        axis.title.x = element_blank(),
        axis.line = element_line(size = 0.25),
        axis.ticks = element_line(size = 0.25),
        axis.title.y = element_blank(),
        axis.text.y = element_blank(),
        axis.text.x = element_blank())

dev.off()

# repeat mtOXPHOS-random nuclear for all tissues combined for TMM normalisation 
# corresponds to Fig S5b

TMM_alltissues_mtOXPHOSrandomdist_spearman <- generate.distribution.mtOXPHOS.random.medians(tissuelist = TMM_tissue_list,
                                                                                            iterations = 100,
                                                                                            method = "spearman",
                                                                                            stat = "rho")

saveRDS(TMM_alltissues_mtOXPHOSrandomdist_spearman, "output/TMM_alltissues_mtOXPHOSrandomdist_spearman.rds")
#TMM_alltissues_mtOXPHOSrandomdist_spearman <- readRDS("output/TMM_alltissues_mtOXPHOSrandomdist_spearman.rds")
#TMM_OXPHOS_spearman_alltissues_combined <- readRDS("output/TMM_OXPHOS_spearman_alltissues_combined.rds")

# perform Wilcox/t test on medians from mt-nuOXPHOS tissues combined vs medians from mtOXPHOS-random nuclear combined

TMM_mtOXPHOSnuOXPHOSorrandom_iterations <- data.frame(cbind(sapply(TMM_OXPHOS_spearman_alltissues_combined$iterations, function(x){median(x[(x[, "Gene1"] %in% mtOXPHOS|x[, "Gene2"] %in% mtOXPHOS) 
                                                                                                                                            & 
                                                                                                                                              (!(x[, "Gene1"] %in% mtOXPHOS)|!(x[, "Gene2"] %in% mtOXPHOS)), "rho"])})),
                                                      TMM_alltissues_mtOXPHOSrandomdist_spearman$mtOXPHOS_random_avg)

colnames(TMM_mtOXPHOSnuOXPHOSorrandom_iterations) <- c("mtOXPHOSnuOXPHOS", "mtOXPHOSrandomnuclear")

TMM_mtOXPHOSnuOXPHOS_wilcoxtest_spearman <- wilcox.test(TMM_mtOXPHOSnuOXPHOSorrandom_iterations$mtOXPHOSnuOXPHOS, TMM_mtOXPHOSnuOXPHOSorrandom_iterations$mtOXPHOSrandom)
TMM_mtOXPHOSnuOXPHOS_ttest_spearman <- t.test(TMM_mtOXPHOSnuOXPHOSorrandom_iterations$mtOXPHOSnuOXPHOS, TMM_mtOXPHOSnuOXPHOSorrandom_iterations$mtOXPHOSrandom)

GTEXforplottogetherTMM <- as.data.frame(cbind(rowMeans(t(TMM_mtOXPHOSnuOXPHOSorrandom_iterations)), rowSds(t(TMM_mtOXPHOSnuOXPHOSorrandom_iterations))))
colnames(GTEXforplottogetherTMM) <- c("mean", "sd")
GTEXforplottogetherTMM[, "group"] <- c("mtOXPHOS - nuOXPHOS", "mtOXPHOS - random nuclear")
GTEXforplottogetherTMM$group <- relevel(factor(GTEXforplottogetherTMM$group), "mtOXPHOS - random nuclear")

# plot a boxplot of the two distributions
pdf("graphics/mtOXPHOS_randomnuclear/TMM_mtOXPHOSrandom_tissuecombined_boxplot.pdf",
    width = 2.2,
    height = 2.2)

ggplot(data = GTEXforplottogetherTMM, aes(x = group, y = mean))+
  geom_col(fill = c("magenta", "grey"),
           lwd = 0.2, size = 0.25, col = "black")+
  geom_hline(yintercept = 0, linetype = "dashed", color = "grey", size = 0.25)+
  geom_errorbar(aes(x = group, ymin = mean - sd, ymax = mean +sd), width = 0.5, size = 0.25)+
  theme_classic() + 
  coord_cartesian(ylim = c(-0.2, 0.2)) +
  theme(panel.border = element_rect(colour = "black", fill = NA, size = 0.25),
        axis.title.x = element_blank(),
        axis.line = element_line(size = 0.25),
        axis.ticks = element_line(size = 0.25),
        axis.title.y = element_blank(),
        axis.text.y = element_blank(),
        axis.text.x = element_blank())

dev.off()

#### PRODUCE SINGLE TISSUE HEATMAPS AND OTHER PLOTS FOR MTOXPHOS - NUOXPHOS CORRELATIONS ####
# corresponds to Fig 1c, Fig 2d, Fig S1b, Fig S1c, Fig S1d, Fig S5c, Suplementary Dataset 1

tissue.mtOXPHOS.nuOXPHOS.heatmaps.from.list <- function(tissuelist, 
                                                        graphics_subfolder = "", 
                                                        spearplotorder = NULL, 
                                                        pearsonplotorder = NULL){

  dir.create(paste0("graphics/", graphics_subfolder), showWarnings = FALSE)
  dir.create(paste0("graphics/", graphics_subfolder, "/spearman"), showWarnings = FALSE)
  dir.create(paste0("graphics/", graphics_subfolder, "/pearson"), showWarnings = FALSE)
  
  # set colours for main plot (mainpal) and sidebar (sidepal)
  mainpal <- (colorRampPalette(c("blue", "white", "red"))(100))
  sidepal <- c("orange", "purple")
  
  # make a vector with the sidebar information (nuOXPHOS or mtOXPHOS)
  sidebar_vec <- OXPHOS_names[, "Sourcelist"]
  
  # replace the values in the sidebar vector with colours specified by sidepal
  frequencies <- table(sidebar_vec)[order(table(sidebar_vec), decreasing = TRUE)]
  categories <- unique(names(frequencies))
  for(j in 1:length(categories)){
    sidebar_vec <- replace(sidebar_vec, which(sidebar_vec == categories[j]), sidepal[j])
  }
  
  # assign name information to the sidebar_vec
  names(sidebar_vec) <- OXPHOS_names[, "hgnc_symbol"]
  
  tissue_mean_corr_df <- as.data.frame(matrix(nrow = length(tissuelist), ncol = 3))
  colnames(tissue_mean_corr_df) <- c("tissue", "total_mt_expr", "no_samples")
  
  tissue_corr_spearman_list <- list()
  tissue_corr_pearson_list <- list()
  
  for(i in 1:length(tissuelist)){
  
  # take normalised counts previously saved in list
  tissuecounts <- tissuelist[[i]]
  
  # put tissue name in data frame
  tissue_mean_corr_df[i, "tissue"] <- names(tissuelist)[i]
  
  # put tissue mean mitochondrial expression (as % TPM) in data frame
  tissue_mean_corr_df[i, "total_mt_expr"] <- mean(colSums(TPMdata[mitogenes$ensembl_gene_id, colnames(tissuecounts)])) / 1e4
  tissue_mean_corr_df[i, "no_samples"] <- ncol(tissuecounts)
  
  # find all correlations using Spearman's rank correlation coefficient (rho)
  tissue_corr_spear <- find.all.correlations(tissuecounts, method = "spearman", stat = "rho")
  tissue_corr_spear[,"Gene1name"] <- OXPHOS_names[tissue_corr_spear$Gene1, "hgnc_symbol"]
  tissue_corr_spear[,"Gene2name"] <- OXPHOS_names[tissue_corr_spear$Gene2, "hgnc_symbol"]
  
  tissue_corr_spearman_list[[i]] <- tissue_corr_spear
  names(tissue_corr_spearman_list)[i] <- names(tissuelist)[i]
  
  # put the mito-nuclear correlation in the dataframe
  tissue_mean_corr_df[i, "median_mtnuOXPHOS_spearman"]  <- median(tissue_corr_spear[(tissue_corr_spear$Gene1 %in% mtOXPHOS | tissue_corr_spear$Gene2 %in% mtOXPHOS) 
                                                                                & 
                                                                                (tissue_corr_spear$Gene1 %in% nuOXPHOS | tissue_corr_spear$Gene2 %in% nuOXPHOS)
                                                                                , "rho"])
  
  tissue_mean_corr_df[i, "Q1_mtnuOXPHOS_spearman"]  <- quantile(tissue_corr_spear[(tissue_corr_spear$Gene1 %in% mtOXPHOS | tissue_corr_spear$Gene2 %in% mtOXPHOS) 
                                                                            &
                                                                              (tissue_corr_spear$Gene1 %in% nuOXPHOS | tissue_corr_spear$Gene2 %in% nuOXPHOS)
                                                                            , "rho"], 0.25)
  
  tissue_mean_corr_df[i, "Q3_mtnuOXPHOS_spearman"]  <- quantile(tissue_corr_spear[(tissue_corr_spear$Gene1 %in% mtOXPHOS | tissue_corr_spear$Gene2 %in% mtOXPHOS) 
                                                                                  &
                                                                                    (tissue_corr_spear$Gene1 %in% nuOXPHOS | tissue_corr_spear$Gene2 %in% nuOXPHOS)
                                                                                  , "rho"], 0.75)
  
  # find all correlations using Pearson's product-moment correlation coefficient
  
  tissue_corr_pearson <- find.all.correlations(tissuecounts, method = "pearson", stat = "r")
  tissue_corr_pearson[,"Gene1name"] <- OXPHOS_names[tissue_corr_pearson$Gene1, "hgnc_symbol"]
  tissue_corr_pearson[,"Gene2name"] <- OXPHOS_names[tissue_corr_pearson$Gene2, "hgnc_symbol"]
  
  tissue_corr_pearson_list[[i]] <- tissue_corr_pearson
  names(tissue_corr_pearson_list)[i] <- names(tissuelist)[i]
  
  # put the mito-nuclear correlation in the dataframe
  tissue_mean_corr_df[i, "mean_mtnuOXPHOS_pearson"]  <- mean(tissue_corr_pearson[(tissue_corr_pearson$Gene1 %in% mtOXPHOS | tissue_corr_pearson$Gene2 %in% mtOXPHOS) 
                                                                                    & 
                                                                                      (tissue_corr_pearson$Gene1 %in% nuOXPHOS | tissue_corr_pearson$Gene2 %in% nuOXPHOS)
                                                                                    , "r"])
  
  tissue_mean_corr_df[i, "sd_mtnuOXPHOS_pearson"]  <- sd(tissue_corr_pearson[(tissue_corr_pearson$Gene1 %in% mtOXPHOS | tissue_corr_pearson$Gene2 %in% mtOXPHOS) 
                                                                            &
                                                                              (tissue_corr_pearson$Gene1 %in% nuOXPHOS | tissue_corr_pearson$Gene2 %in% nuOXPHOS)
                                                                            , "r"])
  
  # make correlation matrix for plotting (with spearman); then
  # change row/column names so that gene symbols appear on plots
  tissue_spear_clustermat <- build.matrix.4.cluster(corr_df = tissue_corr_spear,
                                                    stat = "rho",
                                                    plotorder = spearplotorder)
  
  row.names(tissue_spear_clustermat) <- OXPHOS_names[row.names(tissue_spear_clustermat), "hgnc_symbol"]
  colnames(tissue_spear_clustermat) <- OXPHOS_names[colnames(tissue_spear_clustermat), "hgnc_symbol"]
  
  # make correlation matrix for plotting (with pearson); then
  # change row/column names so that gene symbols appear on plots
  tissue_pearson_clustermat <- build.matrix.4.cluster(corr_df = tissue_corr_pearson, 
                                                      stat = "r", 
                                                      plotorder = pearsonplotorder)
  
  row.names(tissue_pearson_clustermat) <- OXPHOS_names[row.names(tissue_pearson_clustermat), "hgnc_symbol"]
  colnames(tissue_pearson_clustermat) <- OXPHOS_names[colnames(tissue_pearson_clustermat), "hgnc_symbol"]
  
  # ensures correct order of sidebar_vec
  temptissue_spear_sidebar_vec <- sidebar_vec[row.names(tissue_spear_clustermat)]
  
  pdf(paste0(file = "graphics/", graphics_subfolder, "/spearman/", paste0(unlist(str_split(names(tissuelist)[i], pattern = " ")), collapse = ""), ".pdf"))
  
  heatmap.2(tissue_spear_clustermat,
            main = paste0(names(tissuelist)[i], ", n = ", ncol(tissuecounts)),
            symkey = FALSE,
            symbreaks = TRUE,
            breaks = NULL,
            density.info = "none",
            trace = "none",
            margins = c(12,9),
            col = mainpal,
            Rowv = FALSE,
            Colv = "Rowv",
            dendrogram = "none",
            cexRow = 0.2,
            cexCol = 0.2,
            RowSideColors = temptissue_spear_sidebar_vec)
  
  par(lend = 1)
  legend("topright",      
         legend = categories,
         col = unique(names(table(sidebar_vec)[order(table(sidebar_vec), decreasing = TRUE)])),
         lty= 1,
         lwd = 10)
  
  dev.off()
  
  # ensures correct order of sidebar_vec
  temptissue_pearson_sidebar_vec <- sidebar_vec[row.names(tissue_pearson_clustermat)]
  
  pdf(paste0(file = "graphics/", graphics_subfolder, "/pearson/", paste0(unlist(str_split(names(tissuelist)[i], pattern = " ")), collapse = ""), ".pdf"))
  
  heatmap.2(tissue_pearson_clustermat,
            main = paste0(names(tissuelist)[i], ", n = ", ncol(tissuecounts)),
            symkey = FALSE,
            symbreaks = TRUE,
            breaks = NULL,
            density.info = "none",
            trace = "none",
            margins = c(12,9),
            col = mainpal,
            Rowv = FALSE,
            Colv = "Rowv",
            dendrogram = "none",
            cexRow = 0.2,
            cexCol = 0.2,
            RowSideColors = temptissue_pearson_sidebar_vec)
  
  par(lend = 1)           
  legend("topright",      
         legend = categories,
         col = unique(names(table(sidebar_vec)[order(table(sidebar_vec), decreasing = TRUE)])),
         lty= 1,
         lwd = 10)
  
  dev.off()
  
}

  ln_reg_prs <- lm(mean_mtnuOXPHOS_pearson ~ total_mt_expr, data = tissue_mean_corr_df)
  sum_ln_reg_prs <- summary(ln_reg_prs)
  
  pdf(paste0(file = "graphics/", graphics_subfolder,  "/mtOXPHOS-nuOXPHOS-Pearson-across-tissues.pdf"))
  
  print(ggplot(data = tissue_mean_corr_df, aes(x = total_mt_expr, y = mean_mtnuOXPHOS_pearson)) +
    ylab("Mean Pearson's r between mtOXPHOS and nuOXPHOS genes") + 
    xlab("% mitochondrial transcripts") +
    ggtitle("mtOXPHOS-nuOXPHOS correlations") +
    geom_point() +
    geom_smooth(method = "lm", formula = y~ x, se = FALSE) +
    geom_errorbar(aes(x = total_mt_expr, ymax = mean_mtnuOXPHOS_pearson + sd_mtnuOXPHOS_pearson, ymin = mean_mtnuOXPHOS_pearson - sd_mtnuOXPHOS_pearson)) +
    theme_classic() +
    coord_cartesian(ylim = c(-1, 1)) +
    geom_text(x = 60, y = 0.5, label = deparse(bquote(R^2 == .(round(sum_ln_reg_prs$r.squared, 2)))), parse = TRUE) +
    geom_hline(yintercept = 0, linetype = "dashed", color = "grey")
  )
    
  dev.off()
  
  ln_reg_spr <- lm(median_mtnuOXPHOS_spearman ~ total_mt_expr, data = tissue_mean_corr_df)
  sum_ln_reg_spr <- summary(ln_reg_spr)
  
  pdf(paste0(file = "graphics/", graphics_subfolder,  "/mtOXPHOS-nuOXPHOS-Spearman-across-tissues.pdf"))
  
  print(ggplot(data = tissue_mean_corr_df, aes(x = total_mt_expr, y = median_mtnuOXPHOS_spearman)) +
    ylab("Median spearman's rho between mtOXPHOS and nuOXPHOS genes") + 
    xlab("% mitochondrial transcripts") +
    ggtitle("mtOXPHOS-nuOXPHOS correlations") +
    geom_point() +
    geom_smooth(method = "lm", formula = y~ x, se = FALSE) +
    geom_errorbar(aes(x = total_mt_expr, ymax = Q3_mtnuOXPHOS_spearman, ymin = Q1_mtnuOXPHOS_spearman)) +
    theme_classic() +
    coord_cartesian(ylim = c(-1, 1)) +
    geom_text(x = 60, y = 0.5, label = deparse(bquote(R^2 == .(round(sum_ln_reg_spr$r.squared, 2)))), parse = TRUE) +
    geom_hline(yintercept = 0, linetype = "dashed", color = "grey")
  )
  
  dev.off()
  
  mito_cv_vec <- vector()
  
  for(i in 1:length(tissues)){

    tempvec <- mito_expr_long_unchanged[mito_expr_long_unchanged$tissue == tissues[i], "mito_total"]

    mito_cv_vec[i] <- sd(tempvec) / mean(tempvec)
    
  }
  
  names(mito_cv_vec) <- tissues
  
 tissue_mean_corr_df[, "mito_expr_cv"] <- mito_cv_vec
  
 output_list <- list()
 
 output_list[["tissue_corr_spearman_list"]] <- tissue_corr_spearman_list
 output_list[["tissue_corr_pearson_list"]] <- tissue_corr_pearson_list
 output_list[["tissue_mean_corr_df"]] <- tissue_mean_corr_df
 
 
  return(output_list)
  
}

MOR_tissue_correlations <- tissue.mtOXPHOS.nuOXPHOS.heatmaps.from.list(tissuelist = MOR_OXPHOS_lmresiduals_list,
                                                                       graphics_subfolder = "mtOXPHOS-nuOXPHOS-MOR-normalisation-tissue",
                                                                       spearplotorder = MOR_OXPHOS_spearman_plot_order,
                                                                       pearsonplotorder = MOR_OXPHOS_spearman_plot_order)

saveRDS(MOR_tissue_correlations, "output/MOR-OXPHOS-correlation-by-tissue.rds")
# MOR_tissue_correlations <- readRDS("output/MOR-OXPHOS-correlation-by-tissue.rds")

TMM_tissue_correlations <- tissue.mtOXPHOS.nuOXPHOS.heatmaps.from.list(tissuelist = TMM_OXPHOS_lmresiduals_list,
                                                                       graphics_subfolder = "mtOXPHOS-nuOXPHOS-TMM-normalisation-tissue",
                                                                       spearplotorder = MOR_OXPHOS_spearman_plot_order,
                                                                       pearsonplotorder = MOR_OXPHOS_spearman_plot_order)

saveRDS(TMM_tissue_correlations, "output/TMM-OXPHOS-correlation-by-tissue.rds")

TPM_tissue_correlations <- tissue.mtOXPHOS.nuOXPHOS.heatmaps.from.list(tissuelist = TPM_OXPHOS_lmresiduals_list,
                                                                       graphics_subfolder = "mtOXPHOS-nuOXPHOS-TPM-normalisation-tissue",
                                                                       spearplotorder = MOR_OXPHOS_spearman_plot_order,
                                                                       pearsonplotorder = MOR_OXPHOS_spearman_plot_order)

saveRDS(TPM_tissue_correlations, "output/TPM-OXPHOS-correlation-by-tissue.rds")

UQ_tissue_correlations <- tissue.mtOXPHOS.nuOXPHOS.heatmaps.from.list(tissuelist = UQ_OXPHOS_lmresiduals_list,
                                                                       graphics_subfolder = "mtOXPHOS-nuOXPHOS-UQ-normalisation-tissue",
                                                                       spearplotorder = MOR_OXPHOS_spearman_plot_order,
                                                                       pearsonplotorder = MOR_OXPHOS_spearman_plot_order)

saveRDS(UQ_tissue_correlations, "output/UQ-OXPHOS-correlation-by-tissue.rds")

# will replicate analysis for TPM with nuclear factors scaled by mitochondrial exclusion

readcounts_nomito <- countsdata[!(str_remove_all(rownames(countsdata), pattern = "\\..*$") %in% mitogenes$ensembl_gene_id),]

colsums_readcounts_nomito <- colSums(readcounts_nomito)

scalefactor_nuclear <- colsums_readcounts_nomito / 1e6
scalefactor_mito <- colSums(countsdata) / 1e6

TPMnomito_tissue_list <- list()

for (i in 1:length(tissues)){

  sampleIDs <- LUT[LUT$Tissue == tissues[i], "SAMPID"]
  
  mitonormalisation <- t(countsdata[mitogenes$ensembl_gene_id, sampleIDs]) / scalefactor_mito[sampleIDs]
  nuclearnormalisation <- t(countsdata[-match(mitogenes$ensembl_gene_id, row.names(countsdata)), sampleIDs ]) / scalefactor_nuclear[sampleIDs]
  
  TPMnomito_tissue_list[[i]] <- t(cbind(mitonormalisation, nuclearnormalisation))
  
  names(TPMnomito_tissue_list)[i] <- tissues[i]
  
}

saveRDS(TPMnomito_tissue_list, "output/TPMnomito-tissue-list.rds")

TPMnomito_OXPHOS_lmresiduals_list <- lapply(TPMnomito_tissue_list, function(x){apply.lm.to.counts(x[c(mtOXPHOS, nuOXPHOS),])})

TPMnomito_tissue_correlations <- tissue.mtOXPHOS.nuOXPHOS.heatmaps.from.list(tissuelist = TPMnomito_OXPHOS_lmresiduals_list,
                                                                       graphics_subfolder = "mtOXPHOS-nuOXPHOS-TPMnomito-normalisation-tissue",
                                                                       spearplotorder = MOR_OXPHOS_spearman_plot_order,
                                                                       pearsonplotorder = MOR_OXPHOS_spearman_plot_order)

saveRDS(TPMnomito_tissue_correlations, "output/TPMnomito-OXPHOS-correlation-by-tissue.rds")

#### BOOTSTRAP FOR CONFIDENCE INTERVALS ####
# For Fig 1d, Fig S5c

allOXPHOSpairs <- combinations(length(OXPHOS_names$ensembl_gene_id), 2, OXPHOS_names$ensembl_gene_id)
colnames(allOXPHOSpairs) <- c("Gene1", "Gene2")

mtnuOXPHOSpairs <- allOXPHOSpairs[(allOXPHOSpairs[, "Gene1"] %in% mtOXPHOS|allOXPHOSpairs[, "Gene2"] %in% mtOXPHOS) 
                                  & 
                                    (!(allOXPHOSpairs[, "Gene1"] %in% mtOXPHOS)|!(allOXPHOSpairs[, "Gene2"] %in% mtOXPHOS)),]

return.median.correlation <- function(expression_df, genecombo){
  
  output_vec <- vector(length = nrow(genecombo))
  
  for (i in 1:nrow(genecombo)){
    
    x <- unlist(expression_df[unlist(genecombo[i, 1]), ])
    y <- unlist(expression_df[unlist(genecombo[i, 2]), ])
    
    output_vec[i]  <- cor.test(x, 
                               y,
                               method = "spearman",
                               exact = FALSE)$estimate
    
  }
  
  return(median(output_vec, na.rm = TRUE))
  
}

bootstrap.OXPHOS.medians <- function(datalist, no_of_reps, whichLUT = LUT, which.apply.lm = apply.lm.to.counts){
  
  output_matrix <- matrix(nrow = no_of_reps, ncol = length(datalist))
  colnames(output_matrix) <- names(datalist)
  
  lm_list <- lapply(datalist, function(x){
    which.apply.lm(countdata = x[c(mtOXPHOS, nuOXPHOS), colnames(x) %in% whichLUT$SAMPID], lookuptable = whichLUT)
  })
  
  samplesinlist <- unlist(sapply(lm_list, colnames))
  
  samplinglist <-    lapply(datalist, function(x){
    rep_sample_n(data.frame(colnames(x)[colnames(x) %in% samplesinlist]), size = ncol(x), replace = TRUE, reps = no_of_reps)
  })
  
  samplinglist <- lapply(samplinglist, function(x){
    setNames(x, nm = c("replicate", "sample"))
  })
  
  b4 <- Sys.time()
  
  for(i in 1:no_of_reps){
    
    for(j in 1:length(datalist)){
      
      samplecols <- samplinglist[[j]][samplinglist[[j]]$replicate == i, "sample"]
      unlist(samplecols)[!(unlist(samplecols) %in% colnames(lm_list[[j]]))]
      
      expression_df <- lm_list[[j]][c(mtOXPHOS, nuOXPHOS), unlist(samplecols)]
      
      output_matrix[i, j] <- return.median.correlation(expression_df, genecombo = mtnuOXPHOSpairs)
      
    }
    
    if (i %% 10 == 0) {
      
      message(paste("Combination", i, sep = " "))
      
      time_elapsed <- Sys.time() - b4
      message(paste("Time elapsed:", round(as.numeric(time_elapsed), digits = 2), attr(time_elapsed, "units"), sep = " "))
      
      timeperiteration <- time_elapsed / (i)
      
      remaining <- no_of_reps - i
      message(paste("Estimated time remaining:", round((as.numeric(timeperiteration, units = "hours") * remaining), digits = 2), "hours", sep = " "))
      
    }
    
  }
  
  return(output_matrix)
  
}

# MOR_tissue_list <- readRDS("output/MOR-normalisation-by-tissue-list.rds")
# TMM_tissue_list <- readRDS("output/TMM-normalisation-by-tissue-list.rds")
# LUT <- readRDS("output/GTEX-version8-sampleID-LUT.rds")

GTEX_mtnuOXPHOSmedian_bootstrap <- bootstrap.OXPHOS.medians(datalist = MOR_tissue_list,
                                                            no_of_reps = 1000,
                                                            whichLUT = LUT,
                                                            which.apply.lm = apply.lm.to.counts)

saveRDS(GTEX_mtnuOXPHOSmedian_bootstrap, "output/GTEX_mtnuOXPHOSmedian_bootstrap.rds")

GTEX_TMMmtnuOXPHOSmedian_bootstrap <- bootstrap.OXPHOS.medians(datalist = TMM_tissue_list,
                                                               no_of_reps = 1000,
                                                               whichLUT = LUT,
                                                               which.apply.lm = apply.lm.to.counts)

saveRDS(GTEX_TMMmtnuOXPHOSmedian_bootstrap, "output/GTEX_TMMmtnuOXPHOSmedian_bootstrap.rds")

TCGA_mtnuOXPHOSmedian_bootstrap <- bootstrap.OXPHOS.medians(datalist = TCGA_MOR_list, 
                                                            no_of_reps = 1000,
                                                            whichLUT = TCGA_LUT,
                                                            which.apply.lm = TCGA.apply.lm.to.counts)

saveRDS(TCGA_mtnuOXPHOSmedian_bootstrap, "output/TCGA_mtnuOXPHOSmedian_bootstrap.rds")

#### INVESTIGATE mtOXPHOS CORRELATIONS WITH RANDOM NUCLEAR GENES WITHIN TISSUES ####
# corresponds to Fig 1d, Fig S1c, Fig S2a, Fig S4a, Fig S4b, Table S1, Table S2

# loop through and take 100 samples of 126 random genes
# we use 126 because this is the number of nuOXPHOS genes that we previously correlated with the mtOXPHOS genes
# with each random gene sample, loop through the tissues and calculate correlations

# create list to deposit results
# list contains a sublist for each tissue to contain correlation results

correlate.mtOXPHOS.with.random <- function(tissuelist, 
                                           iterations, 
                                           method = c("spearman", "pearson"), 
                                           stat = c("rho", "r"),
                                           graphics_subfolder = "",
                                           filename = "correlations-mtOXPHOS-random-nuclear",
                                           lookuptable = LUT){
  
  method <- match.arg(method)
  stat <- match.arg(stat)
  
  mtOXPHOS_random_corr_list <- list()
  
  # now the loop in the body of the code
  for (i in 1:iterations){

    message(paste0("Starting iteration #", i))
    
    nuclear_sample <- sample(nuclear_gene_expressed, size = 126)
    
    mtOXPHOS_temprandom_residuals_list <- lapply(tissuelist, function(x){apply.lm.to.counts(x[c(mtOXPHOS, nuclear_sample), ], lookuptable = lookuptable)})
    
    mtOXPHOS_tempcorr_list <-   lapply(mtOXPHOS_temprandom_residuals_list,
                                          function(x){
                                            find.all.correlations(x, 
                                                                  method = method, 
                                                                  stat = stat)
                                          })
    
    mtOXPHOS_random_corr_list[[i]] <- mtOXPHOS_tempcorr_list
    
  } # finished looping through iterations
  
  message("Compiling data and producing plot")
  
  tissue_mean_corr_df <- as.data.frame(matrix(nrow = length(tissues), ncol = 3))
  colnames(tissue_mean_corr_df) <- c("tissue", "total_mt_expr", "no_samples")
  
  tissue_mean_corr_df[, "tissue"] <- tissues
  
  for(j in 1:length(tissues)){
    
    sampleIDs <- lookuptable[lookuptable$Tissue == tissues[j], "SAMPID"]
    tissue_mean_corr_df[j, "total_mt_expr"] <- mean(colSums(TPMdata[mitogenes$ensembl_gene_id, sampleIDs])) / 1e4
    tissue_mean_corr_df[j, "no_samples"] <- length(sampleIDs)
    
  } # end of loop over tissues
  
  # ensure graphics_subfolder exists
  dir.create(paste0("graphics/", graphics_subfolder), showWarnings = FALSE)
  
  if(method == "spearman"){
    
    # stick the mean of the medians mito-nuclear correlation in the dataframe
    tissue_mean_corr_df[, "mean_median_mtOXPHOSnuclear_spearman"]  <- rowMeans(sapply(mtOXPHOS_random_corr_list, function(k){
      sapply(k, function(x){median(x[(x[, "Gene1"] %in% mtOXPHOS|x[, "Gene2"] %in% mtOXPHOS) 
                                     & 
                                       (!(x[, "Gene1"] %in% mtOXPHOS)|!(x[, "Gene2"] %in% mtOXPHOS)),
                                     paste(stat)])})
    }))
    
    # stick the sd of the medians mito-nuclear correlation in the dataframe
    tissue_mean_corr_df[, "sd_median_mtOXPHOSnuclear_spearman"]  <- rowSds(sapply(mtOXPHOS_random_corr_list, function(k){
      sapply(k, function(x){median(x[(x$Gene1 %in% mtOXPHOS|x$Gene2 %in% mtOXPHOS) 
                                     & 
                                       (!(x$Gene1 %in% mtOXPHOS)|!(x$Gene2 %in% mtOXPHOS)), paste(stat)])})
    }))
    
    # perform linear model to assess R^2 and add to plot
    ln_reg <- lm(mean_median_mtOXPHOSnuclear_spearman ~ total_mt_expr, data = tissue_mean_corr_df)
    sum_ln_reg <- summary(ln_reg)
    
    pdf(paste0(file = "graphics/", graphics_subfolder,  "/", filename, ".pdf"))
    
    print(ggplot(data = tissue_mean_corr_df,
                 aes(x = total_mt_expr, 
                     y = mean_median_mtOXPHOSnuclear_spearman)) +
            ylab(paste0("Mean ", stat, " between mtOXPHOS and random nuclear genes")) + 
            xlab("% mitochondrial transcripts") +
            ggtitle("Correlation of mtOXPHOS and random nuclear genes") +
            geom_point() +
            geom_smooth(method = "lm", formula = y~ x, se = FALSE) +
            geom_errorbar(aes(x = total_mt_expr,
                              ymax = mean_median_mtOXPHOSnuclear_spearman + (1.96 * sd_median_mtOXPHOSnuclear_spearman / sqrt(iterations)),
                              ymin = mean_median_mtOXPHOSnuclear_spearman - (1.96 * sd_median_mtOXPHOSnuclear_spearman / sqrt(iterations)))) +
            theme_classic() +
            coord_cartesian(ylim = c(-1, 0.5)) +
            geom_text(x = 60, y = 0.5,
                      label = deparse(bquote(R^2 == .(round(sum_ln_reg$r.squared, 2)))),
                      parse = TRUE) +
            geom_hline(yintercept = 0,
                       linetype = "dashed",
                       color = "grey")
    )
    
    dev.off()
    
  } # end of if(spearman)
  
  
  if(method == "pearson"){
    
    # stick the mean of the means mito-nuclear correlation in the dataframe
    tissue_mean_corr_df[, "mean_mean_mtOXPHOSnuclear_pearson"]  <- rowMeans(sapply(mtOXPHOS_random_corr_list, function(k){
      sapply(k, function(x){mean(x[(x$Gene1 %in% mtOXPHOS|x$Gene2 %in% mtOXPHOS) 
                                      & 
                                        (!(x$Gene1 %in% mtOXPHOS)|!(x$Gene2 %in% mtOXPHOS)), paste(stat)])})
    }))
  
    
    
    # stick the sd of the means mito-nuclear correlation in the dataframe
    tissue_mean_corr_df[, "sd_mean_mtOXPHOSnuclear_pearson"]  <- rowSds(sapply(mtOXPHOS_random_corr_list, function(k){
      sapply(k, function(x){mean(x[(x$Gene1 %in% mtOXPHOS|x$Gene2 %in% mtOXPHOS) 
                                   & 
                                     (!(x$Gene1 %in% mtOXPHOS)|!(x$Gene2 %in% mtOXPHOS)), paste(stat)])})
    }))
    
    # perform linear model to assess R^2 and add to plot
    ln_reg <- lm(mean_mean_mtOXPHOSnuclear_pearson ~ total_mt_expr, data = tissue_mean_corr_df)
    sum_ln_reg <- summary(ln_reg)
    
    pdf(paste0(file = "graphics/", graphics_subfolder,  "/", filename, ".pdf"))
    
    print(ggplot(data = tissue_mean_corr_df,
                 aes(x = total_mt_expr, 
                     y = mean_mean_mtOXPHOSnuclear_pearson)) +
            ylab(paste0("Mean ", stat, " between mtOXPHOS and random nuclear genes")) + 
            xlab("% mitochondrial transcripts") +
            ggtitle("Correlation of mtOXPHOS and random nuclear genes") +
            geom_point() +
            geom_smooth(method = "lm", formula = y~ x, se = FALSE) +
            geom_errorbar(aes(x = total_mt_expr,
                              ymax = mean_mean_mtOXPHOSnuclear_pearson + (1.96 * sd_mean_mtOXPHOSnuclear_pearson / sqrt(iterations)),
                              ymin = mean_mean_mtOXPHOSnuclear_pearson - (1.96 * sd_mean_mtOXPHOSnuclear_pearson / sqrt(iterations)))) +
            theme_classic() +
            coord_cartesian(ylim = c(-1, 0.5)) +
            geom_text(x = 60, y = 0.5,
                      label = deparse(bquote(R^2 == .(round(sum_ln_reg$r.squared, 2)))),
                      parse = TRUE) +
            geom_hline(yintercept = 0,
                       linetype = "dashed",
                       color = "grey")
    )
    
    dev.off()
    
  } # end of if(pearson)
  
  output_list <- list(mtOXPHOS_random_corr_list, tissue_mean_corr_df)
  names(output_list) <- c("mito_random_correlations_by_iteration", "tissue_summary_df")
  
  return(output_list)
  
}

test.tissue.mtOXPHOSnuOXPHOS.against.mtOXPHOSrandom <- function(mtOXPHOSrandomnuclear_corr, 
                                                                observedmtnuOXPHOScorr,
                                                                tissues = tissues,
                                                                stat = c("rho", "r")){
  
  stat = match.arg(stat)
  
  output_df <- data.frame(matrix(nrow = length(tissues), ncol = 7))
  colnames(output_df) <- c("tissue", "mtnuOXPHOSmeanavg", "mtOXPHOSrandommeanavg", "mtOXPHOSrandomsd", "Shap_wilk_adj.p", "z_stat", "adj.p")
  
  output_df[, "tissue"] <- tissues
  output_df[, "mtnuOXPHOSmeanavg"] <- observedmtnuOXPHOScorr
  
  # this pulls out a matrix with tissues as columns and iterations as rows, with each median for mtOXPHOS-random nuclear correlations
  
  mtOXPHOSrand_dist <- sapply(1:length(tissues), function(i){
    sapply(mtOXPHOSrandomnuclear_corr, function(x){
      
      median(x[[i]][(x[[i]][, "Gene1"] %in% mtOXPHOS | x[[i]][, "Gene2"] %in% mtOXPHOS) & (!(x[[i]][, "Gene1"] %in% mtOXPHOS) | !(x[[i]][, "Gene2"] %in% mtOXPHOS)),
                    paste(stat)])
      
    })
  })
  
  colnames(mtOXPHOSrand_dist) <- tissues
  
  output_df[, "mtOXPHOSrandommeanavg"] <- rowMeans(t(mtOXPHOSrand_dist))
  output_df[, "mtOXPHOSrandomsd"] <- rowSds(t(mtOXPHOSrand_dist))
  
  # check assumption that distibutions of medians will be normal (according to CLT)
  output_df[, "Shap_wilk_adj.p"] <- p.adjust(apply(mtOXPHOSrand_dist, 2, function(c){
    shapiro.test(c)$p.value
  }), method = "BH")
  
  #calculate parameters of distribution
  mu_dist <- rowMeans(t(mtOXPHOSrand_dist))
  sd_dist <- rowSds(t(mtOXPHOSrand_dist))
  
  output_df[, "z_stat"] <- z_stat <- (observedmtnuOXPHOScorr - mu_dist) / sd_dist
  
  p_value <- 2 * pnorm(-abs(z_stat))
  
  output_df[, "adj.p"] <- p.adjust(p_value, method = "BH")
  
  output_df[, "sd"] <- 
  
  return(output_df)
  
}

dir.create("output/mtOXPHOS_randomnuclear", showWarnings = FALSE)

# run code for Spearman's for MOR, TMM, TPM and TPM (mito excluded for nuclear genes)
MOR_mtOXPHOS_randomnuclear <- correlate.mtOXPHOS.with.random(tissuelist = MOR_tissue_list,
                                                             iterations = 100,
                                                             method = "spearman",
                                                             stat = "rho",
                                                             graphics_subfolder = "mtOXPHOS_randomnuclear",
                                                             filename = "MOR-mtOXPHOS-randomnuclear-by-mito-spearman")

saveRDS(MOR_mtOXPHOS_randomnuclear, "output/mtOXPHOS_randomnuclear/MOR_mtOXPHOS_randomnuclear_spearman.rds")
# MOR_mtOXPHOS_randomnuclear <- readRDS("output/mtOXPHOS_randomnuclear/MOR_mtOXPHOS_randomnuclear_spearman.rds")
# MOR_tissue_correlations <- readRDS("output/MOR-OXPHOS-correlation-by-tissue.rds")

MOR_mtnuOXPHOS_vsrandom_test <- test.tissue.mtOXPHOSnuOXPHOS.against.mtOXPHOSrandom(mtOXPHOSrandomnuclear_corr = MOR_mtOXPHOS_randomnuclear$mito_random_correlations_by_iteration,
                                                                                    observedmtnuOXPHOScorr = MOR_tissue_correlations$median_mtnuOXPHOS_spearman,
                                                                                    tissues = MOR_tissue_correlations$tissue)

write.table(MOR_mtnuOXPHOS_vsrandom_test, "output/MOR_mtnuOXPHOS_vsrandom_test.txt", sep = "\t")
MOR_mtnuOXPHOS_vsrandom_test <- read.table("output/MOR_mtnuOXPHOS_vsrandom_test.txt", sep = "\t")

MOR_bootstrap_sds <- apply(GTEX_mtnuOXPHOSmedian_bootstrap, 2, sd)
MOR_bootstrap_empiricalp <- apply(GTEX_mtnuOXPHOSmedian_bootstrap, 2, function(x){
  if(mean(x) > 0){
    sum(x < 0) / 1000
  } else {
    sum(x > 0) / 1000
  }
  })

MOR_bootstrap_empiricalpadjust <- p.adjust(MOR_bootstrap_empiricalp)

MOR_mtnuOXPHOS_vsrandom_test <- cbind(MOR_mtnuOXPHOS_vsrandom_test, MOR_bootstrap_sds)
MOR_mtnuOXPHOS_vsrandom_test <- cbind(MOR_mtnuOXPHOS_vsrandom_test, MOR_bootstrap_empiricalpadjust)

tissue_select_vec <- c("Adipose \\- Subcutaneous",
                       "Adipose \\- Visceral \\(Omentum\\)",
                       "Adrenal Gland",
                       "Artery \\- Aorta",
                       "Artery \\- Coronary",
                       "Artery  \\- Tibial",
                       "Brain \\- Amygdala",
                       "Brain \\- Anterior cingulate cortex \\(BA24\\)",
                       "Brain \\- Caudate \\(basal ganglia\\)",
                       "Brain \\- Cerebellar Hemisphere",
                       "Brain \\- Cerebellum",
                       "Brain \\- Cortex",
                       "Brain \\- Frontal Cortex \\(BA9\\)",
                       "Brain \\- Hippocampus",
                       "Brain \\- Hypothalamus",
                       "Brain \\- Nucleus accumbens \\(basal ganglia\\)",
                       "Brain \\- Putamen \\(basal ganglia\\)",
                       "Brain \\- Spinal cord \\(cervical c\\-1\\)",
                       "Brain \\- Substantia nigra",
                       "Breast \\- Mammary Tissue",
                       "Cells \\- Cultured fibroblasts",
                       "Cells \\- EBV-transformed lymphocytes",
                       "Colon \\- Sigmoid",
                       "Colon \\- Transverse",
                       "Esophagus \\- Gastroesophageal Junction",
                       "Esophagus \\- Mucosa",
                       "Esophagus \\- Muscularis",
                       "Heart \\- Atrial Appendage",
                       "Heart \\- Left Ventricle",
                       "Liver",
                       "Lung",
                       "Minor Salivary Gland",
                       "Muscle \\- Skeletal",
                       "Nerve \\- Tibial",
                       "Ovary",
                       "Pancreas",
                       "Pituitary",
                       "Prostate",
                       "Skin \\- Not Sun Exposed \\(Suprapubic\\)",
                       "Skin \\- Sun Exposed \\(Lower leg\\)",
                       "Small Intestine \\- Terminal Ileum",
                       "Spleen",
                       "Stomach",
                       "Testis",
                       "Thyroid",
                       "Uterus",
                       "Vagina",
                       "Whole Blood")

tissue_replace_vec <- c("Adipose (Subcut.)",
                        "Adipose (Visc.)",
                        "Adrenal Gland",
                        "Artery (Aorta)",
                        "Artery (Coronary)",
                        "Artery (Tibial)",
                        "Brain (Amygdala)",
                        "Brain (Ant.cing. cortex)",
                        "Brain (Caudate)",
                        "Brain (Cereb. Hemsph.)",
                        "Brain (Cerebellum)",
                        "Brain (Cortex)",
                        "Brain (Frontal Cortex)",
                        "Brain (Hippocampus)",
                        "Brain (Hypothalamus)",
                        "Brain (Nucl. acc.)",
                        "Brain (Putamen)",
                        "Brain (Spinal cord)",
                        "Brain (Subst. nigra)",
                        "Breast (Mammary)",
                        "Cultured fibroblasts",
                        "EBV-transf. lymphocytes",
                        "Colon (Sigmoid)",
                        "Colon (Transverse)",
                        "Esophagus (Gastr. Junc.)",
                        "Esophagus (Mucosa)",
                        "Esophagus (Muscularis)",
                        "Heart (Atrial Appendage)",
                        "Heart (Left Ventricle)",
                        "Liver",
                        "Lung",
                        "Min. Saliv. Gland",
                        "Muscle (Skeletal)",
                        "Nerve (Tibial)",
                        "Ovary",
                        "Pancreas",
                        "Pituitary",
                        "Prostate",
                        "Skin (Suprapubic)",
                        "Skin (Lower leg)",
                        "Small Intestine",
                        "Spleen",
                        "Stomach",
                        "Testis",
                        "Thyroid",
                        "Uterus",
                        "Vagina",
                        "Whole Blood")

for (i in 1:nrow(MOR_mtnuOXPHOS_vsrandom_test)){
  MOR_mtnuOXPHOS_vsrandom_test$tissue <- str_replace_all(MOR_mtnuOXPHOS_vsrandom_test$tissue, pattern = tissue_select_vec[i], replacement = tissue_replace_vec[i])
}

MOR_mtnuOXPHOS_vsrandom_test$tissue <- factor(MOR_mtnuOXPHOS_vsrandom_test$tissue, levels = MOR_mtnuOXPHOS_vsrandom_test[order(MOR_mtnuOXPHOS_vsrandom_test$mtnuOXPHOSmeanavg), "tissue"])
MOR_mtnuOXPHOS_vsrandom_test[MOR_mtnuOXPHOS_vsrandom_test$MOR_bootstrap_empiricalpadjust < 0.05 & MOR_mtnuOXPHOS_vsrandom_test$mtnuOXPHOSmeanavg < 0, "colour"] <- "blue"
MOR_mtnuOXPHOS_vsrandom_test[MOR_mtnuOXPHOS_vsrandom_test$MOR_bootstrap_empiricalpadjust < 0.05 & MOR_mtnuOXPHOS_vsrandom_test$mtnuOXPHOSmeanavg > 0, "colour"] <- "red"
MOR_mtnuOXPHOS_vsrandom_test[MOR_mtnuOXPHOS_vsrandom_test$MOR_bootstrap_empiricalpadjust > 0.05, "colour"] <- "grey"

pdf("graphics/all-tissues-GTEX-mtOXPHOS-nuOXPHOS.pdf",
    height = 5,
    width = 8)

ggplot(data = MOR_mtnuOXPHOS_vsrandom_test, 
       aes(x = tissue, 
           y = mtnuOXPHOSmeanavg)) +
  geom_col(data = MOR_mtnuOXPHOS_vsrandom_test,
           aes(x = tissue,
               y = mtnuOXPHOSmeanavg), 
           fill = MOR_mtnuOXPHOS_vsrandom_test[ ,"colour"],
           col = "black",
           size = 0.5)+
  geom_errorbar(aes(x = tissue,
                    ymin = mtnuOXPHOSmeanavg - 1.96*MOR_bootstrap_sds,
                    ymax = mtnuOXPHOSmeanavg + 1.96*MOR_bootstrap_sds),
                width = 0.3,
                alpha = 0.8,
                col = "black") +
  geom_hline(yintercept = 0,
             linetype = "dashed",
             color = "black") +
  theme_classic() +
  coord_cartesian(ylim=c(-0.5, 0.5)) +
  theme(axis.title.x = element_blank(),
        axis.title.y = element_blank(),
        axis.text.y = element_blank(),
        axis.text.x =  element_text(angle = 90,
                                    vjust = 1,
                                    hjust = 1,
                                    size = 8,
                                    color = "black"))

dev.off()

pdf("graphics/all-tissues-GTEX-mtOXPHOS-random.pdf",
    height = 3.5,
    width = 7)

ggplot(data = MOR_mtnuOXPHOS_vsrandom_test,
       aes(x = tissue,
           y = mtnuOXPHOSmeanavg)) +
  geom_col(data = MOR_mtnuOXPHOS_vsrandom_test,
           aes(x = tissue,
               y = mtOXPHOSrandommeanavg),
           colour = "black",
           fill = "gray") +
  geom_errorbar(aes(x = tissue,
                    ymin = mtOXPHOSrandommeanavg - mtOXPHOSrandomsd,
                    ymax = mtOXPHOSrandommeanavg + mtOXPHOSrandomsd),
                width = 0.3,
                alpha = 0.8,
                col = "black") +
  geom_hline(yintercept = 0,
             linetype = "dashed",
             color = "black") +
  theme_classic() +
  coord_cartesian(ylim=c(-0.5, 0.5)) +
  theme(axis.title.x = element_blank(),
        axis.title.y = element_blank(),
        axis.text.y = element_blank(),
        axis.text.x =  element_text(angle = 90,
                                    vjust = 1,
                                    hjust = 1,
                                    size = 8,
                                    color = "black"))

dev.off()

# Repeat for TMM

TMM_mtOXPHOS_randomnuclear <- correlate.mtOXPHOS.with.random(tissuelist = TMM_tissue_list,
                                                             iterations = 100,
                                                             method = "spearman",
                                                             stat = "rho",
                                                             graphics_subfolder = "mtOXPHOS_randomnuclear",
                                                             filename = "TMM-mtOXPHOS-randomnuclear-by-mito-spearman")

saveRDS(TMM_mtOXPHOS_randomnuclear, "output/mtOXPHOS_randomnuclear/TMM_mtOXPHOS_randomnuclear_spearman.rds")
# TMM_mtOXPHOS_randomnuclear <- readRDS("output/mtOXPHOS_randomnuclear/TMM_mtOXPHOS_randomnuclear_spearman.rds")
# TMM_tissue_correlations <- readRDS("output/TMM-OXPHOS-correlation-by-tissue.rds")

TMM_mtnuOXPHOS_vsrandom_test <- test.tissue.mtOXPHOSnuOXPHOS.against.mtOXPHOSrandom(mtOXPHOSrandomnuclear_corr = TMM_mtOXPHOS_randomnuclear$mito_random_correlations_by_iteration,
                                                                                    observedmtnuOXPHOScorr = TMM_tissue_correlations$median_mtnuOXPHOS_spearman,
                                                                                    tissues = TMM_tissue_correlations$tissue)

TMM_mtnuOXPHOS_vsrandom_test$tissue <- factor(TMM_mtnuOXPHOS_vsrandom_test$tissue, levels = TMM_mtnuOXPHOS_vsrandom_test[order(TMM_mtnuOXPHOS_vsrandom_test$mtnuOXPHOSmeanavg), "tissue"])

TMM_bootstrap_sds <- apply(GTEX_TMMmtnuOXPHOSmedian_bootstrap, 2, sd)
TMM_bootstrap_empiricalp <- apply(GTEX_TMMmtnuOXPHOSmedian_bootstrap, 2, function(x){
  if(mean(x) > 0){
    sum(x < 0) / 1000
  } else {
    sum(x > 0) / 1000
  }
})

TMM_bootstrap_empiricalpadjust <- p.adjust(TMM_bootstrap_empiricalp)

TMM_mtnuOXPHOS_forplot <- cbind(TMM_tissue_correlations, TMM_bootstrap_sds, TMM_bootstrap_empiricalpadjust)

for (i in 1:nrow(TMM_mtnuOXPHOS_vsrandom_test)){
  TMM_mtnuOXPHOS_vsrandom_test$tissue <- str_replace_all(TMM_mtnuOXPHOS_vsrandom_test$tissue, pattern = tissue_select_vec[i], replacement = tissue_replace_vec[i])
}

for (i in 1:nrow(TMM_mtnuOXPHOS_forplot)){
  TMM_mtnuOXPHOS_forplot$tissue <- str_replace_all(TMM_mtnuOXPHOS_forplot$tissue, pattern = tissue_select_vec[i], replacement = tissue_replace_vec[i])
}

TMM_mtnuOXPHOS_forplot$tissue <- factor(TMM_mtnuOXPHOS_forplot$tissue, levels = TMM_mtnuOXPHOS_forplot[order(TMM_mtnuOXPHOS_forplot$median_mtnuOXPHOS_spearman), "tissue"])
TMM_mtnuOXPHOS_forplot[TMM_mtnuOXPHOS_forplot$TMM_bootstrap_empiricalpadjust < 0.05 & TMM_mtnuOXPHOS_forplot$median_mtnuOXPHOS_spearman < 0, "colour"] <- "blue"
TMM_mtnuOXPHOS_forplot[TMM_mtnuOXPHOS_forplot$TMM_bootstrap_empiricalpadjust < 0.05 & TMM_mtnuOXPHOS_forplot$median_mtnuOXPHOS_spearman > 0, "colour"] <- "red"
TMM_mtnuOXPHOS_forplot[TMM_mtnuOXPHOS_forplot$TMM_bootstrap_empiricalpadjust > 0.05, "colour"] <- "grey"

pdf("graphics/all-tissues-GTEX-mtOXPHOS-nuOXPHOS-TMM.pdf",
    height = 5,
    width = 8)

ggplot(data = TMM_mtnuOXPHOS_forplot, aes(x = tissue, y = median_mtnuOXPHOS_spearman))+
  geom_col(data = TMM_mtnuOXPHOS_forplot, aes(x = tissue, y = median_mtnuOXPHOS_spearman), fill = TMM_mtnuOXPHOS_forplot[ ,"colour"], col = "black", size = 0.5)+
  geom_errorbar(aes(x = tissue, ymin = median_mtnuOXPHOS_spearman - 1.96*TMM_bootstrap_sds, ymax = median_mtnuOXPHOS_spearman + 1.96*TMM_bootstrap_sds), width = 0.3, col = "black")+
  geom_hline(yintercept = 0, linetype = "dashed", color = "black")+
  theme_classic()+
  coord_cartesian(ylim=c(-0.5, 0.5))+
  theme(axis.title.x = element_blank(),
        axis.title.y = element_blank(),
        axis.text.y = element_blank(),
        axis.text.x =  element_text(angle = 90, vjust = 1, hjust = 1, size = 8, color = "black"))

dev.off()

pdf("graphics/all-tissues-GTEX-mtOXPHOS-randomnuclear-TMM.pdf",
    height = 3.5,
    width = 7)

ggplot(data = TMM_mtnuOXPHOS_vsrandom_test, aes(x = tissue, y = mtnuOXPHOSmeanavg))+
  geom_col(data = TMM_mtnuOXPHOS_vsrandom_test, aes(x = tissue, y = mtOXPHOSrandommeanavg), colour = "black", fill = "grey")+
  geom_errorbar(aes(x = tissue, ymin = mtOXPHOSrandommeanavg - mtOXPHOSrandomsd, ymax = mtOXPHOSrandommeanavg + mtOXPHOSrandomsd), width = 0.3, col = "black")+
  geom_hline(yintercept = 0, linetype = "dashed", color = "black")+
  theme_classic()+
  coord_cartesian(ylim=c(-0.5, 0.5))+
  theme(axis.title.x = element_blank(),
        axis.title.y = element_blank(),
        axis.text.y = element_blank(),
        axis.text.x =  element_text(angle = 90, vjust = 1, hjust = 1, size = 8, color = "black"))

dev.off()

TPM_mtOXPHOS_randomnuclear <- correlate.mtOXPHOS.with.random(tissuelist = TPM_tissue_list,
                                                             iterations = 3,
                                                             method = "spearman",
                                                             stat = "rho",
                                                             graphics_subfolder = "mtOXPHOS_randomnuclear",
                                                             filename = "TPM-mtOXPHOS-randomnuclear-by-mito-spearman")

saveRDS(TPM_mtOXPHOS_randomnuclear, "output/mtOXPHOS_randomnuclear/TPM_mtOXPHOS_randomnuclear_spearman.rds")

TPMnomito_mtOXPHOS_randomnuclear <- correlate.mtOXPHOS.with.random(tissuelist = TPMnomito_tissue_list,
                                                             iterations = 100,
                                                             method = "spearman",
                                                             stat = "rho",
                                                             graphics_subfolder = "mtOXPHOS_randomnuclear",
                                                             filename = "TPMnomito-mtOXPHOS-randomnuclear-by-mito-spearman")

saveRDS(TPMnomito_mtOXPHOS_randomnuclear, "output/mtOXPHOS_randomnuclear/TPMnomito_mtOXPHOS_randomnuclear_spearman.rds")

UQ_tissue_list <- readRDS("output/UQ-normalisation-by-tissue-list.rds")
UQ_mtOXPHOS_randomnuclear <- correlate.mtOXPHOS.with.random(tissuelist = UQ_tissue_list,
                                                                   iterations = 3,
                                                                   method = "spearman",
                                                                   stat = "rho",
                                                                   graphics_subfolder = "mtOXPHOS_randomnuclear",
                                                                   filename = "UQ-mtOXPHOS-randomnuclear-by-mito-spearman")

saveRDS(UQ_mtOXPHOS_randomnuclear, "output/mtOXPHOS_randomnuclear/UQ_mtOXPHOS_randomnuclear_spearman.rds")

# run code for Pearson's for MOR, TMM, TPM, TPM (mito excluded for nuclear genes) and UQ
MOR_mtOXPHOS_randomnuclear_prs <- correlate.mtOXPHOS.with.random(tissuelist = MOR_tissue_list,
                                                             iterations = 10,
                                                             method = "pearson",
                                                             stat = "r",
                                                             graphics_subfolder = "mtOXPHOS_randomnuclear",
                                                             filename = "MOR-mtOXPHOS-randomnuclear-by-mito-pearson")

saveRDS(MOR_mtOXPHOS_randomnuclear_prs, "output/mtOXPHOS_randomnuclear/MOR_mtOXPHOS_randomnuclear_pearson.rds")

TMM_mtOXPHOS_randomnuclear_prs <- correlate.mtOXPHOS.with.random(tissuelist = TMM_tissue_list,
                                                             iterations = 10,
                                                             method = "pearson",
                                                             stat = "r",
                                                             graphics_subfolder = "mtOXPHOS_randomnuclear",
                                                             filename = "TMM-mtOXPHOS-randomnuclear-by-mito-pearson")

saveRDS(TMM_mtOXPHOS_randomnuclear_prs, "output/mtOXPHOS_randomnuclear/TMM_mtOXPHOS_randomnuclear_pearson.rds")

TPM_mtOXPHOS_randomnuclear_prs <- correlate.mtOXPHOS.with.random(tissuelist = TPM_tissue_list,
                                                             iterations = 10,
                                                             method = "pearson",
                                                             stat = "r",
                                                             graphics_subfolder = "mtOXPHOS_randomnuclear",
                                                             filename = "TPM-mtOXPHOS-randomnuclear-by-mito-pearson")

saveRDS(TPM_mtOXPHOS_randomnuclear_prs, "output/mtOXPHOS_randomnuclear/TPM_mtOXPHOS_randomnuclear_pearson.rds")

TPMnomito_mtOXPHOS_randomnuclear_prs <- correlate.mtOXPHOS.with.random(tissuelist = TPMnomito_tissue_list,
                                                                   iterations = 10,
                                                                   method = "pearson",
                                                                   stat = "r",
                                                                   graphics_subfolder = "mtOXPHOS_randomnuclear",
                                                                   filename = "TPMnomito-mtOXPHOS-randomnuclear-by-mito-pearson")

saveRDS(TPMnomito_mtOXPHOS_randomnuclear_prs, "output/mtOXPHOS_randomnuclear/TPMnomito_mtOXPHOS_randomnuclear_pearson.rds")

UQ_mtOXPHOS_randomnuclear_prs <- correlate.mtOXPHOS.with.random(tissuelist = UQ_tissue_list,
                                                                   iterations = 10,
                                                                   method = "pearson",
                                                                   stat = "r",
                                                                   graphics_subfolder = "mtOXPHOS_randomnuclear",
                                                                   filename = "UQ-mtOXPHOS-randomnuclear-by-mito-pearson")

saveRDS(UQ_mtOXPHOS_randomnuclear_prs, "output/mtOXPHOS_randomnuclear/UQ_mtOXPHOS_randomnuclear_pearson.rds")

# for MOR normalisation, plot mtOXPHOS-nuOXPHOS values together on the plot with the mtOXPHOS-random values

comboplot_temp <- merge(MOR_tissue_correlations[, c("tissue", "total_mt_expr", "median_mtnuOXPHOS_spearman", "Q1_mtnuOXPHOS_spearman", "Q3_mtnuOXPHOS_spearman")], 
                        MOR_mtOXPHOS_randomnuclear$tissue_summary_df[, c("tissue", "mean_median_mtOXPHOSnuclear_spearman", "sd_median_mtOXPHOSnuclear_spearman")])

pdf(file = "graphics/mtOXPHOS_randomnuclear/MOR-mt-nuOXPHOS_with_mtOXPHOS-nurandom.pdf")

print(ggplot(data = comboplot_temp,
             aes(x = total_mt_expr,
                 y = median_mtnuOXPHOS_spearman)) +
        ylab("Correlation") + 
        xlab("% mitochondrial transcripts") +
        ggtitle("mt-nuOXPHOS correlations with mtOXPHOS-random expectation") +
        geom_point(aes(x = total_mt_expr,
                       y = mean_median_mtOXPHOSnuclear_spearman),
                   col = "gray") +
        geom_point() +
        geom_errorbar(aes(x = total_mt_expr,
                          ymax = mean_median_mtOXPHOSnuclear_spearman + (1.96 * sd_median_mtOXPHOSnuclear_spearman / sqrt(iterations)),
                          ymin = mean_median_mtOXPHOSnuclear_spearman - (1.96 * sd_median_mtOXPHOSnuclear_spearman / sqrt(iterations))),
                      col = "gray") +
        geom_errorbar(aes(x = total_mt_expr,
                          ymax = Q3_mtnuOXPHOS_spearman,
                          ymin = Q1_mtnuOXPHOS_spearman)) +
        theme_classic() +
        coord_cartesian(ylim = c(-1, 0.5)) +
        geom_hline(yintercept = 0,
                   linetype = "dashed",
                   color = "grey")
)

dev.off()

# for TPM normalisation, plot mtOXPHOS-nuOXPHOS values together on the plot with the mtOXPHOS-random values

comboplot_temp <- merge(TPM_tissue_correlations[, c("tissue", "total_mt_expr", "median_mtnuOXPHOS_spearman", "Q1_mtnuOXPHOS_spearman", "Q3_mtnuOXPHOS_spearman")], 
                        TPM_mtOXPHOS_randomnuclear$tissue_summary_df[, c("tissue", "mean_median_mtOXPHOSnuclear_spearman", "sd_median_mtOXPHOSnuclear_spearman")])

pdf(file = "graphics/mtOXPHOS_randomnuclear/TPM-mt-nuOXPHOS_with_mtOXPHOS-nurandom.pdf")

print(ggplot(data = comboplot_temp,
             aes(x = total_mt_expr,
                 y = median_mtnuOXPHOS_spearman)) +
        ylab("Correlation") + 
        xlab("% mitochondrial transcripts") +
        ggtitle("mt-nuOXPHOS correlations with mtOXPHOS-random expectation") +
        geom_point(aes(x = total_mt_expr,
                       y = mean_median_mtOXPHOSnuclear_spearman),
                   col = "gray") +
        geom_point() +
        geom_errorbar(aes(x = total_mt_expr,
                          ymax = mean_median_mtOXPHOSnuclear_spearman + (1.96 * sd_median_mtOXPHOSnuclear_spearman / sqrt(iterations)),
                          ymin = mean_median_mtOXPHOSnuclear_spearman - (1.96 * sd_median_mtOXPHOSnuclear_spearman / sqrt(iterations))),
                      col = "gray") +
        geom_errorbar(aes(x = total_mt_expr,
                          ymax = Q3_mtnuOXPHOS_spearman,
                          ymin = Q1_mtnuOXPHOS_spearman)) +
        theme_classic() +
        coord_cartesian(ylim = c(-1, 1)) +
        geom_hline(yintercept = 0,
                   linetype = "dashed",
                   color = "grey")
)

dev.off()

# for TMM normalisation, plot mtOXPHOS-nuOXPHOS values together on the plot with the mtOXPHOS-random values

comboplot_temp <- merge(TMM_tissue_correlations[, c("tissue", "total_mt_expr", "median_mtnuOXPHOS_spearman", "Q1_mtnuOXPHOS_spearman", "Q3_mtnuOXPHOS_spearman")], 
                        TMM_mtOXPHOS_randomnuclear$df_for_plot[, c("tissue", "meancorr_across_iterations", "sdcorr_across_iterations")])

pdf(file = "graphics/mtOXPHOS_randomnuclear/TMM-mt-nuOXPHOS_with_mtOXPHOS-nurandom.pdf")

print(ggplot(data = comboplot_temp, 
             aes(x = total_mt_expr,
                 y = median_mtnuOXPHOS_spearman)) +
        ylab("Correlation") + 
        xlab("% mitochondrial transcripts") +
        ggtitle("mt-nuOXPHOS correlations with mtOXPHOS-random expectation") +
        geom_point(aes(x = total_mt_expr, 
                       y = meancorr_across_iterations),
                   col = "gray") +
        geom_point() +
        geom_errorbar(aes(x = total_mt_expr, 
                          ymax = meancorr_across_iterations + (1.96 * sdcorr_across_iterations / sqrt(iterations)),
                          ymin = meancorr_across_iterations - (1.96 * sdcorr_across_iterations / sqrt(iterations))),
                      col = "gray") +
        geom_errorbar(aes(x = total_mt_expr,
                          ymax = Q3_mtnuOXPHOS_spearman,
                          ymin = Q1_mtnuOXPHOS_spearman)) +
        theme_classic() +
        coord_cartesian(ylim = c(-1, 0.5)) +
        geom_hline(yintercept = 0,
                   linetype = "dashed",
                   color = "grey")
)

dev.off()

#### CORRELATIONS OF RANDOM NUCLEAR GENES ####
# corresponding to Fig 1e, Fig S1d

correlate.random.nuclear.genes <- function(tissuelist,
                                           iterations,
                                           method = c("spearman", "pearson"),
                                           stat = c("rho", "r"),
                                           graphics_subfolder = "",
                                           filename = "random-nuclear-correlations",
                                           lookuptable = LUT) {

method <- match.arg(method)
stat <- match.arg(stat)  
  
correlations_list <- list()

for(i in 1:iterations){
  
  message(paste0("Starting iteration #", i))
  
  nuclear_sample <- sample(nuclear_gene_expressed, 100)
  
  nuclear_temprandom_residuals_list <- lapply(tissuelist, 
                                              function(x){
                                                apply.lm.to.counts(x[nuclear_sample, ], lookuptable = lookuptable)
                                                })
  
  random_corr_list <- lapply(nuclear_temprandom_residuals_list,
                 function(x){
                   find.all.correlations(x[nuclear_sample, ], 
                                         method = method, 
                                         stat = stat)
                 })
  
  correlations_list[[i]] <- random_corr_list
  
} # end of loop over iterations

  message("Compiling data and producing plot")

  tissue_mean_corr_df <- as.data.frame(matrix(nrow = length(tissues), ncol = 3))
  colnames(tissue_mean_corr_df) <- c("tissue", "total_mt_expr", "no_samples")
  
  tissue_mean_corr_df[, "tissue"] <- tissues
  
  for(j in 1:length(tissues)){
    
    sampleIDs <- lookuptable[lookuptable$Tissue == tissues[j], "SAMPID"]
    tissue_mean_corr_df[j, "total_mt_expr"] <- mean(colSums(TPMdata[mitogenes$ensembl_gene_id, sampleIDs])) / 1e4
    tissue_mean_corr_df[j, "no_samples"] <- length(sampleIDs)
    
  } # end of loop over tissues
    
  # ensure graphics_subfolder exists
  dir.create(paste0("graphics/", graphics_subfolder), showWarnings = FALSE)
  
  if(method == "spearman"){
    
    # stick the mean of the medians mito-nuclear correlation in the dataframe
    tissue_mean_corr_df[, "mean_median_randomnuclear_spearman"]  <- rowMeans(sapply(correlations_list, function(k){
      sapply(k, function(x){median(x[, paste(stat)])})
    }))
      
    # stick the sd of the medians mito-nuclear correlation in the dataframe
    tissue_mean_corr_df[, "sd_median_randomnuclear_spearman"]  <- rowSds(sapply(correlations_list, function(k){
      sapply(k, function(x){median(x[, paste(stat)])})
    }))
    
    # perform linear model to assess R^2 for adding to plot
    ln_reg <- lm(mean_median_randomnuclear_spearman ~ total_mt_expr, data = tissue_mean_corr_df)
    sum_ln_reg <- summary(ln_reg)
    
    pdf(paste0(file = "graphics/", graphics_subfolder,  "/", filename, ".pdf"))
    
    # plot mean correlation between random nuclear genes against total mitochondrial expression
    print(ggplot(data = tissue_mean_corr_df,
                 aes(x = total_mt_expr, 
                     y = mean_median_randomnuclear_spearman)) +
            ylab(paste0("Mean ", stat, " between random nuclear genes")) + 
            xlab("% mitochondrial transcripts") +
            ggtitle("Mean random nuclear correlation") +
            geom_point() +
            geom_smooth(method = "lm", formula = y~ x, se = FALSE) +
            geom_errorbar(aes(x = total_mt_expr,
                              ymax = mean_median_randomnuclear_spearman + (1.96 * sd_median_randomnuclear_spearman / sqrt(iterations)),
                              ymin = mean_median_randomnuclear_spearman - (1.96 * sd_median_randomnuclear_spearman / sqrt(iterations)))) +
            theme_classic() +
            coord_cartesian(ylim = c(-0.25, 1)) +
            geom_text(x = 60, y = 0.5,
                      label = deparse(bquote(R^2 == .(round(sum_ln_reg$r.squared, 2)))),
                      parse = TRUE) +
            geom_hline(yintercept = 0,
                       linetype = "dashed",
                       color = "grey")
    )
    
    dev.off()
    
    } # end of if(spearman)
  
    if(method == "pearson"){
      
      # stick the mean of the medians mito-nuclear correlation in the dataframe
      tissue_mean_corr_df[, "mean_mean_randomnuclear_pearson"]  <- rowMeans(sapply(correlations_list, function(k){
        sapply(k, function(x){mean(x[, paste(stat)])})
      }))
      
      # stick the sd of the medians mito-nuclear correlation in the dataframe
      tissue_mean_corr_df[, "sd_mean_randomnuclear_pearson"]  <- rowSds(sapply(correlations_list, function(k){
        sapply(k, function(x){mean(x[, paste(stat)])})
      }))
      
      # perform linear model to assess R^2 for adding to plot
      ln_reg <- lm(mean_mean_randomnuclear_pearson ~ total_mt_expr, data = tissue_mean_corr_df)
      sum_ln_reg <- summary(ln_reg)
      
      pdf(paste0(file = "graphics/", graphics_subfolder,  "/", filename, ".pdf"))
      
      # plot mean correlation between random nuclear genes against total mitochondrial expression
      print(ggplot(data = tissue_mean_corr_df, aes(x = total_mt_expr, y = mean_mean_randomnuclear_pearson)) +
              ylab(paste0("Mean ", stat, " between random nuclear genes")) + 
              xlab("% mitochondrial transcripts") +
              ggtitle("Mean random nuclear correlation") +
              geom_point() +
              geom_smooth(method = "lm",
                          formula = y~ x,
                          se = FALSE) +
              geom_errorbar(aes(x = total_mt_expr, 
                                ymax = mean_mean_randomnuclear_pearson + (1.96 * sd_mean_randomnuclear_pearson / sqrt(iterations)), 
                                ymin = mean_mean_randomnuclear_pearson - (1.96 * sd_mean_randomnuclear_pearson / sqrt(iterations)))) +
              theme_classic() +
              coord_cartesian(ylim = c(-0.25, 0.8)) +
              geom_text(x = 60,
                        y = 0.5,
                        label = deparse(bquote(R^2 == .(round(sum_ln_reg$r.squared, 2)))),
                        parse = TRUE) +
              geom_hline(yintercept = 0,
                         linetype = "dashed",
                         color = "grey")
      )
      
      dev.off()
      
    } # end of if(pearson)

  # put output objects into list, name and return
  output_list <- list(correlations_list, tissue_mean_corr_df)
  
  names(output_list) <- c("correlations_by_iteration", "tissue_summary_df")
  
  return(output_list)
  
}

dir.create("output/random_nuclear_nuclear", showWarnings = FALSE)

MOR_randomnuclear <- correlate.random.nuclear.genes(tissuelist = MOR_tissue_list,
                                                    iterations = 100,
                                                    method = "spearman",
                                                    stat = "rho",
                                                    graphics_subfolder = "random_nuclear_nuclear",
                                                    filename = "MOR_random_nuclear_nuclear_by_mito_expr")

saveRDS(MOR_randomnuclear, "output/random_nuclear_nuclear/MOR_random_nuclear_nuclear.rds")
# MOR_randomnuclear <- readRDS("output/random_nuclear_nuclear/MOR_random_nuclear_nuclear.rds")

TMM_randomnuclear <- correlate.random.nuclear.genes(tissuelist = TMM_tissue_list,
                                                    iterations = 10,
                                                    method = "spearman",
                                                    stat = "rho",
                                                    graphics_subfolder = "random_nuclear_nuclear",
                                                    filename = "TMM_random_nuclear_nuclear_by_mito_expr")

saveRDS(TMM_randomnuclear, "output/random_nuclear_nuclear/TMM_random_nuclear_nuclear.rds")
# TMM_randomnuclear <- readRDS("output/random_nuclear_nuclear/TMM_random_nuclear_nuclear.rds")

TPM_randomnuclear <- correlate.random.nuclear.genes(tissuelist = TPM_tissue_list,
                               iterations = 10,
                               method = "spearman",
                               stat = "rho",
                               graphics_subfolder = "random_nuclear_nuclear",
                               filename = "TPM_random_nuclear_nuclear_by_mito_expr")

saveRDS(TPM_randomnuclear, "output/random_nuclear_nuclear/TPM_random_nuclear_nuclear.rds")
# TPM_randomnuclear <- readRDS("output/random_nuclear_nuclear/TPM_random_nuclear_nuclear.rds")

# linear model to test if cv has influence
TPM_randomnuclear$tissue_summary_df[, "mito_expr_cv"] <- mito_cv_vec
summary(lm(mean_median_randomnuclear_spearman ~ 
             total_mt_expr + mito_expr_cv + total_mt_expr*mito_expr_cv, 
           data = TPM_randomnuclear$tissue_summary_df))

TPMnomito_randomnuclear <- correlate.random.nuclear.genes(tissuelist = TPMnomito_tissue_list,
                                                    iterations = 100,
                                                    method = "spearman",
                                                    stat = "rho",
                                                    graphics_subfolder = "random_nuclear_nuclear",
                                                    filename = "TPMnomito_random_nuclear_nuclear_by_mito_expr")

saveRDS(TPMnomito_randomnuclear, "output/random_nuclear_nuclear/TPMnomito_random_nuclear_nuclear.rds")
# TPMnomito_randomnuclear <- readRDS("output/random_nuclear_nuclear/TPMnomito_random_nuclear_nuclear.rds")

UQ_randomnuclear <- correlate.random.nuclear.genes(tissuelist = UQ_tissue_list,
                                                          iterations = 100,
                                                          method = "spearman",
                                                          stat = "rho",
                                                          graphics_subfolder = "random_nuclear_nuclear",
                                                          filename = "UQ_random_nuclear_nuclear_by_mito_expr")

saveRDS(UQ_randomnuclear, "output/random_nuclear_nuclear/UQ_random_nuclear_nuclear.rds")
# UQ_randomnuclear <- readRDS("output/random_nuclear_nuclear/UQ_random_nuclear_nuclear.rds")

#now pearson
MOR_randomnuclear_pearson <- correlate.random.nuclear.genes(tissuelist = MOR_tissue_list,
                                                    iterations = 100,
                                                    method = "pearson",
                                                    stat = "r",
                                                    graphics_subfolder = "random_nuclear_nuclear",
                                                    filename = "MOR_random_nuclear_nuclear_by_mito_expr_pearson")

saveRDS(MOR_randomnuclear_pearson, "output/random_nuclear_nuclear/MOR_random_nuclear_nuclear_pearson.rds")

TMM_randomnuclear_pearson <- correlate.random.nuclear.genes(tissuelist = TMM_tissue_list,
                                                    iterations = 100,
                                                    method = "pearson",
                                                    stat = "r",
                                                    graphics_subfolder = "random_nuclear_nuclear",
                                                    filename = "TMM_random_nuclear_nuclear_by_mito_expr_pearson")

saveRDS(TMM_randomnuclear, "output/random_nuclear_nuclear/TMM_random_nuclear_nuclear_pearson.rds")

TPM_randomnuclear_pearson <- correlate.random.nuclear.genes(tissuelist = TPM_tissue_list,
                                                    iterations = 100,
                                                    method = "pearson",
                                                    stat = "r",
                                                    graphics_subfolder = "random_nuclear_nuclear",
                                                    filename = "TPM_random_nuclear_nuclear_by_mito_expr_pearson")

saveRDS(TPM_randomnuclear_pearson, "output/random_nuclear_nuclear/TPM_random_nuclear_nuclear_pearson.rds")

#### QTL normalised random nuclear ####
# confirms that GTEx QTL normalised data are appropriate
# results not displayed in manuscript

  QTLcorrelations_list <- list()
  
  for(i in 1:10){
    
    message(paste0("Starting iteration #", i))
    
    QTLnuclear_sample <- sample(nuclear_gene_expressed, 100)
    
    QTLrandom_corr_list <- lapply(GTEXQTLdatalist,
                               function(x){
                                 find.all.correlations(x[row.names(x) %in% QTLnuclear_sample, ], 
                                                       method = "spearman", 
                                                       stat = "rho")
                               })
    
    QTLcorrelations_list[[i]] <- QTLrandom_corr_list
    
  } # end of loop over iterations
  
  QTLtissue_mean_corr_df <- as.data.frame(matrix(nrow = length(GTEXQTLdatalist), ncol = 1))
  colnames(QTLtissue_mean_corr_df) <- c("tissue")
  
  QTLtissue_mean_corr_df[, "tissue"] <- names(GTEXQTLdatalist)
    
    # stick the mean of the medians mito-nuclear correlation in the dataframe
    QTLtissue_mean_corr_df[, "mean_median_randomnuclear_spearman"]  <- rowMeans(sapply(QTLcorrelations_list, function(k){
      sapply(k, function(x){median(x[, paste(stat)])})
    }))
    
    # stick the sd of the medians mito-nuclear correlation in the dataframe
    QTLtissue_mean_corr_df[, "sd_median_randomnuclear_spearman"]  <- rowSds(sapply(QTLcorrelations_list, function(k){
      sapply(k, function(x){median(x[, paste(stat)])})
    }))
    
    splitnames <- str_split(QTLtissue_mean_corr_df$tissue, "_")
    pattern <- sapply(splitnames, function(x){
      x[length(x)]
    })
    
   # the tissues in the QTL set are lacking both Adipose tissue sets and have Kidney in addition
    
    mt_expr <- MOR_tissue_correlations[order(MOR_tissue_correlations[, "tissue"]), c("tissue", "total_mt_expr")]
    mt_expr <- mt_expr[3:nrow(mt_expr), ]
    
    QTLtissue_mean_corr_df <- QTLtissue_mean_corr_df[!(QTLtissue_mean_corr_df$tissue == "Kidney_Cortex"), ]
    QTLtissue_mean_corr_df <- QTLtissue_mean_corr_df[order(QTLtissue_mean_corr_df$tissue), ]
    
    QTLtissue_mean_corr_df <- cbind(QTLtissue_mean_corr_df, mt_expr)
    QTLtissue_mean_corr_df <- QTLtissue_mean_corr_df[, 2:ncol(QTLtissue_mean_corr_df)]
    
    QTLln_reg <- lm(mean_median_randomnuclear_spearman ~ total_mt_expr, data = QTLtissue_mean_corr_df)
    sum_QTLln_reg <- summary(QTLln_reg)
    
    pdf(paste0(file = "graphics/eQTL.pdf"))
    
    # plot mean correlation between random nuclear genes against total mitochondrial expression
    print(ggplot(data = QTLtissue_mean_corr_df,
                 aes(x = total_mt_expr, 
                     y = mean_median_randomnuclear_spearman)) +
            ylab(paste0("Mean ", stat, " between random nuclear genes")) + 
            xlab("% mitochondrial transcripts") +
            ggtitle("Mean random nuclear correlation") +
            geom_point() +
            geom_smooth(method = "lm", formula = y~ x, se = FALSE) +
            geom_errorbar(aes(x = total_mt_expr,
                              ymax = mean_median_randomnuclear_spearman + (1.96 * sd_median_randomnuclear_spearman / sqrt(iterations)),
                              ymin = mean_median_randomnuclear_spearman - (1.96 * sd_median_randomnuclear_spearman / sqrt(iterations)))) +
            theme_classic() +
            coord_cartesian(ylim = c(-0.25, 1)) +
            geom_text(x = 60, y = 0.5,
                      label = deparse(bquote(R^2 == .(round(sum_QTLln_reg$r.squared, 2)))),
                      parse = TRUE) +
            geom_hline(yintercept = 0,
                       linetype = "dashed",
                       color = "grey")
    )
    
    dev.off()
  
  # put output objects into list, name and return
  QTLoutput_list <- list(QTLcorrelations_list, QTLtissue_mean_corr_df)
  
  names(QTLoutput_list) <- c("correlations_by_iteration", "tissue_summary_df")

#### linear model to test if cv has influence ####

TPM_randomnuclear$tissue_summary_df[, "mito_expr_cv"] <- mito_cv_vec
summary(lm(mean_median_randomnuclear_pearson ~ total_mt_expr + mito_expr_cv + total_mt_expr*mito_expr_cv, data = TPM_randomnuclear$tissue_summary_df))

TPMnomito_randomnuclear_pearson <- correlate.random.nuclear.genes(tissuelist = TPMnomito_tissue_list,
                                                          iterations = 10,
                                                          method = "pearson",
                                                          stat = "r",
                                                          graphics_subfolder = "random_nuclear_nuclear",
                                                          filename = "TPMnomito_random_nuclear_nuclear_by_mito_expr_pearson")

saveRDS(TPMnomito_randomnuclear_pearson, "output/random_nuclear_nuclear/TPMnomito_random_nuclear_nuclear_pearson.rds")

UQ_randomnuclear_pearson <- correlate.random.nuclear.genes(tissuelist = UQ_tissue_list,
                                                   iterations = 10,
                                                   method = "pearson",
                                                   stat = "r",
                                                   graphics_subfolder = "random_nuclear_nuclear",
                                                   filename = "UQ_random_nuclear_nuclear_by_mito_expr_pearson")

saveRDS(UQ_randomnuclear_pearson, "output/random_nuclear_nuclear/UQ_random_nuclear_nuclear_pearson.rds")

#### BIMODALITY OF TESTIS CORRELATIONS ####
# corresponds to Fig S3a, Fig S3b

MOR_randomnuclear100 <- correlate.random.nuclear.genes(tissuelist = MOR_tissue_list,
                                                    iterations = 100,
                                                    method = "spearman",
                                                    stat = "rho",
                                                    graphics_subfolder = "random_nuclear_nuclear",
                                                    filename = "MOR_random_nuclear_nuclear_by_mito_expr")

dir.create("graphics/modality_histograms")

for(i in 1:length(MOR_randomnuclear100$correlations_by_iteration[[1]])){

  pdf(paste0("graphics/modality_histograms/", names(MOR_randomnuclear100$correlations_by_iteration[[1]])[i], ".pdf"),
             width = 2.2,
             height = 2.2)
  print(ggplot()+
          geom_density(data = MOR_randomnuclear100$correlations_by_iteration[[1]][[i]], aes(rho), size = 0.2) +
          geom_density(data = MOR_randomnuclear100$correlations_by_iteration[[2]][[i]], aes(rho), size = 0.2) +
    geom_density(data = MOR_randomnuclear100$correlations_by_iteration[[3]][[i]], aes(rho), size = 0.2) +
      geom_density(data = MOR_randomnuclear100$correlations_by_iteration[[4]][[i]], aes(rho), size = 0.2) +
      geom_density(data = MOR_randomnuclear100$correlations_by_iteration[[5]][[i]], aes(rho), size = 0.2) +
      geom_density(data = MOR_randomnuclear100$correlations_by_iteration[[6]][[i]], aes(rho), size = 0.2) +
      geom_density(data = MOR_randomnuclear100$correlations_by_iteration[[7]][[i]], aes(rho), size = 0.2) +
      geom_density(data = MOR_randomnuclear100$correlations_by_iteration[[8]][[i]], aes(rho), size = 0.2) +
      geom_density(data = MOR_randomnuclear100$correlations_by_iteration[[9]][[i]], aes(rho), size = 0.2) +
      geom_density(data = MOR_randomnuclear100$correlations_by_iteration[[10]][[i]], aes(rho), size = 0.2) +
      
    theme_classic() + 
    coord_cartesian(xlim = c(-1,1), ylim = c(0, 2.5))+
      theme(axis.title.x = element_blank(),
            axis.text.x = element_blank(),
            axis.title.y = element_blank(),
            axis.text.y = element_blank(),
            axis.ticks.y = element_blank()))
    
  dev.off()
}

tissue_diptests <- sapply(1:length(MOR_randomnuclear100$correlations_by_iteration[[1]]), function(i){
  sapply(MOR_randomnuclear100$correlations_by_iteration, function(x){
    diptest::dip.test(x[[i]][, 3])$p.value
  })
})

colnames(tissue_diptests) <- names(MOR_randomnuclear100$correlations_by_iteration[[1]])

tissue_diptests_adj <- apply(tissue_diptests, 2, p.adjust)

# heatmaps for random nuclear genes

sample_for_random_heatmap <- sample(nuclear_gene_expressed, 100)

nuclear_random_residuals_list <- lapply(MOR_tissue_list, 
                                            function(x){
                                              apply.lm.to.counts(x[sample_for_random_heatmap, ], lookuptable = LUT)
                                            })

random_corr_heatmap_list <- lapply(nuclear_random_residuals_list,
                           function(x){
                             find.all.correlations(x[sample_for_random_heatmap, ], 
                                                   method = "spearman", 
                                                   stat = "rho")
                           })

random_clustermat_list <- lapply(random_corr_heatmap_list,
                                 function(x){
                                   build.matrix.4.cluster(corr_df = x,
                                                  stat = "rho")
                                 })

dir.create("graphics/random-nuclear-heatmaps")
mainpal <- (colorRampPalette(c("blue", "white", "red"))(100))

for(i in 1:length(random_clustermat_list)){
  
pdf(paste0(file = "graphics/random-nuclear-heatmaps/", paste0(unlist(str_split(names(random_clustermat_list)[i], pattern = " ")), collapse = ""), ".pdf"))

heatmap.2(random_clustermat_list[[i]],
          main = paste0(names(random_clustermat_list)[i]),
          symkey = FALSE,
          symbreaks = TRUE,
          breaks = NULL,
          density.info = "none",
          trace = "none",
          margins = c(12, 9),
          col = mainpal,
          Rowv = TRUE,
          Colv = "Rowv",
          dendrogram = "none",
          cexRow = 0.2,
          cexCol = 0.2)

dev.off()

}

#### EXAMINATION OF OTHER LIBRARY COMPOSITION BIASES ####

# do top10 non-mito gene expression for all tissues

tissue_top10_df <- data.frame(matrix(nrow = length(tissues), ncol  = 23))
colnames(tissue_top10_df) <- c("tissue", "samples", "top10_expression", paste0("gene", 1:10, "_expr"), paste0("gene", 1:10, "_name"))

tissue_top10_df$tissue <- tissues

for(i in 1:length(tissues)){
  
  sampleIDs <- LUT[LUT$Tissue == tissues[i], "SAMPID"]
  
  tissue_top10_df[i, "samples"] <- length(sampleIDs)
  
  tissuedata <- TPMdata[, sampleIDs]
  
  gene_means <- rowMeans(tissuedata)
  
  # here I will cut out the mito genes
  gene_means <- gene_means[(!names(gene_means) %in% mitogenes$ensembl_gene_id)]
  
  gene_means <- gene_means[order(gene_means, decreasing = TRUE)]
  
  tissue_top10_df[i, "top10_expression"] <- sum(gene_means[1:10])
  
  for(j in 1:10){
    
    tissue_top10_df[i, paste0("gene", j, "_expr")] <- gene_means[j]
    tissue_top10_df[i, paste0("gene", j, "_name")] <- names(gene_means)[j]
    
  }
  
}

# get names for top 10 genes
top10genes <- unique(unlist(tissue_top10_df[, 14:23]))

top10names <- getBM(attributes = c("ensembl_gene_id", "hgnc_symbol", "description"),
                    filters = "ensembl_gene_id",
                    values = top10genes,
                    mart = ensembl)

tissue_top10_df$gene1_name <- top10names[match(tissue_top10_df$gene1_name, top10names$ensembl_gene_id), "hgnc_symbol"]
tissue_top10_df$gene2_name <- top10names[match(tissue_top10_df$gene2_name, top10names$ensembl_gene_id), "hgnc_symbol"]
tissue_top10_df$gene3_name <- top10names[match(tissue_top10_df$gene3_name, top10names$ensembl_gene_id), "hgnc_symbol"]
tissue_top10_df$gene4_name <- top10names[match(tissue_top10_df$gene4_name, top10names$ensembl_gene_id), "hgnc_symbol"]
tissue_top10_df$gene5_name <- top10names[match(tissue_top10_df$gene5_name, top10names$ensembl_gene_id), "hgnc_symbol"]
tissue_top10_df$gene6_name <- top10names[match(tissue_top10_df$gene6_name, top10names$ensembl_gene_id), "hgnc_symbol"]
tissue_top10_df$gene7_name <- top10names[match(tissue_top10_df$gene7_name, top10names$ensembl_gene_id), "hgnc_symbol"]
tissue_top10_df$gene8_name <- top10names[match(tissue_top10_df$gene8_name, top10names$ensembl_gene_id), "hgnc_symbol"]
tissue_top10_df$gene9_name <- top10names[match(tissue_top10_df$gene9_name, top10names$ensembl_gene_id), "hgnc_symbol"]
tissue_top10_df$gene10_name <- top10names[match(tissue_top10_df$gene10_name, top10names$ensembl_gene_id), "hgnc_symbol"]

# total in pancreas from PRSS1 & 2
sum(tissue_top10_df[tissue_top10_df$tissue == "Pancreas", c("gene1_expr", "gene2_expr")])

# total in blood from HBA1&2, HBB, HBD
sum(tissue_top10_df[tissue_top10_df$tissue == "Whole Blood", c("gene1_expr", "gene2_expr", "gene4_expr", "gene5_expr")])

#### EXCLUDE PRSS1 & 2 FROM PANCREAS TPM COUNTS ####
# corresponds to Fig 1f, Fig S1e

pancreas_ids <- LUT[LUT$Tissue == "Pancreas", "SAMPID"]
pancreascounts <- countsdata[, pancreas_ids]

excludedgenes <- top10names[top10names$hgnc_symbol %in% c("PRSS1", "PRSS2"), "ensembl_gene_id"]

samplescale_excl <- colSums(pancreascounts[!(row.names(pancreascounts) %in% excludedgenes), ]) / 1e6

pancreascounts_excl_normal <- t(t(pancreascounts[!(row.names(pancreascounts) %in% excludedgenes), ]) / samplescale_excl)

#restrict to ones with TPM > 5.0

pancreascounts_excl_normal <- pancreascounts_excl_normal[row.names(pancreascounts_excl_normal) %in% nuclear_gene_expressed, ]

pancreas_nuclear_nuclear_noPRSS_spearman <- vector(mode = "numeric", length = 10)
pancreas_nuclear_nuclear_noPRSS_pearson <- vector(mode = "numeric", length = 10)

pancreas_nuclear_nuclear_TPM_spearman <- vector(mode = "numeric", length = 10)
pancreas_nuclear_nuclear_TPM_pearson <- vector(mode = "numeric", length = 10)

for(i in 1:10){

  genesample <- sample(row.names(pancreascounts_excl_normal)[!(row.names(pancreascounts_excl_normal) %in% mitogenes$ensembl_gene_id)], 100)
  
  tempdata <- pancreascounts_excl_normal[genesample, ]
  
  tissuecorr <- find.all.correlations(tempdata, method = "spearman")
  
  pancreas_nuclear_nuclear_noPRSS_spearman[i] <-  median(tissuecorr$rho)
  
  # for this tissue and sample, do pearson and report mean
  tissuecorr <- find.all.correlations(tempdata, method = "pearson")
  
  pancreas_nuclear_nuclear_noPRSS_pearson[i] <-  mean(tissuecorr$rho)
  
  tempTPMdata <- TPMdata[genesample, pancreas_ids]
  
  tissuecorr <- find.all.correlations(tempTPMdata, method = "spearman")
  
  pancreas_nuclear_nuclear_TPM_spearman[i] <-  median(tissuecorr$rho)
  
  # for this tissue and sample, do pearson and report mean
  tissuecorr <- find.all.correlations(tempTPMdata, method = "pearson")
  
  pancreas_nuclear_nuclear_TPM_pearson[i] <-  mean(tissuecorr$rho)
  
}

pancreas_sp_mat <- matrix(nrow = 2, ncol = 2)
row.names(pancreas_sp_mat) <- c("TPM", "Excl. PRSS1/2")
colnames(pancreas_sp_mat) <- c("mean", "sd")

pancreas_sp_mat["TPM", "mean"] <- mean(pancreas_nuclear_nuclear_TPM_spearman)
pancreas_sp_mat["TPM", "sd"] <- sd(pancreas_nuclear_nuclear_TPM_spearman)

pancreas_sp_mat["Excl. PRSS1/2", "mean"] <- mean(pancreas_nuclear_nuclear_noPRSS_spearman)
pancreas_sp_mat["Excl. PRSS1/2", "sd"] <- sd(pancreas_nuclear_nuclear_noPRSS_spearman)

pancreas_sp_df <- as.data.frame(pancreas_sp_mat)
pancreas_sp_df[, "group"] <- row.names(pancreas_sp_mat)

pancreas_sp_plot_df <- cbind(pancreas_nuclear_nuclear_noPRSS_spearman, pancreas_nuclear_nuclear_TPM_spearman)
pancreas_sp_plot_df <- melt(pancreas_sp_plot_df, value.name = "rho", )

pancreas_sp_plot_df$Var2 <- relevel(factor(pancreas_sp_plot_df$Var2), ref = "pancreas_nuclear_nuclear_TPM_spearman")

# test significance
wilcox.test(pancreas_nuclear_nuclear_noPRSS_spearman, pancreas_nuclear_nuclear_TPM_spearman)

pdf("graphics/pancreas-spearman-noPRSS.pdf",
    width = 2.2,
    height = 2.2)

ggplot(data = pancreas_sp_plot_df, aes(x = Var2, y = rho), ) + 
  geom_boxplot(fill = c("grey", "magenta"),
               lwd = 0.25,
               fatten = 1)+
  theme_classic()+
  coord_cartesian(ylim = c(0, 1))+
  theme(axis.title = element_blank(),
        axis.text = element_blank(),
        axis.line = element_line(size = 0.25),
        axis.ticks = element_line(size = 0.25),
        panel.border = element_rect(colour = "black", fill = NA, size = 0.25)
  ) + 
  scale_y_continuous(breaks = c(0, 0.5, 1))

dev.off()


pancreas_prs_mat <- matrix(nrow = 2, ncol = 2)
row.names(pancreas_prs_mat) <- c("TPM", "Excl. PRSS1/2")
colnames(pancreas_prs_mat) <- c("mean", "sd")

pancreas_prs_mat["TPM", "mean"] <- mean(pancreas_nuclear_nuclear_TPM_pearson)
pancreas_prs_mat["TPM", "sd"] <- sd(pancreas_nuclear_nuclear_TPM_pearson)

pancreas_prs_mat["Excl. PRSS1/2", "mean"] <- mean(pancreas_nuclear_nuclear_noPRSS_pearson)
pancreas_prs_mat["Excl. PRSS1/2", "sd"] <- sd(pancreas_nuclear_nuclear_noPRSS_pearson)

pancreas_prs_df <- as.data.frame(pancreas_prs_mat)
pancreas_prs_df[, "group"] <- row.names(pancreas_prs_mat)
pancreas_prs_df$group <- relevel(factor(pancreas_prs_df$group), ref = "TPM")

pancreas_prs_plot_df <- cbind(pancreas_nuclear_nuclear_noPRSS_pearson, pancreas_nuclear_nuclear_TPM_pearson)
pancreas_prs_plot_df <- melt(pancreas_prs_plot_df, value.name = "r", )

pancreas_prs_plot_df$Var2 <- relevel(factor(pancreas_prs_plot_df$Var2), ref = "pancreas_nuclear_nuclear_TPM_pearson")

t.test(pancreas_nuclear_nuclear_noPRSS_pearson, pancreas_nuclear_nuclear_TPM_pearson)

pdf("graphics/pancreas-pearson-noPRSS.pdf",
    width = 2.2,
    height = 2.2)

ggplot(data = pancreas_prs_plot_df, aes(x = Var2, y = r), ) + 
  geom_boxplot(fill = c("grey", "magenta"),
               lwd = 0.25,
               outlier.size = 0.5,
               fatten = 1)+
  theme_classic()+
  coord_cartesian(ylim = c(0, 1))+
  theme(axis.title = element_blank(),
        axis.text = element_blank(),
        axis.line = element_line(size = 0.25),
        axis.ticks = element_line(size = 0.25),
        panel.border = element_rect(colour = "black", fill = NA, size = 0.25) 
  ) + 
  scale_y_continuous(breaks = c(0, 0.5, 1))

dev.off()

#### EXCLUDE HB GENES FROM BLOOD TPM COUNTS ####
# corresponds to Figure 1f, Fig S1e

blood_ids <- LUT[LUT$Tissue == "Whole Blood", "SAMPID"]
bloodcounts <- countsdata[, blood_ids]

excludedgenes <- top10names[top10names$hgnc_symbol %in% c("HBB", "HBA1", "HBA2", "HBD"), "ensembl_gene_id"]

samplescale_excl <- colSums(bloodcounts[!(row.names(bloodcounts) %in% excludedgenes), ]) / 1e6

bloodcounts_excl_normal <- t(t(bloodcounts[!(row.names(bloodcounts) %in% excludedgenes), ]) / samplescale_excl)

#restrict to ones with TPM > 5.0

bloodcounts_excl_normal <- bloodcounts_excl_normal[row.names(bloodcounts_excl_normal) %in% nuclear_gene_expressed, ]

blood_nuclear_nuclear_noPRSS_spearman <- vector(mode = "numeric", length = 10)
blood_nuclear_nuclear_noPRSS_pearson <- vector(mode = "numeric", length = 10)

blood_nuclear_nuclear_TPM_spearman <- vector(mode = "numeric", length = 10)
blood_nuclear_nuclear_TPM_pearson <- vector(mode = "numeric", length = 10)

for(i in 1:10){
  
  genesample <- sample(row.names(bloodcounts_excl_normal)[!(row.names(bloodcounts_excl_normal) %in% mitogenes$ensembl_gene_id)], 100)
  
  tempdata <- bloodcounts_excl_normal[genesample, ]
  
  tissuecorr <- find.all.correlations(tempdata, method = "spearman")
  
  blood_nuclear_nuclear_noPRSS_spearman[i] <-  median(tissuecorr$rho)
  
  # for this tissue and sample, do pearson and report mean
  tissuecorr <- find.all.correlations(tempdata, method = "pearson")
  
  blood_nuclear_nuclear_noPRSS_pearson[i] <-  mean(tissuecorr$rho)
  
  tempTPMdata <- TPMdata[genesample, blood_ids]
  
  tissuecorr <- find.all.correlations(tempTPMdata, method = "spearman")
  
  blood_nuclear_nuclear_TPM_spearman[i] <-  median(tissuecorr$rho)
  
  # for this tissue and sample, do pearson and report mean
  tissuecorr <- find.all.correlations(tempTPMdata, method = "pearson")
  
  blood_nuclear_nuclear_TPM_pearson[i] <-  mean(tissuecorr$rho)
  
}

blood_sp_mat <- matrix(nrow = 2, ncol = 2)
row.names(blood_sp_mat) <- c("TPM", "Excl. HB")
colnames(blood_sp_mat) <- c("mean", "sd")

blood_sp_mat["TPM", "mean"] <- mean(blood_nuclear_nuclear_TPM_spearman)
blood_sp_mat["TPM", "sd"] <- sd(blood_nuclear_nuclear_TPM_spearman)

blood_sp_mat["Excl. HB", "mean"] <- mean(blood_nuclear_nuclear_noPRSS_spearman)
blood_sp_mat["Excl. HB", "sd"] <- sd(blood_nuclear_nuclear_noPRSS_spearman)

blood_sp_df <- as.data.frame(blood_sp_mat)
blood_sp_df[, "group"] <- row.names(blood_sp_mat)

blood_sp_plot_df <- cbind(blood_nuclear_nuclear_noPRSS_spearman, blood_nuclear_nuclear_TPM_spearman)
blood_sp_plot_df <- melt(blood_sp_plot_df, value.name = "rho", )

blood_sp_plot_df$Var2 <- relevel(factor(blood_sp_plot_df$Var2), ref = "blood_nuclear_nuclear_TPM_spearman")

# test significance
wilcox.test(blood_nuclear_nuclear_noPRSS_spearman, blood_nuclear_nuclear_TPM_spearman)

pdf("graphics/blood-spearman-noHB.pdf",
    width = 2.2,
    height = 2.2)

ggplot(data = blood_sp_plot_df, aes(x = Var2, y = rho), ) + 
  geom_boxplot(fill = c("grey", "magenta"),
               lwd = 0.25,
               fatten = 1,
               outlier.size = 0.5)+
  theme_classic()+
  coord_cartesian(ylim = c(0, 1))+
  theme(axis.title = element_blank(),
        axis.text = element_blank(),
        axis.line = element_line(size = 0.25),
        axis.ticks = element_line(size = 0.25),
        panel.border = element_rect(colour = "black", fill = NA, size = 0.25)
  ) + 
  scale_y_continuous(breaks = c(0, 0.5, 1))

dev.off()

blood_prs_mat <- matrix(nrow = 2, ncol = 2)
row.names(blood_prs_mat) <- c("TPM", "Excl. HB")
colnames(blood_prs_mat) <- c("mean", "sd")

blood_prs_mat["TPM", "mean"] <- mean(blood_nuclear_nuclear_TPM_pearson)
blood_prs_mat["TPM", "sd"] <- sd(blood_nuclear_nuclear_TPM_pearson)

blood_prs_mat["Excl. HB", "mean"] <- mean(blood_nuclear_nuclear_noPRSS_pearson)
blood_prs_mat["Excl. HB", "sd"] <- sd(blood_nuclear_nuclear_noPRSS_pearson)

blood_prs_df <- as.data.frame(blood_prs_mat)
blood_prs_df[, "group"] <- row.names(blood_prs_mat)
blood_prs_df$group <- relevel(factor(blood_prs_df$group), ref = "TPM")

blood_prs_plot_df <- cbind(blood_nuclear_nuclear_noPRSS_pearson, blood_nuclear_nuclear_TPM_pearson)
blood_prs_plot_df <- melt(blood_prs_plot_df, value.name = "r", )

blood_prs_plot_df$Var2 <- relevel(factor(blood_prs_plot_df$Var2), ref = "blood_nuclear_nuclear_TPM_pearson")

t.test(blood_nuclear_nuclear_noPRSS_pearson, blood_nuclear_nuclear_TPM_pearson)

pdf("graphics/blood-pearson-noHB.pdf",
    width = 2.2,
    height = 2.2)

ggplot(data = blood_prs_plot_df, aes(x = Var2, y = r), ) + 
  geom_boxplot(fill = c("grey", "magenta"),
               lwd = 0.25,
               fatten = 1,
               outlier.size = 0.5)+
  theme_classic()+
  coord_cartesian(ylim = c(0, 1))+
  theme(axis.title = element_blank(),
        axis.text = element_blank(),
        axis.line = element_line(size = 0.25),
        axis.ticks = element_line(size = 0.25),
        panel.border = element_rect(colour = "black", fill = NA, size = 0.25)
  ) + 
  scale_y_continuous(breaks = c(0, 0.5, 1))

dev.off()

#### CORRELATION EXAMPLE INSR and TSPAN5 ####
# corresponds to Fig 1g, Fig S1f

# find a confident example that is negative in most tissues in MOR normalisation
# read in network for INSR from brain tissue from GIANT project at HumanBase
brainINSRnetwork <- read.table("input/GIANT-brain-coexpression-INSR-conf0.2.txt", sep = ",", header = TRUE)
brainINSRnetwork <- brainINSRnetwork[(brainINSRnetwork$GENE1 == "INSR") | (brainINSRnetwork$GENE2 == "INSR"), ]

tempcombo <- paste(brainINSRnetwork$GENE1, brainINSRnetwork$GENE2, sep = "")
INSRinteractors <- str_remove(tempcombo, pattern = "INSR")
INSRinteractors <- c(INSRinteractors, "INSR")

genesselected <- getBM(attributes = c("ensembl_gene_id", "hgnc_symbol"),
                       filters = "external_gene_name",
                       values = INSRinteractors,
                       mart = ensembl)

genesselected[genesselected$hgnc_symbol == "INSR", "ensembl_gene_id"]
genesselected <- genesselected[!(genesselected$hgnc_symbol == "INSR"), ]

# remove 6(!) DDR1 duplicate ensembl gene ids not found in the dataframe

genesselected <- genesselected[7:nrow(genesselected), ]

interactorcandidate_df <- as.data.frame(matrix(nrow = nrow(genesselected), ncol = 9))
colnames(interactorcandidate_df) <- c("gene", "MOR_median", "MOR_max", "MOR_min", "MOR_negtissues", "TPM_median", "TPM_max", "TPM_min", "TPM_negtissues")

interactorcandidate_df[, "gene"] <- genesselected$hgnc_symbol

for (j in 1:nrow(genesselected)){

  genepair <- c("ENSG00000171105", paste(genesselected[j, "ensembl_gene_id"]))
  
  example_df <- as.data.frame(matrix(nrow = length(tissues), ncol = 2))
  colnames(example_df) <- c("tissue", "MORcor")
  
  for(i in 1:length(tissues)){

    # subset the dataframe of residuals by sample ids corresponding to tissue i
    tissuedata <- MOR_tissue_list[[paste(tissues[i])]]
    
    tissuecounts <- tissuedata[genepair, ]
    
    tissueresiduals <- t(apply.lm.to.counts(tissuecounts))
    
    example_df[i, "MORcor"] <- cor.test(unlist(tissueresiduals[, 1]), unlist(tissueresiduals[, 2]))$estimate
    
  }
  
  example_df[, "tissue"] <- tissues
  
  for(i in 1:length(tissues)){
    
    # find samples corresponding to tissue i from lookup table
    tissuesamples <- LUT[LUT$Tissue == tissues[i], "SAMPID"]
    
    # subset the dataframe of residuals by sample ids corresponding to tissue i
    tissuecounts <- TPMdata[genepair, colnames(TPMdata) %in% tissuesamples]
    
    tissueresiduals <- t(apply.lm.to.counts(tissuecounts))
    
    example_df[i, "TPMcor"] <-  cor.test(unlist(tissueresiduals[, 1]), unlist(tissueresiduals[, 2]))$estimate
    
  }
  
  interactorcandidate_df[j, "TPM_median"] <- median(example_df$TPMcor)
  interactorcandidate_df[j, "TPM_max"] <- max(example_df$TPMcor)
  interactorcandidate_df[j, "TPM_min"] <- min(example_df$TPMcor)
  interactorcandidate_df[j, "TPM_negtissues"] <- sum(example_df$TPMcor < 0)
  
  interactorcandidate_df[j, "MOR_median"] <- median(example_df$MORcor)
  interactorcandidate_df[j, "MOR_max"] <- max(example_df$MORcor)
  interactorcandidate_df[j, "MOR_min"] <- min(example_df$MORcor)
  interactorcandidate_df[j, "MOR_negtissues"] <- sum(example_df$MORcor < 0)
  
}

# take TSPAN5 as example

genepair <- c("ENSG00000171105", paste(genesselected[genesselected$hgnc_symbol == "TSPAN5", "ensembl_gene_id"]))

example_df <- as.data.frame(matrix(nrow = length(tissues), ncol = 2))
colnames(example_df) <- c("tissue", "MORcor")

example_df[, "tissue"] <- tissues

for(i in 1:length(tissues)){
 
  tissuedata <- MOR_tissue_list[[tissues[i]]]
  
  # subset the dataframe of residuals by sample ids corresponding to tissue i
  tissuecounts <- tissuedata[genepair, ]
  
  tissueresiduals <- t(apply.lm.to.counts(tissuecounts))
  
  example_df[i, "MORcor"] <- cor.test(unlist(tissueresiduals[, 1]), unlist(tissueresiduals[, 2]), method = "spearman")$estimate
  
}

for(i in 1:length(tissues)){
  
  # find samples corresponding to tissue i from lookup table
  tissuesamples <- LUT[LUT$Tissue == tissues[i], "SAMPID"]
  
  # subset the dataframe of residuals by sample ids corresponding to tissue i
  tissuecounts <- TPMdata[genepair, colnames(TPMdata) %in% tissuesamples]
  
  tissueresiduals <- t(apply.lm.to.counts(tissuecounts))
  
  example_df[i, "TPMcor"] <-  cor.test(unlist(tissueresiduals[, 1]), unlist(tissueresiduals[, 2]), method = "spearman")$estimate
  
}

example_df <- example_df[order(example_df$MORcor, decreasing = TRUE),]
example_df[, "MORder"] <- 1:nrow(example_df)

example_df <- merge(example_df, MOR_tissue_correlations[, c("total_mt_expr", "tissue")])

dir.create("graphics/genepairexamples")

pdf("graphics/genepairexamples/INSR-TSPAN5-spearman.pdf",
    height = 2.2,
    width = 2.2)

ggplot(data = example_df, aes(x = MORcor, y = TPMcor, col = total_mt_expr)) + 
  geom_point(size = 0.8) + 
  scale_color_gradient2(high = "red", mid = "yellow", low = "green", midpoint = 40) + 
  theme_classic() + 
  geom_hline(yintercept = 0, linetype = "dashed", color = "grey")+
  geom_vline(xintercept = 0, linetype = "dashed", color = "grey")+
  geom_abline(slope = 1, linetype = "dashed", color = "black") +
  labs(color = "% mito transcripts", size = 0.25) + 
  coord_fixed(xlim = c(-0.6, 0.8), ylim = c(-0.6, 0.8)) +
  scale_x_continuous(breaks=seq(-0.4, 0.8, 0.4)) + 
  scale_y_continuous(breaks=seq(-0.4, 0.8, 0.4)) + 
  theme(axis.title = element_blank(),
        axis.text = element_blank(),
        axis.line = element_line(size = 0.25),
        axis.ticks = element_line(size = 0.25),
        panel.border = element_rect(colour = "black", fill = NA, size = 0.25),
        legend.position = "none"
  )


dev.off()

pdf("graphics/genepairexamples/INSR-TSPAN5-colour-key.pdf",
    height = 2.2,
    width = 2.2)

ggplot(data = example_df, aes(x = MORcor, y = TPMcor, col = total_mt_expr)) + 
  geom_point(size = 0.8) + 
  scale_color_gradient2(high = "red", mid = "yellow", low = "green", midpoint = 40) + 
  theme_classic() + 
  geom_hline(yintercept = 0, linetype = "dashed", color = "grey")+
  geom_vline(xintercept = 0, linetype = "dashed", color = "grey")+
  geom_abline(slope = 1, linetype = "dashed", color = "black") +
  labs(color = "% mito transcripts", size = 0.25) + 
  coord_fixed(xlim = c(-0.6, 0.8), ylim = c(-0.6, 0.8)) +
  scale_x_continuous(breaks=seq(-0.4, 0.8, 0.4)) + 
  scale_y_continuous(breaks=seq(-0.4, 0.8, 0.4)) + 
  theme(axis.title = element_blank(),
        axis.text = element_blank(),
        axis.line = element_line(size = 0.25),
        axis.ticks = element_line(size = 0.25),
        panel.border = element_rect(colour = "black", fill = NA, size = 0.25),
  )


dev.off()

# repeat for pearson's correlation

example_df <- as.data.frame(matrix(nrow = length(tissues), ncol = 2))
colnames(example_df) <- c("tissue", "MORcor")

example_df[, "tissue"] <- tissues

for(i in 1:length(tissues)){
  
  tissuedata <- MOR_tissue_list[[tissues[i]]]
  
  # subset the dataframe of residuals by sample ids corresponding to tissue i
  tissuecounts <- tissuedata[genepair, ]
  
  tissueresiduals <- t(apply.lm.to.counts(tissuecounts))
  
  example_df[i, "MORcor"] <- cor.test(unlist(tissueresiduals[, 1]), unlist(tissueresiduals[, 2]), method = "pearson")$estimate
  
}

for(i in 1:length(tissues)){
  
  # find samples corresponding to tissue i from lookup table
  tissuesamples <- LUT[LUT$Tissue == tissues[i], "SAMPID"]
  
  # subset the dataframe of residuals by sample ids corresponding to tissue i
  tissuecounts <- TPMdata[genepair, colnames(TPMdata) %in% tissuesamples]
  
  tissueresiduals <- t(apply.lm.to.counts(tissuecounts))
  
  example_df[i, "TPMcor"] <-  cor.test(unlist(tissueresiduals[, 1]), unlist(tissueresiduals[, 2]), method = "pearson")$estimate
  
}

example_df <- example_df[order(example_df$MORcor, decreasing = TRUE),]
example_df[, "MORder"] <- 1:nrow(example_df)

example_df <- merge(example_df, MOR_tissue_correlations[, c("total_mt_expr", "tissue")])

dir.create("graphics/genepairexamples")

pdf("graphics/genepairexamples/INSR-TSPAN5-pearson.pdf",
    height = 2.2,
    width = 2.2)

ggplot(data = example_df, aes(x = MORcor, y = TPMcor, col = total_mt_expr)) + 
  geom_point(size = 0.8) + 
  scale_color_gradient2(high = "red", mid = "yellow", low = "green", midpoint = 40) + 
  theme_classic() + 
  geom_hline(yintercept = 0, linetype = "dashed", color = "grey")+
  geom_vline(xintercept = 0, linetype = "dashed", color = "grey")+
  geom_abline(slope = 1, linetype = "dashed", color = "black") +
  labs(color = "% mito transcripts", size = 0.25) + 
  coord_fixed(xlim = c(-0.6, 0.8), ylim = c(-0.6, 0.8)) +
  scale_x_continuous(breaks=seq(-0.4, 0.8, 0.4)) + 
  scale_y_continuous(breaks=seq(-0.4, 0.8, 0.4)) + 
  theme(axis.title = element_blank(),
        axis.text = element_blank(),
        axis.line = element_line(size = 0.25),
        axis.ticks = element_line(size = 0.25),
        panel.border = element_rect(colour = "black", fill = NA, size = 0.25),
        legend.position = "none"
  )


dev.off()

#### rpkm/TPM from GTEX V6p ####
# corresponds to Fig S2

# for v6rpkm put the samples into a list
v6rpkm_tissue_list <- list()

for(i in 1:length(v6tissues)){
  
  sampleIDs <- v6LUT[v6LUT$Tissue == v6tissues[i], "SAMPID"]
  tissuerpkm <- v6prpkmdata[, sampleIDs]
  v6rpkm_tissue_list[[i]] <- tissuerpkm
  names(v6rpkm_tissue_list)[i] <- v6tissues[i]
  
}

names(v6rpkm_tissue_list) <- v6tissues

# for v6 TPM put the samples into a list
v6TPM_tissue_list <- list()

for(i in 1:length(v6tissues)){
  
  sampleIDs <- v6LUT[v6LUT$Tissue == v6tissues[i], "SAMPID"]
  tissueTPM <- v6TPMdata[, sampleIDs]
  v6TPM_tissue_list[[i]] <- tissueTPM
  names(v6TPM_tissue_list)[i] <- v6tissues[i]
  
}

names(v6TPM_tissue_list) <- v6tissues

# have to generate a new nuclear gene expressed list for v6p because the genes change from v6 to v8
v6nuclear_gene_expressed <- matrixStats::rowMedians(as.matrix(v6TPMdata))
names(v6nuclear_gene_expressed) <- row.names(v6TPMdata)

v6nuclear_gene_expressed <- names(v6nuclear_gene_expressed[v6nuclear_gene_expressed > 5])

v6nuclear_gene_expressed <- v6nuclear_gene_expressed[!(v6nuclear_gene_expressed %in% mitogenes$ensembl_gene_id)]

v6nuclear_gene_expressed <- v6nuclear_gene_expressed[!(v6nuclear_gene_expressed %in% OXPHOS_names$ensembl_gene_id)]

v6pkm_mtOXPHOS_randomnuclear <- correlate.mtOXPHOS.with.random(tissuelist = v6rpkm_tissue_list,
                                                             iterations = 10,
                                                             method = "spearman",
                                                             stat = "rho",
                                                             graphics_subfolder = "mtOXPHOS_randomnuclear",
                                                             filename = "v6rpkm-mtOXPHOS-randomnuclear-by-mito-spearman",
                                                             lookuptable = v6LUT,
                                                             tissues = v6tissues,
                                                             nuclear_gene_expressed = v6nuclear_gene_expressed,
                                                             TPMdata = v6TPMdata)

saveRDS(v6pkm_mtOXPHOS_randomnuclear, "output/mtOXPHOS_randomnuclear/v6rpkm_mtOPHOS_randomnuclear_spearman.rds")

v6TPM_mtOXPHOS_randomnuclear <- correlate.mtOXPHOS.with.random(tissuelist = v6TPM_tissue_list,
                                                               iterations = 10,
                                                               method = "spearman",
                                                               stat = "rho",
                                                               graphics_subfolder = "mtOXPHOS_randomnuclear",
                                                               filename = "v6TPM-mtOXPHOS-randomnuclear-by-mito-spearman",
                                                               lookuptable = v6LUT,
                                                               tissues = v6tissues,
                                                               nuclear_gene_expressed = v6nuclear_gene_expressed,
                                                               TPMdata = v6TPMdata)

saveRDS(v6TPM_mtOXPHOS_randomnuclear, "output/mtOXPHOS_randomnuclear/v6TPM_mtOPHOS_randomnuclear_spearman.rds")

v6MOR_mtOXPHOS_randomnuclear <- correlate.mtOXPHOS.with.random(tissuelist = v6MOR_tissue_list,
                                                               iterations = 10,
                                                               method = "spearman",
                                                               stat = "rho",
                                                               graphics_subfolder = "mtOXPHOS_randomnuclear",
                                                               filename = "v6MOR-mtOXPHOS-randomnuclear-by-mito-spearman",
                                                               lookuptable = v6LUT,
                                                               tissues = v6tissues,
                                                               nuclear_gene_expressed = v6nuclear_gene_expressed,
                                                               TPMdata = v6TPMdata)

saveRDS(v6MOR_mtOXPHOS_randomnuclear, "output/mtOXPHOS_randomnuclear/v6MOR_mtOPHOS_randomnuclear_spearman.rds")

v6pkm_randomnuclear <- correlate.random.nuclear.genes(tissuelist = v6rpkm_tissue_list,
                                                               iterations = 10,
                                                               method = "spearman",
                                                               stat = "rho",
                                                               graphics_subfolder = "random_nuclear_nuclear",
                                                               filename = "v6rpkm_random_nuclear_nuclear_by_mito_spearman",
                                                               lookuptable = v6LUT,
                                                               tissues = v6tissues,
                                                               nuclear_gene_expressed = v6nuclear_gene_expressed,
                                                               TPMdata = v6TPMdata)

saveRDS(v6pkm_randomnuclear, "output/random_nuclear_nuclear/v6rpkm_random_nuclear_nuclear_spearman.rds")


v6TPM_randomnuclear <- correlate.random.nuclear.genes(tissuelist = v6TPM_tissue_list,
                                                               iterations = 10,
                                                               method = "spearman",
                                                               stat = "rho",
                                                               graphics_subfolder = "random_nuclear_nuclear",
                                                               filename = "v6TPM-random_nuclear_nuclear_by_mito_spearman",
                                                               lookuptable = v6LUT,
                                                               tissues = v6tissues,
                                                               nuclear_gene_expressed = v6nuclear_gene_expressed,
                                                               TPMdata = v6TPMdata)

saveRDS(v6TPM_randomnuclear, "output/random_nuclear_nuclear/v6TPM_random_nuclear_nuclear_spearman.rds")

v6MOR_randomnuclear <- correlate.random.nuclear.genes(tissuelist = v6MOR_tissue_list,
                                                               iterations = 10,
                                                               method = "spearman",
                                                               stat = "rho",
                                                               graphics_subfolder = "random_nuclear_nuclear",
                                                               filename = "v6MOR-random_nuclear_nuclear_by_mito_spearman",
                                                               lookuptable = v6LUT,
                                                               tissues = v6tissues,
                                                               nuclear_gene_expressed = v6nuclear_gene_expressed,
                                                               TPMdata = v6TPMdata)

saveRDS(v6MOR_randomnuclear, "output/random_nuclear_nuclear/v6MOR_random_nuclear_nuclear_spearman.rds")

#### HEATMAPS FOR CANCER TYPES TCGA ####
# corresponds to Fig 3c, Supplementary Dataset 2

TCGA.cancertype.mtOXPHOS.nuOXPHOS.heatmaps.from.list <- function(cancerlist, 
                                                        graphics_subfolder = "", 
                                                        spearplotorder = NULL){
  
  dir.create(paste0("graphics/", graphics_subfolder), showWarnings = FALSE)
  dir.create(paste0("graphics/", graphics_subfolder, "/spearman"), showWarnings = FALSE)

  # set colours for main plot (mainpal) and sidebar (sidepal)
  mainpal <- (colorRampPalette(c("blue", "white", "red"))(100))
  sidepal <- c("orange", "purple")
  
  # make a vector with the sidebar information (nuOXPHOS or mtOXPHOS)
  sidebar_vec <- OXPHOS_names[, "Sourcelist"]
  
  # replace the values in the sidebar vector with colours specified by sidepal
  frequencies <- table(sidebar_vec)[order(table(sidebar_vec), decreasing = TRUE)]
  categories <- unique(names(frequencies))
  for(j in 1:length(categories)){
    sidebar_vec <- replace(sidebar_vec, which(sidebar_vec == categories[j]), sidepal[j])
  }
  
  # assign name information to the sidebar_vec
  names(sidebar_vec) <- OXPHOS_names[, "hgnc_symbol"]
  
  cancer_mean_corr_df <- as.data.frame(matrix(nrow = length(cancerlist), ncol = 3))
  colnames(cancer_mean_corr_df) <- c("cancertype", "total_mt_expr", "no_samples")
  
  cancer_corr_list <- list()
  
  for(i in 1:length(cancerlist)){

    # take normalised counts previously saved in list
    cancercounts <- cancerlist[[i]]
    
    # put tissue name in data frame
    cancer_mean_corr_df[i, "cancertype"] <- names(cancerlist)[i]
    
    # put tissue mean mitochondrial expression (as % TPM) in data frame
    cancer_mean_corr_df[i, "total_mt_expr"] <- mean(colSums(TCGA_TPM_data[mitogenes$ensembl_gene_id, colnames(cancercounts)])) / 1e4
    cancer_mean_corr_df[i, "no_samples"] <- ncol(cancercounts)
    
    # find all correlations using Spearman's rank correlation coefficient (rho)
    cancer_corr_spear <- find.all.correlations(cancercounts, method = "spearman", stat = "rho")
    cancer_corr_spear[,"Gene1name"] <- OXPHOS_names[cancer_corr_spear$Gene1, "hgnc_symbol"]
    cancer_corr_spear[,"Gene2name"] <- OXPHOS_names[cancer_corr_spear$Gene2, "hgnc_symbol"]
    
    cancer_corr_list[[i]] <- cancer_corr_spear
    names(cancer_corr_list)[i] <- names(cancerlist)[i]
    
    # put the mito-nuclear correlation in the dataframe
    cancer_mean_corr_df[i, "median_mtnuOXPHOS_spearman"]  <- median(cancer_corr_spear[(cancer_corr_spear$Gene1 %in% mtOXPHOS | cancer_corr_spear$Gene2 %in% mtOXPHOS) 
                                                                                      & 
                                                                                        (cancer_corr_spear$Gene1 %in% nuOXPHOS | cancer_corr_spear$Gene2 %in% nuOXPHOS)
                                                                                      , "rho"])
    
    cancer_mean_corr_df[i, "Q1_mtnuOXPHOS_spearman"]  <- quantile(cancer_corr_spear[(cancer_corr_spear$Gene1 %in% mtOXPHOS | cancer_corr_spear$Gene2 %in% mtOXPHOS) 
                                                                                    &
                                                                                      (cancer_corr_spear$Gene1 %in% nuOXPHOS | cancer_corr_spear$Gene2 %in% nuOXPHOS)
                                                                                    , "rho"], 0.25)
    
    cancer_mean_corr_df[i, "Q3_mtnuOXPHOS_spearman"]  <- quantile(cancer_corr_spear[(cancer_corr_spear$Gene1 %in% mtOXPHOS | cancer_corr_spear$Gene2 %in% mtOXPHOS) 
                                                                                    &
                                                                                      (cancer_corr_spear$Gene1 %in% nuOXPHOS | cancer_corr_spear$Gene2 %in% nuOXPHOS)
                                                                                    , "rho"], 0.75)
    
    # make correlation matrix for plotting (with spearman); then
    # change row/column names so that gene symbols appear on plots
    cancer_spear_clustermat <- build.matrix.4.cluster(corr_df = cancer_corr_spear,
                                                      stat = "rho",
                                                      plotorder = spearplotorder)
    
    row.names(cancer_spear_clustermat) <- OXPHOS_names[row.names(cancer_spear_clustermat), "hgnc_symbol"]
    colnames(cancer_spear_clustermat) <- OXPHOS_names[colnames(cancer_spear_clustermat), "hgnc_symbol"]
    
    # ensures correct order of sidebar_vec
    tempcancer_spear_sidebar_vec <- sidebar_vec[row.names(cancer_spear_clustermat)]
    
    pdf(paste0(file = "graphics/", graphics_subfolder, "/spearman/", paste0(unlist(str_split(names(cancerlist)[i], pattern = " ")), collapse = ""), ".pdf"))
    
    heatmap.2(cancer_spear_clustermat,
              main = paste0(names(cancerlist)[i], ", n = ", ncol(cancercounts)),
              symkey = FALSE,
              symbreaks = TRUE,
              breaks = NULL,
              density.info = "none",
              trace = "none",
              margins = c(12,9),
              col = mainpal,
              Rowv = FALSE,
              Colv = "Rowv",
              dendrogram = "none",
              cexRow = 0.2,
              cexCol = 0.2,
              RowSideColors = tempcancer_spear_sidebar_vec)
    
    par(lend = 1)
    legend("topright",      
           legend = categories,
           col = unique(names(table(sidebar_vec)[order(table(sidebar_vec), decreasing = TRUE)])),
           lty= 1,
           lwd = 10)
    
    dev.off()
    
  }
  
  ln_reg_spr <- lm(median_mtnuOXPHOS_spearman ~ total_mt_expr, data = cancer_mean_corr_df)
  sum_ln_reg_spr <- summary(ln_reg_spr)
  
  pdf(paste0(file = "graphics/", graphics_subfolder,  "/TCGA-mtOXPHOS-nuOXPHOS-Spearman-across-cancers.pdf"))
  
  print(ggplot(data = cancer_mean_corr_df, aes(x = total_mt_expr, y = median_mtnuOXPHOS_spearman)) +
          ylab("Median spearman's rho between mtOXPHOS and nuOXPHOS genes") + 
          xlab("% mitochondrial transcripts") +
          ggtitle("mtOXPHOS-nuOXPHOS correlations") +
          geom_point() +
          geom_smooth(method = "lm", formula = y ~ x, se = FALSE) +
          geom_errorbar(aes(x = total_mt_expr, ymax = Q3_mtnuOXPHOS_spearman, ymin = Q1_mtnuOXPHOS_spearman)) +
          theme_classic() +
          coord_cartesian(ylim = c(-1, 1)) +
          geom_text(x = 60, y = 0.5, label = deparse(bquote(R^2 == .(round(sum_ln_reg_spr$r.squared, 2)))), parse = TRUE) +
          geom_hline(yintercept = 0, linetype = "dashed", color = "grey")
  )
  
  dev.off()
  
  output_list <- list()
  
  output_list[["cancer_corr_list"]] <- cancer_corr_list
  output_list[["cancer_mean_corr_df"]] <- cancer_mean_corr_df
  
  return(output_list)
  
}

#MOR_OXPHOS_spearman_alltissues_combined <- readRDS("output/MOR_OXPHOS_spearman_alltissues_combined.rds")
#MOR_OXPHOS_spearman_plot_order <- OXPHOS_names[match(MOR_OXPHOS_spearman_alltissues_combined$heatmap_plot_order, OXPHOS_names$hgnc_symbol), "ensembl_gene_id"]

TCGA_TPM_list <- readRDS("output/TCGA_TPM_list.rds")

# apply for TPM
TCGA_TPM_OXPHOS_residuals_list <- lapply(TCGA_TPM_list, function(x){
  
  TCGA.apply.lm.to.counts(x[c(mtOXPHOS, nuOXPHOS), TCGA_LUT[TCGA_LUT$SAMPID %in% colnames(x), "SAMPID"]], lookuptable = TCGA_LUT)
  
})

TCGA_TPM_OXPHOS_residuals_cut50_list <- TCGA_TPM_OXPHOS_residuals_list[(which(sapply(TCGA_TPM_OXPHOS_residuals_list, ncol) > 50))]

TCGA_TPM_mtOXPHOS_nuOXPHOS <- TCGA.cancertype.mtOXPHOS.nuOXPHOS.heatmaps.from.list(cancerlist = TCGA_TPM_OXPHOS_residuals_cut50_list,
                                                     graphics_subfolder = "TCGA_mtnuOXPHOS_TPM",
                                                     spearplotorder = MOR_OXPHOS_spearman_plot_order)

saveRDS(TCGA_TPM_mtOXPHOS_nuOXPHOS, "output/TCGA_TPM_mtOXPHOS_nuOXPHOS_correlations.rds")

# here remove large object from memory
rm(TCGA_TPM_list)

# apply for MOR
TCGA_MOR_list <- readRDS("output/TCGA_MOR_list.rds")

TCGA_MOR_OXPHOS_residuals_list <- lapply(TCGA_MOR_list, function(x){
  
  TCGA.apply.lm.to.counts(countdata = x[c(mtOXPHOS, nuOXPHOS), colnames(x) %in% TCGA_LUT$SAMPID], lookuptable = TCGA_LUT)
  
})

TCGA_MOR_OXPHOS_residuals_cut50_list <- TCGA_MOR_OXPHOS_residuals_list[(which(sapply(TCGA_MOR_OXPHOS_residuals_list, ncol) > 50))]

TCGA_MOR_mtOXPHOS_nuOXPHOS <- TCGA.cancertype.mtOXPHOS.nuOXPHOS.heatmaps.from.list(cancerlist = TCGA_MOR_OXPHOS_residuals_cut50_list,
                                                     graphics_subfolder = "TCGA_mtnuOXPHOS_MOR",
                                                     spearplotorder = MOR_OXPHOS_spearman_plot_order)

saveRDS(TCGA_MOR_mtOXPHOS_nuOXPHOS, "output/TCGA_MOR_mtOXPHOS_nuOXPHOS_correlations.rds")
# TCGA_MOR_mtOXPHOS_nuOXPHOS <- readRDS("output/TCGA_MOR_mtOXPHOS_nuOXPHOS_correlations.rds")

# here remove large object from memory
rm(TCGA_MOR_list)

# apply for TMM
TCGA_TMM_list <- readRDS("output/TCGA_TMM_list.rds")


TCGA_TMM_OXPHOS_residuals_list <- lapply(TCGA_TMM_list, function(x){
  
  TCGA.apply.lm.to.counts(x[c(mtOXPHOS, nuOXPHOS), TCGA_LUT[TCGA_LUT$SAMPID %in% colnames(x), "SAMPID"]], lookuptable = TCGA_LUT)
  
})

TCGA_TMM_OXPHOS_residuals_cut50_list <- TCGA_TMM_OXPHOS_residuals_list[(which(sapply(TCGA_TMM_OXPHOS_residuals_list, ncol) > 50))]

TCGA_TMM_mtOXPHOS_nuOXPHOS <- TCGA.cancertype.mtOXPHOS.nuOXPHOS.heatmaps.from.list(cancerlist = TCGA_TMM_OXPHOS_residuals_cut50_list,
                                                                                   graphics_subfolder = "TCGA_mtnuOXPHOS_TMM",
                                                                                   spearplotorder = MOR_OXPHOS_spearman_plot_order)

saveRDS(TCGA_TMM_mtOXPHOS_nuOXPHOS, "output/TCGA_TMM_mtOXPHOS_nuOXPHOS_correlations.rds")

# here remove large object from memory
rm(TCGA_TMM_list)

# apply for UQ
TCGA_UQ_list <- readRDS("output/TCGA_UQ_list.rds")

TCGA_UQ_OXPHOS_residuals_list <- lapply(TCGA_UQ_list, function(x){
  
  TCGA.apply.lm.to.counts(x[c(mtOXPHOS, nuOXPHOS), TCGA_LUT[TCGA_LUT$SAMPID %in% colnames(x), "SAMPID"]], lookuptable = TCGA_LUT)
  
})

TCGA_UQ_OXPHOS_residuals_cut50_list <- TCGA_UQ_OXPHOS_residuals_list[(which(sapply(TCGA_UQ_OXPHOS_residuals_list, ncol) > 50))]

TCGA_UQ_mtOXPHOS_nuOXPHOS <- TCGA.cancertype.mtOXPHOS.nuOXPHOS.heatmaps.from.list(cancerlist = TCGA_UQ_OXPHOS_residuals_cut50_list,
                                                                                   graphics_subfolder = "TCGA_mtnuOXPHOS_UQ",
                                                                                   spearplotorder = MOR_OXPHOS_spearman_plot_order)

saveRDS(TCGA_UQ_mtOXPHOS_nuOXPHOS, "output/TCGA_UQ_mtOXPHOS_nuOXPHOS_correlations.rds")

# here remove large object from memory
rm(TCGA_UQ_list)

# apply for FPKM
TCGA_FPKM_list <- readRDS("output/TCGA_FPKM_list.rds")

TCGA_FPKM_OXPHOS_residuals_list <- lapply(TCGA_FPKM_list, function(x){
  
  TCGA.apply.lm.to.counts(x[c(mtOXPHOS, nuOXPHOS), TCGA_LUT[TCGA_LUT$SAMPID %in% colnames(x), "SAMPID"]], lookuptable = TCGA_LUT)
  
})

TCGA_FPKM_OXPHOS_residuals_cut50_list <- TCGA_FPKM_OXPHOS_residuals_list[(which(sapply(TCGA_FPKM_OXPHOS_residuals_list, ncol) > 50))]

TCGA_FPKM_mtOXPHOS_nuOXPHOS <- TCGA.cancertype.mtOXPHOS.nuOXPHOS.heatmaps.from.list(cancerlist = TCGA_FPKM_OXPHOS_residuals_cut50_list,
                                                                                  graphics_subfolder = "TCGA_mtnuOXPHOS_FPKM",
                                                                                  spearplotorder = MOR_OXPHOS_spearman_plot_order)

saveRDS(TCGA_FPKM_mtOXPHOS_nuOXPHOS, "output/TCGA_FPKM_mtOXPHOS_nuOXPHOS_correlations.rds")

# here remove large object from memory
rm(TCGA_FPKM_list)

# will replicate analysis for TPM with nuclear factors scaled by mitochondrial exclusion

TCGA_nomito_list <- lapply(TCGA_counts_list, function(x){
  nomitoscalefactor <- colSums(x[!(row.names(x) %in% mitogenes$ensembl_gene_id),]) / 1e6
  nuc_norm <- t(x[-match(mitogenes$ensembl_gene_id, row.names(x)), ]) / nomitoscalefactor
  mit_norm <- t(x[match(mitogenes$ensembl_gene_id, row.names(x)), ]) / (colSums(x) / 1e6)
  return(t(cbind(nuc_norm, mit_norm)))
})

saveRDS(TCGA_nomito_list, "output/TCGA_nomito_list.rds")

# apply for TCGA_nomito
TCGA_nomito_list <- readRDS("output/TCGA_nomito_list.rds")

TCGA_nomito_OXPHOS_residuals_list <- lapply(TCGA_nomito_list, function(x){
  
  TCGA.apply.lm.to.counts(x[c(mtOXPHOS, nuOXPHOS), TCGA_LUT[TCGA_LUT$SAMPID %in% colnames(x), "SAMPID"]], lookuptable = TCGA_LUT)
  
})

TCGA_nomito_OXPHOS_residuals_cut50_list <- TCGA_nomito_OXPHOS_residuals_list[(which(sapply(TCGA_nomito_OXPHOS_residuals_list, ncol) > 50))]

TCGA_nomito_mtOXPHOS_nuOXPHOS <- TCGA.cancertype.mtOXPHOS.nuOXPHOS.heatmaps.from.list(cancerlist = TCGA_nomito_OXPHOS_residuals_cut50_list,
                                                                                    graphics_subfolder = "TCGA_mtnuOXPHOS_nomito",
                                                                                    spearplotorder = MOR_OXPHOS_spearman_plot_order)

saveRDS(TCGA_nomito_mtOXPHOS_nuOXPHOS, "output/TCGA_nomito_mtOXPHOS_nuOXPHOS_correlations.rds")

# here remove large object from memory
rm(TCGA_TPMnomito_list)

#### TGCA mtOXPHOS CORRELATIONS WITH RANDOM NUCLEAR GENES ####
# corresponds to Fig 3d, Table S2

# have to generate a new nuclear gene expressed list for v6p because the genes change from v6 to v8
TCGAnuclear_gene_expressed <- matrixStats::rowMedians(as.matrix(TCGA_TPM_data))
names(TCGAnuclear_gene_expressed) <- row.names(TCGA_TPM_data)

TCGAnuclear_gene_expressed <- names(TCGAnuclear_gene_expressed[TCGAnuclear_gene_expressed > 5])

TCGAnuclear_gene_expressed <- TCGAnuclear_gene_expressed[!(TCGAnuclear_gene_expressed %in% mitogenes$ensembl_gene_id)]

TCGAnuclear_gene_expressed <- TCGAnuclear_gene_expressed[!(TCGAnuclear_gene_expressed %in% OXPHOS_names$ensembl_gene_id)]

# loop through and take x samples of 126 random genes
# we use 126 because this is the number of nuOXPHOS genes that we previously correlated with the mtOXPHOS genes
# with each random gene sample, loop through the tissues and calculate correlations

# create list to deposit results
# list contains a sublist for each tissue to contain correlation results

TCGA.correlate.mtOXPHOS.with.random <- function(cancerlist, 
                                           iterations, 
                                           method = c("spearman", "pearson"), 
                                           stat = c("rho", "r"),
                                           graphics_subfolder = "",
                                           filename = "correlations-mtOXPHOS-random-nuclear",
                                           lookuptable = TCGA_LUT,
                                           nuclear_gene_expressed = TCGAnuclear_gene_expressed,
                                           TPMdata = TCGA_TPM_data){
  
  method <- match.arg(method)
  stat <- match.arg(stat)
  
  mtOXPHOS_random_corr_list <- list()
  
  # now the loop in the body of the code
  for (i in 1:iterations){
    
    message(paste0("Starting iteration #", i))
    
    nuclear_sample <- sample(nuclear_gene_expressed, size = 126)
    
    mtOXPHOS_temprandom_residuals_list <- lapply(cancerlist, function(x){TCGA.apply.lm.to.counts(x[c(mtOXPHOS, nuclear_sample), lookuptable[lookuptable$SAMPID %in% colnames(x), "SAMPID"]], lookuptable = lookuptable)})
    
    mtOXPHOS_tempcorr_list <-   lapply(mtOXPHOS_temprandom_residuals_list,
                                       function(x){
                                         find.all.correlations(x, 
                                                               method = method, 
                                                               stat = stat)
                                       })
    
    mtOXPHOS_random_corr_list[[i]] <- mtOXPHOS_tempcorr_list
    
  } # finished looping through iterations
  
  message("Compiling data and producing plot")
  
  cancer_mean_corr_df <- as.data.frame(matrix(nrow = length(cancers), ncol = 3))
  colnames(cancer_mean_corr_df) <- c("cancer", "total_mt_expr", "no_samples")
  
  # many lists have underscores instead of - that features in lookup table
  cancers <- str_replace_all(names(cancerlist), pattern = "_", replacement = "-")
  
  cancer_mean_corr_df[, "cancer"] <- cancers
  
  for(j in 1:length(cancers)){
    
    sampleIDs <- lookuptable[lookuptable$cancer == cancers[j], "SAMPID"]
    cancer_mean_corr_df[j, "total_mt_expr"] <- mean(colSums(TPMdata[mitogenes$ensembl_gene_id, sampleIDs[sampleIDs %in% colnames(TPMdata)]])) / 1e4
    cancer_mean_corr_df[j, "no_samples"] <- sum(sampleIDs %in% colnames(TPMdata))
    
  } # end of loop over tissues
  
  # ensure graphics_subfolder exists
  dir.create(paste0("graphics/", graphics_subfolder), showWarnings = FALSE)

  if(method == "spearman"){
    
    # stick the mean of the medians mito-nuclear correlation in the dataframe
    cancer_mean_corr_df[, "mean_median_mtOXPHOSnuclear_spearman"]  <- rowMeans(sapply(mtOXPHOS_random_corr_list, function(k){
      sapply(k, function(x){median(x[(x[, "Gene1"] %in% mtOXPHOS|x[, "Gene2"] %in% mtOXPHOS) 
                                     & 
                                       (!(x[, "Gene1"] %in% mtOXPHOS)|!(x[, "Gene2"] %in% mtOXPHOS)),
                                     paste(stat)])})
    }))
    
    # stick the sd of the medians mito-nuclear correlation in the dataframe
    cancer_mean_corr_df[, "sd_median_mtOXPHOSnuclear_spearman"]  <- rowSds(sapply(mtOXPHOS_random_corr_list, function(k){
      sapply(k, function(x){median(x[(x$Gene1 %in% mtOXPHOS|x$Gene2 %in% mtOXPHOS) 
                                     & 
                                       (!(x$Gene1 %in% mtOXPHOS)|!(x$Gene2 %in% mtOXPHOS)), paste(stat)])})
    }))
    
    # perform linear model to assess R^2 and add to plot
    ln_reg <- lm(mean_median_mtOXPHOSnuclear_spearman ~ total_mt_expr, data = cancer_mean_corr_df)
    sum_ln_reg <- summary(ln_reg)
    
    pdf(paste0(file = "graphics/", graphics_subfolder,  "/", filename, ".pdf"))
    
    print(ggplot(data = cancer_mean_corr_df,
                 aes(x = total_mt_expr, 
                     y = mean_median_mtOXPHOSnuclear_spearman)) +
            ylab(paste0("Mean ", stat, " between mtOXPHOS and random nuclear genes")) + 
            xlab("% mitochondrial transcripts") +
            ggtitle("Correlation of mtOXPHOS and random nuclear genes") +
            geom_point() +
            geom_smooth(method = "lm", formula = y~ x, se = FALSE) +
            geom_errorbar(aes(x = total_mt_expr,
                              ymax = mean_median_mtOXPHOSnuclear_spearman + (1.96 * sd_median_mtOXPHOSnuclear_spearman / sqrt(iterations)),
                              ymin = mean_median_mtOXPHOSnuclear_spearman - (1.96 * sd_median_mtOXPHOSnuclear_spearman / sqrt(iterations)))) +
            theme_classic() +
            coord_cartesian(ylim = c(-1, 0.5)) +
            geom_text(x = 60, y = 0.5,
                      label = deparse(bquote(R^2 == .(round(sum_ln_reg$r.squared, 2)))),
                      parse = TRUE) +
            geom_hline(yintercept = 0,
                       linetype = "dashed",
                       color = "grey")
    )
    
    dev.off()
    
  } # end of if(spearman)
  
  if(method == "pearson"){
    
    # stick the mean of the means mito-nuclear correlation in the dataframe
    cancer_mean_corr_df[, "mean_mean_mtOXPHOSnuclear_pearson"]  <- rowMeans(sapply(mtOXPHOS_random_corr_list, function(k){
      sapply(k, function(x){mean(x[(x$Gene1 %in% mtOXPHOS|x$Gene2 %in% mtOXPHOS) 
                                   & 
                                     (!(x$Gene1 %in% mtOXPHOS)|!(x$Gene2 %in% mtOXPHOS)), paste(stat)])})
    }))
    
    
    
    
    
    
    # stick the sd of the means mito-nuclear correlation in the dataframe
    cancer_mean_corr_df[, "sd_mean_mtOXPHOSnuclear_pearson"]  <- rowSds(sapply(mtOXPHOS_random_corr_list, function(k){
      sapply(k, function(x){mean(x[(x$Gene1 %in% mtOXPHOS|x$Gene2 %in% mtOXPHOS) 
                                   & 
                                     (!(x$Gene1 %in% mtOXPHOS)|!(x$Gene2 %in% mtOXPHOS)), paste(stat)])})
    }))
    
    # perform linear model to assess R^2 and add to plot
    ln_reg <- lm(mean_mean_mtOXPHOSnuclear_pearson ~ total_mt_expr, data = cancer_mean_corr_df)
    sum_ln_reg <- summary(ln_reg)
    
    pdf(paste0(file = "graphics/", graphics_subfolder,  "/", filename, ".pdf"))
    
    print(ggplot(data = cancer_mean_corr_df,
                 aes(x = total_mt_expr, 
                     y = mean_mean_mtOXPHOSnuclear_pearson)) +
            ylab(paste0("Mean ", stat, " between mtOXPHOS and random nuclear genes")) + 
            xlab("% mitochondrial transcripts") +
            ggtitle("Correlation of mtOXPHOS and random nuclear genes") +
            geom_point() +
            geom_smooth(method = "lm", formula = y~ x, se = FALSE) +
            geom_errorbar(aes(x = total_mt_expr,
                              ymax = mean_mean_mtOXPHOSnuclear_pearson + (1.96 * sd_mean_mtOXPHOSnuclear_pearson / sqrt(iterations)),
                              ymin = mean_mean_mtOXPHOSnuclear_pearson - (1.96 * sd_mean_mtOXPHOSnuclear_pearson / sqrt(iterations)))) +
            theme_classic() +
            coord_cartesian(ylim = c(-1, 0.5)) +
            geom_text(x = 60, y = 0.5,
                      label = deparse(bquote(R^2 == .(round(sum_ln_reg$r.squared, 2)))),
                      parse = TRUE) +
            geom_hline(yintercept = 0,
                       linetype = "dashed",
                       color = "grey")
    )
    
    dev.off()
    
  } # end of if(pearson)
  
  output_list <- list(mtOXPHOS_random_corr_list, cancer_mean_corr_df)
  names(output_list) <- c("mito_random_correlations_by_iteration", "cancer_summary_df")
  
  return(output_list)
  
}

# TCGA_MOR_list <- readRDS("output/TCGA_MOR_list.rds")

TCGA_MOR_mtOXPHOS_randomnuclear <- TCGA.correlate.mtOXPHOS.with.random(cancerlist = TCGA_MOR_list,
                                    iterations = 100,
                                    method = "spearman",
                                    stat = "rho",
                                    graphics_subfolder = "mtOXPHOS_randomnuclear",
                                    filename = "TCGA_MOR_mtOXPHOS_randomnuclear_spearman",
                                    lookuptable = TCGA_LUT,
                                    nuclear_gene_expressed = TCGAnuclear_gene_expressed,
                                    TPMdata = TCGA_TPM_data)
                                    
saveRDS(TCGA_MOR_mtOXPHOS_randomnuclear, "output/mtOXPHOS_randomnuclear/TCGA_MOR_mtOXPHOS_randomnuclear.rds")
# TCGA_MOR_mtOXPHOS_randomnuclear <- readRDS("output/mtOXPHOS_randomnuclear/TCGA_MOR_mtOXPHOS_randomnuclear.rds")

# remove large object from memory
rm(TCGA_MOR_list)

# TCGA_TMM_list <- readRDS("output/TCGA_TMM_list.rds")

TCGA_TMM_mtOXPHOS_randomnuclear <- TCGA.correlate.mtOXPHOS.with.random(cancerlist = TCGA_TMM_list,
                                                                       iterations = 100,
                                                                       method = "spearman",
                                                                       stat = "rho",
                                                                       graphics_subfolder = "mtOXPHOS_randomnuclear",
                                                                       filename = "TCGA_TMM_mtOXPHOS_randomnuclear_spearman",
                                                                       lookuptable = TCGA_LUT,
                                                                       nuclear_gene_expressed = TCGAnuclear_gene_expressed,
                                                                       TPMdata = TCGA_TPM_data)

saveRDS(TCGA_TMM_mtOXPHOS_randomnuclear, "output/mtOXPHOS_randomnuclear/TCGA_TMM_mtOXPHOS_randomnuclear.rds")

# remove large object from memory
rm(TCGA_TMM_list)

# TCGA_UQ_list <- readRDS("output/TCGA_UQ_list.rds")

TCGA_UQ_mtOXPHOS_randomnuclear <- TCGA.correlate.mtOXPHOS.with.random(cancerlist = TCGA_UQ_list,
                                                                       iterations = 100,
                                                                       method = "spearman",
                                                                       stat = "rho",
                                                                       graphics_subfolder = "mtOXPHOS_randomnuclear",
                                                                       filename = "TCGA_UQ_mtOXPHOS_randomnuclear_spearman",
                                                                       lookuptable = TCGA_LUT,
                                                                       nuclear_gene_expressed = TCGAnuclear_gene_expressed,
                                                                       TPMdata = TCGA_TPM_data)

saveRDS(TCGA_UQ_mtOXPHOS_randomnuclear, "output/mtOXPHOS_randomnuclear/TCGA_UQ_mtOXPHOS_randomnuclear.rds")

# remove large object from memory
rm(TCGA_UQ_list)

# TCGA_FPKM_list <- readRDS("output/TCGA_FPKM_list.rds")

TCGA_FPKM_mtOXPHOS_randomnuclear <- TCGA.correlate.mtOXPHOS.with.random(cancerlist = TCGA_FPKM_list,
                                                                      iterations = 10,
                                                                      method = "spearman",
                                                                      stat = "rho",
                                                                      graphics_subfolder = "mtOXPHOS_randomnuclear",
                                                                      filename = "TCGA_FPKM_mtOXPHOS_randomnuclear_spearman",
                                                                      lookuptable = TCGA_LUT,
                                                                      nuclear_gene_expressed = TCGAnuclear_gene_expressed,
                                                                      TPMdata = TCGA_TPM_data)

saveRDS(View(TCGA_FPKM_mtOXPHOS_randomnuclear$cancer_summary_df), "output/mtOXPHOS_randomnuclear/TCGA_FPKM_mtOXPHOS_randomnuclear.rds")

# remove large object from memory
rm(TCGA_FPKM_list)

# TCGA_TPM_list <- readRDS("output/TCGA_TPM_list.rds")

TCGA_TPM_mtOXPHOS_randomnuclear <- TCGA.correlate.mtOXPHOS.with.random(cancerlist = TCGA_TPM_list,
                                                                        iterations = 10,
                                                                        method = "spearman",
                                                                        stat = "rho",
                                                                        graphics_subfolder = "mtOXPHOS_randomnuclear",
                                                                        filename = "TCGA_TPM_mtOXPHOS_randomnuclear_spearman",
                                                                        lookuptable = TCGA_LUT,
                                                                        nuclear_gene_expressed = TCGAnuclear_gene_expressed,
                                                                        TPMdata = TCGA_TPM_data)

saveRDS(TCGA_TPM_mtOXPHOS_randomnuclear, "output/mtOXPHOS_randomnuclear/TCGA_TPM_mtOXPHOS_randomnuclear.rds")

# remove large object from memory
rm(TCGA_TPM_list)

# TCGA_nomito_list <- readRDS("output/TCGA_nomito_list.rds")

TCGA_nomito_mtOXPHOS_randomnuclear <- TCGA.correlate.mtOXPHOS.with.random(cancerlist = TCGA_nomito_list,
                                                                       iterations = 10,
                                                                       method = "spearman",
                                                                       stat = "rho",
                                                                       graphics_subfolder = "mtOXPHOS_randomnuclear",
                                                                       filename = "TCGA_nomito_mtOXPHOS_randomnuclear_spearman",
                                                                       lookuptable = TCGA_LUT,
                                                                       nuclear_gene_expressed = TCGAnuclear_gene_expressed,
                                                                       TPMdata = TCGA_TPM_data)

saveRDS(TCGA_nomito_mtOXPHOS_randomnuclear, "output/mtOXPHOS_randomnuclear/TCGA_nomito_mtOXPHOS_randomnuclear.rds")

# remove large object from memory
rm(TCGA_nomito_list)

#### TCGA TEST CANCERS VS RANDOM ####
# corresponds to Table S2

TCGA.test.cancer.mtOXPHOSnuOXPHOS.against.mtOXPHOSrandom <- function(mtOXPHOSrandomnuclear_corr, 
                                                                observedmtnuOXPHOScorr,
                                                                cancers = cancertypes,
                                                                stat = c("rho", "r")){
  
  stat = match.arg(stat)
  
  output_df <- data.frame(matrix(nrow = length(cancers), ncol = 7))
  colnames(output_df) <- c("cancer", "mtnuOXPHOSmeanavg", "mtOXPHOSrandommeanavg", "mtOXPHOSrandomsd", "Shap_wilk_adj.p", "z_stat", "adj.p")
  
  output_df[, "cancer"] <- cancers
  output_df[, "mtnuOXPHOSmeanavg"] <- observedmtnuOXPHOScorr
  
  # this pulls out a matrix with tissues as columns and iterations as rows, with each median for mtOXPHOS-random nuclear correlations
  
  mtOXPHOSrand_dist <- sapply(1:length(cancers), function(i){
    sapply(mtOXPHOSrandomnuclear_corr, function(x){
      
      median(x[[i]][(x[[i]][, "Gene1"] %in% mtOXPHOS | x[[i]][, "Gene2"] %in% mtOXPHOS) & (!(x[[i]][, "Gene1"] %in% mtOXPHOS) | !(x[[i]][, "Gene2"] %in% mtOXPHOS)),
                    paste(stat)])
      
    })
  })
  
  colnames(mtOXPHOSrand_dist) <- cancers
  
  output_df[, "mtOXPHOSrandommeanavg"] <- rowMeans(t(mtOXPHOSrand_dist))
  output_df[, "mtOXPHOSrandomsd"] <- rowSds(t(mtOXPHOSrand_dist))
  
  # check assumption that distibutions of medians will be normal (according to CLT)
  output_df[, "Shap_wilk_adj.p"] <- p.adjust(apply(mtOXPHOSrand_dist, 2, function(c){
    shapiro.test(c)$p.value
  }), method = "BH")
  
  #calculate parameters of distribution
  mu_dist <- rowMeans(t(mtOXPHOSrand_dist))
  sd_dist <- rowSds(t(mtOXPHOSrand_dist))
  
  output_df[, "z_stat"] <- z_stat <- (observedmtnuOXPHOScorr - mu_dist) / sd_dist
  
  p_value <- 2 * pnorm(-abs(z_stat))
  
  output_df[, "adj.p"] <- p.adjust(p_value, method = "BH")
  
  return(output_df)
  
}


TCGA_MOR_mtOXPHOSnuOXPHOS_cancer_test <- TCGA.test.cancer.mtOXPHOSnuOXPHOS.against.mtOXPHOSrandom(mtOXPHOSrandomnuclear_corr = TCGA_MOR_mtOXPHOS_randomnuclear$mito_random_correlations_by_iteration,
                                                         observedmtnuOXPHOScorr = TCGA_MOR_mtOXPHOS_nuOXPHOS$cancer_mean_corr_df$median_mtnuOXPHOS_spearman,
                                                         cancers = TCGA_MOR_mtOXPHOS_nuOXPHOS$cancer_mean_corr_df$cancertype)

write.table(TCGA_MOR_mtOXPHOSnuOXPHOS_cancer_test, "output/TCGA_MOR_mtOXPHOSnuOXPHOS_cancer_test.txt", sep = "\t")
# TCGA_MOR_mtOXPHOSnuOXPHOS_cancer_test <- read.table("output/TCGA_MOR_mtOXPHOSnuOXPHOS_cancer_test.txt", sep = "\t")

TCGA_TMM_mtOXPHOSnuOXPHOS_cancer_test <- TCGA.test.cancer.mtOXPHOSnuOXPHOS.against.mtOXPHOSrandom(mtOXPHOSrandomnuclear_corr = TCGA_TMM_mtOXPHOS_randomnuclear$mito_random_correlations_by_iteration,
                                                                                                  observedmtnuOXPHOScorr = TCGA_TMM_mtOXPHOS_nuOXPHOS$cancer_mean_corr_df$median_mtnuOXPHOS_spearman,
                                                                                                  cancers = TCGA_TMM_mtOXPHOS_nuOXPHOS$cancer_mean_corr_df$cancertype)

write.table(TCGA_TMM_mtOXPHOSnuOXPHOS_cancer_test, "output/TCGA_TMM_mtOXPHOSnuOXPHOS_cancer_test.txt", sep = "\t")
# with TMM, 27 positive, 2 negative, 2 ns.
# with MOR, 26, 1, 4.

# TCGA_mtnuOXPHOSmedian_bootstrap <- readRDS("output/TCGA_mtnuOXPHOSmedian_bootstrap.rds")

TCGA_bootstrap_sds <- apply(TCGA_mtnuOXPHOSmedian_bootstrap, 2, sd)
TCGA_bootstrap_empiricalp <- apply(TCGA_mtnuOXPHOSmedian_bootstrap, 2, function(x){
  if(mean(x) > 0){
    sum(x < 0) / 1000
  } else {
    sum(x > 0) / 1000
  }
})

TCGA_bootstrap_empiricalpadjust <- p.adjust(TCGA_bootstrap_empiricalp[TCGA_MOR_mtOXPHOSnuOXPHOS_cancer_test$cancer])

TCGA_MOR_mtOXPHOSnuOXPHOS_forplot <- cbind(TCGA_MOR_mtOXPHOS_nuOXPHOS$cancer_mean_corr_df, TCGA_bootstrap_sds[TCGA_MOR_mtOXPHOS_nuOXPHOS$cancer_mean_corr_df$cancertype])
TCGA_MOR_mtOXPHOSnuOXPHOS_forplot <- cbind(TCGA_MOR_mtOXPHOSnuOXPHOS_forplot, TCGA_bootstrap_empiricalp[TCGA_MOR_mtOXPHOS_nuOXPHOS$cancer_mean_corr_df$cancertype])

colnames(TCGA_MOR_mtOXPHOSnuOXPHOS_forplot)[7:8] <- c("bootstrap_sd", "bootstrap_p")

# plot all correlations 

TCGA_MOR_mtOXPHOSnuOXPHOS_forplot$cancertype <- str_remove(TCGA_MOR_mtOXPHOSnuOXPHOS_forplot$cancertype, pattern = "^TCGA-")

TCGA_MOR_mtOXPHOSnuOXPHOS_forplot$cancer <- factor(TCGA_MOR_mtOXPHOSnuOXPHOS_forplot$cancertype, levels = TCGA_MOR_mtOXPHOSnuOXPHOS_forplot[order(TCGA_MOR_mtOXPHOSnuOXPHOS_forplot$median_mtnuOXPHOS_spearman), "cancertype"])
TCGA_MOR_mtOXPHOSnuOXPHOS_forplot[TCGA_MOR_mtOXPHOSnuOXPHOS_forplot$bootstrap_p < 0.05 & TCGA_MOR_mtOXPHOSnuOXPHOS_forplot$median_mtnuOXPHOS_spearman < 0, "colour"] <- "blue"
TCGA_MOR_mtOXPHOSnuOXPHOS_forplot[TCGA_MOR_mtOXPHOSnuOXPHOS_forplot$bootstrap_p < 0.05 & TCGA_MOR_mtOXPHOSnuOXPHOS_forplot$median_mtnuOXPHOS_spearman > 0, "colour"] <- "red"
TCGA_MOR_mtOXPHOSnuOXPHOS_forplot[TCGA_MOR_mtOXPHOSnuOXPHOS_forplot$bootstrap_p > 0.05, "colour"] <- "grey"

pdf("graphics/all-cancers-TCGA-mtOXPHOS-nuOXPHOS.pdf",
    height = 3.75,
    width = 7.25)

ggplot(data = TCGA_MOR_mtOXPHOSnuOXPHOS_forplot, 
       aes(x = cancer,
           y = median_mtnuOXPHOS_spearman)) +
  geom_col(data = TCGA_MOR_mtOXPHOSnuOXPHOS_forplot,
           aes(x = cancer,
               y = median_mtnuOXPHOS_spearman),
           fill = TCGA_MOR_mtOXPHOSnuOXPHOS_forplot[ ,"colour"],
           col = "black",
           size = 0.5) +
  geom_errorbar(aes(x = cancer,
                    ymin = median_mtnuOXPHOS_spearman - 1.96*bootstrap_sd,
                    ymax = median_mtnuOXPHOS_spearman + 1.96*bootstrap_sd),
                width = 0.3,
                alpha = 0.8,
                col = "black") +
  geom_hline(yintercept = 0,
             linetype = "dashed",
             color = "black") +
  theme_classic() +
  coord_cartesian(ylim = c(-0.5, 0.5)) +
  theme(axis.title.x = element_blank(),
        axis.title.y = element_blank(),
        axis.text.y = element_blank(),
        axis.text.x =  element_text(angle = 90,
                                    vjust = 1,
                                    hjust = 1,
                                    size = 8,
                                    color = "black"))

dev.off()

TCGA_MOR_mtOXPHOSnuOXPHOS_cancer_test$cancer <- str_remove(TCGA_MOR_mtOXPHOSnuOXPHOS_cancer_test$cancer, pattern = "^TCGA-")

pdf("graphics/all-cancers-TCGA-mtOXPHOS-randomnuclear.pdf",
    height = 2.57,
    width = 7)

ggplot(data = TCGA_MOR_mtOXPHOSnuOXPHOS_cancer_test, aes(x = cancer, y = mtnuOXPHOSmeanavg))+
  geom_col(data = TCGA_MOR_mtOXPHOSnuOXPHOS_cancer_test, aes(x = cancer, y = mtOXPHOSrandommeanavg), colour = "black", fill = "grey")+
  geom_errorbar(aes(x = cancer, ymin = mtOXPHOSrandommeanavg - 1.96*mtOXPHOSrandomsd, ymax = mtOXPHOSrandommeanavg + 1.96*mtOXPHOSrandomsd), width = 0.3, alpha = 0.8, col = "black")+
  geom_hline(yintercept = 0, linetype = "dashed", color = "black")+
  theme_classic()+
  coord_cartesian(ylim=c(-0.5, 0.5))+
  theme(axis.title.x = element_blank(),
        axis.title.y = element_blank(),
        axis.text.y = element_blank(),
        axis.text.x =  element_text(angle = 90, vjust = 1, hjust = 1, size = 8, color = "black"))

dev.off()

#### TCGA RANDOM NUCLEAR GENES ####
# corresponds to Fig 3e

TCGA.correlate.random.nuclear.genes <- function(cancerlist,
                                           iterations,
                                           method = c("spearman", "pearson"),
                                           stat = c("rho", "r"),
                                           graphics_subfolder = "",
                                           filename = "random-nuclear-correlations",
                                           lookuptable = LUT,
                                           nuclear_gene_expressed = TCGAnuclear_gene_expressed,
                                           TPMdata = TCGA_TPM_data) {
  
  method <- match.arg(method)
  stat <- match.arg(stat)  
  
  correlations_list <- list()
  
  for(i in 1:iterations){
    
    message(paste0("Starting iteration #", i))
    
    nuclear_sample <- sample(nuclear_gene_expressed, 100)
    
    nuclear_temprandom_residuals_list <- lapply(cancerlist, 
                                                function(x){
                                                  TCGA.apply.lm.to.counts(x[nuclear_sample, lookuptable[lookuptable$SAMPID %in% colnames(x), "SAMPID"]], lookuptable = lookuptable)
                                                })
    
    random_corr_list <- lapply(nuclear_temprandom_residuals_list,
                               function(x){
                                 find.all.correlations(x[nuclear_sample, ], 
                                                       method = method, 
                                                       stat = stat)
                               })
    
    correlations_list[[i]] <- random_corr_list
    
  } # end of loop over iterations
  
  message("Compiling data and producing plot")
  
  cancer_mean_corr_df <- as.data.frame(matrix(nrow = length(cancers), ncol = 3))
  colnames(cancer_mean_corr_df) <- c("cancer", "total_mt_expr", "no_samples")
  
  cancers <- str_replace_all(names(cancerlist), pattern = "_", replacement = "-")
  
  cancer_mean_corr_df[, "cancer"] <- cancers
  
  for(j in 1:length(cancers)){
    
    sampleIDs <- lookuptable[lookuptable$cancer == cancers[j], "SAMPID"]
    cancer_mean_corr_df[j, "total_mt_expr"] <- mean(colSums(TPMdata[mitogenes$ensembl_gene_id, sampleIDs[sampleIDs %in% colnames(TPMdata)]])) / 1e4
    cancer_mean_corr_df[j, "no_samples"] <- sum(sampleIDs %in% colnames(TPMdata))
  
  } # end of loop over tissues
  
  # ensure graphics_subfolder exists
  dir.create(paste0("graphics/", graphics_subfolder), showWarnings = FALSE)
  
  if(method == "spearman"){
    
    # stick the mean of the medians mito-nuclear correlation in the dataframe
    cancer_mean_corr_df[, "mean_median_randomnuclear_spearman"]  <- rowMeans(sapply(correlations_list, function(k){
      sapply(k, function(x){median(x[, paste(stat)])})
    }))
    
    # stick the sd of the medians mito-nuclear correlation in the dataframe
    cancer_mean_corr_df[, "sd_median_randomnuclear_spearman"]  <- rowSds(sapply(correlations_list, function(k){
      sapply(k, function(x){median(x[, paste(stat)])})
    }))
    
    # perform linear model to assess R^2 for adding to plot
    ln_reg <- lm(mean_median_randomnuclear_spearman ~ total_mt_expr, data = cancer_mean_corr_df)
    sum_ln_reg <- summary(ln_reg)
    
    pdf(paste0(file = "graphics/", graphics_subfolder,  "/", filename, ".pdf"))
    
    # plot mean correlation between random nuclear genes against total mitochondrial expression
    print(ggplot(data = cancer_mean_corr_df,
                 aes(x = total_mt_expr, 
                     y = mean_median_randomnuclear_spearman)) +
            ylab(paste0("Mean ", stat, " between random nuclear genes")) + 
            xlab("% mitochondrial transcripts") +
            ggtitle("Mean random nuclear correlation") +
            geom_point() +
            geom_smooth(method = "lm", formula = y~ x, se = FALSE) +
            geom_errorbar(aes(x = total_mt_expr,
                              ymax = mean_median_randomnuclear_spearman + (1.96 * sd_median_randomnuclear_spearman / sqrt(iterations)),
                              ymin = mean_median_randomnuclear_spearman - (1.96 * sd_median_randomnuclear_spearman / sqrt(iterations)))) +
            theme_classic() +
            coord_cartesian(ylim = c(-0.25, 1)) +
            geom_text(x = 60, y = 0.5,
                      label = deparse(bquote(R^2 == .(round(sum_ln_reg$r.squared, 2)))),
                      parse = TRUE) +
            geom_hline(yintercept = 0,
                       linetype = "dashed",
                       color = "grey")
    )
    
    dev.off()
    
  } # end of if(spearman)
  
  if(method == "pearson"){
    
    # stick the mean of the medians mito-nuclear correlation in the dataframe
    cancer_mean_corr_df[, "mean_mean_randomnuclear_pearson"]  <- rowMeans(sapply(correlations_list, function(k){
      sapply(k, function(x){mean(x[, paste(stat)])})
    }))
    
    # stick the sd of the medians mito-nuclear correlation in the dataframe
    cancer_mean_corr_df[, "sd_mean_randomnuclear_pearson"]  <- rowSds(sapply(correlations_list, function(k){
      sapply(k, function(x){mean(x[, paste(stat)])})
    }))
    
    # perform linear model to assess R^2 for adding to plot
    ln_reg <- lm(mean_mean_randomnuclear_pearson ~ total_mt_expr, data = cancer_mean_corr_df)
    sum_ln_reg <- summary(ln_reg)
    
    pdf(paste0(file = "graphics/", graphics_subfolder,  "/", filename, ".pdf"))
    
    # plot mean correlation between random nuclear genes against total mitochondrial expression
    print(ggplot(data = cancer_mean_corr_df, aes(x = total_mt_expr, y = mean_mean_randomnuclear_pearson)) +
            ylab(paste0("Mean ", stat, " between random nuclear genes")) + 
            xlab("% mitochondrial transcripts") +
            ggtitle("Mean random nuclear correlation") +
            geom_point() +
            geom_smooth(method = "lm",
                        formula = y~ x,
                        se = FALSE) +
            geom_errorbar(aes(x = total_mt_expr, 
                              ymax = mean_mean_randomnuclear_pearson + (1.96 * sd_mean_randomnuclear_pearson / sqrt(iterations)), 
                              ymin = mean_mean_randomnuclear_pearson - (1.96 * sd_mean_randomnuclear_pearson / sqrt(iterations)))) +
            theme_classic() +
            coord_cartesian(ylim = c(-0.25, 0.8)) +
            geom_text(x = 60,
                      y = 0.5,
                      label = deparse(bquote(R^2 == .(round(sum_ln_reg$r.squared, 2)))),
                      parse = TRUE) +
            geom_hline(yintercept = 0,
                       linetype = "dashed",
                       color = "grey")
    )
    
    dev.off()
    
  } # end of if(pearson)
  
  # put output objects into list, name and return
  output_list <- list(correlations_list, cancer_mean_corr_df)
  
  names(output_list) <- c("correlations_by_iteration", "cancer_summary_df")
  
  return(output_list)
  
}

# TCGA_MOR_list <- readRDS("output/TCGA_MOR_list.rds")

TCGA_MOR_random_nuclear_nuclear <- TCGA.correlate.random.nuclear.genes(cancerlist = TCGA_MOR_list,
                                                                       iterations = 100,
                                                                       method = "spearman",
                                                                       stat = "rho",
                                                                       graphics_subfolder = "random_nuclear_nuclear",
                                                                       filename = "TCGA_MOR_random_nuclear_nuclear_spearman",
                                                                       lookuptable = TCGA_LUT,
                                                                       nuclear_gene_expressed = TCGAnuclear_gene_expressed,
                                                                       cancers = cancertypes,
                                                                       TPMdata = TCGA_TPM_data)

saveRDS(TCGA_MOR_random_nuclear_nuclear, "output/random_nuclear_nuclear/TCGA_MOR_random_nuclear_nuclear.rds")

# remove large object from memory
rm(TCGA_MOR_list)

# TCGA_TPM_list <- readRDS("output/TCGA_TPM_list.rds")

TCGA_TPM_random_nuclear_nuclear <- TCGA.correlate.random.nuclear.genes(cancerlist = TCGA_TPM_list,
                                                                       iterations = 100,
                                                                       method = "spearman",
                                                                       stat = "rho",
                                                                       graphics_subfolder = "random_nuclear_nuclear",
                                                                       filename = "TCGA_TPM_random_nuclear_nuclear_spearman",
                                                                       lookuptable = TCGA_LUT,
                                                                       nuclear_gene_expressed = TCGAnuclear_gene_expressed,
                                                                       cancers = cancertypes,
                                                                       TPMdata = TCGA_TPM_data)

saveRDS(TCGA_TPM_random_nuclear_nuclear, "output/random_nuclear_nuclear/TCGA_TPM_random_nuclear_nuclear.rds")

# remove large object from memory
rm(TCGA_TPM_list)

#TCGA_TMM_list <- readRDS("output/TCGA_TMM_list.rds")

TCGA_TMM_random_nuclear_nuclear <- TCGA.correlate.random.nuclear.genes(cancerlist = TCGA_TMM_list,
                                                                       iterations = 100,
                                                                       method = "spearman",
                                                                       stat = "rho",
                                                                       graphics_subfolder = "random_nuclear_nuclear",
                                                                       filename = "TCGA_TMM_random_nuclear_nuclear_spearman",
                                                                       lookuptable = TCGA_LUT,
                                                                       nuclear_gene_expressed = TCGAnuclear_gene_expressed,
                                                                       cancers = cancertypes,
                                                                       TPMdata = TCGA_TPM_data)

saveRDS(TCGA_TMM_random_nuclear_nuclear, "output/random_nuclear_nuclear/TCGA_TMM_random_nuclear_nuclear.rds")

# remove large object from memory
rm(TCGA_TMM_list)

#TCGA_FPKM_list <- readRDS("output/TCGA_FPKM_list.rds")

TCGA_FPKM_random_nuclear_nuclear <- TCGA.correlate.random.nuclear.genes(cancerlist = TCGA_FPKM_list,
                                                                       iterations = 100,
                                                                       method = "spearman",
                                                                       stat = "rho",
                                                                       graphics_subfolder = "random_nuclear_nuclear",
                                                                       filename = "TCGA_FPKM_random_nuclear_nuclear_spearman",
                                                                       lookuptable = TCGA_LUT,
                                                                       nuclear_gene_expressed = TCGAnuclear_gene_expressed,
                                                                       cancers = cancertypes,
                                                                       TPMdata = TCGA_TPM_data)

saveRDS(TCGA_FPKM_random_nuclear_nuclear, "output/random_nuclear_nuclear/TCGA_FPKM_random_nuclear_nuclear.rds")

# remove large object from memory
rm(TCGA_FPKM_list)

#TCGA_UQ_list <- readRDS("output/TCGA_UQ_list.rds")

TCGA_UQ_random_nuclear_nuclear <- TCGA.correlate.random.nuclear.genes(cancerlist = TCGA_UQ_list,
                                                                       iterations = 100,
                                                                       method = "spearman",
                                                                       stat = "rho",
                                                                       graphics_subfolder = "random_nuclear_nuclear",
                                                                       filename = "TCGA_UQ_random_nuclear_nuclear_spearman",
                                                                       lookuptable = TCGA_LUT,
                                                                       nuclear_gene_expressed = TCGAnuclear_gene_expressed,
                                                                       cancers = cancertypes,
                                                                       TPMdata = TCGA_TPM_data)

saveRDS(TCGA_UQ_random_nuclear_nuclear, "output/random_nuclear_nuclear/TCGA_UQ_random_nuclear_nuclear.rds")

# remove large object from memory
rm(TCGA_UQ_list)

#TCGA_nomito_list <- readRDS("output/TCGA_nomito_list.rds")

TCGA_nomito_random_nuclear_nuclear <- TCGA.correlate.random.nuclear.genes(cancerlist = TCGA_nomito_list,
                                                                      iterations = 3,
                                                                      method = "spearman",
                                                                      stat = "rho",
                                                                      graphics_subfolder = "random_nuclear_nuclear",
                                                                      filename = "TCGA_nomito_random_nuclear_nuclear_spearman",
                                                                      lookuptable = TCGA_LUT,
                                                                      nuclear_gene_expressed = TCGAnuclear_gene_expressed,
                                                                      TPMdata = TCGA_TPM_data)

saveRDS(TCGA_nomito_random_nuclear_nuclear, "output/random_nuclear_nuclear/TCGA_nomito_random_nuclear_nuclear.rds")

# remove large object from memory
rm(TCGA_nomito_list)

#### TCGA all tissues combined ####  
# corresponds to Fig 3a, Fig S4a

TCGA.make.combined.tissue.mtOXPHOS.nuOXPHOS.heatmaps <- function(cancerlist, 
                                                            iterations, 
                                                            graphics_subfolder = "", 
                                                            filename = "combined_heatmap",
                                                            additionalplotorder = NULL,
                                                            orderedfilename = "combined_heatmap_ordered",
                                                            cancers = cancertypes,
                                                            lookuptable = TCGA_LUT){
  
  # create graphics subfolder, if it doesn't exist, to deposit plots
  dir.create(paste0("graphics/", graphics_subfolder), showWarnings = FALSE)
  
  # set colours for main plot (mainpal) and sidebar (sidepal)
  mainpal <- (colorRampPalette(c("blue", "white", "red"))(100))
  sidepal <- c("orange", "purple")
  
  # make a vector with the sidebar information (nuOXPHOS or mtOXPHOS)
  sidebar_vec <- OXPHOS_names[, "Sourcelist"]
  
  # replace the values in the sidebar vector with colours specified by sidepal
  frequencies <- table(sidebar_vec)[order(table(sidebar_vec), decreasing = TRUE)]
  categories <- unique(names(frequencies))
  for(j in 1:length(categories)){
    sidebar_vec <- replace(sidebar_vec, which(sidebar_vec == categories[j]), sidepal[j])
  }
  
  # assign name information to the sidebar_vec
  names(sidebar_vec) <- OXPHOS_names[, "hgnc_symbol"]
  
  # create list to deposit iterations
  allcancers_list <- list()
  
  for (i in 1:iterations) {
    
    cancersamples_rankspercent_df <- as.data.frame(matrix(nrow = nrow(OXPHOS_names), ncol = 0))
    row.names(cancersamples_rankspercent_df) <- OXPHOS_names$ensembl_gene_id
    
    message(paste0("Starting iteration #", i))
    
    for(j in 1:length(cancerlist)){

      cancercounts <- cancerlist[[j]]
      
      cancersampleIDs <- lookuptable[lookuptable$SAMPID %in% colnames(cancercounts), "SAMPID"]
      
      if(!(length(cancersampleIDs) > 50)){
        next
      }
      
      # take 50 random samples from tissue and restrict to OXPHOS genes
      samplecounts <- cancercounts[c(mtOXPHOS, nuOXPHOS), sample(cancersampleIDs, 50)]
      
      # rank the samples
      tempdata_ranks <- matrix(nrow = nrow(samplecounts),
                               ncol = ncol(samplecounts))
      row.names(tempdata_ranks) <- row.names(samplecounts)
      colnames(tempdata_ranks) <- colnames(samplecounts)
      
      for (k in 1:nrow(samplecounts)){
        tempdata_ranks[k, ] <- rank(samplecounts[k, ])
      } 
      
      # here we add the ranks for this tissue to those previously calculated
      cancersamples_rankspercent_df <- merge(cancersamples_rankspercent_df, tempdata_ranks, all.x = TRUE, all.y = FALSE, by = "row.names")
      row.names(cancersamples_rankspercent_df) <- cancersamples_rankspercent_df[, "Row.names"]
      cancersamples_rankspercent_df <- cancersamples_rankspercent_df[, 2:ncol(cancersamples_rankspercent_df)]
      
    }
    
    combinecorr <-find.all.correlations(cancersamples_rankspercent_df,
                                        method = "spearman")
    
    allcancers_list[[i]] <- combinecorr
    
  }
  
  iterations_matrix <- sapply(1:nrow(combinecorr), function(i){
    sapply(allcancers_list, function(x){x[i, 3]})
  })
  
  
  alliterations_mediancorr <- cbind(combinecorr[, 1:2], rowMedians(t(iterations_matrix)))
  colnames(alliterations_mediancorr)[3] <- "MedianSpearman"
  
  # build matrix for plotting heatmap
  alliterations_clustermat <- build.matrix.4.cluster(corr_df = alliterations_mediancorr, 
                                                     stat = "MedianSpearman")
  
  # rename rows and columns so gene symbols appear in plot
  row.names(alliterations_clustermat) <- OXPHOS_names[row.names(alliterations_clustermat), "hgnc_symbol"]
  colnames(alliterations_clustermat) <- OXPHOS_names[colnames(alliterations_clustermat), "hgnc_symbol"]
  
  # ensures correct order of sidebar_vec
  temptissue_sidebar_vec <- sidebar_vec[row.names(alliterations_clustermat)]
  
  pdf(paste0(file = "graphics/", graphics_subfolder, "/", filename, ".pdf"))
  
  heatmap <- heatmap.2(alliterations_clustermat,
                       main = "All cancers combined",
                       symkey = FALSE,
                       symbreaks = TRUE,
                       breaks = NULL,
                       density.info = "none",
                       trace = "none",
                       margins = c(12,9),
                       col = mainpal,
                       dendrogram = "none",
                       cexRow = 0.2,
                       cexCol = 0.2,
                       RowSideColors = temptissue_sidebar_vec)
  
  par(lend = 1)           
  legend("topright",      
         legend = categories,
         col = unique(names(table(sidebar_vec)[order(table(sidebar_vec), decreasing = TRUE)])),
         lty= 1,
         lwd = 10)
  
  dev.off()
  
  # save plot order according to dendogram produced with this method
  plotorder <- row.names(alliterations_clustermat)[heatmap$rowInd]
  
  functionoutput <- list()
  
  functionoutput[["iterations"]] <- allcancers_list
  functionoutput[["median_matrix"]] <- alliterations_clustermat
  functionoutput[["heatmap_plot_order"]] <- plotorder
  
  # if desired, produce ADDITIONAL plot ordered by a predefined order supplied in the arguments
  if(!(is.null(additionalplotorder))){
    
    # build matrix for plotting heatmap
    alliterations_clustermat <- build.matrix.4.cluster(corr_df = alliterations_mediancorr, 
                                                       stat = "MedianSpearman",
                                                       plotorder = additionalplotorder)
    
    # rename rows and columns so gene symbols appear in plot
    row.names(alliterations_clustermat) <- OXPHOS_names[row.names(alliterations_clustermat), "hgnc_symbol"]
    colnames(alliterations_clustermat) <- OXPHOS_names[colnames(alliterations_clustermat), "hgnc_symbol"]
    
    # ensures correct order of sidebar_vec
    temptissue_sidebar_vec <- sidebar_vec[row.names(alliterations_clustermat)]
    
    pdf(paste0(file = "graphics/", graphics_subfolder, "/", orderedfilename, ".pdf"))
    
    heatmap <- heatmap.2(alliterations_clustermat,
                         main = "All cancers combined (reordered)",
                         symkey = FALSE,
                         symbreaks = TRUE,
                         breaks = NULL,
                         density.info = "none",
                         trace = "none",
                         margins = c(12,9),
                         col = mainpal,
                         Rowv = FALSE,
                         Colv = "Rowv",
                         dendrogram = "none",
                         cexRow = 0.2,
                         cexCol = 0.2,
                         RowSideColors = temptissue_sidebar_vec)
    
    par(lend = 1)           
    legend("topright",      
           legend = categories,
           col = unique(names(table(sidebar_vec)[order(table(sidebar_vec), decreasing = TRUE)])),
           lty= 1,
           lwd = 10)
    
    dev.off()
    
  }
  
  return(functionoutput)
  
}

# apply linear model correction to MOR data. restrict to samples in the lookuptable to exclude healthy samples
TCGA_MOR_OXPHOS_lmresiduals_list <- lapply(TCGA_MOR_list, function(x){
                                          TCGA.apply.lm.to.counts(x[c(mtOXPHOS, nuOXPHOS), TCGA_LUT[TCGA_LUT$SAMPID %in% colnames(x), "SAMPID"]],
                                                                  lookuptable = TCGA_LUT)
})
  

TCGA_MOR_allcancerscombined <- TCGA.make.combined.tissue.mtOXPHOS.nuOXPHOS.heatmaps(cancerlist = TCGA_MOR_OXPHOS_lmresiduals_list, 
                                                                 iterations = 100, 
                                                                 graphics_subfolder = "TCGA_combined", 
                                                                 filename = "TCGA_combined_heatmap",
                                                                 additionalplotorder = MOR_OXPHOS_spearman_plot_order,
                                                                 orderedfilename = "TCGA_combined_heatmap_GTEXMORorder",
                                                                 cancers = cancertypes,
                                                                 lookuptable = TCGA_LUT)

saveRDS(TCGA_MOR_allcancerscombined, "output/TCGA_MOR_allcancerscombined.rds")

# apply linear model correction to MOR data. restrict to samples in the lookuptable to exclude healthy samples
TCGA_TPM_OXPHOS_lmresiduals_list <- lapply(TCGA_TPM_list, function(x){
  TCGA.apply.lm.to.counts(x[c(mtOXPHOS, nuOXPHOS), TCGA_LUT[TCGA_LUT$SAMPID %in% colnames(x), "SAMPID"]],
                          lookuptable = TCGA_LUT)
})

TCGA_TPM_allcancerscombined <- TCGA.make.combined.tissue.mtOXPHOS.nuOXPHOS.heatmaps(cancerlist = TCGA_TPM_OXPHOS_lmresiduals_list, 
                                                                                    iterations = 10, 
                                                                                    graphics_subfolder = "TCGA_combined", 
                                                                                    filename = "TCGA_TPM_combined_heatmap",
                                                                                    additionalplotorder = MOR_OXPHOS_spearman_plot_order,
                                                                                    orderedfilename = "TCGA_TPM_combined_heatmap_GTEXMORorder",
                                                                                    cancers = cancertypes,
                                                                                    lookuptable = TCGA_LUT)

saveRDS(TCGA_MOR_allcancerscombined, "output/TCGA_MOR_allcancerscombined.rds")

#### DISTRIBUTION FOR ALL CANCERS COMBINED WITH mtOXPHOS AND RANDOM NUCLEAR FACTORS ####
# corresponds to Fig 4b

# here put code in a function which can be applied to both lists of normalised values
TCGA.generate.distribution.mtOXPHOS.random.medians <- function(cancerlist, 
                                                          iterations, 
                                                          method = c("spearman", "pearson"),
                                                          stat = c("rho", "r"),
                                                          nuclear_gene_expressed = TCGAnuclear_gene_expressed,
                                                          lookuptable = TCGA_LUT)
{
  
  method <- match.arg(method)
  stat <- match.arg(stat)
  
  # create list to deposit iterations
  allcancers_list <- list()

  for (i in 1:iterations) {

    message(paste0("Starting iteration #", i))
    
    # take nuclear sample of 126 non-mito genes (same size as nuOXPHOS group)
    nuclear_sample <- sample(nuclear_gene_expressed, size = 126)
    
    randomnuclearsample_residuals <- lapply(cancerlist, function(x){
      TCGA.apply.lm.to.counts(x[c(mtOXPHOS, nuclear_sample), lookuptable[lookuptable$SAMPID %in% colnames (x), "SAMPID"]])
    })
    
    cancersamples_rankspercent_df <- data.frame(matrix(nrow = length(c(mtOXPHOS, nuclear_sample)), ncol = 0))
    row.names(cancersamples_rankspercent_df) <- c(mtOXPHOS, nuclear_sample)
    
    for(j in 1:length(randomnuclearsample_residuals)){
    
      cancersampleIDs <- lookuptable[lookuptable$SAMPID %in% colnames(randomnuclearsample_residuals[[j]]), "SAMPID"]
      
      if(!(length(cancersampleIDs) > 50)){
        next
      }
      
      # take 50 random samples from the tissue
      samplecounts <- randomnuclearsample_residuals[[j]][, sample(cancersampleIDs, 50)]
      
      # rank the samples
      tempdata_ranks <- matrix(nrow = nrow(samplecounts), ncol = ncol(samplecounts))
      row.names(tempdata_ranks) <- row.names(samplecounts)
      colnames(tempdata_ranks) <- colnames(samplecounts)
      
      for (k in 1:nrow(samplecounts)){
        tempdata_ranks[k, ] <- rank(samplecounts[k, ])
      } 
      
      # here we add the ranks for this tissue to those previously calculated
      cancersamples_rankspercent_df <- merge(cancersamples_rankspercent_df, tempdata_ranks, all.x = TRUE, all.y = FALSE, by = "row.names")
      row.names(cancersamples_rankspercent_df) <- cancersamples_rankspercent_df[, "Row.names"]
      cancersamples_rankspercent_df <- cancersamples_rankspercent_df[, 2:ncol(cancersamples_rankspercent_df)]
      
    }
    
    combinecorr <- find.all.correlations(cancersamples_rankspercent_df, 
                                         method = method,
                                         stat = stat)

    allcancers_list[[i]] <- combinecorr
    
  } # end of looping over iterations
  
  if(method == "spearman"){
    average <- function(x){
      median(x)
    }
  }
  
  if(method == "pearson"){
    average <- function(x){
      mean(x)
    }
  }
  
  iteration_avg <- sapply(allcancers_list, function(x){average(x[(x[, "Gene1"] %in% mtOXPHOS|x[, "Gene2"] %in% mtOXPHOS) 
                                                                 & 
                                                                   (!(x[, "Gene1"] %in% mtOXPHOS)|!(x[, "Gene2"] %in% mtOXPHOS)), paste(stat)])})
  
  functionoutput <- list()
  
  functionoutput[["iterations"]] <- allcancers_list
  functionoutput[["mtOXPHOS_random_avg"]] <- iteration_avg
  
  return(functionoutput)
  
}

TCGA_MOR_alltissues_mtnuOXPHOS_randomdist <- TCGA.generate.distribution.mtOXPHOS.random.medians(cancerlist = TCGA_MOR_list,
                                                   iterations = 100,
                                                   method = "spearman",
                                                   stat = "rho",
                                                   nuclear_gene_expressed = TCGAnuclear_gene_expressed)

saveRDS(TCGA_MOR_alltissues_mtnuOXPHOS_randomdist, "output/TCGA_MOR_alltissues_mtnuOXPHOS_randomdist.rds")

TCGA_MOR_mtOXPHOSnuOXPHOSorrandom_iterations <- data.frame(cbind(sapply(TCGA_MOR_allcancerscombined$iterations, function(x){median(x[(x[, "Gene1"] %in% mtOXPHOS|x[, "Gene2"] %in% mtOXPHOS) 
                                                                                                                                            & 
                                                                                                                                              (!(x[, "Gene1"] %in% mtOXPHOS)|!(x[, "Gene2"] %in% mtOXPHOS)), "rho"])})),
                                                           TCGA_MOR_alltissues_mtnuOXPHOS_randomdist$mtOXPHOS_random_avg)

colnames(TCGA_MOR_mtOXPHOSnuOXPHOSorrandom_iterations) <- c("mtOXPHOSnuOXPHOS", "mtOXPHOSrandomnuclear")

TCGA_MOR_mtOXPHOSnuOXPHOS_wilcoxtest_spearman <- wilcox.test(TCGA_MOR_mtOXPHOSnuOXPHOSorrandom_iterations$mtOXPHOSnuOXPHOS, TCGA_MOR_mtOXPHOSnuOXPHOSorrandom_iterations$mtOXPHOSrandom)
TCGA_MOR_mtOXPHOSnuOXPHOS_ttest_spearman <- t.test(TCGA_MOR_mtOXPHOSnuOXPHOSorrandom_iterations$mtOXPHOSnuOXPHOS, TCGA_MOR_mtOXPHOSnuOXPHOSorrandom_iterations$mtOXPHOSrandom)

forplottogether <- as.data.frame(cbind(rowMeans(t(TCGA_MOR_mtOXPHOSnuOXPHOSorrandom_iterations)), rowSds(t(TCGA_MOR_mtOXPHOSnuOXPHOSorrandom_iterations))))
colnames(forplottogether) <- c("mean", "sd")
forplottogether[, "group"] <- row.names(forplottogether)
forplottogether$group <- str_replace_all(string = forplottogether$group,
                                              pattern = "mtOXPHOSnuOXPHOS", 
                                              replacement = "mtOXPHOS - nuOXPHOS")

forplottogether$group <- str_replace_all(string = forplottogether$group,
                                              pattern = "mtOXPHOSrandomnuclear", 
                                              replacement = "mtOXPHOS - random nuclear")
forplottogether$group <- relevel(factor(forplottogether$group), "mtOXPHOS - random nuclear")

# plot a boxplot of the two distributions
pdf("graphics/mtOXPHOS_randomnuclear/TCGA_MOR_mtOXPHOSrandom_cancerscombined_boxplot.pdf",
    height = 2.2,
    width = 2.2)

ggplot(data = forplottogether, aes(x = group,
                                   y = mean)) +
  geom_col(fill = c("magenta", "grey"),
           col = "black",
           size = 0.25) +
  geom_hline(yintercept = 0,
             linetype = "dashed",
             color = "grey",
             size = 0.25) +
  geom_errorbar(aes(x = group,
                    ymin = mean - sd,
                    ymax = mean + sd),
                width = 0.5,
                size = 0.25) +
  theme_classic() + 
  coord_cartesian(ylim = c(-0.2, 0.2)) +
  theme(panel.border = element_rect(colour = "black",
                                    fill = NA,
                                    size = 0.25),
        axis.title.x = element_blank(),
        axis.line = element_line(size = 0.25),
        axis.ticks = element_line(size = 0.25),
        axis.title.y = element_blank(),
        axis.text.y = element_blank(),
        axis.text.x = element_blank())

dev.off()

#### TCGA MATCHED NORMAL/CANCER SAMPLES ####
# corresponds to Fig 4d, Supplementary Dataset 3, Supplementary Dataset 4

# TCGA_TPM_data <- readRDS("output/TCGA_TPM_data.rds")

# normal samples found in expression data

normsamples <- colnames(TCGA_TPM_data)[match(metadata_for_lm[metadata_for_lm$sample_type == "Solid Tissue Normal", "SAMPID"], colnames(TCGA_TPM_data))]
normsamples <- normsamples[!is.na(normsamples)]

normtissues <- sort(table(metadata_for_lm[metadata_for_lm$SAMPID %in% normsamples, "cancer"]), decreasing = TRUE)
normtissues <- names(normtissues[normtissues > 10])

# 15 tissues with at least 10 normal samples

normsamples <- metadata_for_lm[metadata_for_lm$sample_type == "Solid Tissue Normal", "SAMPID"]
normsamples <- normsamples[normsamples %in% colnames(TCGA_TPM_data)]
normsamplesLUT <- metadata_for_lm[metadata_for_lm$SAMPID %in% normsamples, ]
normsamplesLUT <- normsamplesLUT[normsamplesLUT$cancer %in% normtissues, ]
row.names(normsamplesLUT) <- normsamplesLUT$SAMPID
  
normsamples_MOR_list <- list()

# TCGA_MOR_list <- readRDS("output/TCGA_MOR_list.rds")

for(i in 1:length(unique(normsamplesLUT$cancer))){

  temptype <- unique(normsamplesLUT$cancer)[i]
  
  temptypeIDs <- normsamplesLUT[normsamplesLUT$cancer == temptype, "SAMPID"]

  normsamples_MOR_list[[i]] <- TCGA_MOR_list[[temptype]][, temptypeIDs]
  names(normsamples_MOR_list)[i] <- temptype
  
}

normsamples_mtOXPHOS_residuals_list <- lapply(normsamples_MOR_list, function(x){
  TCGA.apply.lm.to.counts(countdata = x[c(mtOXPHOS, nuOXPHOS), ], lookuptable = normsamplesLUT)
})
 
# extract donors from sample IDs to find matched tumour samples
normaldonor_ID_list <- lapply(normsamples_MOR_list, function(x) {
  str_extract(string = colnames(x), pattern = "^TCGA-[:alnum:]{2}-[:alnum:]{4}")
})

normaldonor_SAMPID_list <- lapply(normsamples_MOR_list, colnames)

# find donor IDs (first three barcode entries) for matched tumour samples
matchedcancer_ID_list <- lapply(normaldonor_ID_list, function(x){
  metadata_for_lm[metadata_for_lm$sample_type == "Primary Tumor" 
                  & str_extract(string = metadata_for_lm$SAMPID, pattern = "^TCGA-[:alnum:]{2}-[:alnum:]{4}") %in% x, "SAMPID"]
})

# some normal samples have multiple matching cancer samples. Will just take the first
matchedcancer_ID_list <- lapply(matchedcancer_ID_list, function(x){
  x[!duplicated(str_extract(string = x, pattern = "^TCGA-[:alnum:]{2}-[:alnum:]{4}"))]
})

# and now also cull some normal samples that don't have matching tumour samples
for(i in 1:length(normaldonor_SAMPID_list)){
  normaldonor_SAMPID_list[[i]] <- normaldonor_SAMPID_list[[i]][str_extract(string = normaldonor_SAMPID_list[[i]], pattern = "^TCGA-[:alnum:]{2}-[:alnum:]{4}") %in% str_extract(string = matchedcancer_ID_list[[i]], pattern = "^TCGA-[:alnum:]{2}-[:alnum:]{4}")]
}

for(i in 1:length(normsamples_mtOXPHOS_residuals_list)){

  tempcancertype <- names(normsamples_mtOXPHOS_residuals_list)[[i]]
  
  normsamples_mtOXPHOS_residuals_list[[tempcancertype]] <- normsamples_mtOXPHOS_residuals_list[[tempcancertype]][, colnames(normsamples_mtOXPHOS_residuals_list[[tempcancertype]]) %in% normaldonor_SAMPID_list[[tempcancertype]]]
}

matchedcancer_mtOXPHOS_residuals_list <- list()

for(i in 1:length(normtissues)){
  
  tempcancertype <- normtissues[i]

  matchedcancer_mtOXPHOS_residuals_list[[i]] <- TCGA_MOR_OXPHOS_lmresiduals_list[[tempcancertype]][, colnames(TCGA_MOR_OXPHOS_lmresiduals_list[[tempcancertype]]) %in% matchedcancer_ID_list[[tempcancertype]]]

  names(matchedcancer_mtOXPHOS_residuals_list)[i] <- tempcancertype
  
  }

# some of the matched cancer IDs were not in the RNA seq data.
# so one last step to cull out the normal samples that correspond

for(i in 1:length(normtissues)){
  
  tempcancertype <- normtissues[i]

  normsamples_mtOXPHOS_residuals_list[[tempcancertype]] <- normsamples_mtOXPHOS_residuals_list[[tempcancertype]][, str_extract(string = colnames(normsamples_mtOXPHOS_residuals_list[[tempcancertype]]), pattern = "^TCGA-[:alnum:]{2}-[:alnum:]{4}") %in% str_extract(string = colnames(matchedcancer_mtOXPHOS_residuals_list[[tempcancertype]]), pattern = "^TCGA-[:alnum:]{2}-[:alnum:]{4}")]
  
}

# now they match
# but now ESCA has few datapoints and will be removed

normsamples_mtOXPHOS_residuals_list <- normsamples_mtOXPHOS_residuals_list[!(names(normsamples_mtOXPHOS_residuals_list) == "TCGA-ESCA")]
matchedcancer_mtOXPHOS_residuals_list <- matchedcancer_mtOXPHOS_residuals_list[!(names(matchedcancer_mtOXPHOS_residuals_list) == "TCGA-ESCA")]

# bootstrapping confidence intervals

bootstrap.mtnuOXPHOS.corr <- function(inputlist){

bootstrap_df <- data.frame(matrix(nrow = length(inputlist), ncol = 6))
colnames(bootstrap_df) <- c("cancertype", "median", "mean", "sd", "min", "max")

for(i in 1:length(inputlist)){
  
tempsampleIDs <- colnames(inputlist[[i]])
sizeofsubsample <- round(length(tempsampleIDs)/10, digits=0)

Ncomb <- choose(length(tempsampleIDs), sizeofsubsample)

if(Ncomb<1000){
  SubSamples <- combn(tempsampleIDs, sizeofsubsample)
  } else{
  SubSamples <- list()
  for(j in 1:100){
    SubSamples[[j]] <- sample(tempsampleIDs, sizeofsubsample)
    }
}

if(is.list(SubSamples)){

  median_vec <- c()
  
  for (k in 1:100){
    
    tempdata <- inputlist[[i]][, !(colnames(inputlist[[i]]) %in% SubSamples[[k]])]
    
    tempcorr <- find.all.correlations(tempdata)
    
    OXPHOScorr_median <- median(tempcorr[(tempcorr[, "Gene1"] %in% mtOXPHOS|tempcorr[, "Gene2"] %in% mtOXPHOS) 
                               & 
                                 (!(tempcorr[, "Gene1"] %in% mtOXPHOS)|!(tempcorr[, "Gene2"] %in% mtOXPHOS)), "rho"])
    
    median_vec[k] <- OXPHOScorr_median
  }
  
} else {
  
  if(ncol(SubSamples) > 100){
    
    SubSamples <- SubSamples[, sample(ncol(SubSamples), 100)]
    
  }
  
  median_vec <- c()
  
  for (k in 1:ncol(SubSamples)){
    
    tempdata <- inputlist[[i]][, !(colnames(inputlist[[i]]) %in% SubSamples[, k])]
    
    tempcorr <- find.all.correlations(tempdata)
    
    OXPHOScorr_median <- median(tempcorr[(tempcorr[, "Gene1"] %in% mtOXPHOS|tempcorr[, "Gene2"] %in% mtOXPHOS) 
                                         & 
                                           (!(tempcorr[, "Gene1"] %in% mtOXPHOS)|!(tempcorr[, "Gene2"] %in% mtOXPHOS)), "rho"])
    
    median_vec[k] <- OXPHOScorr_median
  
}

}

bootstrap_df[i, "cancertype"] <- names(inputlist)[i]
bootstrap_df[i, "mean"] <- mean(median_vec)
bootstrap_df[i, "median"] <- median(median_vec)
bootstrap_df[i, "sd"] <- sd(median_vec)
bootstrap_df[i, "max"] <- max(median_vec)
bootstrap_df[i, "min"] <- min(median_vec)

}

return(bootstrap_df)

}

normalbootstrap_df <- bootstrap.mtnuOXPHOS.corr(normsamples_mtOXPHOS_residuals_list)

saveRDS(normalbootstrap_df, "output/TCGA_normal_bootstrap_df.rds")

normalsamples_mtnuOXPHOS_corr <- TCGA.cancertype.mtOXPHOS.nuOXPHOS.heatmaps.from.list(cancerlist = normsamples_mtOXPHOS_residuals_list,
                                                                                      graphics_subfolder = "TCGA normal heatmaps",
                                                                                      spearplotorder = MOR_OXPHOS_spearman_plot_order)

cancerbootstrap_df <- bootstrap.mtnuOXPHOS.corr(matchedcancer_mtOXPHOS_residuals_list)

saveRDS(cancerbootstrap_df, "output/TCGA_matchedcancer_bootstrap_df.rds")

matchedcancer_mtnuOXPHOS_corr <- TCGA.cancertype.mtOXPHOS.nuOXPHOS.heatmaps.from.list(cancerlist = matchedcancer_mtOXPHOS_residuals_list,
                                                                                      graphics_subfolder = "TCGA matched cancer heatmaps",
                                                                                      spearplotorder = MOR_OXPHOS_spearman_plot_order)

normalforplot <- merge(normalsamples_mtnuOXPHOS_corr$cancer_mean_corr_df, normalbootstrap_df, by = "cancertype")
normalforplot$cancertype <- str_remove(normalforplot$cancertype, "^TCGA-")
normalforplot$cancertype <- factor(normalforplot$cancertype, levels = normalforplot[order(normalforplot$median_mtnuOXPHOS_spearman), "cancertype"])

cancerforplot <- merge(matchedcancer_mtnuOXPHOS_corr$cancer_mean_corr_df, cancerbootstrap_df, by = "cancertype")
cancerforplot$cancertype <- str_remove(cancerforplot$cancertype, "^TCGA-")
cancerforplot$cancertype <- factor(cancerforplot$cancertype, levels = normalforplot[order(normalforplot$median_mtnuOXPHOS_spearman), "cancertype"])

pdf("graphics/TCGA-normal-vs-cancer-mtnuOXPHOS.pdf",
width = 5,
height = 3)

ggplot(data = normalforplot, aes(x = cancertype, y = median_mtnuOXPHOS_spearman))+
  geom_col(width = 0.3, position = position_dodge(0.7), col = "black", fill = "green", size = 0.3) +
  geom_col(data = cancerforplot, aes(x = cancertype, y = median_mtnuOXPHOS_spearman), width = 0.3, position = position_nudge(x = 0.3), fill = "magenta", col = "black", size = 0.3) +
  theme_classic()+
  geom_errorbar(aes(x = cancertype, ymin = median_mtnuOXPHOS_spearman - sd, ymax = median_mtnuOXPHOS_spearman + sd), width = 0.3, size = 0.3) +
  geom_errorbar(data = cancerforplot, aes(x = cancertype, ymin = median_mtnuOXPHOS_spearman - sd, ymax = median_mtnuOXPHOS_spearman + sd), width = 0.3, position = position_nudge(x = 0.3), size = 0.3) +
  theme(
    axis.text.x = element_text(angle = 30, vjust = 1, hjust = 1, size = 8, color = "black"),
    axis.title.x = element_blank(),
    axis.title.y = element_blank(),
    axis.text.y = element_blank()
  )+
  geom_hline(yintercept = 0, linetype = "solid", color = "black", size = 0.25)+
  coord_cartesian(ylim = c(-0.5, 0.5))

dev.off()

#### GTEx TOP/BOTTOM 1000 GENES WITH EXPRESSION CORRELATING TO MTOXPHOS-NUOXPHOS CORRELATION ####
# corresponds to Supplementary Dataset 5, Supplementary Dataset 6

# expression across tissues for each gene

# MOR_across_tissues <- readRDS("output/MOR-normalisation-across-tissues.rds")

tissue_means <- matrix(nrow = nrow(MOR_across_tissues),
                       ncol = length(tissues))

row.names(tissue_means) <- row.names(MOR_across_tissues)
colnames(tissue_means) <- tissues

for(i in 1:length(tissues)){

  samplesIDs <- LUT[LUT$Tissue == tissues[i], "SAMPID"]
  tempdata <- MOR_across_tissues[, samplesIDs]
  
  for(j in 1:nrow(tempdata)){
    
    tissue_means[j, i] <- mean(tempdata[j, ]) 
    
  }
  
}

# MOR_tissue_correlations <- readRDS("output/MOR-OXPHOS-correlation-by-tissue.rds")

genecorrs <- sapply(1:nrow(tissue_means), function(i){
  tempcor <- cor.test(tissue_means[i, ], MOR_tissue_correlations$median_mtnuOXPHOS_spearman, method = "spearman")
  tempcor$estimate
})

genepvalues <- sapply(1:nrow(tissue_means), function(i){
  tempcor <- cor.test(tissue_means[i, ], MOR_tissue_correlations$median_mtnuOXPHOS_spearman, method = "spearman")
  tempcor$p.value
})

names(genecorrs) <- row.names(tissue_means)
names(genepvalues) <- row.names(tissue_means)

genep_adjust <- p.adjust(genepvalues, method = "BH")

gene_corr_p <- data.frame(cbind(genecorrs, genep_adjust))

# try top 1000

gene_corr_top1000 <- gene_corr_p[order(gene_corr_p$genecorrs, decreasing = TRUE), ]
gene_corr_top1000 <- gene_corr_top1000[1:1000, ]

# try bottom 1000 correlated genes 

gene_corr_bottom1000 <- gene_corr_p[order(gene_corr_p$genecorrs, decreasing = FALSE), ]
gene_corr_bottom1000 <- gene_corr_bottom1000[1:1000, ]

# do the mito genes pop out?
mtOXPHOS %in% row.names(gene_corr_top1000)
mtOXPHOS %in% row.names(gene_corr_bottom1000)

nuOXPHOS %in% row.names(gene_corr_top1000)
nuOXPHOS %in% row.names(gene_corr_bottom1000)

# remove the mitochondrial genes

gene_corr_top1000 <- gene_corr_top1000[!(row.names(gene_corr_top1000) %in% c(mtOXPHOS, nuOXPHOS)), ]
gene_corr_bottom1000 <- gene_corr_bottom1000[!(row.names(gene_corr_bottom1000) %in% c(mtOXPHOS, nuOXPHOS)), ]

genesymbolspos1000 <- getBM(mart = ensembl,
                                attributes = c("hgnc_symbol", "ensembl_gene_id"),
                                filters = "ensembl_gene_id",
                                values = row.names(gene_corr_top1000))

genesymbolsneg1000 <- getBM(mart = ensembl,
                                attributes = c("hgnc_symbol", "ensembl_gene_id"),
                                filters = "ensembl_gene_id",
                                values = row.names(gene_corr_bottom1000))

dbs <- c("KEGG_2021_Human",
         "GO_Molecular_Function_2018",
         "GO_Cellular_Component_2018",
         "GO_Biological_Process_2018",
         "Genome_Browser_PWMs",
         "TRANSFAC_and_JASPAR_PWMs",
         "ENCODE_TF_ChIP-seq_2014")

pos1000enriched <- enrichr(unlist(genesymbolspos1000$hgnc_symbol), dbs)
neg1000enriched <- enrichr(unlist(genesymbolsneg1000$hgnc_symbol), dbs)

View(neg1000enriched$TRANSFAC_and_JASPAR_PWMs)

write.xlsx(list("FDR0.05_KEGG_2021_Human" = pos1000enriched$KEGG_2021_Human, 
                "FDR0.05_GO_Molecular_Function_2018" = pos1000enriched$GO_Molecular_Function_2018,
                "FDR0.05_GO_Cellular_Component_2018" = pos1000enriched$GO_Cellular_Component_2018,
                "FDR0.05_GO_Biological_Process_2018" = pos1000enriched$GO_Biological_Process_2018),
           file = "output/GTEX_genewise_expression_corr_enrichment_top1000.xlsx")

write.xlsx(list("FDR0.05_KEGG_2021_Human" = neg1000enriched$KEGG_2021_Human, 
                "FDR0.05_GO_Molecular_Function_2018" = neg1000enriched$GO_Molecular_Function_2018,
                "FDR0.05_GO_Cellular_Component_2018" = neg1000enriched$GO_Cellular_Component_2018,
                "FDR0.05_GO_Biological_Process_2018" = neg1000enriched$GO_Biological_Process_2018),
           file = "output/GTEX_genewise_expression_corr_enrichment_bottom1000.xlsx")

# check out IG loci

# locus information downloaded from: [https://www.genenames.org/data/genegroup/#!/group/348]
IGKorphons <- read.table("input/IGKorphons.txt", sep = "\t", header = TRUE, stringsAsFactors = FALSE, quote = "")
IGHlocus <-read.table("input/IGHlocus14.txt", sep = "\t", header = TRUE, stringsAsFactors = FALSE, quote = "")
IGHorphons <-read.table("input/IGHorphons.txt", sep = "\t", header = TRUE, stringsAsFactors = FALSE, quote = "")
IGLlocus <-read.table("input/IGLlocus22.txt", sep = "\t", header = TRUE, stringsAsFactors = FALSE, quote = "")
IGLorphons <-read.table("input/IGLorphons.txt", sep = "\t", header = TRUE, stringsAsFactors = FALSE, quote = "")
IGKlocus <- read.table("input/IGKlocus2.txt", sep = "\t", header = TRUE, stringsAsFactors = FALSE, quote = "")

IGKlocus_ensembl <- getBM(attributes = c("ensembl_gene_id", "hgnc_symbol"),
                          values = IGKlocus$Approved.symbol,
                          filters = "hgnc_symbol",
                          mart = ensembl)

IGHlocus_ensembl <- getBM(attributes = c("ensembl_gene_id", "hgnc_symbol"),
                          values = IGHlocus$Approved.symbol,
                          filters = "hgnc_symbol",
                          mart = ensembl)

IGLlocus_ensembl <- getBM(attributes = c("ensembl_gene_id", "hgnc_symbol"),
                          values = IGLlocus$Approved.symbol,
                          filters = "hgnc_symbol",
                          mart = ensembl)

IGKor_ensembl <- getBM(attributes = c("ensembl_gene_id", "hgnc_symbol"),
                          values = IGKorphons$Approved.symbol,
                          filters = "hgnc_symbol",
                          mart = ensembl)

IGHor_ensembl <- getBM(attributes = c("ensembl_gene_id", "hgnc_symbol"),
                          values = IGHorphons$Approved.symbol,
                          filters = "hgnc_symbol",
                          mart = ensembl)

# none of the 6 IGL orphons present in top genes.

sum(genesymbolspos1000$hgnc_symbol %in% IGKlocus$Approved.symbol)
sum(genesymbolspos1000$hgnc_symbol %in% IGKorphons$Approved.symbol)
sum(genesymbolspos1000$hgnc_symbol %in% IGHlocus$Approved.symbol)
sum(genesymbolspos1000$hgnc_symbol %in% IGHorphons$Approved.symbol)
sum(genesymbolspos1000$hgnc_symbol %in% IGLlocus$Approved.symbol)
sum(genesymbolspos1000$hgnc_symbol %in% IGLorphons$Approved.symbol)

IGKpresent <- genesymbolspos1000[genesymbolspos1000$hgnc_symbol %in% IGKlocus[IGKlocus$Approved.symbol %in% genesymbolspos1000$hgnc_symbol, "Approved.symbol"], "ensembl_gene_id"]
IGLpresent <- genesymbolspos1000[genesymbolspos1000$hgnc_symbol %in% IGLlocus[IGLlocus$Approved.symbol %in% genesymbolspos1000$hgnc_symbol, "Approved.symbol"], "ensembl_gene_id"]
IGHpresent <- genesymbolspos1000[genesymbolspos1000$hgnc_symbol %in% IGHlocus[IGHlocus$Approved.symbol %in% genesymbolspos1000$hgnc_symbol, "Approved.symbol"], "ensembl_gene_id"]
IGKorpresent <- genesymbolspos1000[genesymbolspos1000$hgnc_symbol %in% IGKorphons[IGKorphons$Approved.symbol %in% genesymbolspos1000$hgnc_symbol, "Approved.symbol"], "ensembl_gene_id"]
IGLorpresent <- genesymbolspos1000[genesymbolspos1000$hgnc_symbol %in% IGLorphons[IGLorphons$Approved.symbol %in% genesymbolspos1000$hgnc_symbol, "Approved.symbol"], "ensembl_gene_id"]
IGHorpresent <- genesymbolspos1000[genesymbolspos1000$hgnc_symbol %in% IGHorphons[IGHorphons$Approved.symbol %in% genesymbolspos1000$hgnc_symbol, "Approved.symbol"], "ensembl_gene_id"]

IGloci <- cbind(MOR_tissue_correlations$median_mtnuOXPHOS_spearman,
      colSums(tissue_means[IGKpresent, ]),
      colSums(tissue_means[IGHpresent, ]),
      colSums(tissue_means[IGLpresent, ]),
      colSums(tissue_means[IGHorpresent, ]),
      colSums(tissue_means[IGKorpresent, ]))

cor.test(IGloci[,1], IGloci[, 2])
cor.test(IGloci[,1], IGloci[, 3])
cor.test(IGloci[,1], IGloci[, 4])
cor.test(IGloci[,1], IGloci[, 5])
cor.test(IGloci[,1], IGloci[, 6])

plot(log10(colSums(tissue_means[IGKpresent, ])), MOR_tissue_correlations$median_mtnuOXPHOS_spearman, pch = MOR_tissue_correlations$tissue)
plot(log10(colSums(tissue_means[IGHpresent, ])), MOR_tissue_correlations$median_mtnuOXPHOS_spearman, pch = MOR_tissue_correlations$tissue)
plot(log10(colSums(tissue_means[IGLpresent, ])), MOR_tissue_correlations$median_mtnuOXPHOS_spearman, pch = MOR_tissue_correlations$tissue)

plot(log10(colSums(tissue_means[IGKorpresent, ])), MOR_tissue_correlations$median_mtnuOXPHOS_spearman, pch = MOR_tissue_correlations$tissue)
plot(log10(colSums(tissue_means[IGHorpresent, ])), MOR_tissue_correlations$median_mtnuOXPHOS_spearman, pch = MOR_tissue_correlations$tissue)

plot(log10(colSums(tissue_means[c(IGKpresent, IGHpresent, IGLpresent, IGHorpresent, IGKorpresent), ])), MOR_tissue_correlations$median_mtnuOXPHOS_spearman, pch = MOR_tissue_correlations$tissue)

cor.test(colSums(tissue_means[c(IGKpresent, IGHpresent, IGLpresent, IGHorpresent, IGKorpresent), ]), MOR_tissue_correlations$median_mtnuOXPHOS_spearman,)

# MOR_across_tissues <- readRDS('output/MOR-normalisation-across-tissues.rds')
# MOR_tissue_correlations <- readRDS("output/MOR-OXPHOS-correlation-by-tissue.rds")

IGs_long <- matrix(ncol = 8, nrow = 0)
colnames(IGs_long) <- c("Tissue", "Total_IG", "TotalIGHlocus", "TotalIGHorphons", "TotalIGKlocus", "TotalIGKorphons", "TotalIGLlocus", "OXPHOScorr")

for(i in 1:length(tissues)){

  sampIDS <- LUT[LUT$Tissue == tissues[i], "SAMPID"]
  tempdata <- MOR_across_tissues[row.names(MOR_across_tissues) %in% c(IGHlocus_ensembl$ensembl_gene_id, IGKlocus_ensembl$ensembl_gene_id, IGLlocus_ensembl$ensembl_gene_id, IGHor_ensembl$ensembl_gene_id, IGKor_ensembl$ensembl_gene_id), sampIDS]
  
  tissuelabelvec <- vector(length = length(sampIDS))
  tissuelabelvec[] <- tissues[i]
  
  tissueOXPHOScorr <- vector(length = length(sampIDS))
  tissueOXPHOScorr[] <- MOR_tissue_correlations[MOR_tissue_correlations$tissue == tissues[i], "median_mtnuOXPHOS_spearman"]
  
  temp_df <- cbind(tissuelabelvec,
                   colSums(tempdata[, ]),
                   colSums(tempdata[row.names(tempdata) %in% IGHlocus_ensembl$ensembl_gene_id, ]),
                   colSums(tempdata[row.names(tempdata) %in% IGHor_ensembl$ensembl_gene_id, ]),
                   colSums(tempdata[row.names(tempdata) %in% IGKlocus_ensembl$ensembl_gene_id, ]),
                   colSums(tempdata[row.names(tempdata) %in% IGKor_ensembl$ensembl_gene_id, ]),
                   colSums(tempdata[row.names(tempdata) %in% IGLlocus_ensembl$ensembl_gene_id, ]),
                   tissueOXPHOScorr)

  colnames(temp_df) <- c("Tissue", "Total_IG", "TotalIGHlocus", "TotalIGHorphons", "TotalIGKlocus", "TotalIGKorphons", "TotalIGLlocus", "OXPHOScorr")
  
  IGs_long <- rbind(IGs_long, temp_df)
  
}

IGs_long <- data.frame(IGs_long)

totalIG_linefit <- lm(log10(as.numeric(IGs_long$Total_IG)) ~ as.numeric(IGs_long$OXPHOScorr))
sum_totalIG_linefit <- summary(totalIG_linefit)

pdf("graphics/allIGsviolinplot.pdf", width = 8, height = 4)

ggplot(data = IGs_long,
       aes(x = as.numeric(OXPHOScorr),
           y = as.numeric(Total_IG))) +
  geom_violin(aes(group = Tissue,
                  fill = Tissue),
              show.legend = FALSE,
              scale = 1,
              width = 0.015,
              position = "identity",
              lwd = 0.2) + 
  ylab("Total IG expression (pseudocounts)") +
  xlab("mtOXPHOS-nuOXPHOS correlation") +
  theme_classic()+
  theme(axis.text.x = element_text(angle = 60,
                                   vjust = 1,
                                   hjust=1,
                                   family = "sans",
                                   color = "black",
                                   size = 5),
        axis.line = element_line(size = 0.25),
        axis.ticks = element_line(size = 0.25),
        legend.position = "none") +
  coord_cartesian(xlim = c(-0.4, 0.4)) +
  geom_text(x = -0.2,
            y = 7,
            label = deparse(bquote(R^2 == .(round(sum_totalIG_linefit$r.squared, 2)))),
            parse = TRUE,
            size = 2.3) +
  geom_text(x = -0.2,
            y = 6.5,
            label = deparse(bquote('p < 2 x' ~10^-16)),
            parse = TRUE,
            size = 2.3) +
  geom_abline(color='red',
              intercept = sum_totalIG_linefit$coefficients[1,1],
              slope = sum_totalIG_linefit$coefficients[2,1]) +
  scale_y_continuous(trans = "log10",
                     labels = function(x) format(x, scientific = FALSE))
  
dev.off()

IGH_linefit <- lm(log10(as.numeric(IGs_long$TotalIGHlocus)) ~ as.numeric(IGs_long$OXPHOScorr))
sum_IGH_linefit <- summary(IGH_linefit)

pdf("graphics/IGHviolinplot.pdf", width = 8, height = 4)

ggplot(data = IGs_long, aes(x = as.numeric(OXPHOScorr), y = as.numeric(TotalIGHlocus))) +
  geom_violin(aes(group = Tissue, fill = Tissue), show.legend = FALSE, scale = 1, width = 0.015, position = "identity", lwd = 0.2) + 
  ylab("Total IGH expression (pseudocounts)") +
  xlab("mtOXPHOS-nuOXPHOS correlation") +
  theme_classic()+
  theme(axis.text.x = element_text(angle = 60,
                                   vjust = 1,
                                   hjust=1,
                                   family = "sans",
                                   color = "black",
                                   size = 5),
        axis.line = element_line(size = 0.25),
        axis.ticks = element_line(size = 0.25),
        legend.position = "none") +
  coord_cartesian(xlim = c(-0.4, 0.4)) +
  geom_text(x = -0.12, y = 6, label = deparse(bquote(R^2 == .(round(sum_IGH_linefit$r.squared, 2)))), parse = TRUE, size = 2.3)+
  geom_text(x = -0.12, y = 5.5, label = deparse(bquote('p < 2 x' ~10^-16)), parse = TRUE, size = 2.3)+
  geom_abline(color='red', intercept = sum_IGH_linefit$coefficients[1,1], slope = sum_IGH_linefit$coefficients[2,1]) +
  scale_y_continuous(trans = "log10", labels = function(x) format(x, scientific = FALSE))

dev.off()

IGK_linefit <- lm(log10(as.numeric(IGs_long$TotalIGKlocus)+0.0001) ~ as.numeric(IGs_long$OXPHOScorr))
sum_IGK_linefit <- summary(IGK_linefit)

pdf("graphics/IGKviolinplot.pdf", width = 8, height = 4)

ggplot(data = IGs_long, aes(x = as.numeric(OXPHOScorr), y = as.numeric(TotalIGKlocus))) +
  geom_violin(aes(group = Tissue, fill = Tissue), show.legend = FALSE, scale = 1, width = 0.015, position = "identity", lwd = 0.2) + 
  ylab("Total IGK expression (pseudocounts)") +
  xlab("mtOXPHOS-nuOXPHOS correlation") +
  theme_classic()+
  theme(axis.text.x = element_text(angle = 60,
                                   vjust = 1,
                                   hjust=1,
                                   family = "sans",
                                   color = "black",
                                   size = 5),
        axis.line = element_line(size = 0.25),
        axis.ticks = element_line(size = 0.25),
        legend.position = "none") +
  coord_cartesian(xlim = c(-0.4, 0.4)) +
  geom_text(x = -0.2, y = 6, label = deparse(bquote(R^2 == .(round(sum_IGK_linefit$r.squared, 2)))), parse = TRUE, size = 2.3)+
  geom_text(x = -0.2, y = 5.5, label = deparse(bquote('p < 2 x' ~10^-16)), parse = TRUE, size = 2.3)+
  geom_abline(color='red', intercept = sum_IGK_linefit$coefficients[1,1], slope = sum_IGK_linefit$coefficients[2,1]) +  
  scale_y_continuous(trans = "log10", labels = function(x) format(x, scientific = FALSE))

dev.off()

IGL_linefit <- lm(log10(as.numeric(IGs_long$TotalIGLlocus)+0.0001) ~ as.numeric(IGs_long$OXPHOScorr))
sum_IGL_linefit <- summary(IGL_linefit)

pdf("graphics/IGLviolinplot.pdf", width = 8, height = 4)

ggplot(data = IGs_long, aes(x = as.numeric(OXPHOScorr), y = as.numeric(TotalIGLlocus))) +
  geom_violin(aes(group = Tissue, fill = Tissue), show.legend = FALSE, scale = 1, width = 0.015, position = "identity", lwd = 0.2) + 
  ylab("Total IGL expression (pseudocounts)") +
  xlab("mtOXPHOS-nuOXPHOS correlation") +
  theme_classic()+
  theme(axis.text.x = element_text(angle = 60,
                                   vjust = 1,
                                   hjust=1,
                                   family = "sans",
                                   color = "black",
                                   size = 5),
        axis.line = element_line(size = 0.25),
        axis.ticks = element_line(size = 0.25),
        legend.position = "none") +
  coord_cartesian(xlim = c(-0.4, 0.4)) +
  geom_text(x = -0.2, y = 6, label = deparse(bquote(R^2 == .(round(sum_IGL_linefit$r.squared, 2)))), parse = TRUE, size = 2.3)+
  geom_text(x = -0.2, y = 5.5, label = deparse(bquote('p < 2 x' ~10^-16)), parse = TRUE, size = 2.3)+
  geom_abline(color='red', intercept = sum_IGL_linefit$coefficients[1,1], slope = sum_IGL_linefit$coefficients[2,1])+
  scale_y_continuous(trans = "log10", labels = function(x) format(x, scientific = FALSE))

dev.off()

#### TCGA TOP/BOTTOM 1000 GENES WITH EXPRESSION CORRELATING TO MTOXPHOS-NUOXPHOS CORRELATION ####
# corresponds to Supplementary Dataset 7, Supplementary Dataset 8

# expression across tissues for each gene

TCGA_MOR_allcancerscombined <- readRDS("output/TCGA-MOR-normalisation-across-cancers.rds")

tumour_means <- matrix(nrow = nrow(TCGA_MOR_allcancerscombined),
                       ncol = length(cancertypes))

row.names(tumour_means) <- row.names(TCGA_MOR_allcancerscombined)
colnames(tumour_means) <- cancertypes

for(i in 1:length(cancertypes)){

  samplesIDs <- TCGA_LUT[TCGA_LUT$cancer == cancertypes[i], "SAMPID"]
  tempdata <- TCGA_MOR_allcancerscombined[, colnames(TCGA_MOR_allcancerscombined) %in% samplesIDs]
  
  for(j in 1:nrow(tempdata)){
    
    tumour_means[j, i] <- mean(tempdata[j, ]) 
    
  }
  
}

MOR_tumour_correlations <- readRDS("output/TCGA_MOR_mtOXPHOS_nuOXPHOS_correlations.rds")
MOR_tumour_correlations <- MOR_tumour_correlations$cancer_mean_corr_df

# restrict mean to cancertypes present in OXPHOS corr analysis
tumour_means <- tumour_means[, MOR_tumour_correlations$cancertype]

TCGAgenecorrs <- sapply(1:nrow(tumour_means), function(i){
  tempcor <- cor.test(tumour_means[i, ], MOR_tumour_correlations$median_mtnuOXPHOS_spearman, method = "spearman")
  tempcor$estimate
})

TCGAgenepvalues <- sapply(1:nrow(tumour_means), function(i){
  tempcor <- cor.test(tumour_means[i, ], MOR_tumour_correlations$median_mtnuOXPHOS_spearman, method = "spearman")
  tempcor$p.value
})

names(TCGAgenecorrs) <- row.names(tumour_means)
names(TCGAgenepvalues) <- row.names(tumour_means)

TCGAgenep_adjust <- p.adjust(TCGAgenepvalues, method = "BH")

TCGAgene_corr_p <- data.frame(cbind(TCGAgenecorrs, TCGAgenep_adjust))

# try top 1000 correlated genes 

TCGAgene_corr_topp1000 <- TCGAgene_corr_p[order(TCGAgene_corr_p$TCGAgenecorrs, decreasing = TRUE), ]
TCGAgene_corr_topp1000 <- TCGAgene_corr_topp1000[1:1000, ]

# try bottom 1000 correlated genes 

TCGAgene_corr_bottom1000 <- TCGAgene_corr_p[order(TCGAgene_corr_p$TCGAgenecorrs, decreasing = FALSE), ]
TCGAgene_corr_bottom1000 <- TCGAgene_corr_bottom1000[1:1000, ]

# do the mito genes pop out?
mtOXPHOS %in% row.names(TCGAgene_corr_topp1000)
mtOXPHOS %in% row.names(TCGAgene_corr_bottom1000)

nuOXPHOS %in% row.names(TCGAgene_corr_topp1000)
nuOXPHOS %in% row.names(TCGAgene_corr_bottom1000)

# remove the mitochondrial genes

TCGAgene_corr_topp1000 <- TCGAgene_corr_topp1000[!(row.names(TCGAgene_corr_topp1000) %in% c(mtOXPHOS, nuOXPHOS)), ]
TCGAgene_corr_tbottom1000 <- TCGAgene_corr_bottom1000[!(row.names(TCGAgene_corr_bottom1000) %in% c(mtOXPHOS, nuOXPHOS)), ]

TCGAgenesymbolspos1000 <- getBM(mart = ensembl,
                            attributes = c("hgnc_symbol", "ensembl_gene_id"),
                            filters = "ensembl_gene_id",
                            values = row.names(TCGAgene_corr_topp1000))

TCGAgenesymbolsneg1000 <- getBM(mart = ensembl,
                                attributes = c("hgnc_symbol", "ensembl_gene_id"),
                                filters = "ensembl_gene_id",
                                values = row.names(TCGAgene_corr_bottom1000))

TCGApos1000enriched <- enrichr(unlist(TCGAgenesymbolspos1000$hgnc_symbol), dbs)
TCGAneg1000enriched <- enrichr(unlist(TCGAgenesymbolsneg1000$hgnc_symbol), dbs)

write.xlsx(list("FDR0.05_KEGG_2021_Human" = TCGApos1000enriched$KEGG_2021_Human, 
                "FDR0.05_GO_Molecular_Function_2018" = TCGApos1000enriched$GO_Molecular_Function_2018,
                "FDR0.05_GO_Cellular_Component_2018" = TCGApos1000enriched$GO_Cellular_Component_2018,
                "FDR0.05_GO_Biological_Process_2018" = TCGApos1000enriched$GO_Biological_Process_2018),
           file = "output/TCGA_genewise_expression_corr_enrichment_top1000.xlsx")

write.xlsx(list("FDR0.05_KEGG_2021_Human" = TCGAneg1000enriched$KEGG_2021_Human, 
                "FDR0.05_GO_Molecular_Function_2018" = TCGAneg1000enriched$GO_Molecular_Function_2018,
                "FDR0.05_GO_Cellular_Component_2018" = TCGAneg1000enriched$GO_Cellular_Component_2018,
                "FDR0.05_GO_Biological_Process_2018" = TCGAneg1000enriched$GO_Biological_Process_2018),
           file = "output/TCGA_genewise_expression_corr_enrichment_bottom1000.xlsx")

#### PROLIFERATIVE INDEX FOR GTEX ####
# corresponds to Fig 5b

# DESeq2 wants a colData object. Not actually used for the normalisation. Here we can use the sample IDs with tissue.
col_tissue <- LUT[colnames(countsdata), "Tissue"]
col_tissue <- as.matrix(col_tissue)
  
rownames(col_tissue) <- colnames(countsdata)
colnames(col_tissue) <- c("Tissue")
  
# need to create a DESeq2 object. Design set to ~1 allows for use of estimateSizeFactors
tempdds <- DESeqDataSetFromMatrix(countData = countsdata, colData = col_tissue, design = ~ 1)
  
tempvsd <- varianceStabilizingTransformation(tempdds, blind = TRUE)

# retrieve ensembl gene id and HGNC gene symbols for genes on mitochondrial DNA from biomart
replace_gene_symbols <- getBM(mart = ensembl,
                            attributes = c("hgnc_symbol", "ensembl_gene_id"),
                            filters = "ensembl_gene_id",
                            values = row.names(tempvsd))

replace_gene_symbols <- replace_gene_symbols[!(duplicated(replace_gene_symbols$ensembl_gene_id)), ]

row.names(replace_gene_symbols) <- replace_gene_symbols$ensembl_gene_id

MOR_vst_forPI <- data.frame(assay(tempvsd[row.names(tempvsd) %in% replace_gene_symbols$ensembl_gene_id, ]))
names <- replace_gene_symbols[row.names(MOR_vst_forPI), "hgnc_symbol"]
names[is.na(names)] <- paste("na", 1:sum(is.na(names)))
names[names == ""] <- paste(1:sum(names == ""))
names[duplicated(names)] <- paste0(names[duplicated(names)], 1:length(names[duplicated(names)]))
row.names(MOR_vst_forPI) <- names
  
GTEX_PIobject <- readDataForPI(MOR_vst_forPI, modelIDs = c("SCYL3"))
GTEX_PIs <- calculatePI(GTEX_PIobject)

OXPHOScorrforbind <- MOR_tissue_correlations$median_mtnuOXPHOS_spearman
names(OXPHOScorrforbind) <- MOR_tissue_correlations$tissue

tissuePIS_long <- data.frame(matrix(ncol = 2, nrow = 0))
colnames(tissuePIS_long) <- c("PI", "tissue")

for(i in 1:length(tissues)){

tempsampIDs <- LUT[LUT$Tissue == tissues[i], "SAMPID"]
  
tissuelabelvec <- vector(length = length(tempsampIDs))
tissuelabelvec[] <- tissues[i]

temp_df <- cbind(GTEX_PIs[tempsampIDs], tissuelabelvec)
colnames(temp_df) <- c("PI", "tissue")

tissuePIS_long <- rbind(tissuePIS_long, temp_df)

}

tissuePIS_long[, "OXPHOScorr"] <- OXPHOScorrforbind[paste(tissuePIS_long$tissue)]

saveRDS(tissuePIS_long, "output/tissuePIS_long.rds")

linefit <- lm(as.numeric(tissuePIS_long$PI) ~ tissuePIS_long$OXPHOScorr )
sum_linefit <- summary(linefit)

pdf("graphics/GTEXPIvolinplot.pdf", width = 8, height = 3)

ggplot(data = tissuePIS_long, aes(x = OXPHOScorr, y = as.numeric(PI))) +
  geom_violin(aes(group = tissue, fill = tissue), show.legend = FALSE, scale = 1, width = 0.015, position = "identity", lwd = 0.2) + 
  ylab("Proliferative Index") +
  xlab("mtOXPHOS-nuOXPHOS correlation") +
  theme_classic() +
  theme(axis.text.x = element_text(angle = 60,
                                   vjust = 1,
                                   hjust=1,
                                   family = "sans",
                                   color = "black",
                                   size = 5),
        axis.line = element_line(size = 0.25),
        axis.ticks = element_line(size = 0.25),
        legend.position = "none")+
  coord_cartesian(xlim = c(-0.4, 0.4)) +
  geom_abline(color='red', slope = sum_linefit$coefficients[2,1], intercept = sum_linefit$coefficients[1,1])+
  geom_vline(xintercept = 0, linetype = "dashed", color = "gray") + 
  geom_text(x = -0.2, y = 12, label = deparse(bquote(R^2 == .(round(sum_linefit$r.squared, 2)))), parse = TRUE, size = 2.3)+
  geom_text(x = -0.2, y = 11.4, label = deparse(bquote('p < 2 x' ~10^-16)), parse = TRUE, size = 2.3)

dev.off()

#### NFKB EXPRESSION FOR GTEx ####
# corresponding to Fig 5a, Table S4

NFKB1expression <- MOR_across_tissues["ENSG00000109320", ]

OXPHOScorr_forbind <- vector()
tissuevec_forbind <- vector()

for(i in 1:length(NFKB1expression)){

  temptish <- LUT[names(NFKB1expression)[i], "Tissue"]
  tissuevec_forbind[i] <- temptish
  
  if(temptish %in% MOR_tissue_correlations$tissue){
  
OXPHOScorr_forbind[i] <- MOR_tissue_correlations[MOR_tissue_correlations$tissue == temptish, "median_mtnuOXPHOS_spearman"]

}
}

tissue_long_NFKB1 <- cbind(NFKB1expression, OXPHOScorr_forbind, tissuevec_forbind)
tissue_long_NFKB1 <- data.frame(tissue_long_NFKB1)
colnames(tissue_long_NFKB1) <- c("NFKB1expression", "OXPHOScorr", "Tissue")

tissue_long_NFKB1 <- tissue_long_NFKB1[tissue_long_NFKB1$Tissue %in% MOR_tissue_correlations$tissue, ]

NFKB1linefit <- lm(log10(as.numeric(tissue_long_NFKB1$NFKB1expression)) ~ as.numeric(tissue_long_NFKB1$OXPHOScorr))
sum_NFKB1linefit <- summary(NFKB1linefit)

pdf("graphics/NFKB1violinplot.pdf", width = 8, height = 3)

ggplot(data = tissue_long_NFKB1,
       aes(x = as.numeric(OXPHOScorr),
           y = as.numeric(NFKB1expression))) +
  geom_violin(aes(group = Tissue,
                  fill = Tissue),
              show.legend = FALSE,
              scale = 1,
              width = 0.015,
              position = "identity",
              lwd = 0.2) + 
  ylab("NFKB1 expression") +
  xlab("mtOXPHOS-nuOXPHOS correlation") +
  theme_classic()+
  theme(axis.text.x = element_text(angle = 60,
                                   vjust = 1,
                                   hjust=1,
                                   family = "sans",
                                   color = "black",
                                   size = 5),
        axis.line = element_line(size = 0.25),
        axis.ticks = element_line(size = 0.25),
        legend.position = "none") +
  coord_cartesian(xlim = c(-0.4, 0.4)) +
  geom_text(x = -0.12,
            y = 4.25,
            label = deparse(bquote(R^2 == .(round(sum_NFKB1linefit$r.squared, 2)))),
            parse = TRUE,
            size = 2.3) +
  geom_text(x = -0.12,
            y = 4.05,
            label = deparse(bquote('p < 2 x' ~10^-16)),
            parse = TRUE,
            size = 2.3) +
  geom_abline(color='red',
              intercept = sum_NFKB1linefit$coefficients[1,1],
              slope = sum_NFKB1linefit$coefficients[2,1])+ 
  geom_vline(xintercept = 0,
             linetype = "dashed",
             color = "gray") +
  scale_y_continuous(trans = "log10")

dev.off()

# NFKB2
NFKB2expression <- MOR_across_tissues["ENSG00000077150", ]

OXPHOScorr_forbind <- vector()
tissuevec_forbind <- vector()

for(i in 1:length(NFKB2expression)){
  
  temptish <- LUT[names(NFKB2expression)[i], "Tissue"]
  tissuevec_forbind[i] <- temptish
  
  if(temptish %in% MOR_tissue_correlations$tissue){
    
    OXPHOScorr_forbind[i] <- MOR_tissue_correlations[MOR_tissue_correlations$tissue == temptish, "median_mtnuOXPHOS_spearman"]
    
  }
}

tissue_long_NFKB2 <- cbind(NFKB2expression, OXPHOScorr_forbind, tissuevec_forbind)
tissue_long_NFKB2 <- data.frame(tissue_long_NFKB2)
colnames(tissue_long_NFKB2) <- c("NFKB2expression", "OXPHOScorr", "Tissue")

tissue_long_NFKB2 <- tissue_long_NFKB2[tissue_long_NFKB2$Tissue %in% MOR_tissue_correlations$tissue, ]

NFKB2linefit <- lm(log10(as.numeric(tissue_long_NFKB2$NFKB2expression)) ~ as.numeric(tissue_long_NFKB2$OXPHOScorr))
sum_NFKB2linefit <- summary(NFKB2linefit)

pdf("graphics/NFKB2violinplot.pdf", width = 8, height = 4)

ggplot(data = tissue_long_NFKB2,
       aes(x = as.numeric(OXPHOScorr),
           y = as.numeric(NFKB2expression))) +
  geom_violin(aes(group = Tissue,
                  fill = Tissue),
              show.legend = FALSE,
              scale = 1,
              width = 0.015,
              position = "identity",
              lwd = 0.2) + 
  ylab("NFKB2 expression") +
  xlab("mtOXPHOS-nuOXPHOS correlation") +
  theme_classic() +
  theme(axis.text.x = element_text(angle = 60,
                                   vjust = 1,
                                   hjust=1,
                                   family = "sans",
                                   color = "black",
                                   size = 5),
        axis.line = element_line(size = 0.25),
        axis.ticks = element_line(size = 0.25),
        legend.position = "none") +
  coord_cartesian(xlim = c(-0.4, 0.4)) +
  geom_text(x = -0.12,
            y = 4.25,
            label = deparse(bquote(R^2 == .(round(sum_NFKB2linefit$r.squared, 2)))),
            parse = TRUE,
            size = 2.3)+
  geom_text(x = -0.12,
            y = 4.05,
            label = deparse(bquote('p < 2 x' ~10^-16)),
            parse = TRUE, size = 2.3) +
  geom_abline(color='red',
              intercept = sum_NFKB2linefit$coefficients[1,1],
              slope = sum_NFKB2linefit$coefficients[2,1]) +
  scale_y_continuous(trans = "log10")

dev.off()

#RELA   

RELAexpression <- MOR_across_tissues["ENSG00000173039", ]

OXPHOScorr_forbind <- vector()
tissuevec_forbind <- vector()

for(i in 1:length(RELAexpression)){
  
  temptish <- LUT[names(RELAexpression)[i], "Tissue"]
  tissuevec_forbind[i] <- temptish
  
  if(temptish %in% MOR_tissue_correlations$tissue){
    
    OXPHOScorr_forbind[i] <- MOR_tissue_correlations[MOR_tissue_correlations$tissue == temptish, "median_mtnuOXPHOS_spearman"]
    
  }
}

tissue_long_RELA <- cbind(RELAexpression, OXPHOScorr_forbind, tissuevec_forbind)
tissue_long_RELA <- data.frame(tissue_long_RELA)
colnames(tissue_long_RELA) <- c("RELAexpression", "OXPHOScorr", "Tissue")

tissue_long_RELA <- tissue_long_RELA[tissue_long_RELA$Tissue %in% MOR_tissue_correlations$tissue, ]

RELAlinefit <- lm(log10(as.numeric(tissue_long_RELA$RELAexpression)) ~ as.numeric(tissue_long_RELA$OXPHOScorr))
sum_RELAlinefit <- summary(RELAlinefit)

pdf("graphics/RELAviolinplot.pdf", width = 8, height = 4)

ggplot(data = tissue_long_RELA,
       aes(x = as.numeric(OXPHOScorr),
           y = as.numeric(RELAexpression))) +
  geom_violin(aes(group = Tissue,
                  fill = Tissue), 
              show.legend = FALSE, 
              scale = 1, 
              width = 0.015, 
              position = "identity", 
              lwd = 0.2) + 
  ylab("RELA expression") +
  xlab("mtOXPHOS-nuOXPHOS correlation") +
  theme_classic()+
  theme(axis.text.x = element_text(angle = 60,
                                   vjust = 1,
                                   hjust=1,
                                   family = "sans",
                                   color = "black",
                                   size = 5),
        axis.line = element_line(size = 0.25),
        axis.ticks = element_line(size = 0.25),
        legend.position = "none") +
  coord_cartesian(xlim = c(-0.4, 0.4)) +
  geom_text(x = -0.12,
            y = 4.25,
            label = deparse(bquote(R^2 == .(round(sum_RELAlinefit$r.squared, 2)))),
            parse = TRUE,
            size = 2.3) +
  geom_text(x = -0.12,
            y = 4.05,
            label = deparse(bquote('p < 2 x' ~10^-16)),
            parse = TRUE,
            size = 2.3) +
  geom_abline(color='red',
              intercept = sum_RELAlinefit$coefficients[1,1],
              slope = sum_RELAlinefit$coefficients[2,1]) +
  scale_y_continuous(trans = "log10")

dev.off()

# RELB  

RELBexpression <- MOR_across_tissues["ENSG00000104856", ]

OXPHOScorr_forbind <- vector()
tissuevec_forbind <- vector()

for(i in 1:length(RELBexpression)){
  
  temptish <- LUT[names(RELBexpression)[i], "Tissue"]
  tissuevec_forbind[i] <- temptish
  
  if(temptish %in% MOR_tissue_correlations$tissue){
    
    OXPHOScorr_forbind[i] <- MOR_tissue_correlations[MOR_tissue_correlations$tissue == temptish, "median_mtnuOXPHOS_spearman"]
    
  }
}

tissue_long_RELB <- cbind(RELBexpression, OXPHOScorr_forbind, tissuevec_forbind)
tissue_long_RELB <- data.frame(tissue_long_RELB)
colnames(tissue_long_RELB) <- c("RELBexpression", "OXPHOScorr", "Tissue")

tissue_long_RELB <- tissue_long_RELB[tissue_long_RELB$Tissue %in% MOR_tissue_correlations$tissue, ]

RELBlinefit <- lm(log10(as.numeric(tissue_long_RELB$RELBexpression)) ~ as.numeric(tissue_long_RELB$OXPHOScorr))
sum_RELBlinefit <- summary(RELBlinefit)

pdf("graphics/RELBviolinplot.pdf", width = 8, height = 4)

ggplot(data = tissue_long_RELB, 
       aes(x = as.numeric(OXPHOScorr),
           y = as.numeric(RELBexpression))) +
  geom_violin(aes(group = Tissue, 
                  fill = Tissue),
              show.legend = FALSE,
              scale = 1,
              width = 0.015,
              position = "identity",
              lwd = 0.2) + 
  ylab("RELB expression") +
  xlab("mtOXPHOS-nuOXPHOS correlation") +
  theme_classic()+
  theme(axis.text.x = element_text(angle = 60,
                                   vjust = 1,
                                   hjust=1,
                                   family = "sans",
                                   color = "black",
                                   size = 5),
        axis.line = element_line(size = 0.25),
        axis.ticks = element_line(size = 0.25),
        legend.position = "none") +
  coord_cartesian(xlim = c(-0.4, 0.4)) +
  geom_text(x = -0.12,
            y = 4.25,
            label = deparse(bquote(R^2 == .(round(sum_RELBlinefit$r.squared, 2)))),
            parse = TRUE,
            size = 2.3) +
  geom_text(x = -0.12,
            y = 4.05,
            label = deparse(bquote('p < 2 x' ~10^-16)),
            parse = TRUE,
            size = 2.3) +
  geom_abline(color='red',
              intercept = sum_RELBlinefit$coefficients[1,1],
              slope = sum_RELBlinefit$coefficients[2,1]) +
  scale_y_continuous(trans = "log10")

dev.off()

#REL  

RELexpression <- MOR_across_tissues["ENSG00000162924", ]

OXPHOScorr_forbind <- vector()
tissuevec_forbind <- vector()

for(i in 1:length(RELexpression)){
  
  temptish <- LUT[names(RELexpression)[i], "Tissue"]
  tissuevec_forbind[i] <- temptish
  
  if(temptish %in% MOR_tissue_correlations$tissue){
    
    OXPHOScorr_forbind[i] <- MOR_tissue_correlations[MOR_tissue_correlations$tissue == temptish, "median_mtnuOXPHOS_spearman"]
    
  }
}

tissue_long_REL <- cbind(RELexpression, OXPHOScorr_forbind, tissuevec_forbind)
tissue_long_REL <- data.frame(tissue_long_REL)
colnames(tissue_long_REL) <- c("RELexpression", "OXPHOScorr", "Tissue")

tissue_long_REL <- tissue_long_REL[tissue_long_REL$Tissue %in% MOR_tissue_correlations$tissue, ]

RELlinefit <- lm(log10(as.numeric(tissue_long_REL$RELexpression)) ~ as.numeric(tissue_long_REL$OXPHOScorr))
sum_RELlinefit <- summary(RELlinefit)

pdf("graphics/RELviolinplot.pdf", width = 8, height = 4)

ggplot(data = tissue_long_REL,
       aes(x = as.numeric(OXPHOScorr),
           y = as.numeric(RELexpression))) +
  geom_violin(aes(group = Tissue,
                  fill = Tissue),
              show.legend = FALSE,
              scale = 1,
              width = 0.015,
              position = "identity",
              lwd = 0.2) + 
  ylab("REL expression") +
  xlab("mtOXPHOS-nuOXPHOS correlation") +
  theme_classic()+
  theme(axis.text.x = element_text(angle = 60,
                                   vjust = 1,
                                   hjust=1,
                                   family = "sans",
                                   color = "black",
                                   size = 5),
        axis.line = element_line(size = 0.25),
        axis.ticks = element_line(size = 0.25),
        legend.position = "none") +
  coord_cartesian(xlim = c(-0.4, 0.4)) +
  geom_text(x = -0.12, y = 4.05,
            label = deparse(bquote(R^2 == .(round(sum_RELlinefit$r.squared, 2)))),
            parse = TRUE,
            size = 2.3)+
  geom_text(x = -0.12,
            y = 3.85,
            label = deparse(bquote('p < 2 x' ~10^-16)),
            parse = TRUE,
            size = 2.3) +
  geom_abline(color='red',
              intercept = sum_RELlinefit$coefficients[1,1],
              slope = sum_RELlinefit$coefficients[2,1]) +
  scale_y_continuous(trans = "log10")

dev.off()

tissuelong_NFKBmembers <- cbind(as.numeric(tissue_long_NFKB1$NFKB1expression),
                                as.numeric(tissue_long_NFKB1$OXPHOScorr),
                                as.numeric(tissue_long_NFKB2$NFKB2expression),
                                as.numeric(tissue_long_REL$RELexpression),
                                as.numeric(tissue_long_RELA$RELAexpression),
                                as.numeric(tissue_long_RELB$RELBexpression))

tissuelong_NFKBmembers <- data.frame(tissuelong_NFKBmembers)
tissuelong_NFKBmembers[, "Tissue"] <- tissue_long_NFKB1$Tissue
tissuelong_NFKBmembers[, "PI"] <- as.numeric(tissuePIS_long[row.names(tissue_long_NFKB1), "PI"])

colnames(tissuelong_NFKBmembers) <- c("NFKB1",
                                      "OXPHOScorr",
                                      "NFKB2", 
                                      "REL",
                                      "RELA",
                                      "RELB",
                                      "Tissue",
                                      "PI")

GTEX_NFKB_PI_lmsum <- summary(lm(formula = OXPHOScorr ~ 
                                   log10(NFKB1) + log10(NFKB2) + log10(REL) + log10(RELA) + log10(RELB) + as.numeric(PI), 
     data = tissuelong_NFKBmembers))

summary(lm(formula = OXPHOScorr ~ 
             log10(NFKB1),
           data = tissuelong_NFKBmembers))

sink("output/GTEX_NFKB_PI_lmsum.txt")
print(GTEX_NFKB_PI_lmsum)
sink()

#### TCGA PROLIFERATIVE INDEX ####
# corresponding to Fig 5d, Table S4

# TCGA_counts_list <- readRDS("output/TCGA_counts_list.rds")
# TCGA_LUT <- readRDS("output/TCGA_LUT.rds")
# TCGA_MOR_mtOXPHOS_nuOXPHOS <- readRDS("output/TCGA_MOR_mtOXPHOS_nuOXPHOS_correlations.rds")

TCGA_counts_df <- do.call(cbind, TCGA_counts_list)

TCGA_counts_df <- TCGA_counts_df[, colnames(TCGA_counts_df) %in% TCGA_LUT$SAMPID]
col_data <- TCGA_LUT[colnames(TCGA_counts_df), "cancer"]
names(col_data) <- TCGA_LUT[colnames(TCGA_counts_df), "SAMPID"]
col_data <- data.frame(col_data)

# need to create a DESeq2 object. Design set to ~1 allows for use of estimateSizeFactors
TCGAdds <- DESeqDataSetFromMatrix(countData = TCGA_counts_df, colData = col_data, design = ~ 1)

TCGAvsd <- varianceStabilizingTransformation(TCGAdds, blind = TRUE)

TCGA_vst_df <- data.frame(assay(TCGAvsd))

# retrieve ensembl gene id and HGNC gene symbols for genes on mitochondrial DNA from biomart
TCGA_replace_gene_symbols <- getBM(mart = ensembl,
                                   attributes = c("hgnc_symbol", "ensembl_gene_id"),
                                   filters = "ensembl_gene_id",
                                   values = row.names(TCGA_vst_df))

TCGA_replace_gene_symbols <- TCGA_replace_gene_symbols[!(duplicated(TCGA_replace_gene_symbols$ensembl_gene_id)), ]
row.names(TCGA_replace_gene_symbols) <- TCGA_replace_gene_symbols$ensembl_gene_id

TCGA_vst_df <- data.frame(assay(TCGAvsd[row.names(TCGAvsd) %in% TCGA_replace_gene_symbols$ensembl_gene_id, ]))
names <- TCGA_replace_gene_symbols[row.names(TCGA_vst_df), "hgnc_symbol"]
names[is.na(names)] <- paste("na.", 1:sum(is.na(names)))
names[names == ""] <- paste(1:sum(names == ""))
names[duplicated(names)] <- paste0(names[duplicated(names)], 1:length(names[duplicated(names)]))
row.names(TCGA_vst_df) <- names

TCGA_PIobject <- readDataForPI(TCGA_vst_df, modelIDs = c("SCYL3"))
TCGA_PI <- calculatePI(TCGA_PIobject)

TCGAforbind <- TCGA_MOR_mtOXPHOS_nuOXPHOS$cancer_mean_corr_df$median_mtnuOXPHOS_spearman
names(TCGAforbind) <- TCGA_MOR_mtOXPHOS_nuOXPHOS$cancer_mean_corr_df$cancertype

TCGAcancerPIS_long <- data.frame(matrix(ncol = 2, nrow = 0))
colnames(TCGAcancerPIS_long) <- c("PI", "cancer")

for(i in 1:length(TCGA_MOR_mtOXPHOS_nuOXPHOS$cancer_mean_corr_df$cancertype)){
  
  tempsampIDs <- TCGA_LUT[TCGA_LUT$cancer == TCGA_MOR_mtOXPHOS_nuOXPHOS$cancer_mean_corr_df$cancertype[i], "SAMPID"]
  presentsampIDs <- tempsampIDs[tempsampIDs %in% str_replace_all(names(TCGA_PI), pattern = "\\.", replacement = "-")]
  
  cancerlabelvec <- vector(length = length(presentsampIDs))
  cancerlabelvec[] <- TCGA_MOR_mtOXPHOS_nuOXPHOS$cancer_mean_corr_df$cancertype[i]
  
  temp_df <- cbind(TCGA_PI[str_replace_all(presentsampIDs, pattern = "-", replacement = "\\.")], cancerlabelvec)
  colnames(temp_df) <- c("PI", "cancer")
  
  TCGAcancerPIS_long <- rbind(TCGAcancerPIS_long, temp_df)
  
}

TCGAcancerPIS_long[, "OXPHOScorr"] <- TCGAforbind[paste(TCGAcancerPIS_long$cancer)]

saveRDS(TCGAcancerPIS_long, "output/TCGAcancerPIS_long.rds")

TCGAlinefit <- lm(as.numeric(TCGAcancerPIS_long$PI) ~ TCGAcancerPIS_long$OXPHOScorr)
TCGAsum_linefit <- summary(TCGAlinefit)

pdf("graphics/TCGA_OXPHOScorr_vs_PI_volinplot.pdf", width = 8, height = 3)

ggplot(data = TCGAcancerPIS_long,
       aes(x = OXPHOScorr,
           y = as.numeric(PI))) +
  geom_violin(aes(group = cancer,
                  fill = cancer),
              show.legend = FALSE,
              scale = 1,
              width = 0.015,
              position = "identity",
              lwd = 0.2) + 
  ylab("Proliferative Index") +
  xlab("mtOXPHOS-nuOXPHOS correlation") +
  theme_classic() +
  theme(axis.text.x = element_text(angle = 60,
                                   vjust = 1,
                                   hjust=1,
                                   family = "sans",
                                   color = "black",
                                   size = 5),
        axis.line = element_line(size = 0.25),
        axis.ticks = element_line(size = 0.25),
        legend.position = "none") +
  coord_cartesian(xlim = c(-0.05, 0.4)) +
  geom_vline(xintercept = 0,
             linetype = "dashed",
             color = "gray") +
  geom_abline(color='red',
              intercept = TCGAsum_linefit$coefficients[1,1],
              slope = TCGAsum_linefit$coefficients[2,1]) +
  geom_text(x = 0.25,
            y = 12,
            label = deparse(bquote(R^2 == .(round(TCGAsum_linefit$r.squared, 2)))),
            parse = TRUE,
            size = 2.3)+
  geom_text(x = 0.25,
            y = 11.6,
            label = deparse(bquote('p < 2.2 x' ~10^-16)),
            parse = TRUE,
            size = 2.3)

dev.off()

#### NFKB EXPRESSION FOR TCGA ####
# corresponds to Fig 5c

# TCGA_MORnorm_acrosscancers <- readRDS("output/TCGA-MOR-normalisation-across-cancers.rds")

# TCGA_MOR_list <- readRDS("output/TCGA_MOR_list.rds")
# TCGA_counts_list <- readRDS("output/TCGA_counts_list.rds")

TCGA_MOR_mtOXPHOS_nuOXPHOS <- readRDS("output/TCGA_MOR_mtOXPHOS_nuOXPHOS_correlations.rds")
row.names(TCGA_MOR_mtOXPHOS_nuOXPHOS$cancer_mean_corr_df) <- TCGA_MOR_mtOXPHOS_nuOXPHOS$cancer_mean_corr_df$cancertype

# TCGA_LUT <- readRDS("output/TCGA_LUT.rds")

TCGA_NFKB1expression <- TCGA_MORnorm_acrosscancers["ENSG00000109320", ]
TCGA_NFKB1expression <- TCGA_NFKB1expression[names(TCGA_NFKB1expression) %in% TCGA_LUT$SAMPID]

TCGA_NFKB2expression <- TCGA_MORnorm_acrosscancers["ENSG00000077150", ]
TCGA_NFKB2expression <- TCGA_NFKB2expression[names(TCGA_NFKB2expression) %in% TCGA_LUT$SAMPID]

TCGA_RELexpression <- TCGA_MORnorm_acrosscancers["ENSG00000162924", ]
TCGA_RELexpression <- TCGA_RELexpression[names(TCGA_RELexpression) %in% TCGA_LUT$SAMPID]

TCGA_RELAexpression <- TCGA_MORnorm_acrosscancers["ENSG00000173039", ]
TCGA_RELAexpression <- TCGA_RELAexpression[names(TCGA_RELAexpression) %in% TCGA_LUT$SAMPID]

TCGA_RELBexpression <- TCGA_MORnorm_acrosscancers["ENSG00000104856", ]
TCGA_RELBexpression <- TCGA_RELBexpression[names(TCGA_RELBexpression) %in% TCGA_LUT$SAMPID]

TCGA_OXPHOS_NFKBmembers <- cbind(TCGA_LUT[names(TCGA_NFKB1expression), "cancer"],
                                 TCGA_NFKB1expression,
                                 TCGA_NFKB2expression,
                                 TCGA_RELexpression,
                                 TCGA_RELAexpression,
                                 TCGA_RELBexpression,
                                 TCGAcancerPIS_long[str_replace_all(names(TCGA_NFKB1expression), pattern = "-", replacement = "."), "PI"])

TCGA_OXPHOS_NFKBmembers <- data.frame(TCGA_OXPHOS_NFKBmembers)

colnames(TCGA_OXPHOS_NFKBmembers) <- c("cancertype",
                                       "NFKB1expression",
                                       "NFKB2expression",
                                       "RELexpression",
                                       "RELAexpression",
                                       "RELBexpression",
                                       "PI")

TCGA_OXPHOS_NFKBmembers <- TCGA_OXPHOS_NFKBmembers[TCGA_OXPHOS_NFKBmembers$cancertype %in% unique(TCGA_MOR_mtOXPHOS_nuOXPHOS$cancer_mean_corr_df$cancertype), ]
TCGA_OXPHOS_NFKBmembers[, "OXPHOScorr"] <- TCGA_MOR_mtOXPHOS_nuOXPHOS$cancer_mean_corr_df[paste(TCGA_OXPHOS_NFKBmembers$cancertype), "median_mtnuOXPHOS_spearman"]

TCGA_NFKB1_lm <- lm(log10(as.numeric(TCGA_OXPHOS_NFKB_long$NFKB1expression)) ~ as.numeric(TCGA_OXPHOS_NFKB_long$OXPHOScorr))
sum_TCGA_NFKB1 <- summary(TCGA_NFKB1_lm)

TCGA_NFKB_PI_lmsum <- summary(lm(formula = OXPHOScorr ~ 
                                   log10(as.numeric(NFKB1expression)) + log10(as.numeric(NFKB2expression)) + log10(as.numeric(RELexpression)) + log10(as.numeric(RELAexpression)) + log10(as.numeric(RELBexpression)) + as.numeric(PI), 
                                 data = TCGA_OXPHOS_NFKBmembers))

sink("output/TCGA_NFKB_PI_lmsum.txt")
print(TCGA_NFKB_PI_lmsum)
sink()

pdf("graphics/TCGA_NFKB1volinplot.pdf", width = 8, height = 3)

ggplot(data = TCGA_OXPHOS_NFKBmembers,
       aes(x = OXPHOScorr,
           y = as.numeric(NFKB1expression))) +
  geom_violin(aes(group = cancertype,
                  fill = cancertype),
              show.legend = FALSE,
              scale = 1,
              width = 0.015,
              position = "identity",
              lwd = 0.2) + 
  ylab("NFKB1 expression") +
  xlab("mtOXPHOS-nuOXPHOS correlation") +
  theme_classic() +
  theme(axis.text.x = element_text(angle = 60,
                                   vjust = 1,
                                   hjust=1,
                                   family = "sans",
                                   color = "black",
                                   size = 5),
        axis.line = element_line(size = 0.25),
        axis.ticks = element_line(size = 0.25),
        legend.position = "none")+
  coord_cartesian(xlim = c(-0.1, 0.4)) +
  geom_abline(color = "red",
              intercept = sum_TCGA_NFKB1$coefficients[1,1],
              slope = sum_TCGA_NFKB1$coefficients[2,1])+
  geom_vline(xintercept = 0, linetype = "dashed",
             color = "gray") +
  geom_text(x = -0.05,
            y = 4.3,
            label = deparse(bquote(R^2 == .(round(sum_TCGA_NFKB1$r.squared, 2)))),
            parse = TRUE,
            size = 2.3)+
  geom_text(x = -0.05,
            y = 4.1,
            label = deparse(bquote('p < 2 x' ~10^-16)),
            parse = TRUE,
            size = 2.3) +
  scale_y_continuous(trans = "log10")

dev.off()

#### IMMUNE CELL PROPORTION ####
# corresponds to Fig S6

# this file obtained from [https://github.com/BNadel/GEDIT/blob/master/Application/GTEX/2_Predictions_GTEXNames/CombinedPredictions.tsv]
GEDITpredictions <- read.table("input/CombinedPredictions.txt", sep = ",", header = TRUE)

TotalImmune_long <- matrix(ncol = 4, nrow = 0)
colnames(TotalImmune_long) <- c("Tissue", "TotalImmune", "OXPHOScorr", "NFKB1expression")

for(i in 1:length(tissues)){

  sampIDS <- LUT[LUT$Tissue == tissues[i], "SAMPID"]
  tempdata <- GEDITpredictions[GEDITpredictions$Sample %in% sampIDS, "TotalImmune"]
  names(tempdata) <- GEDITpredictions[GEDITpredictions$Sample %in% sampIDS, "Sample"]
  
  tissuelabelvec <- vector(length = length(sampIDS))
  tissuelabelvec[] <- tissues[i]
  
  NFKB1_vec <- NFKB1expression[sampIDS]

  tissueOXPHOScorr <- vector(length = length(sampIDS))
  tissueOXPHOScorr[] <- MOR_tissue_correlations[MOR_tissue_correlations$tissue == tissues[i], "median_mtnuOXPHOS_spearman"]
  
  temp_df <- cbind(tissuelabelvec, tempdata, tissueOXPHOScorr, NFKB1_vec)
  
  colnames(temp_df) <- c("Tissue", "TotalImmune", "OXPHOScorr", "NFKB1expression")
  
  TotalImmune_long <- rbind(TotalImmune_long, temp_df)
  
}

TotalImmune_long <- data.frame(TotalImmune_long)

TotalImmune_linefit <- lm(log10(as.numeric(TotalImmune_long$TotalImmune)) ~ as.numeric(TotalImmune_long$OXPHOScorr))
sum_TotalImmune_linefit <- summary(TotalImmune_linefit)

pdf("graphics/TotalImmuneviolinplot.pdf", width = 8, height = 4)

ggplot(data = TotalImmune_long,
       aes(x = as.numeric(OXPHOScorr),
           y = as.numeric(TotalImmune))) +
  geom_violin(aes(group = Tissue,
                  fill = Tissue),
              show.legend = FALSE,
              scale = 1,
              width = 0.015,
              position = "identity",
              lwd = 0.2) + 
  ylab("Total immmune cell fraction") +
  xlab("mtOXPHOS-nuOXPHOS correlation") +
  theme_classic() +
  theme(axis.text.x = element_text(angle = 60,
                                   vjust = 1,
                                   hjust=1,
                                   family = "sans",
                                   color = "black",
                                   size = 5),
        axis.line = element_line(size = 0.25),
        axis.ticks = element_line(size = 0.25),
        legend.position = "none") +
  coord_cartesian(xlim = c(-0.4, 0.4)) +
  geom_text(x = -0.2,
            y = 6,
            label = deparse(bquote(R^2 == .(round(sum_TotalImmune_linefit$r.squared, 2)))),
            parse = TRUE,
            size = 2.3) +
  geom_text(x = -0.2,
            y = 5.5,
            label = deparse(bquote('p < 2 x' ~10^-16)),
            parse = TRUE,
            size = 2.3) +
  geom_abline(color = 'red',
              intercept = sum_TotalImmune_linefit $coefficients[1,1],
              slope = sum_TotalImmune_linefit $coefficients[2,1]) +
  scale_y_continuous(trans = "log10",
                     labels = function(x) format(x, scientific = FALSE))

dev.off()
```
